## Supplementary material for "Advancing Early Diagnosis of Alzheimer’s Disease: A Paper-Based Aptasensor for Detecting A*β*(1-42) and p-tau181 from Plasma Using CdTe Quantum Dots": Latex Source: cas-dc-template.tex

%% 
%% Copyright 2019-2021 Elsevier Ltd
%% 
%% This file is part of the 'CAS Bundle'.
%% --------------------------------------
%% 
%% It may be distributed under the conditions of the LaTeX Project Public
%% License, either version 1.2 of this license or (at your option) any
%% later version.  The latest version of this license is in
%%    http://www.latex-project.org/lppl.txt
%% and version 1.2 or later is part of all distributions of LaTeX
%% version 1999/12/01 or later.
%% 
%% The list of all files belonging to the 'CAS Bundle' is
%% given in the file `manifest.txt'.
%% 
%% Template article for cas-dc documentclass for 
%% double column output.

\documentclass[a4paper,fleqn]{cas-dc}


% If the frontmatter runs over more than one page
% use the longmktitle option.

%\documentclass[a4paper,fleqn,longmktitle]{cas-dc}

\usepackage{subfigure}
\usepackage{caption}
\usepackage{subcaption}
\usepackage{float}
\usepackage[numbers]{natbib}
%\usepackage[authoryear]{natbib}


%%%Author macros
\def\tsc#1{\csdef{#1}{\textsc{\lowercase{#1}}\xspace}}
\tsc{WGM}
\tsc{QE}
%%%

% Uncomment and use as if needed
%\newtheorem{theorem}{Theorem}
%\newtheorem{lemma}[theorem]{Lemma}
%\newdefinition{rmk}{Remark}
%\newproof{pf}{Proof}
%\newproof{pot}{Proof of Theorem \ref{thm}}

\begin{document}
\let\WriteBookmarks\relax
\def\floatpagepagefraction{1}
\def\textpagefraction{.001}

% Short title
\shorttitle{}    

% Short author
%\shortauthors{<short author list for running head>}  

% Main title of the paper
\title [mode = title]{Advancing Early Diagnosis of Alzheimer's Disease: A Paper-Based Aptasensor for Detecting A$\beta$(1-42) and p-tau181 from Plasma Using CdTe Quantum Dots}  

% Title footnote mark
% eg: \tnotemark[1]
%\tnotemark[<tnote number>] 

% Title footnote 1.
% eg: \tnotetext[1]{Title footnote text}
%\tnotetext[<tnote number>]{<tnote text>} 

% First author
%
% Options: Use if required
% eg: \author[1,3]{Author Name}[type=editor,
%       style=chinese,
%       auid=000,
%       bioid=1,
%       prefix=Sir,
%       orcid=0000-0000-0000-0000,
%       facebook=<facebook id>,
%       twitter=<twitter id>,
%       linkedin=<linkedin id>,
%       gplus=<gplus id>]

\author[1]{Ebrar Balci}[]
\ead{}
% Corresponding author indication
\cormark[1]
% Corresponding author text

% Footnote of the first author
% Email id of the first author
% \ead{}

\affiliation[1]{organization={Molecular Biology and Genetics Department, Istanbul Technical University},%Department and Organization
            addressline={ITU Ayazaga Campus}, 
            city={Istanbul},
            postcode={34469}, 
            %state={Istanbul},
            country={Turkey}}
\affiliation[2]{organization={Electronics and Communication Engineering Department, Istanbul Technical University},%Department and Organization
            city={Istanbul},
            postcode={34469}, 
            %state={Istanbul},
            country={Turkey}}

\affiliation[3]{organization={Physics Engineering Department, Istanbul Technical University},%Department and Organization
            addressline={ITU Ayazaga Campus}, 
            city={Istanbul},
            postcode={34469}, 
            %state={Istanbul},
            country={Turkey}}
            
\affiliation[4]{organization={Chemical Engineering Department, Istanbul Technical University},%Department and Organization
            addressline={ITU Ayazaga Campus}, 
            city={Istanbul},
            postcode={34469}, 
            %state={Istanbul},
            country={Turkey}}

% \affiliation[5]{organization={Chemistry Department, Istanbul Technical University},
%             addressline={ITU Ayazaga Campus}, 
%             city={Istanbul},
%             postcode={34469}, 
%             %state={Istanbul},
%             country={Turkey}}
            
%     \affiliation[6]{organization={Biomedical Engineering Department, Ankara Yildirim Beyazit University},
%             addressline={Etlik-Kecioren}, 
%             city={Ankara},
%             postcode={06220}, 
%             %state={Istanbul},
%             country={Turkey}}    
            
\author[1]{Elif Nur Yildiz}[]
\author[1]{Sevval Sueda Oksuz}[]
\author[2]{Nihat Ahmadli}[orcid = 0009-0005-2008-4533]
\author[3]{Miray Kargidan}[]
\author[1]{Nilay Ayyildiz}[]
\author[1]{Esra Alemdar}[]
\author[1]{Irem Uludag}[]
\author[4]{Umut Hasirci}[]


% For a title note without a number/mark
%\nonumnote{}

% Here goes the abstract
\begin{abstract}
Alzheimer's Disease (AD), characterized by a gradual onset and a lack of exact therapeutic interventions, underscores the imperative for the development of uncomplicated and cost-effective biosensors capable of detecting its biomarkers. This necessity arises in anticipation of a projected surge in the incidence of AD. Quantum dots (QDs) represent the promising new generation of luminophores owing to their size, composition, and surface-dependent tunable photoluminescence (PL) and photochemical stability. In this study, a paper-based QD aptasensor for the early detection of AD by targeting amyloid beta (A$\beta$-42) and p-tau181 proteins using Förster Resonance Energy Transfer (FRET) is developed. The sensor employs a Whatman paper with six sensing wells, integrating hydrophobic and hydrophilic regions, hydrophobic parts created through wax. Blood samples are placed in the inlet, dispersing into six sensing wells containing QD-aptamer-AuNP complexes. Target proteins induce conformational changes in aptamers, leading to fluorescence quenching in CdTe QDs. Two wells target p-tau181, two target amyloid beta-42, and two serve as references. Fluorescence emission spectra from each well are recorded, showing a linear correlation between fluorescence quenching and protein concentration. Values from each pair of wells are then averaged, and the average values from the pairs targeting (A$\beta$-42) and p-tau181 are compared to the average value of the reference wells. This paper-based aptasensor holds promise for early diagnosis of Alzheimer's Disease and opens the avenue of personalized medicine for the diagnosis of Alzheimer’s.
\end{abstract}

% Use if graphical abstract is present
%\begin{graphicalabstract}
%\includegraphics{}
%\end{graphicalabstract}

% Research highlights
% \begin{highlights}
% \item 
% \item 
% \item 
% \end{highlights}

% Keywords
% Each keyword is seperated by \sep
\begin{keywords}
Paper-based biosensor \sep Micropad \sep Quantum Dots \sep Amyloid beta 42 \sep Aptamer \sep p-tau181 \sep Alzheimer's Disease \sep Fluorescence \sep FRET
\end{keywords}
\maketitle

% Main text
\section{Introduction}\label{intro}
Alzheimer's Disease (AD) is a neurodegenerative disorder that primarily affects the brain, leading to progressive cognitive decline and may proceed to loss of capacity to converse and respond to the surroundings \cite{knopman2021alzheimer}. AD affects brain regions that govern cognition, memory, and language \cite{knopman2021alzheimer}. The urgency of AD is underscored by its increasing global prevalence. Namely, the number of AD cases, stemming from various causes, is projected to surge from 50 million in 2010 to an alarming 113 million by 2050 \cite{schneider2009neuropathology}.  Furthermore, in 2019, the anticipated worldwide economic burden of Alzheimer's Disease and Related Dementias (ADRDs) was projected at \$2.8 trillion. Projections indicate a substantial escalation, with anticipated costs reaching \$4.7 trillion in 2030, \$8.5 trillion in 2040, and a staggering \$16.9 trillion in 2050 \cite{nandi2022global}.

Neuropathological markers of AD mainly include the presence of neuritic (senile) plaques and neurofibrillary tangles, as well as neuronal loss, dystrophic neuritis, and inflammation. Neuritic plaques are extracellular lesions containing the amyloid beta 42 (A$\beta$-42) peptide \cite{sipe2005amyloid}. While the cause of neurofibrillary tangles, tau protein, contains 2-3 moles of phosphate per mole of tau protein in a healthy adult human brain. Brain tau is three to four times more hyperphosphorylated in AD than in healthy mature brain tau, which reduces neurosynaptic activities \cite{iqbal2010tau}. The degenerative cascade of Alzheimer's Disease lasts at least 10-15 years before clinical symptoms appear \cite{lu2022high}. The A$\beta$-42/A$\beta$-40 ratio allows for the identification of patients at the earlier and presymptomatic phase of illness \cite{findeis2007role}. While this ratio provides the earliest diagnosis, the tau level (in terms of pg/ml) in blood and cerebrospinal fluid  (CSF) provides the most specific detection \cite{zetterberg2010amyloid}. It is also found that p-tau181 and p-tau217 have the highest selectivity for diagnosis of the disease as blood biomarkers \cite{barthelemy2020blood}. The early detection of the disease promotes early intervention and treatment by providing individuals with access to medications and services that can help them manage symptoms. The development of a low-cost test capable of diagnosing AD at especially early stages is thus imperative.

In contemporary technology, neuroimaging and clinical techniques are used for the diagnosis of Alzheimer’s Disease.  While Magnetic Resonance Imaging (MRI) and Positron Emission Tomography (PET) play pivotal roles in assessing and monitoring the disease's progression,  MRI falls short in distinguishing AD from other neurodegenerative disorders associated with dementia, and PET is a high-cost process \cite{petrella2003neuroimaging}. Furthermore, these screening techniques prove inefficient in early-stage detection even though their time-consuming process. In contrast, blood biomarkers offer the advantage of detecting AD at its early stages by accumulating p-tau181 and A$\beta$(1-42) monomers \cite{valkova2021novel}. 

Biosensors emerge as a promising alternative to above traditional diagnostic methods, offering an effective and accessible solution for detection at all stages \cite{chepyala2020applications}. In the past decade, significant research has focused on biosensor development for Alzheimer's Disease, utilizing techniques such as ELISA, colorimetric assays, and electrochemical assays \cite{brazaca2020applications, kim2020clinically, shinohara2021elisa}. However, current biosensors face limitations in achieving early and precise AD diagnosis due to their high costs and complex instrument setups \cite{karki2021advances}. Aptamers, single-stranded DNA or RNA oligonucleotides adopting specific three-dimensional structures when interacting with specific targets, offer distinct advantages as biorecognition elements. These include easy chemical synthesis, high stability, reproducibility, and sensitivity, setting them apart from traditional antibodies \cite{kim2016aptamer, liu2022aptamers}.

\begin{figure*}[t]
	\centering
		\includegraphics[width=0.70\linewidth, height=0.70\linewidth]{Images/desn.png}
	  \caption{Graph of wavelength and intensity of the QD-Aptamer combination demonstrating the estimated change of intensity.}\label{fig: summary}
\end{figure*}

Fluorescence-based aptasensing assays have garnered considerable interest in monitoring biomarkers due to their exceptional features. The Förster Resonance Energy Transfer (FRET) process, which facilitates the transmission of energy from a "donor" to an "acceptor", stands out as an effective technique for designing fluorescence-based aptasensors, ensuring excellent sensitivity and selectivity in target detection \cite{khoshbin2020low, ghosh2020study}. Any changes in the environment donor that alter the distance between the two molecules also alter the fluorescence of the acceptor molecule since it is only active when both the donor and acceptor fluorophore molecules are in close proximity. This eliminates the need to explicitly label or alter the target biomolecule to detect its existence \cite{verma2023fret}. 

Quantum dots (QDs) are minute semiconductor particles or nanocrystals that exhibit unique optical properties and are characterized by diameters ranging from 2 to 10 nm, comprising 10 to 50 atoms \cite{day2012fluorescent}. Due to their small size and capacity to function within confined spaces, they exhibit the quantum confinement effect, resulting in distinct energy levels. QDs have been widely utilized in FRET systems due to their size-dependent photoluminescence characteristics, making them excellent donors \cite{singh2020quantum}. On the other hand, gold nanoparticles (AuNPs) have recently emerged as superior fluorescence acceptors in FRET systems, replacing traditional quenchers and offering new possibilities for sensitive detection of various substances due to their distinctive properties \cite{frigerio2012application}. The QDs-AuNPs FRET approach has found applications in assays for detecting several analytes, such as glycoproteins \cite{saha2012gold}, enzymes \cite{chang2017fluorescent}, and DNA \cite{fu2017fluorescence}. 

Furthermore, there is a notable interest in developing paper-based aptasensors for applications in a wide range. Paper-based biosensors offer advantages such as low cost, low reagent consumption, rapid response, and a high surface area \cite{zhao2007adaption, weihs2021resonance}. Fluorescent paper-based assays, particularly those incorporating the FRET process, have gained attention due to their easy fabrication, high sensitivity, and selectivity. Additionally, the use of inorganic nano quenchers, such as AuNps, enhances the functionality of paper-based FRET aptasensors \cite{nie2012low, khoshbin2019simple}. 

In this study, a novel paper-based aptasensor containing CdTe QD-aptamer-AuNp complex for early diagnosis of Alzheimer’s Disease is introduced. Targeting the key plasma biomarkers of the disease, p-tau181 proteins and A$\beta$-42 peptides, a less invasive detection method for AD compared to CSF is aimed. In this design, AuNp serves as the acceptor, and CdTe QD serves as the donor, both attached to opposite ends of a DNA aptamer. The binding of the analyte with a specific aptamer induces conformational changes in the aptamer, altering the distance between CdTe QD and AuNp, and subsequently affecting the emitted light intensity from QDs due to FRET. The change in the emitted light intensity is detected by a fluorometry spectrometer, where the change in intensity is expected to be inversely proportional to the analyte concentration. This fluorescent paper-based aptasensor is expected to demonstrate effective monitoring of ultra-trace targets within minutes owing to the properties of QDs and aptamers. With its lower Limit of Detection (LoD), this paper-based FRET aptasensor outperforms traditional biosensors.


\textbf{Aim of the Study}
The first aim of this study is to enhance the early detection of AD through a multifaceted approach that addresses distinctive aspects. Differentiation of AD from other neurodegenerative disorders is pursued, ensuring specificity in diagnostic protocols. Novel treatment avenues are explored, contributing to the development of innovative interventions for this debilitating condition. The second aim is the creation of a practical and cutting-edge biosensor for detection purposes is a primary focus, involving the construction of a FRET system with QDs combined with aptamers and AuNsP for high selectivity. Paper-based systems are explored for enhanced detection efficiency. The third and final aim is the investigation extends to human plasma, where dual analyte detection strategies with high accuracy are devised, emphasizing minimal invasiveness compared to traditional methods. Through these interconnected sub-aims, a significant contribution is aimed at the advancement of early diagnosis of AD methodologies, integrating diverse approaches for increased precision, innovation, and clinical applicability.

\section{Materials and Methods}
\subsection{Aptamer Design}
Aptamers are short single-chain nucleic acid or protein molecules with specific binding properties to a particular target molecule. These molecules are used as an alternative to antibodies, which have recently become popular in scientific studies. In addition to being an alternative, it also promises a promising future with some advantages in different areas. Features of aptamers such as thermal stability, low cost, non-toxicity, and unlimited application are some of the advantages \cite{song2012aptamers}. Aptamers are produced more easily than antibodies. While many colonies need to be screened and examined to produce antibodies, aptamers can be produced by selective expansion of ligands (SELEX) in in vitro conditions \cite{syed2010advances}. The SELEX process is divided into three steps. The first step is pool generation. In this step, a library containing random DNA and RNA sequences is developed. These random sequences consist of 20-40 bases \cite{syed2010advances}. The second step is selection. In this step, the target molecule is processed with its nucleotide sequence. Those with a strong affinity for their goals leave. In the third step, the amplification step, the target to which the sequences are attached is amplified by polymerase chain reaction (PCR) \cite{syed2010advances}. Aptamers can be used in many areas. Aptamer-based biosensors, drugs, drug delivery systems, and bioimaging are some of them. While designing an aptamer-based biosensor, which is one of the usage areas of aptamer, it is taken into consideration the features of aptamers such as low cost, high stability, high affinity for the target molecule, and non-toxicity.
Optimum aptamer sequences specific to p-tau181 proteins and amyloid beta 42 peptides which are 5'-AGT CTA GGA TTC GGC GTG GGT TAA TTT TTT GCG GAG CGT GGC AGG-3' and  5'-AGT CTA GGA TTC GGC GTG GGT TAA TTT TTT GCT GCC TGT GGT GTT GGG GCG GGT GCG-3' are taken from  Chan et al. study and a modified aptamer is obtained by adding thiol to one end and the amino group to the other end to bind with AuNP and QD \cite{chan2019highly}. Ultrapure water (18.2 M$\Omega$.cm) is used to prepare the aqueous solution (5$\mu$M) of aptamers \cite{kim2021aptamer}.

\subsection{AuNPs synthesis}
AuNPs can be easily synthesized and functionalized, making them applicable in sensing, imaging, and photothermal therapy. Their stability and biocompatibility make them suitable for integration into biological systems 
\cite{saha2012gold}. In fluorescence-based analyses, AuNPs increase fluorescence sensitivity and enable the study of interparticle distances by efficiently quenching fluorescence \cite{samanta2014fluorescence}. With easy functionalization using biomolecules, they facilitate targeted delivery and specific interactions in biological contexts. The different optical properties of AuNPs, especially surface plasmon resonance, contribute to imaging and sensing applications \cite{li2016aptamer}. Gold nanoparticles (AuNPs) are prepared according to the paper of Yu et al, which shares similarities with the type of QDs used and is an aptamer-based study \cite{yu2022turn}. For the preparation of AuNPs, each glassware is well cleaned with newly prepared aqua regia (HNO$_3$:HCl = 1:3), thoroughly rinsed with deionized water, and after that, for 2-3 hours oven dried at 373 K \cite{wang2014ligand}. The process, using the citrate reduction method, involves adding 3.5 mL of 1\% trisodium citrate solution to a boiling solution of 0.01\% HAuCl$_4$ (100 mL) under vigorous stirring in a 250 mL round-bottom flask. As the solution changes from light yellow to burgundy, it continues to boil while stirring for 10 minutes before slowly cooling to room temperature. The final step includes purifying the solution through a 0.45 $\mu$m microporous membrane and storing it at 4$^{\circ}$C for later use \cite{yu2022turn}.

\subsection{QD synthesis}

Synthesis of CdTe QDs using chemical reduction Aqueously dispersible MPA-stabilized CdTe QDs were produced in a single chemical reduction. During synthesis, the addition of the thiol stabilizer, MPA to the aqueous CdCl$_2$ solution protects the cadmium ions from oxidation, which might result in insoluble CdTeO$_3$ and Cd(OH)$_2$. Thus, K$_2$TeO$_3$ is added after MPA. The addition of NaBH$_4$ (18 mM) and citric acid (26 mM) initiates the reduction by producing CdTe seed particles. However, due to the acidic pH caused by citric acid and MPA, the liquid becomes dark brown. With dropwise NaOH addition (> pH 7), the brown precipitate became a translucent solution that promotes QD development. Thus, the reaction pH was kept between 8 and 12. 

To start the growth, the resultant clear brown solution was heated for two hours at 80 °C. As would be predicted for CdTe QDs, both the solution and the powders fluoresced brightly when stimulated by a 405 nm diode laser. Te$^{4+}$ was reduced to Te$^{2-}$ during synthesis by either or both of the reducing agents, citric acid and NaBH$_4$. Deprotonation of thiol molecules by an alkaline pH causes Cd$^{2+}$ ions to coordinate with MPA and citrate ions, potentially forming Cd-S-(MPA/citrate) overhangs on the QD surface. Thorough adjustment of the precursor molar ratio, temperature, and pH results in the easy production of photostable and aqueously dispersible CdTe QD powders. Such one-pot operations are very important from a business standpoint. Additionally, it would be beneficial to precisely tune QD photoluminescence to a desired wavelength for multi-color assay applications. The ideal proportions of Cd:MPA:Te: NaBH$_2$:Citric acid precursors have been determined to be 1:2:0.2:3.2:5.2, respectively. Research on how the precursor ratio affects the development of CdTe QDs has been covered in a later section. When kept at 4 °C, the resulting CdTe QDs are both photostable and stable as a colloidal solution for over six months.
\cite{kini2019dual}.


 

\textbf{QD-aptamer-AuNP complex}
According to Azizi et al.’s method \cite{azizi2014plasmonic},the prepared AuNPs and QDs solutions are combined in equal volumes of 250 $\mu$L each.   Afterward, both amyloid beta 42 and p-tau181 specific aptamers at a concentration of 5 $\mu$m are added, with a volume of 25 $\mu$L. The resulting mixture is then subjected to a temperature of 95°C for 10 minutes using a thermoblock. Subsequently, the solution is cooled down to 25°C, which leads to the self-assembly of the plexcitonic hybrid system (PQ). To remove impurities, the PQ is placed in a centrifuge and spun at 14,000 rpm for 10 minutes, after which the supernatant is discarded.   The resulting brownish plate is dissolved in 500 $\mu$L of PBS (1 M, pH = 8). It is then sonicated for 2 minutes and stored at 4°C until needed. Chitosan solution is added to the solution by taking advantage of the solution stabilization feature of chitosan \cite{rinaudo2006chitin}. While preparing the chitosan solution, the procedures in the Rinauda article are applied \cite{rinaudo2006chitin}.


\begin{figure}[t]
	\centering
		\includegraphics[width=1.0\linewidth, height=0.75\linewidth]{Images/complex.png}
	  \caption{The QD-Aptamer-AuNP combination and the possible configurational change resulting from its interaction with the analyte. }\label{fig: complex}
\end{figure}

\subsection{Paper Fabrication}
Whatman No. 1 filter paper is chosen for the 3D $\mu$PAD system because it is a standard filter paper with medium retention and flow rate commonly used in many studies \cite{liana2012recent}. The process of hydrophobization of the lower base of the paper in $\mu$PAD is as follows based on the processes in the study of Songjaroen et al. \cite{songjaroen2011novel}: First, a glass slide is permanently affixed with a magnet on its back. The process of hydrophobization of the lower base of the paper in $\mu$PAD is as follows: First, a glass slide is permanently affixed with a magnet on its back. Then, both of the paper layers, cut to the same size (38 ×60 mm²), are placed over it. Iron molds, cut into the desired shapes of channels and sensing wells, are attached to create a temporary assembly. The magnetic force generated by the magnet ensures a secure connection. Next, the assembly is quickly dipped in and out of wax to create a barrier and heated to 120-130 $^{\circ}$C for 1 second. The base layer of the paper is then left to dry, and after this process, the paper is separated from the glass slide.
In addition to the wax dipping process, the wax printing method using a solid ink printer is also suggested for its ease of use and practicality in paper-based sensors. For the wax printing process, channel patterns (0.2 mm in width and 20 mm in length) of the base layer can be designed using software such as AutoCAD or Adobe Illustrator (CS6). The design can then be printed onto the Whatman No. 1 paper using a Xerox 8560DN printer. The double-sided patterning approach can be used to enhance the resolution of the wax for precise penetration into the paper. The upper base layer is cut by laser using an HP Color LaserJet 4520 on the matching wells of the inlet (4 mm diameter) and sensing wells (3 mm diameter) on the base layer \cite{martinez2011microfluidic, nishat2021based, tenda2016high, pollock2012paper}.

\begin{figure*}[t]
	\centering
		\includegraphics[width=0.65\linewidth, height=0.45\linewidth]{Images/paper.png}
	  \caption{The overall Paper Design. Lamination films, two layers of Whatman paper, the filter, and sensing wells are shown.}\label{fig: paper}
\end{figure*}

Following pattern completion and paper layer cutting, a lamination procedure is carried out to make a semi-enclosed 3D $\mu$PAD. This step helps to reduce contamination and prevent the evaporation of samples. Additionally, lamination enhances the capillary wicking effect, which is important in paper-based systems where the channel width is limited to micrometers, thus limiting the capillary action. To begin, the lower base of the waxed paper is aligned with the one layer of lamination film on the back side, leaving an additional 2 mm on the edges of the lower base paper. Then, the paper with the lamination aligned is exposed to a hot laminator at around 150°C twice, using a non-adhesive polyester sheet to protect the lamination film. The same procedure is done for the upper base on the top surface, except the lamination film is cut through laser cut with a 4 mm diameter on the inlet and 3 mm diameter on the sensing wells as illustrated in Fig. \ref{fig: paper}. This allows for quick melting of the wax in the layers (necessary for wax printing), closure of the channels in later steps, and accurate measurement without any layer that could potentially affect the results in the sensing wells \cite{tenda2016high, de2017simple}.

After treating the papers with hydrophobic material wax and applying lamination, the mixture of QDs, aptamers, and AuNP complex that visualized in Fig.\ref{fig: complex} is introduced into the system. Following the preparation of the CdTe QD-Aptamer specifically designed for A$\beta$-42 and the CdTe QD-Aptamer specifically designed for p-tau181 protein, samples are added to 4 different wells. The specified amount of QD-Aptamer for A$\beta$-42 solutions is added to the first and third well, while the specified amount of QD-Aptamer for p-tau181 analyte solution is added to the second and fourth well (Fig.\ref{fig: paper}). The remaining wells, which serve as the control group, only receive CdTe QD. After an appropriate waiting period for a solution to dry, the wells are thoroughly washed with ultrapure water multiple times for 1 minute to remove any non-adhering quantum dots and purify the system from potential measurement noises \cite{li2021analyte}.


In addition to the main layers of Whatman chromatography paper, another layer of nitrocellulose (25 $mm^2$) with a pore size of 0.45 $\mu$m is incorporated as a filter on the top to complete the three-dimensional (3D) structure of the $\mu$PAD. The nitrocellulose filter is precisely laser cut to cover only 25 millimeters on the sides of the inlet area, ensuring a controlled flow of analytes. This innovative filter allows the passage of substances with a molecular weight equal to or smaller than 80 kDa.

To assemble these 2D layers into 3D $\mu$PAD and ensure their structural integrity and functionality, a high-quality spray adhesive (3M Super 77 Multi-Purpose) is carefully applied. The adhesive is sprayed for precisely 1 second, maintaining a distance of 24 cm from each paper layer's edges. This application method guarantees that only a thin and fine amount of adhesive is used, preventing any compromise to the layers' performance. With these carefully stacked layers to take up average 10 $\mu$L of the sample, the microPAD facilitates the vertical and horizontal movement of samples through an efficient filtration system. This filtration system is illustrated in detail in Fig.\ref{fig: paper}, providing a visual representation of how the analytes traverse the microPAD \cite{wang2019kirigami, lewis2012high}.

\subsection{Preparation of Samples}
To test the accuracy of the biosensor, artificial solutions prepared with different concentrations of analytes are used instead of human plasma in the system.

\textbf{Expression of A$\beta$(M1-42) and p-tau-181}
The protocols are carried out accordingly of \cite{prakash2019rapid, yoo2018efficient, liu2022generation, bolos2017soluble} is shown in \ref{fig: sample}.
A$\beta$(1-42) peptides and p-tau181 proteins are expressed. The A$\beta$(M1-42) variant that contains an N-terminal methionine residue as a start codon is used to express A$\beta$(1-42) peptides \cite{prakash2019rapid}. Moreover; recombinant human A$\beta$(M1-42) and synthetic A$\beta$(1-42) both exhibit aggregate-forming properties \cite{prakash2019rapid}, so A$\beta$(M1-42) will be used for analyses. 

\textbf{Plasmid Isolation}
Firstly, pET-Sac-A$\beta$(M1-42) plasmid and pTXB1 plasmid are isolated. LB medium is prepared and 100 mg/L Ampicillin is added for antibiotic selection during amplification of the plasmids. Then, \textit{Escherichia coli} cells are plated and incubated for approximately 16 hours at 37°C. After incubation, a single colony is selected and inoculated into a 5 mL liquid LB medium containing 100 mg/L ampicillin. The next day, the cells in the tube are inoculated in the 1L flask and incubated until the optical density (OD) reaches 0.45. The plasmids are isolated from the culture using the plasmid isolation miniprep kit.


\textbf{Transformation}
pET-Sac-A$\beta$(M1-42) plasmid is transformed into BL21(DE3)-pLysS competent\textit{ E. coli} by heat shock method. First, the frozen vials of BL21(DE3)pLysS competent cells are thawed. Approximately 1–2 $\mu$L of the isolated plasmid (totaling 50–100 ng DNA) is added to the thawed cells and the tube is gently flicked to mix the plasmid with the cells. The cells and plasmid mixture are incubated on ice for 20 minutes, then placed in a 42 °C water bath for 45 seconds to facilitate plasmid transformation into the cells via the heat shock method. After the heat shock, the tubes are immediately placed on ice for 2 minutes. Transformed bacteria are inoculated into 500 $\mu$L of liquid LB media without antibiotics and the tube is kept in a 37°C shaker for 1 hour at 220 rpm \cite{yoo2018efficient}. For p-tau181 proteins, cDNA encoding tau protein 1-174-(Cys 175) is cloned into pTXB1 plasmid, and the recombinant plasmid is transformed into \textit{E. coli} BL21(DE3) strain. 

\begin{figure*}[t]
	\centering
		\includegraphics[width=0.8\linewidth, height=0.4\linewidth]{Images/sample.png}
	  \caption{Preparation of Samples: Plasmid isolation, transformation, purification, cell lysis, and protein expression steps.}\label{fig: sample}
\end{figure*}

\textbf{Protein Expression}
After transforming the A$\beta$(M1-42) plasmid into \textit{E. coli}, 25-30 $\mu$L of transformed cells are spread onto LB agar plates and incubated overnight. Then, a single colony selected from the plate is inoculated into liquid LB medium and incubated at 220 rpm and 37 °C overnight. The following day, the culture is transferred to a 1 L flask and protein expression is initiated by adding 0.1 mM IPTG. The culture is further incubated at 220 rpm and 37 °C for 4 hours. After that, the culture is centrifuged at 7068X g for 25 minutes at 4 °C. The resulting cell pellet is resuspended in 1X PBS and stored for the next step \cite{yoo2018efficient}. Whereas for p-tau181, transformed cells are cultured in 500 ml of LB media containing 100 mg/ml Ampicillin at 37ºC and 300 rpm of shaking until the culture reaches an optical density of 0.6 - 1.0 at 600 nm. The protocol is followed for p-tau181 with appropriate adjustments on the concentration of IPTG (0.4mM) and centrifugation (3000 g for 15 min at 4°C). The cell pellet is frozen in liquid nitrogen and kept at 80°C  after resuspended in a 10 ml extraction buffer for the next step. 

\textbf{Cell Lysis}
For the cell lysis process, cells for both A$\beta$(M1-42) peptides and p-tau181 proteins are mechanically disrupted by dissolving them in a 25mL Buffer A solution. The sonicator procedure is applied to obtain a homogeneous lysate. It takes place at 60\% amplitude with 30-second pulses, for a total of 4 cycles. After centrifugation at 7068X g for 25 minutes at 4 °C, the obtained pellet is dissolved in 20 mL Buffer B solution, and sonication is performed until the solution appears clear. Finally, the resulting supernatant is filtered through a 0.22 $\mu$m hydrophilic PVDF syringe filter to obtain a clear solution. 

\textbf{Purification Using Reverse-Phase HPLC}
The A$\beta$(M1-42) peptide is purified using reverse-phase HPLC. Firstly, a water bath heated between 60 to 80 °C is prepared using sous vide containing water. The guard and primary columns are fully immersed in this heated water bath. The system is connected to Solvent A and Solvent B bottles through solvent lines, and it is cleaned sequentially with Solvent B and Solvent A. Subsequently, 4 mL from the obtained solution is injected into 18 × 150 mm glass test tubes, and peaks detected at 214 nm are collected with a solvent gradient. Once the purification protocol is completed, fractions of A$\beta$(M1-42) are combined \cite{yoo2018efficient}. 
Additionally to the A$\beta$(M1-42) procedure, affinity purification, and on-column cleavage are carried on with the chitin column for purification of p-Tau181. Then, the protocol given above is followed with the IMPACT Kit E6901.


\textbf{Peptide and Protein Dilution}
A$\beta$(M1-42) peptide samples with concentrations of 25, 50, 75, 100, 125, and 150 pg/mL are prepared according to the human A$\beta$(1-42) protocol provided by Abcam. Before starting the experimental procedure, it is essential to initiate the process by dissolving the peptide in a 1\% PBS-containing solution. Approximately 80 $\mu$L PBS is sufficient for 1 mg of peptide. Using the specified buffer, the solution is diluted to a concentration of 1 mg/mL or less. The peptide cannot be kept in 1\% PBS for an extended time, hence immediate dilution is essential. Vortexing is done quickly since this could promote peptide seeding and additional aggregation. p-tau181 protein samples with the same concentrations as A$\beta$(M1-42) peptide samples are prepared following the procedure recommended by Abcam. 1\% PBS (pH 7.4) solution is used to dissolve proteins, and vortexing is conducted gently. 


\section{Measurement}\label{sec: Measurement}
A typical FRET-based analysis of A$\beta$-42 and p-tau181 is performed as follows: different concentrations of A$\beta$-42 and p-tau181 are added to the sensing well containing QD-aptamer-AuNP complex via samples containing analytes in concentrations of 25, 50, 75, 100, 125, and 150 pg/mL. Afterward, the fluorescence emission spectra are recorded with the excitation of 410 nm. With increasing concentrations of both proteins, fluorescence intensity emitted from CdTe is expected to regularly decrease as a result of an increase in fluorescence quenching of QDs. Thus, the correlation between the degree of fluorescence quenching and the concentration of PSA is expected to be linear to a certain extent.

The UV–vis absorption spectra and fluorescence emission spectra of QDs are shown in Fig. \ref{fig: emission}a. As can be seen, QDs demonstrate broad absorption spectra, whereas the fluorescence emission spectra are relatively narrow, giving them desirable optical characteristics. Under optimized conditions, various concentrations of A$\beta$-42 and p-tau181 are added to the complex of aptamer-QDs-AuNPs in the sensing wells to verify the sensitivity of the assay. The fluorescence intensity at 655 nm is to be measured. Then, values from each pair of wells are averaged, and the average values from the pairs targeting A$\beta$-42 and p-tau181 are compared to the average value of the reference wells. As previously mentioned, it is expected to get a decrease in the fluorescence intensity with increased target molecule concentration as a result of QD quenching (Fig. \ref{fig: emission}b). This expectation is the result of the increase in the energy transfer from QDs to AuNPs due to the distance reduction from aptamer-molecule binding.


\begin{figure*}[t]
	\centering
    \subfigure[]{
    \includegraphics[width=0.45\linewidth,height= 0.35\linewidth]{Images/absorp_emissi_qd.png}}
    \hfill
     \subfigure[]{
    \includegraphics[width=0.5\linewidth,height= 0.35\linewidth]{Images/emission.png}}
	\caption{ (a) UV-vis absorption spectra of QDs (b)Emission spectra of QDs with different analyte concentrations}
	\label{fig: emission}
\end{figure*} 


\section{Conclusion}
In conclusion, this work proposes a novel approach to improving the diagnosis Alzheimer's Disease through the invention of a novel paper-based aptasensor. The combination of Cd-Te QDs, aptamers, and AuNps in a multidimensional design enables the detection of key plasma biomarkers—p-tau181 proteins and A$\beta$(1-42) peptides—in a less invasive way. Also, the FRET method, which is aided by conformational changes in the aptamer caused by analyte binding, enables efficient monitoring of ultra-trace targets. Furthermore, the proposed work contributes to the larger field by investigating innovative therapeutic possibilities for AD and distinguishing it from other neurodegenerative disorders. The creation of a cutting-edge biosensor not only addresses the multifaceted aspects of early detection but also offers a promising technology for facilitating the process of the disease.

\section{Future Directions}
As the designed aptosensor progresses towards mass production and practical application, several avenues for future research and development emerge. One critical aspect deserving attention is the refinement of sensor sensitivity and specificity. Fine-tuning the surface chemistry of the QD-AuNp system will be essential to ensure high-affinity binding, minimizing potential interference from other biomolecules. Moreover, the potential for multi-target detection should be explored, allowing the sensor to simultaneously identify various biomarkers associated with Alzheimer's Disease. This expansion would provide a more holistic diagnostic profile, enhancing the sensor's overall utility.

Moreover, it is essential to investigate ways to further enhance the signal intensity from the designed paper-based aptosensor, potentially enabling a calorimetric sensing method. This enhancement opens the possibility of training an Artificial Intelligence (AI)-based regression model for the detection of A$\beta$-42 and p-tau181 concentrations using images of color changes in the sensing wells of the sensor. Once the model is trained, a patient at risk can capture an image of the biosensor after measurement and upload it to an app integrated with the AI model. The digital screen within the app, coupled with AI capabilities, will convey the concentration of A$\beta$-42 and p-tau181 in the blood. This facilitates the tracking of their levels over time, making early detection and individualized patient care for Alzheimer’s Disease a tangible possibility.

\section{Biosafety}
This project focuses on biosafety and biosecurity aspects. The\textit{ E. coli} BL21 strain, employed during recombinant protein expression, is non-pathogenic and falls under Biosafety Level 1 (BSL1). Processes such as cell seeding and cell harvesting must be carried out within a biosafety cabinet in full compliance with biosafety rules. Special precautions should be taken when working with toxic compounds, such as Cadmium Telluride used in Quantum Dot synthesis. While recycling toxic, chemical, and biological waste, laboratory rules should be observed. Before the experimental stage, all researchers should undergo brief laboratory training to ensure the safe handling of equipment and materials. This is crucial to enhance compliance with biosafety and biosecurity standards.


% To print the credit authorship contribution details
\printcredits

%% Loading bibliography style file
%\bibliographystyle{model1-num-names}
% \bibliographystyle{cas-model2-names}

% Loading bibliography database
% \bibliography{cas-refs}

\bibliographystyle{elsarticle-num} 
\bibliography{cas-refs}


% % Biography
% \bio{}
% % Here goes the biography details.
% \endbio

% \bio{pic1}
% % Here goes the biography details.
% \endbio

\end{document}
