## Supplementary material for "Advancing Early Diagnosis of Alzheimer’s Disease: A Paper-Based Aptasensor for Detecting A*β*(1-42) and p-tau181 from Plasma Using CdTe Quantum Dots": Latex Source: cas-refs.bib

1-@article{knopman2021alzheimer,
  title={Alzheimer disease},
  author={Knopman, David S and Amieva, Helene and Petersen, Ronald C and Ch{\'e}telat, G{\"a}el and Holtzman, David M and Hyman, Bradley T and Nixon, Ralph A and Jones, David T},
  journal={Nature reviews Disease primers},
  volume={7},
  number={1},
  pages={33},
  year={2021},
  publisher={Nature Publishing Group UK London}
}

2-@article{schneider2009neuropathology,
  title={The neuropathology of probable Alzheimer disease and mild cognitive impairment},
  author={Schneider, Julie A and Arvanitakis, Zoe and Leurgans, Sue E and Bennett, David A},
  journal={Annals of Neurology: Official Journal of the American Neurological Association and the Child Neurology Society},
  volume={66},
  number={2},
  pages={200--208},
  year={2009},
  publisher={Wiley Online Library}
}

3-@article{nandi2022global,
  title={Global and regional projections of the economic burden of Alzheimer's disease and related dementias from 2019 to 2050: A value of statistical life approach},
  author={Nandi, Arindam and Counts, Nathaniel and Chen, Simiao and Seligman, Benjamin and Tortorice, Daniel and Vigo, Daniel and Bloom, David E},
  journal={EClinicalMedicine},
  volume={51},
  year={2022},
  publisher={Elsevier}
}


4-@article{sipe2005amyloid,
  title={Amyloid proteins: the beta sheet conformation and disease},
  author={Sipe, Jean D},
  journal={(No Title)},
  year={2005}
}


5-
@article{iqbal2010tau,
  title={Tau in Alzheimer disease and related tauopathies},
  author={Iqbal, Khalid and Liu, Fei and Gong, C-X and Grundke-Iqbal, Inge},
  journal={Current Alzheimer Research},
  volume={7},
  number={8},
  pages={656--664},
  year={2010},
  publisher={Bentham Science Publishers}
}


6-
@article{lu2022high,
  title={A High-Quality CdSe/CdS/ZnS Quantum-Dot-Based FRET Aptasensor for the Simultaneous Detection of Two Different Alzheimer’s Disease Core Biomarkers},
  author={Lu, Xingchang and Hou, Xiaoqi and Tang, Hailin and Yi, Xinyao and Wang, Jianxiu},
  journal={Nanomaterials},
  volume={12},
  number={22},
  pages={4031},
  year={2022},
  publisher={MDPI}
}


7- 
@article{findeis2007role,
  title={The role of amyloid $\beta$ peptide 42 in Alzheimer's disease},
  author={Findeis, Mark A},
  journal={Pharmacology \& therapeutics},
  volume={116},
  number={2},
  pages={266--286},
  year={2007},
  publisher={Elsevier}
}

8-
@article{zetterberg2010amyloid,
  title={Amyloid $\beta$ and APP as biomarkers for Alzheimer’s disease},
  author={Zetterberg, Henrik and Blennow, Kaj and Hanse, Eric},
  journal={Experimental gerontology},
  volume={45},
  number={1},
  pages={23--29},
  year={2010},
  publisher={Elsevier}
}

9-
@article{barthelemy2020blood,
  title={Blood plasma phosphorylated-tau isoforms track CNS change in Alzheimer’s disease},
  author={Barth{\'e}lemy, Nicolas R and Horie, Kanta and Sato, Chihiro and Bateman, Randall J},
  journal={Journal of Experimental Medicine},
  volume={217},
  number={11},
  pages={e20200861},
  year={2020},
  publisher={Rockefeller University Press}
}

10- 
@article{petrella2003neuroimaging,
  title={Neuroimaging and early diagnosis of Alzheimer disease: a look to the future},
  author={Petrella, Jeffrey R and Coleman, R Edward and Doraiswamy, P Murali},
  journal={Radiology},
  volume={226},
  number={2},
  pages={315--336},
  year={2003},
  publisher={Radiological Society of North America}
}

11- @article{valkova2021novel,
  title={Novel trends in electrochemical biosensors for early diagnosis of alzheimer’s disease},
  author={Valkova, Pavla and Pohanka, Miroslav},
  journal={International Journal of Analytical Chemistry},
  volume={2021},
  year={2021},
  publisher={Hindawi}
}

12-@incollection{chepyala2020applications,
  title={Applications and success of MIPs in optical-based nanosensors},
  author={Chepyala, Ramchander},
  booktitle={Nanofabrication for Smart Nanosensor Applications},
  pages={89--121},
  year={2020},
  publisher={Elsevier}
}
13-@article{brazaca2020applications,
  title={Applications of biosensors in Alzheimer's disease diagnosis},
  author={Brazaca, La{\'\i}s Canniatti and Sampaio, Isabella and Zucolotto, Valtencir and Janegitz, Bruno Campos},
  journal={Talanta},
  volume={210},
  pages={120644},
  year={2020},
  publisher={Elsevier}
}
14-@article{kim2020clinically,
  title={Clinically accurate diagnosis of Alzheimer’s disease via multiplexed sensing of core biomarkers in human plasma},
  author={Kim, Kayoung and Kim, Min-Ji and Kim, Da Won and Kim, Su Yeong and Park, Steve and Park, Chan Beum},
  journal={Nature communications},
  volume={11},
  number={1},
  pages={119},
  year={2020},
  publisher={Nature Publishing Group UK London}
}
15-@article{shinohara2021elisa,
  title={ELISA evaluation of tau accumulation in the brains of patients with Alzheimer disease},
  author={Shinohara, Mitsuru and Hirokawa, Junko and Shimodaira, Akemi and Tashiro, Yoshitaka and Suzuki, Kaoru and Gheni, Ghupurjan and Fukumori, Akio and Matsubara, Tomoyasu and Morishima, Maho and Saito, Yuko and others},
  journal={Journal of Neuropathology \& Experimental Neurology},
  volume={80},
  number={7},
  pages={652--662},
  year={2021},
  publisher={Oxford University Press}
}
16- @article{karki2021advances,
  title={Advances in the development paradigm of biosample-based biosensors for early ultrasensitive detection of alzheimer’s disease},
  author={Karki, Hem Prakash and Jang, Yeongseok and Jung, Jinmu and Oh, Jonghyun},
  journal={Journal of nanobiotechnology},
  volume={19},
  pages={1--24},
  year={2021},
  publisher={Springer}
}
17-@article{kim2016aptamer,
  title={Aptamer-based nanobiosensors},
  author={Kim, Yeon Seok and Raston, Nurul Hanun Ahmad and Gu, Man Bock},
  journal={Biosensors and Bioelectronics},
  volume={76},
  pages={2--19},
  year={2016},
  publisher={Elsevier}
}
18-@article{liu2022aptamers,
  title={Aptamers against pathogenic bacteria: Selection strategies and apta-assay/aptasensor application for food safety},
  author={Liu, Mengyue and Yue, Fengling and Kong, Qianqian and Liu, Zhanli and Guo, Yemin and Sun, Xia},
  journal={Journal of Agricultural and Food Chemistry},
  volume={70},
  number={18},
  pages={5477--5498},
  year={2022},
  publisher={ACS Publications}
}
19-@article{khoshbin2020low,
  title={A low-cost paper-based aptasensor for simultaneous trace-level monitoring of mercury (II) and silver (I) ions},
  author={Khoshbin, Zahra and Housaindokht, Mohammad Reza and Verdian, Asma},
  journal={Analytical biochemistry},
  volume={597},
  pages={113689},
  year={2020},
  publisher={Elsevier}
}
20-@article{ghosh2020study,
  title={A study on the response of FRET based DNA aptasensors in intracellular environment},
  author={Ghosh, Shreya and Chen, Yinghua and Sebastian, Jesvin and George, Anne and Dutta, Mitra and Stroscio, Michael A},
  journal={Scientific Reports},
  volume={10},
  number={1},
  pages={13250},
  year={2020},
  publisher={Nature Publishing Group UK London}
}
21-@article{verma2023fret,
  title={FRET Based Biosensor: Principle Applications Recent Advances and Challenges},
  author={Verma, Awadhesh Kumar and Noumani, Ashab and Yadav, Amit K and Solanki, Pratima R},
  journal={Diagnostics},
  volume={13},
  number={8},
  pages={1375},
  year={2023},
  publisher={MDPI}
}
22-@article{day2012fluorescent,
  title={Fluorescent proteins for FRET microscopy: monitoring protein interactions in living cells},
  author={Day, Richard N and Davidson, Michael W},
  journal={Bioessays},
  volume={34},
  number={5},
  pages={341--350},
  year={2012},
  publisher={Wiley Online Library}
}
23-@article{singh2020quantum,
  title={Quantum dots: An emerging tool for point-of-care testing},
  author={Singh, Suchita and Dhawan, Aksha and Karhana, Sonali and Bhat, Madhusudan and Dinda, Amit Kumar},
  journal={Micromachines},
  volume={11},
  number={12},
  pages={1058},
  year={2020},
  publisher={MDPI}
}
24-@article{frigerio2012application,
  title={Application of quantum dots as analytical tools in automated chemical analysis: a review},
  author={Frigerio, Christian and Ribeiro, David SM and Rodrigues, S Sofia M and Abreu, Vera LRG and Barbosa, Jo{\~a}o AC and Prior, Jo{\~a}o AV and Marques, Karine L and Santos, Jo{\~a}o LM},
  journal={Analytica chimica acta},
  volume={735},
  pages={9--22},
  year={2012},
  publisher={Elsevier}
}
25-@article{saha2012gold,
  title={Gold nanoparticles in chemical and biological sensing},
  author={Saha, Krishnendu and Agasti, Sarit S and Kim, Chaekyu and Li, Xiaoning and Rotello, Vincent M},
  journal={Chemical reviews},
  volume={112},
  number={5},
  pages={2739--2779},
  year={2012},
  publisher={ACS Publications}
}
26-@article{chang2017fluorescent,
  title={A fluorescent sensing for glycoproteins based on the FRET between quantum dots and Au nanoparticles},
  author={Chang, Lifang and He, Xiwen and Chen, Langxing and Zhang, Yukui},
  journal={Sensors and Actuators B: Chemical},
  volume={250},
  pages={17--23},
  year={2017},
  publisher={Elsevier}
}
27-@article{fu2017fluorescence,
  title={Fluorescence switch biosensor based on quantum dots and gold nanoparticles for discriminative detection of lysozyme},
  author={Fu, Xuan and Fu, Xing and Wang, Qi and Sheng, Long and Huang, Xi and Ma, Meihu and Cai, Zhaoxia},
  journal={International Journal of Biological Macromolecules},
  volume={103},
  pages={1155--1161},
  year={2017},
  publisher={Elsevier}
}
28-@article{zhao2007adaption,
  title={Adaption of Au nanoparticles and CdTe quantum dots in DNA detection},
  author={Zhao, DAI and Zhang, Jimei and Quanxi, DONG and Ning, GUO and Shichao, XU and Bo, SUN and Yuehua, BU},
  journal={Chinese Journal of Chemical Engineering},
  volume={15},
  number={6},
  pages={791--794},
  year={2007},
  publisher={Elsevier}
}
29-@article{weihs2021resonance,
  title={Resonance energy transfer-based biosensors for point-of-need diagnosis—progress and perspectives},
  author={Weihs, Felix and Anderson, Alisha and Trowell, Stephen and Caron, Karine},
  journal={Sensors},
  volume={21},
  number={2},
  pages={660},
  year={2021},
  publisher={MDPI}
}
30-@article{nie2012low,
  title={Low-cost fabrication of paper-based microfluidic devices by one-step plotting},
  author={Nie, Jinfang and Zhang, Yun and Lin, Liwen and Zhou, Caibin and Li, Shuhuai and Zhang, Lianming and Li, Jianping},
  journal={Analytical chemistry},
  volume={84},
  number={15},
  pages={6331--6335},
  year={2012},
  publisher={ACS Publications}
}
31-@article{khoshbin2019simple,
  title={A simple paper-based aptasensor for ultrasensitive detection of lead (II) ion},
  author={Khoshbin, Zahra and Housaindokht, Mohammad Reza and Izadyar, Mohammad and Verdian, Asma and Bozorgmehr, Mohammad Reza},
  journal={Analytica chimica acta},
  volume={1071},
  pages={70--77},
  year={2019},
  publisher={Elsevier}
}

[1] @article{song2012aptamers,
  title={Aptamers and their biological applications},
  author={Song, Kyung-Mi and Lee, Seonghwan and Ban, Changill},
  journal={Sensors},
  volume={12},
  number={1},
  pages={612--631},
  year={2012},
  publisher={Molecular Diversity Preservation International (MDPI)}
}
[2] @article{syed2010advances,
  title={Advances in aptamers},
  author={Syed, Muhammad Ali and Pervaiz, Saima},
  journal={Oligonucleotides},
  volume={20},
  number={5},
  pages={215--224},
  year={2010},
  publisher={Mary Ann Liebert, Inc. 140 Huguenot Street, 3rd Floor New Rochelle, NY 10801 USA}
}
[5] @article{rinaudo2006chitin,
  title={Chitin and chitosan: Properties and applications},
  author={Rinaudo, Marguerite},
  journal={Progress in polymer science},
  volume={31},
  number={7},
  pages={603--632},
  year={2006},
  publisher={Elsevier}
}
[3] @article{chan2019highly,
  title={Highly sensitive quantification of Alzheimer's disease biomarkers by aptamer-assisted amplification},
  author={Chan, Hei-Nga and Xu, Di and Ho, See-Lok and He, Dinggeng and Wong, Man Shing and Li, Hung-Wing},
  journal={Theranostics},
  volume={9},
  number={10},
  pages={2939},
  year={2019},
  publisher={Ivyspring International Publisher}
}
[4] @article{kim2021aptamer,
  title={Aptamer-conjugated quantum dot optical biosensors: Strategies and applications},
  author={Kim, Dongmin and Yoo, Seungmin},
  journal={Chemosensors},
  volume={9},
  number={11},
  pages={318},
  year={2021},
  publisher={MDPI}
}

[6] @article{azizi2014plasmonic,
  title={Plasmonic circular dichroism study of DNA--gold nanoparticles bioconjugates},
  author={Azizi, Azadeh and Ranjbar, Bijan and Moghadam, Tahereh Tohidi and Bagheri, Zeinab},
  journal={Plasmonics},
  volume={9},
  pages={273--281},
  year={2014},
  publisher={Springer}
}

[1] @article{yu2022turn,
  title={A turn-on fluorescent aptasensor for ampicillin detection based on gold nanoparticles and CdTe QDs},
  author={Yu, Weidao and Hao, Aiyue and Mei, Yanzhen and Yang, Yaqiong and Dai, Chuanchao},
  journal={Microchemical Journal},
  volume={179},
  pages={107454},
  year={2022},
  publisher={Elsevier}
}
[2] @article{wang2014ligand,
  title={Ligand exchange on the surface of cadmium telluride quantum dots with fluorosurfactant-capped gold nanoparticles: synthesis, characterization and toxicity evaluation},
  author={Wang, Lingyun and Zhang, Hongxia and Lu, Chao and Zhao, Lixia},
  journal={Journal of colloid and interface science},
  volume={413},
  pages={140--146},
  year={2014},
  publisher={Elsevier}
}
@article{gaponik2002thiol,
  title={Thiol-capping of CdTe nanocrystals: an alternative to organometallic synthetic routes},
  author={Gaponik, Nikolai and Talapin, Dmitri V and Rogach, Andrey L and Hoppe, Kathrin and Shevchenko, Elena V and Kornowski, Andreas and Eychm{\"u}ller, Alexander and Weller, Horst},
  journal={The Journal of Physical Chemistry B},
  volume={106},
  number={29},
  pages={7177--7185},
  year={2002},
  publisher={ACS Publications}
}

[a] @article{martinez2011microfluidic,
  title={Microfluidic paper-based analytical devices: from POCKET to paper-based ELISA},
  author={Martinez, Andres W},
  journal={Bioanalysis},
  volume={3},
  number={23},
  pages={2589--2592},
  year={2011},
  publisher={Future Science}
}
[b] @article{nishat2021based,
  title={based microfluidics: Simplified fabrication and assay methods},
  author={Nishat, Sumaira and Jafry, Ali Turab and Martinez, Andres W and Awan, Fazli Rabbi},
  journal={Sensors and Actuators B: Chemical},
  volume={336},
  pages={129681},
  year={2021},
  publisher={Elsevier}
}

[c] @article{tenda2016high,
  title={High-resolution microfluidic paper-based analytical devices for sub-microliter sample analysis},
  author={Tenda, Keisuke and Ota, Riki and Yamada, Kentaro and Henares, Terence G and Suzuki, Koji and Citterio, Daniel},
  journal={Micromachines},
  volume={7},
  number={5},
  pages={80},
  year={2016},
  publisher={MDPI}
}

[d] @article{pollock2012paper,
  title={A paper-based multiplexed transaminase test for low-cost, point-of-care liver function testing},
  author={Pollock, Nira R and Rolland, Jason P and Kumar, Shailendra and Beattie, Patrick D and Jain, Sidhartha and Noubary, Farzad and Wong, Vicki L and Pohlmann, Rebecca A and Ryan, Una S and Whitesides, George M},
  journal={Science translational medicine},
  volume={4},
  number={152},
  pages={152ra129--152ra129},
  year={2012},
  publisher={American Association for the Advancement of Science}
}

[e] @article{de2017simple,
  title={A simple method to produce 2D and 3D microfluidic paper-based analytical devices for clinical analysis},
  author={de Oliveira, Ricardo AG and Camargo, Fiamma and Pesquero, Naira C and Faria, Ronaldo Censi},
  journal={Analytica chimica acta},
  volume={957},
  pages={40--46},
  year={2017},
  publisher={Elsevier}
}

[f] @article{wang2019kirigami,
  title={From kirigami to three-dimensional paper-based micro-analytical device: cut-and-paste fabrication and mobile app quantitation},
  author={Wang, Jianhua and Zhang, Lishen and Li, Xiaochun and Zhang, Xiaoliang and Yu, Hua-Zhong},
  journal={RSC advances},
  volume={9},
  number={40},
  pages={23267--23275},
  year={2019},
  publisher={Royal Society of Chemistry}
}

[g] @article{lewis2012high,
  title={High throughput method for prototyping three-dimensional, paper-based microfluidic devices},
  author={Lewis, Gregory G and DiTucci, Matthew J and Baker, Matthew S and Phillips, Scott T},
  journal={Lab on a Chip},
  volume={12},
  number={15},
  pages={2630--2633},
  year={2012},
  publisher={Royal Society of Chemistry}
}
@article{songjaroen2011novel,
  title={Novel, simple and low-cost alternative method for fabrication of paper-based microfluidics by wax dipping},
  author={Songjaroen, Temsiri and Dungchai, Wijitar and Chailapakul, Orawon and Laiwattanapaisal, Wanida},
  journal={Talanta},
  volume={85},
  number={5},
  pages={2587--2593},
  year={2011},
  publisher={Elsevier}
}

@article{li2021analyte,
  title={Analyte-enhanced photocatalytic activity of CdSe/ZnS quantum dots for paper-based colorimetric sensing of Hg2+ under visible light},
  author={Li, Haiyu and Wang, Wan and Wang, Zhijuan and Lv, Qing and Bai, Hua and Zhang, Qing},
  journal={Microchemical Journal},
  volume={164},
  pages={106037},
  year={2021},
  publisher={Elsevier}
}

@article{liana2012recent,
  title={Recent advances in paper-based sensors},
  author={Liana, Devi D and Raguse, Burkhard and Gooding, J Justin and Chow, Edith},
  journal={sensors},
  volume={12},
  number={9},
  pages={11505--11526},
  year={2012},
  publisher={Molecular Diversity Preservation International (MDPI)}
}

1- @article{prakash2019rapid,
  title={Rapid, refined, and robust method for expression, purification, and characterization of recombinant human amyloid beta 1-42},
  author={Prakash, Priya and Lantz, Travis C and Jethava, Krupal P and Chopra, Gaurav},
  journal={Methods and Protocols},
  volume={2},
  number={2},
  pages={48},
  year={2019},
  publisher={MDPI}
}

2- @article{yoo2018efficient,
  title={An efficient method for the expression and purification of A$\beta$ (M1--42)},
  author={Yoo, Stan and Zhang, Sheng and Kreutzer, Adam G and Nowick, James S},
  journal={Biochemistry},
  volume={57},
  number={26},
  pages={3861--3866},
  year={2018},
  publisher={ACS Publications}
}


3- @article{liu2022generation,
  title={Generation and application of semi-synthetic p-Tau181 calibrator for immunoassay calibration},
  author={Liu, Lei and Cai, Yuqi and Lauro, Bianca M and Meunier, Angela L and Chhatwal, Jasmeer and Selkoe, Dennis J},
  journal={Biochemical and Biophysical Research Communications},
  volume={611},
  pages={85--90},
  year={2022},
  publisher={Elsevier}
}
4- @article{bolos2017soluble,
  title={Soluble Tau has devastating effects on the structural plasticity of hippocampal granule neurons},
  author={Bol{\'o}s, Marta and Pallas-Bazarra, Noem{\'\i} and Terreros-Roncal, J and Perea, JR and Jurado-Arjona, Jer{\'o}nimo and {\'A}vila, Jes{\'u}s and Llorens-Mart{\'\i}n, Mar{\'\i}a},
  journal={Translational psychiatry},
  volume={7},
  number={12},
  pages={1267},
  year={2017},
  publisher={Nature Publishing Group UK London}
}


[2] @article{samanta2014fluorescence,
  title={Fluorescence quenching of quantum dots by gold nanoparticles: a potential long range spectroscopic ruler},
  author={Samanta, Anirban and Zhou, Yadong and Zou, Shengli and Yan, Hao and Liu, Yan},
  journal={Nano letters},
  volume={14},
  number={9},
  pages={5052--5057},
  year={2014},
  publisher={ACS Publications}
}
[3] @article{li2016aptamer,
  title={Aptamer-based fluorescent detection of bisphenol A using nonconjugated gold nanoparticles and CdTe quantum dots},
  author={Li, Ying and Xu, Jingyue and Wang, Luokai and Huang, Yanjun and Guo, Jiajia and Cao, Xianyi and Shen, Fei and Luo, Yeli and Sun, Chunyan},
  journal={Sensors and Actuators B: Chemical},
  volume={222},
  pages={815--822},
  year={2016},
  publisher={Elsevier}
}
[4] @incollection{maxwell2020quantum,
  title={Quantum dots},
  author={Maxwell, Tyler and Campos, Maria Gabriela Nogueira and Smith, Stephen and Doomra, Mitsushita and Thwin, Zon and Santra, Swadeshmukul},
  booktitle={Nanoparticles for Biomedical Applications},
  pages={243--265},
  year={2020},
  publisher={Elsevier}
}
@article{kini2019dual,
  title={Dual functionalized, stable and water dispersible CdTe quantum dots: facile, one-pot aqueous synthesis, optical tuning and energy transfer applications},
  author={Kini, Sudarshan and Kulkarni, Suresh D and Ganiga, Vinitha and Nagarakshit, TK and Chidangil, Santhosh},
  journal={Materials Research Bulletin},
  volume={110},
  pages={57--66},
  year={2019},
  publisher={Elsevier}
}
