## Supplementary material for "Advancing Early Diagnosis of Alzheimer’s Disease: A Paper-Based Aptasensor for Detecting A*β*(1-42) and p-tau181 from Plasma Using CdTe Quantum Dots": Latex Source: cas-sc-template.tex

%% 
%% Copyright 2019-2021 Elsevier Ltd
%% 
%% This file is part of the 'CAS Bundle'.
%% --------------------------------------
%% 
%% It may be distributed under the conditions of the LaTeX Project Public
%% License, either version 1.2 of this license or (at your option) any
%% later version.  The latest version of this license is in
%%    http://www.latex-project.org/lppl.txt
%% and version 1.2 or later is part of all distributions of LaTeX
%% version 1999/12/01 or later.
%% 
%% The list of all files belonging to the 'CAS Bundle' is
%% given in the file `manifest.txt'.
%% 
%% Template article for cas-sc documentclass for 
%% single column output.

\documentclass[a4paper,fleqn]{cas-sc}

% If the frontmatter runs over more than one page
% use the longmktitle option.

%\documentclass[a4paper,fleqn,longmktitle]{cas-sc}

%\usepackage[numbers]{natbib}
%\usepackage[authoryear]{natbib}
\usepackage[authoryear,longnamesfirst]{natbib}

%%%Author macros
\def\tsc#1{\csdef{#1}{\textsc{\lowercase{#1}}\xspace}}
\tsc{WGM}
\tsc{QE}
%%%

% Uncomment and use as if needed
%\newtheorem{theorem}{Theorem}
%\newtheorem{lemma}[theorem]{Lemma}
%\newdefinition{rmk}{Remark}
%\newproof{pf}{Proof}
%\newproof{pot}{Proof of Theorem \ref{thm}}

\begin{document}
\let\WriteBookmarks\relax
\def\floatpagepagefraction{1}
\def\textpagefraction{.001}

% Short title
\shorttitle{<short title of the paper for running head>}    

% Short author
\shortauthors{<short author list for running head>}  

% Main title of the paper
\title [mode = title]{<main title>}  

% Title footnote mark
% eg: \tnotemark[1]
\tnotemark[<tnote number>] 

% Title footnote 1.
% eg: \tnotetext[1]{Title footnote text}
\tnotetext[<tnote number>]{<tnote text>} 

% First author
%
% Options: Use if required
% eg: \author[1,3]{Author Name}[type=editor,
%       style=chinese,
%       auid=000,
%       bioid=1,
%       prefix=Sir,
%       orcid=0000-0000-0000-0000,
%       facebook=<facebook id>,
%       twitter=<twitter id>,
%       linkedin=<linkedin id>,
%       gplus=<gplus id>]

%\author[<aff no>]{<author name>}[<options>]

% Corresponding author indication
%\cormark[<corr mark no>]

% Footnote of the first author
%\fnmark[<footnote mark no>]

% Email id of the first author
\ead{<email address>}

% URL of the first author
\ead[url]{<URL>}

% Credit authorship
% eg: \credit{Conceptualization of this study, Methodology, Software}
\credit{<Credit authorship details>}

% Address/affiliation
\affiliation[<aff no>]{organization={},
            addressline={}, 
            city={},
%          citysep={}, % Uncomment if no comma needed between city and postcode
            postcode={}, 
            state={},
            country={}}

%\author[<aff no>]{<author name>}[<options>]

% Footnote of the second author
%\fnmark[2]

% Email id of the second author
\ead{}

% URL of the second author
\ead[url]{}

% Credit authorship
\credit{}

% Address/affiliation
\affiliation[<aff no>]{organization={},
            addressline={}, 
            city={},
%          citysep={}, % Uncomment if no comma needed between city and postcode
            postcode={}, 
            state={},
            country={}}

% Corresponding author text
\cortext[1]{Corresponding author}

% Footnote text
\fntext[1]{}

% For a title note without a number/mark
%\nonumnote{}

% Here goes the abstract
\begin{abstract}
frdhghjhjdg
\end{abstract}

% Use if graphical abstract is present
%\begin{graphicalabstract}
%\includegraphics{}
%\end{graphicalabstract}

% Research highlights
\begin{highlights}
\item 
\item 
\item 
\end{highlights}

% Keywords
% Each keyword is seperated by \sep
\begin{keywords}
 \sep \sep \sep
\end{keywords}

\maketitle

% Main text
\section{}\label{}

% Numbered list
% Use the style of numbering in square brackets.
% If nothing is used, default style will be taken.
%\begin{enumerate}[a)]
%\item 
%\item 
%\item 
%\end{enumerate}  

% Unnumbered list
%\begin{itemize}
%\item 
%\item 
%\item 
%\end{itemize}  

% Description list
%\begin{description}
%\item[]
%\item[] 
%\item[] 
%\end{description}  

% Figure
\begin{figure}[<options>]
	\centering
		\includegraphics[<options>]{}
	  \caption{}\label{fig1}
\end{figure}


\begin{table}[<options>]
\caption{}\label{tbl1}
\begin{tabular*}{\tblwidth}{@{}LL@{}}
\toprule
  &  \\ % Table header row
\midrule
 & \\
 & \\
 & \\
 & \\
\bottomrule
\end{tabular*}
\end{table}

% Uncomment and use as the case may be
%\begin{theorem} 
%\end{theorem}

% Uncomment and use as the case may be
%\begin{lemma} 
%\end{lemma}

%% The Appendices part is started with the command \appendix;
%% appendix sections are then done as normal sections
%% \appendix

\section{}\label{}

% To print the credit authorship contribution details
\printcredits

%% Loading bibliography style file
%\bibliographystyle{model1-num-names}
\bibliographystyle{cas-model2-names}

% Loading bibliography database
\bibliography{}

% Biography
\bio{}
% Here goes the biography details.
\endbio

\bio{pic1}
% Here goes the biography details.
\endbio

\end{document}
