## Supplementary material for "Advancing Early Diagnosis of Alzheimer’s Disease: A Paper-Based Aptasensor for Detecting A*β*(1-42) and p-tau181 from Plasma Using CdTe Quantum Dots": Latex Source: dc-sample.pdf

### This is a specimen $a_b$ title[⋆,⋆⋆]

Sir J.K. Krishnan[a,c,∗,1] (Researcher), Han Thane[b,d], William J. Hansen Jr[b,c,2] (Co-ordinator) and T. Rafeeq[a,c,∗∗,1,3]

[a]*Department of Physics, J.K. Institute of Science, Jawahar Nagar, Trivandrum, 695013, Kerala, India*
[b]*World Scientific University, Street 29, 1011 NX Amsterdam, The Netherlands*
[c]*University of Intelligent Studies, Street 15, Jabaldesh, 825001, Orissa, India*

ARTICLE INFO

*Keywords*:
quadrupole exciton
polariton
WGM
BEC

ABSTRACT

This template helps you to create a properly formatted LaTeX manuscript.
`\beginabstract ... \endabstract` and `\begin{keyword} ... \end{keyword}` which contain the abstract and keywords respectively.
Each keyword shall be separated by a `\sep` command.

#### 1. Introduction

The Elsevier cas-dc class is based on the standard article class and supports almost all of the functionality of that class. In addition, it features commands and options to format the

- document style
- baselineskip
- front matter
- keywords and MSC codes
- theorems, definitions and proofs
- lables of enumerations
- citation style and labeling.

This class depends on the following packages for its proper functioning:

1. natbib.sty for citation processing;
2. geometry.sty for margin settings;
3. fleqn.clo for left aligned equations;
4. graphicx.sty for graphics inclusion;
5. hyperref.sty optional packages if hyperlinking is required in the document;

All the above packages are part of any standard LaTeX installation. Therefore, the users need not be bothered about downloading any extra packages.

#### 2. Installation

The package is available at author resources page at Elsevier (`http://www.elsevier.com/locate/latex`). The class may be moved or copied to a place, usually, `$TEXMF/tex/latex/elsevier/`, or a folder which will be read by LaTeX during document compilation. The TeX file database needs updation after moving/copying class file. Usually, we use commands like `mktexlsr` or `texhash` depending upon the distribution and operating system.

#### 3. Front matter

The author names and affiliations could be formatted in two ways:

(1) Group the authors per affiliation.
(2) Use footnotes to indicate the affiliations.

See the front matter of this document for examples. You are recommended to conform your choice to the journal you are submitting to.

#### 4. Bibliography styles

There are various bibliography styles available. You can select the style of your choice in the preamble of this document. These styles are Elsevier styles based on standard styles like Harvard and Vancouver. Please use BibTeX to generate your bibliography and include DOIs whenever available.

Here are two sample references: [1] [1, 2] [1, 3]

⋆ This document is the results of the research project funded by the National Science Foundation.

⋆⋆ The second title footnote which is a longer text matter to fill through the whole text width and overflow into another line in the footnotes area of the first page.

This note has no numbers. In this work we demonstrate $a_b$ the formation Y_1 of a new type of polariton on the interface between a cuprous oxide slab and a polystyrene micro-sphere placed on the slab.

∗ Corresponding author
∗∗ Principal corresponding author
 (J.K. Krishnan); (W. J. Hansen); (T. Rafeeq)
www.jkkrishnan.in (J.K. Krishnan); https://www.university.org (W. J. Hansen); www.campus.in (T. Rafeeq)
ORCID(s): 0000-0001-0000-0000 (J.K. Krishnan)

[1] This is the first author footnote. but is common to third author as well.

[2] Another author footnote, this is a very long footnote and it should be a really long footnote. But this footnote is not yet sufficiently long enough to make two lines of footnote text.
