## Supplementary material for "Advancing Early Diagnosis of Alzheimer’s Disease: A Paper-Based Aptasensor for Detecting A*β*(1-42) and p-tau181 from Plasma Using CdTe Quantum Dots": Latex Source: elsdoc-cas.pdf

### Documentation for Elsevier's CAS LaTeX template

ELSEVIER LTD

QUICK LINKS
▶ Introduction ▶ Front matter
▶ Main Matter ▶ CRediT...
▶ Bibliography

#### 1. Introduction

This bundle provides two classfiles, namely `cas-sc.cls` and `cas-dc.cls` and corresponding template files for typesetting journal articles supposed to go through Elsevier's updated workflow. `cas-sc.cls` is meant for one-column, the other `cas-dc.cls` for two-column layout. These are now accepted for submitting articles both in Elsevier's electronic submission system and elsewhere.

##### 1.1. Usage

1. `cas-sc.cls` for single column journals.

```
\documentclass[<options>]{cas-sc}
```

2. `cas-dc.cls` for single column journals.

```
\documentclass[<options>]{cas-dc}
```

and have an option `longmktitle` to handle long front matter.

#### 2. Front matter

```
\title [mode = title]{This is a specimen $a_b$ title}
\tnotemark[1,2]

\tnotetext[1]{This document is the results of the research
  project funded by the National Science Foundation.}

\tnotetext[2]{The second title footnote which is a longer text
  matter to fill through the whole text width and overflow
  into another line in the footnotes area of the first page.}
```

© 2021, Elsevier Ltd. Bugs, feature requests, suggestions and comments shall be mailed to <>.

```
\author[1,3]{J.K. Krishnan}[type=editor,
                 auid=000,bioid=1,
                 prefix=Sir,
                 role=Researcher,
                 orcid=0000-0001-0000-0000]
\cormark[1]
\fnmark[1]
\ead{}
\ead[url]{www.jkkrishnan.in}

\credit{Conceptualization of this study,
       Methodology, Software}

\affiliation[1]{organization={Department of Physics,
             J.K. Institute of Science},
            addressline={Jawahar Nagar},
            city={Trivandrum},
%           citysep={}, % Uncomment if no comma needed
%                       between city and postcode
            postcode={695013},
            state={Kerala},
            country={India}}

\author[2,4]{Han Thane}[style=chinese]

\author[2,3]{William {J. Hansen}}[%
  role=Co-ordinator,
  suffix=Jr,
  ]
\fnmark[2]
\ead{}
\ead[URL]{https://www.university.org}

\credit{Data curation, Writing - Original draft preparation}
```

```
\affiliation[2]{organization={World Scientific University},
            addressline={Street 29},
            postcode={1011 NX},
            postcodesep={},
            city={Amsterdam},
            country={The Netherlands}}

\author[1,3]{T. Rafeeq}
\cormark[2]
\fnmark[1,3]
\ead{}
\ead[URL]{www.campus.in}

\affiliation[3]{organization={University of Intelligent
               Studies},
            addressline={Street 15},
            city={Jabaldesh},
            postcode={825001},
            state={Orissa},
            country={India}}

\cortext[cor1]{Corresponding author}
\cortext[cor2]{Principal corresponding author}
\fntext[fn1]{This is the first author footnote, but is common
  to third author as well.}
\fntext[fn2]{Another author footnote, this is a very long
 footnote and it should be a really long footnote. But this
 footnote is not yet sufficiently long enough to make two
 lines of footnote text.}

\nonumnote{This note has no numbers. In this work we
 demonstrate $a_b$ the formation Y\_1 of a new type of
 polariton on the interface between a cuprous oxide slab
 and a polystyrene micro-sphere placed on the slab.
 }
```

```
\begin{abstract}[S U M M A R Y]
This template helps you to create a properly formatted
 \LaTeX\ manuscript.

\begin{abstract} ... \end{abstract} and \begin{keyword}
... \end{keyword} which contain the abstract and keywords
respectively.  Each keyword shall be separated by
a \sep command.
\end{abstract}

\begin{keywords}
quadrupole exciton \sep polariton \sep WGM \sep BEC
\end{keywords}

\maketitle
```

##### 2.1. Title

`\title` command have the below options:

1. `title:` Document title
2. `alt:` Alternate title
3. `sub:` Sub title
4. `trans:` Translated title
5. `transsub:` Translated sub title

```
\title[mode=title]{This is a title}
\title[mode=alt]{This is a alternate title}
\title[mode=sub]{This is a sub title}
\title[mode=trans]{This is a translated title}
\title[mode=transsub]{This is a translated sub title}
```

##### 2.2. Author

`\author` command have the below options:

1. `auid:` Author id
2. `bioid:` Biography id

3. `alt:` Alternate author
4. `style:` Style of author name, eg. chinese
5. `prefix:` Prefix, eg. Sir
6. `suffix:` Suffix
7. `degree:` Degree
8. `role:` Role
9. `orcid:` ORCID
10. `collab:` Collaboration
11. `anon:` Anonymous author
12. `deceased:` Deceased author
13. `twitter:` Twitter account
14. `facebook:` Facebook account
15. `linkedin:` LinkedIn account
16. `plus:` Google plus account
17. `gplus:` Google plus account

```
\author[1,3]{Author Name}[type=editor,
   auid=000,bioid=1,
   prefix=Sir,
   role=Researcher,
   orcid=0000-0001-0000-0000,
   facebook=<facebook id>,
   twitter=<twitter id>,
   linkedin=<linkedin id>,
   gplus=<gplus id>]
```

##### 2.3. Various Marks in the Front Matter

The front matter becomes complicated due to various kinds of notes and marks to the title and author names. Marks in the title will be denoted by a star ($\star$) mark; footnotes are denoted by super scripted Arabic numerals, corresponding author by an Conformal asterisk (*) mark.



Clip 1: Single column output (classfile: cas-sc.cls).

© 2021, Elsevier Ltd. Bugs, feature requests, suggestions and comments shall be mailed to <>.

### This is a specimen $a_b$ title[⋆,⋆⋆]


Sir J.K. Krishnan[a,c,∗,1] (Researcher), Han Thane[b,d], William J. Hansen Jr[b,c,2] (Co-ordinator) and T. Rafeeq[a,c,∗∗,1,3]

J.K. Krishnan et al.: *Preprint submitted to Elsevier* Page 1 of 3

Clip 2: Double column output (classfile: cas-dc.cls).

© 2021, Elsevier Ltd. Bugs, feature requests, suggestions and comments shall be mailed to <>.

##### 2.3.1. Title marks

Title mark can be entered by the command, `\tnotemark[<num>]` and the corresponding text can be entered with the command `\tnotetext[<num>]{<text>}`. An example will be:

```
\title[mode=title]{Leveraging social media news to predict
                stock index movement using RNN-boost}

\tnotemark[1,2]

\tnotetext[1]{This document is the results of the research
  project funded by the National Science Foundation.}

\tnotetext[2]{The second title footnote which is a longer
  text matter to fill through the whole text width and
  overflow into another line in the footnotes area of
  the first page.}
```

`\tnotemark` and `\tnotetext` can be anywhere in the front matter, but should be before `\maketitle` command.

##### 2.3.2. Author marks

Author names can have many kinds of marks and notes:

```
    footnote mark : \fnmark[<num>]
    footnote text : \fntext[<num>]{<text>}
    affiliation mark : \author[<num>]
    email : \ead{<emailid>}
    url : \ead[url]{<url>}
    corresponding author mark : \cormark[<num>]
    corresponding author text : \cortext[<num>]{<text>}
```

##### 2.3.3. Other marks

At times, authors want footnotes which leave no marks in the author names. The note text shall be listed as part of the front matter notes. Class files provides `\nonumnote` for this purpose. The usage

```
\nonumnote{<text>}
```

and should be entered anywhere before the `\maketitle` command for this to take effect.

##### 2.4. Abstract and Keywords

Abstract shall be entered in an environment that starts with `\begin{abstract}` and ends with `\end{abstract}`. Longer abstracts spanning more than one page is also possible in slass file even in double column mode. We need to invoke `longmktitle` option in the class loading line for this to happen smoothly.

The key words are enclosed in a `{keyword}` environment.

```
\begin{abstract}
 This is an abstract. \lipsum[3]
\end{abstract}

\begin{keywords}
 First keyword \sep Second keyword \sep Third
   keyword \sep Fourth keyword
\end{keywords}
```

#### 3. Main Matter

Main matter contains sections, paragraphs, equations and floats like tables, figures, textboxes etc.

##### 3.1. Tables

###### 3.1.1. Normal tables

```
\begin{table}
  \caption{This is a test caption.}
  \begin{tabular*}{\tblwidth}{@{} LLLL@{} }
   \toprule
    Col 1 & Col 2\\
   \midrule
    12345 & 12345\\
    12345 & 12345\\
    12345 & 12345\\
    12345 & 12345\\
    12345 & 12345\\
    12345 & 12345\\
   \bottomrule
  \end{tabular*}
\end{table}
```

###### 3.1.2. Span tables

```
\begin{table*}[width=.9\textwidth,cols=4,pos=h]
 \caption{This is a test caption.}
 \begin{tabular*}{\tblwidth}{@{} LLLLLL@{} }
  \toprule
   Col 1 & Col 2 & Col 3 & Col4 & Col5 & Col6 & Col7\\
  \midrule
   12345 & 12345 & 123 & 12345 & 123 & 12345 & 123 \\
   12345 & 12345 & 123 & 12345 & 123 & 12345 & 123 \\
   12345 & 12345 & 123 & 12345 & 123 & 12345 & 123 \\
   12345 & 12345 & 123 & 12345 & 123 & 12345 & 123 \\
   12345 & 12345 & 123 & 12345 & 123 & 12345 & 123 \\
   12345 & 12345 & 123 & 12345 & 123 & 12345 & 123 \\
   12345 & 12345 & 123 & 12345 & 123 & 12345 & 123 \\
  \bottomrule
 \end{tabular*}
\end{table*}
```

##### 3.2. Figures

###### 3.2.1. Normal figures

```
\begin{figure}
\centering
\includegraphics[scale=.75]{Fig1.pdf}
\caption{The evanescent light - $1S$ quadrupole coupling.
  See also Fig. \protect\ref{FIG:2}).}
\label{FIG:1}
\end{figure}
```

###### 3.2.2. Span figures

```
\begin{figure*}
\centering
 \includegraphics[width=\textwidth,height=2in]{Fig2.pdf}
\caption{Schematic of formation of the evanescent polariton on
linear chain of \PMS.}
 \label{FIG:2}
\end{figure*}
```

© 2021, Elsevier Ltd. Bugs, feature requests, suggestions and comments shall be mailed to <>.

##### 3.3. Theorem and theorem like environments

CAS class file provides a few hooks to format theorems and theorem like environments with ease. All commands the options that are used with `\newtheorem` command will work exactly in the same manner. Class file provides three commands to format theorem or theorem like environments:

1. `\newtheorem` command formats a theorem in LaTeX's default style with italicized font for theorem statement, bold weight for theorem heading and theorem number typeset at the right of theorem heading. It also optionally accepts an argument which will be printed as an extra heading in parentheses. Here is an example coding and output:

```
\newtheorem{theorem}{Theorem}
\begin{theorem}\label{thm}
 The \WGM evanescent field penetration depth into the
 cuprous oxide adjacent crystal is much larger than the
 \QE radius:
 \begin{equation*}
  \lambda_{1S}/2 \pi \left({\epsilon_{Cu2O}-1}
    \right)^{1/2} = 414 \mbox{ \AA} \gg a_B = 4.6
    \mbox{ \AA}
 \end{equation*}
\end{theorem}
```

2. `\newdefinition` command does exactly the same thing as with except that the body font is up-shape instead of italic. See the example below:

```
\newdefinition{definition}{Definition}
\begin{definition}
 The bulk and evanescent polaritons in cuprous oxide
 are formed through the quadrupole part of the light-matter
 interaction:
 \begin{equation*}
  H_{int} = \frac{i e }{m \omega_{1S}} {\bf E}_{i,s}
    \cdot {\bf p}
 \end{equation*}
\end{definition}
```

3. `\newproof` command helps to define proof and custom proof environments without counters as provided in the example code. Given below is an example of proof of theorem kind.

```
\newproof{pot}{Proof of Theorem \ref{thm}}
\begin{pot}
 The photon part of the polariton trapped inside the \PMS
 moves as it would move in a micro-cavity of the effective
 modal volume $V \ll 4 \pi r_{0}^{3} /3$. Consequently, it
 can escape through the evanescent field. This evanescent
 field essentially has a quantum origin and is due to
 tunneling through the potential caused by dielectric
 mismatch on the \PMS surface. Therefore, we define the
 \emph{evanescent} polariton (\EP) as an evanescent light -
 \QE coherent superposition.
\end{pot}
```

##### 3.4. Enumerated and Itemized Lists

CAS class files provides an extended list processing macros which makes the usage a bit more user friendly than the default LaTeX list macros. With an optional argument to the `\begin{enumerate}` command, you can change the list counter type and its attributes. You can see the coding and typeset copy.

```
\begin{enumerate}[1.]
 \item The enumerate environment starts with an optional
     argument ‘1.’ so that the item counter will be suffixed
     by a period as in the optional argument.
 \item If you provide a closing parenthesis to the number in the
     optional argument, the output will have closing
     parenthesis for all the item counters.
 \item You can use ‘(a)’ for alphabetical counter and ‘(i)’ for
     roman counter.
 \begin{enumerate}[a)]
  \item Another level of list with alphabetical counter.
  \item One more item before we start another.
  \begin{enumerate}[(i)]
   \item This item has roman numeral counter.
```

© 2021, Elsevier Ltd. Bugs, feature requests, suggestions and comments shall be mailed to <>.

```
    \item Another one before we close the third level.
   \end{enumerate}
   \item Third item in second level.
  \end{enumerate}
  \item All list items conclude with this step.
\end{enumerate}

\section{Biography}

\verb+\bio+ command have the below options:
\begin{enumerate}
 \item \verb+width:+ Width of the author photo (default is 1in).
 \item \verb+pos:+ Position of author photo.
\end{enumerate}

\begin{vquote}
\bio[width=10mm,pos=l]{tuglogo.jpg}
 \textbf{Another Biography:}
  Recent experimental \cite{HARA:2005} and theoretical
  \cite{DEYCH:2006} studies have shown that the \WGM can travel
  along the chain as "heavy photons". Therefore the \WGM
  acquires the spatial dispersion, and the evanescent
  quadrupole polariton has the form (See Fig.\ref{FIG:3}):
\endbio
```

#### 4. CRediT authorship contribution statement

Give the authorship contribution after each author as

```
 \credit{Conceptualization of this study, Methodology,
      Software}
```

To print the details use `\printcredits`

```
\author[1,3]{J.K. Krishnan}[type=editor,
                auid=000,bioid=1,
                prefix=Sir,
                role=Researcher,
                orcid=0000-0001-0000-0000]
```

```
\cormark[1]
\fnmark[1]
\ead{}
\ead[url]{www.jkkrishnan.in}

\credit{Conceptualization of this study, Methodology, Software}

\affiliation[1]{organization={Department of Physics,
                J.K. Institute of Science},
            addressline={Jawahar Nagar},
            city={Trivandrum},
%           citysep={}, % Uncomment if no comma needed
%                   between city and postcode
            postcode={695013},
            state={Kerala},
            country={India}}

\author[2,4]{Han Thane}[style=chinese]

\author[2,3]{William {J. Hansen}}[%
  role=Co-ordinator,
  suffix=Jr,
  ]
\fnmark[2]
\ead{}
\ead[URL]{https://www.university.org}

\credit{Data curation, Writing - Original draft preparation}

 . . .
 . . .
 . . .
 \printcredits
```

#### 5. Bibliography

For CAS categories, two reference models are recommended. They are model1-num-names.bst and cas-model2-names.bst. Former will format the reference list and their citations according to numbered scheme whereas the latter will format according name-date or author-year style. Authors are requested to choose any one of these according to the journal style. You may download these from

The above bsts are available in the following location for you to download:

https://support.stmdocs.in/wiki/index.php?title=Model-wise_bibliographic_style_files □
