## Supplementary material for "Advancing Early Diagnosis of Alzheimer’s Disease: A Paper-Based Aptasensor for Detecting A*β*(1-42) and p-tau181 from Plasma Using CdTe Quantum Dots": Latex Source: elsdoc-cas.tex

%% 
%% Copyright 2019-2021 Elsevier Ltd
%% 
%% This file is part of the 'CAS Bundle'.
%% ---------------------------------------------
%% 
%% It may be distributed under the conditions of the LaTeX Project Public
%% License, either version 1.2 of this license or (at your option) any
%% later version.  The latest version of this license is in
%%    http://www.latex-project.org/lppl.txt
%% and version 1.2 or later is part of all distributions of LaTeX
%% version 1999/12/01 or later.
%% 
%% The list of all files belonging to the 'CAS Bundle' is
%% given in the file `manifest.txt'.
%%
%% $Id: elsdoc-cas.tex 71 2021-05-12 06:14:05Z rishi $
%%
\documentclass[a4paper,12pt]{article}

\input{glyphtounicode}
\pdfgentounicode=1

\usepackage[xcolor,qtwo]{rvdtx}
\usepackage{multicol}
\usepackage{color}
\usepackage{xspace}
\usepackage{pdfwidgets}
\usepackage{enumerate}

\def\ttdefault{cmtt}

\headsep4pc

\makeatletter
\def\bs{\expandafter\@gobble\string\\}
\def\lb{\expandafter\@gobble\string\{}
\def\rb{\expandafter\@gobble\string\}}
\def\@pdfauthor{C.V.Radhakrishnan}
\def\@pdftitle{CAS templates: A documentation}
\def\@pdfsubject{Document formatting with CAS template}
\def\@pdfkeywords{LaTeX, Elsevier Ltd, document class}
\def\file#1{\textsf{#1}\xspace}

%\def\LastPage{19}

\DeclareRobustCommand{\LaTeX}{L\kern-.26em%
        {\sbox\z@ T%
         \vbox to\ht\z@{\hbox{\check@mathfonts
           \fontsize\sf@size\z@
           \math@fontsfalse\selectfont
          A\,}%
         \vss}%
        }%
     \kern-.15em%
    \TeX}
\makeatother

\def\figurename{Clip}

\setcounter{tocdepth}{1}


\AtBeginDocument{
 \setcounter{topnumber}{2}
 \setcounter{bottomnumber}{2}
 \setcounter{totalnumber}{4}
 \renewcommand{\topfraction}{0.85}
 \renewcommand{\bottomfraction}{0.85}
 \renewcommand{\textfraction}{0.15}
 \renewcommand{\floatpagefraction}{0.7}
}

\begin{document}

\def\testa{This is a specimen document. }
\def\testc{\testa\testa\testa\testa}
\def\testb{\testc\testc\testc\testc\testc}
\long\def\test{\testb\par\testb\par\testb\par}

\pinclude{\copy\contbox\printSq{\LastPage}}

\title{Documentation for Elsevier's CAS \LaTeX\ template}

\author{Elsevier Ltd}
\contact{}

\version{2.3}
\date{\today}
\maketitle

\section{Introduction}
This bundle provides two classfiles, namely \verb+cas-sc.cls+ and
\verb+cas-dc.cls+  and corresponding template files for typesetting
journal articles supposed to go through Elsevier's updated workflow.
\verb+cas-sc.cls+ is meant for one-column, the other 
\verb+cas-dc.cls+ for two-column layout. These are now accepted for
submitting articles both in Elsevier's electronic submission system and
elsewhere.

\subsection{Usage}
\begin{enumerate}
\item \verb+cas-sc.cls+ for single column journals. 

\begin{vquote}
 \documentclass[<options>]{cas-sc}
\end{vquote}
\item \verb+cas-dc.cls+ for single column journals. 

\begin{vquote}
 \documentclass[<options>]{cas-dc}
\end{vquote}
\end{enumerate}
and have an option \verb+longmktitle+ to handle long front matter. 

\section{Front matter}

\begin{vquote}
\title [mode = title]{This is a specimen $a_b$ title}                      
\tnotemark[1,2]

\tnotetext[1]{This document is the results of the research
   project funded by the National Science Foundation.}

\tnotetext[2]{The second title footnote which is a longer text 
   matter to fill through the whole text width and overflow 
   into another line in the footnotes area of the first page.}

\end{vquote}

\begin{vquote}

\author[1,3]{J.K. Krishnan}[type=editor,
                        auid=000,bioid=1,
                        prefix=Sir,
                        role=Researcher,
                        orcid=0000-0001-0000-0000]
\cormark[1]
\fnmark[1]
\ead{}
\ead[url]{www.jkkrishnan.in}

\credit{Conceptualization of this study, 
          Methodology, Software}

\affiliation[1]{organization={Department of Physics, 
                  J.K. Institute of Science},
                addressline={Jawahar Nagar}, 
                city={Trivandrum},
%               citysep={}, % Uncomment if no comma needed 
%                               between city and postcode
                postcode={695013}, 
                state={Kerala},
                country={India}}

\author[2,4]{Han Thane}[style=chinese]

\author[2,3]{William {J. Hansen}}[%
   role=Co-ordinator,
   suffix=Jr,
   ]
\fnmark[2]
\ead{}
\ead[URL]{https://www.university.org}

\credit{Data curation, Writing - Original draft preparation}

\end{vquote}

\begin{vquote}

\affiliation[2]{organization={World Scientific University},
                addressline={Street 29}, 
                postcode={1011 NX}, 
                postcodesep={}, 
                city={Amsterdam},
                country={The Netherlands}}

\author[1,3]{T. Rafeeq}
\cormark[2]
\fnmark[1,3]
\ead{}
\ead[URL]{www.campus.in}

\affiliation[3]{organization={University of Intelligent 
                    Studies},
                addressline={Street 15}, 
                city={Jabaldesh},
                postcode={825001}, 
                state={Orissa}, 
                country={India}}

\cortext[cor1]{Corresponding author}
\cortext[cor2]{Principal corresponding author}
\fntext[fn1]{This is the first author footnote, but is common 
   to third author as well.}
\fntext[fn2]{Another author footnote, this is a very long 
  footnote and it should be a really long footnote. But this 
  footnote is not yet sufficiently long enough to make two 
  lines of footnote text.}

\nonumnote{This note has no numbers. In this work we 
  demonstrate $a_b$ the formation Y\_1 of a new type of 
  polariton on the interface between a cuprous oxide slab 
  and a polystyrene micro-sphere placed on the slab.
  }
\end{vquote}

\begin{vquote}

\begin{abstract}[S U M M A R Y]
This template helps you to create a properly formatted 
 \LaTeX\ manuscript.

\begin{abstract} ... \end{abstract} and \begin{keyword} 
... \end{keyword} which contain the abstract and keywords 
respectively.  Each keyword shall be separated by 
a \sep command.
\end{abstract}

\begin{keywords}
quadrupole exciton \sep polariton \sep WGM \sep BEC
\end{keywords}

\maketitle
\end{vquote}

\subsection{Title}

\verb+\title+ command have the below options:
\begin{enumerate}
\item \verb+title:+ Document title
\item \verb+alt:+ Alternate title
\item \verb+sub:+ Sub title
\item \verb+trans:+ Translated title
\item \verb+transsub:+ Translated sub title
\end{enumerate}

\begin{vquote}
 \title[mode=title]{This is a title}
 \title[mode=alt]{This is a alternate title}
 \title[mode=sub]{This is a sub title}
 \title[mode=trans]{This is a translated title}
 \title[mode=transsub]{This is a translated sub title}
\end{vquote}


\subsection{Author}
\verb+\author+ command have the below options: 

\begin{enumerate}
\item \verb+auid:+ Author id
\item \verb+bioid:+ Biography id
\item \verb+alt:+ Alternate author
\item \verb+style:+ Style of author name, eg.\ chinese
\item \verb+prefix:+ Prefix, eg.\ Sir
\item \verb+suffix:+ Suffix
\item \verb+degree:+ Degree
\item \verb+role:+ Role
\item \verb+orcid:+ ORCID
\item \verb+collab:+ Collaboration
\item \verb+anon:+ Anonymous author
\item \verb+deceased:+ Deceased author
\item \verb+twitter:+ Twitter account
\item \verb+facebook:+ Facebook account
\item \verb+linkedin:+ LinkedIn account
\item \verb+plus:+ Google plus account
\item \verb+gplus:+ Google plus account
\end{enumerate}

\begin{vquote}
\author[1,3]{Author Name}[type=editor,
    auid=000,bioid=1,
    prefix=Sir,
    role=Researcher,
    orcid=0000-0001-0000-0000,
    facebook=<facebook id>,
    twitter=<twitter id>,
    linkedin=<linkedin id>,
    gplus=<gplus id>]
\end{vquote}

\begin{figure}
\includegraphics[width=\textwidth]{sc-sample.pdf}
\caption{Single column output (classfile: cas-sc.cls).}
\end{figure}

\begin{figure}
\includegraphics[width=\textwidth]{dc-sample.pdf}
\caption{Double column output (classfile: cas-dc.cls).}
\end{figure}

\subsection{Various Marks in the Front Matter}

The front matter becomes complicated due to various kinds
of notes and marks to the title and author names. Marks in
the title will be denoted by a star ($\star$) mark;
footnotes are denoted by super scripted Arabic numerals,
corresponding author by an Conformal asterisk (*) mark.

\subsubsection{Title marks}

Title mark can be entered by the command, \verb+\tnotemark[<num>]+
and the corresponding text can be entered with the command
\verb+\tnotetext[<num>]+ \verb+{<text>}+. An example will be:

\begin{vquote}
\title[mode=title]{Leveraging social media news to predict
                      stock index movement using RNN-boost}

\tnotemark[1,2]

\tnotetext[1]{This document is the results of the research
   project funded by the National Science Foundation.}

\tnotetext[2]{The second title footnote which is a longer 
   text matter to fill through the whole text width and 
   overflow into another line in the footnotes area of 
   the first page.}
\end{vquote}

\verb+\tnotemark+ and \verb+\tnotetext+ can be anywhere in
the front matter, but should be before \verb+\maketitle+ command.

\subsubsection{Author marks}

Author names can have many kinds of marks and notes:

\begin{vquote}
    footnote mark : \fnmark[<num>]
    footnote text : \fntext[<num>]{<text>}
    affiliation mark : \author[<num>]
    email : \ead{<emailid>}
    url : \ead[url]{<url>}
    corresponding author mark : \cormark[<num>]
    corresponding author text : \cortext[<num>]{<text>}
\end{vquote}

\subsubsection{Other marks}

At times, authors want footnotes which leave no marks in
the author names. The note text shall be listed as part of
the front matter notes. Class files provides
\verb+\nonumnote+ for this purpose. The usage

\begin{vquote}
\nonumnote{<text>}
\end{vquote}

\noindent and should be entered anywhere before the \verb+\maketitle+
command for this to take effect. 

\subsection{Abstract and Keywords}

Abstract shall be entered in an environment that starts
with\break \verb+\begin{abstract}+ and ends with
\verb+\end{abstract}+. Longer abstracts spanning more than
one page is also possible in slass file even in double
column mode. We need to invoke \verb+longmktitle+ option in the
class loading line for this to happen smoothly.

The key words are enclosed in a \verb+{keyword}+
environment.

\begin{vquote}
\begin{abstract}
 This is an abstract. \lipsum[3]
\end{abstract}

\begin{keywords}
 First keyword \sep Second keyword \sep Third 
    keyword \sep Fourth keyword
\end{keywords}
\end{vquote}

\section{Main Matter}
Main matter contains sections, paragraphs, equations and floats like
tables, figures, textboxes etc.
\subsection{Tables}
\subsubsection{Normal tables}

\begin{vquote}
\begin{table}
  \caption{This is a test caption.}
  \begin{tabular*}{\tblwidth}{@{} LLLL@{} }
   \toprule
    Col 1 & Col 2\\
   \midrule
    12345 & 12345\\
    12345 & 12345\\
    12345 & 12345\\
    12345 & 12345\\
    12345 & 12345\\
    12345 & 12345\\
   \bottomrule
  \end{tabular*}
\end{table}
\end{vquote}

\subsubsection{Span tables}

\begin{vquote}
\begin{table*}[width=.9\textwidth,cols=4,pos=h]
  \caption{This is a test caption.}
  \begin{tabular*}{\tblwidth}{@{} LLLLLL@{} }
   \toprule
    Col 1 & Col 2 & Col 3 & Col4 & Col5 & Col6 & Col7\\
   \midrule
    12345 & 12345 & 123 & 12345 & 123 & 12345 & 123 \\
    12345 & 12345 & 123 & 12345 & 123 & 12345 & 123 \\
    12345 & 12345 & 123 & 12345 & 123 & 12345 & 123 \\
    12345 & 12345 & 123 & 12345 & 123 & 12345 & 123 \\
    12345 & 12345 & 123 & 12345 & 123 & 12345 & 123 \\
    12345 & 12345 & 123 & 12345 & 123 & 12345 & 123 \\
    12345 & 12345 & 123 & 12345 & 123 & 12345 & 123 \\
   \bottomrule
  \end{tabular*}
\end{table*}
\end{vquote}

\subsection{Figures}
\subsubsection{Normal figures}
\begin{vquote}
\begin{figure}
	\centering
		\includegraphics[scale=.75]{Fig1.pdf}
	\caption{The evanescent light - $1S$ quadrupole coupling. 
   See also Fig. \protect\ref{FIG:2}).}
	\label{FIG:1}
\end{figure}
\end{vquote}

\subsubsection{Span figures}

\begin{vquote}
\begin{figure*}
	\centering
	  \includegraphics[width=\textwidth,height=2in]{Fig2.pdf}
	\caption{Schematic of formation of the evanescent polariton on
	linear chain of \PMS.}
  \label{FIG:2}
\end{figure*}\end{vquote}

\subsection{Theorem and theorem like environments}

CAS class file provides a few hooks to format theorems and
theorem like environments with ease. All commands the
options that are used with \verb+\newtheorem+ command will work
exactly in the same manner. Class file provides three
commands to format theorem or theorem like environments:

\begin{enumerate}
\item \verb+\newtheorem+ command formats a theorem in
\LaTeX's default style with italicized font for theorem
statement, bold weight for theorem heading and theorem
number typeset at the right of theorem heading. It also
optionally accepts an argument which will be printed as an
extra heading in parentheses. Here is an example coding and
output:

\begin{vquote}
\newtheorem{theorem}{Theorem}
\begin{theorem}\label{thm}
 The \WGM evanescent field penetration depth into the 
 cuprous oxide adjacent crystal is much larger than the 
 \QE radius: 
 \begin{equation*}
  \lambda_{1S}/2 \pi \left({\epsilon_{Cu2O}-1}
    \right)^{1/2} = 414 \mbox{ \AA} \gg a_B = 4.6 
    \mbox{ \AA}  
 \end{equation*}
\end{theorem}
\end{vquote}

\item \verb+\newdefinition+ command does exactly the same
thing as with except that the body font is up-shape instead
of italic. See the example below:

\begin{vquote}
\newdefinition{definition}{Definition}
\begin{definition}
 The bulk and evanescent polaritons in cuprous oxide
 are formed through the quadrupole part of the light-matter
 interaction:
 \begin{equation*}
  H_{int} = \frac{i e }{m \omega_{1S}} {\bf E}_{i,s} 
    \cdot {\bf p}
 \end{equation*}
\end{definition}
\end{vquote}

\item \verb+\newproof+ command helps to define proof and
custom proof environments without counters as provided in
the example code. Given below is an example of proof of
theorem kind.

\begin{vquote}
\newproof{pot}{Proof of Theorem \ref{thm}}
\begin{pot}
 The photon part of the polariton trapped inside the \PMS
 moves as it would move in a micro-cavity of the effective
 modal volume $V \ll 4 \pi r_{0}^{3} /3$. Consequently, it
 can escape through the evanescent field. This evanescent
 field essentially has a quantum origin and is due to
 tunneling through the potential caused by dielectric
 mismatch on the \PMS surface. Therefore, we define the
 \emph{evanescent} polariton (\EP) as an evanescent light -
 \QE coherent superposition.
\end{pot}
\end{vquote}

\end{enumerate}

\subsection{Enumerated and Itemized Lists}

CAS class files provides an extended list processing macros
which makes the usage a bit more user friendly than the
default LaTeX list macros. With an optional argument to the
\verb+\begin{enumerate}+ command, you can change the list
counter type and its attributes. You can see the coding and
typeset copy. 

\begin{vquote}
\begin{enumerate}[1.]
  \item The enumerate environment starts with an optional
        argument `1.' so that the item counter will be suffixed
        by a period as in the optional argument.
  \item If you provide a closing parenthesis to the number in the
        optional argument, the output will have closing 
        parenthesis for all the item counters.
  \item You can use `(a)' for alphabetical counter and `(i)' for
        roman counter.
  \begin{enumerate}[a)]
    \item Another level of list with alphabetical counter.
    \item One more item before we start another.
    \begin{enumerate}[(i)]
      \item This item has roman numeral counter.
\end{vquote}

\begin{vquote}
      \item Another one before we close the third level.
    \end{enumerate}
    \item Third item in second level.
  \end{enumerate}
  \item All list items conclude with this step.
\end{enumerate}

\section{Biography}

\verb+\bio+ command have the below options:
\begin{enumerate}
 \item \verb+width:+ Width of the author photo (default is 1in).
 \item \verb+pos:+ Position of author photo.
\end{enumerate}

\begin{vquote}
\bio[width=10mm,pos=l]{tuglogo.jpg}
 \textbf{Another Biography:}
  Recent experimental \cite{HARA:2005} and theoretical
  \cite{DEYCH:2006} studies have shown that the \WGM can travel
  along the chain as "heavy photons". Therefore the \WGM 
  acquires the spatial dispersion, and the evanescent 
  quadrupole polariton has the form (See Fig.\ref{FIG:3}):
\endbio
\end{vquote}

\section[CRediT...]{CRediT authorship contribution statement}

Give the authorship contribution after each author as 

\begin{vquote}
 \credit{Conceptualization of this study, Methodology, 
         Software}
\end{vquote}

To print the details use \verb+\printcredits+ 

\begin{vquote}
\author[1,3]{J.K. Krishnan}[type=editor,
                        auid=000,bioid=1,
                        prefix=Sir,
                        role=Researcher,
                        orcid=0000-0001-0000-0000]
\end{vquote}

\begin{vquote}
\cormark[1]
\fnmark[1]
\ead{}
\ead[url]{www.jkkrishnan.in}

\credit{Conceptualization of this study, Methodology, Software}

\affiliation[1]{organization={Department of Physics, 
                      J.K. Institute of Science},
                addressline={Jawahar Nagar}, 
                city={Trivandrum},
%               citysep={}, % Uncomment if no comma needed 
%                           between city and postcode
                postcode={695013}, 
                state={Kerala},
                country={India}}

\author[2,4]{Han Thane}[style=chinese]

\author[2,3]{William {J. Hansen}}[%
   role=Co-ordinator,
   suffix=Jr,
   ]
\fnmark[2]
\ead{}
\ead[URL]{https://www.university.org}

\credit{Data curation, Writing - Original draft preparation}

  . . .
  . . .
  . . .
  \printcredits
\end{vquote}

\section{Bibliography}

For CAS categories, two reference models are recommended.
They are \file{model1-num-names.bst} and \file{cas-model2-names.bst}.
Former will format the reference list and their citations according to
numbered scheme whereas the latter will format according name-date or
author-year style. Authors are requested to choose any one of these
according to the journal style. You may download these from 

The above bsts are available in the following location for you to
download:

\url{https://support.stmdocs.in/wiki/index.php?title=Model-wise_bibliographic_style_files} 
\hfill $\Box$

\end{document}
