## Supplementary material for "Advancing Early Diagnosis of Alzheimer’s Disease: A Paper-Based Aptasensor for Detecting A*β*(1-42) and p-tau181 from Plasma Using CdTe Quantum Dots": Latex Source: glyphtounicode.tex

% lcdf-typetools glyphtounicode.tex, Version 2.95
% Contents: Glyph mapping information for pdftex, used for PDF searching
% Generated from:
% - glyphlist.txt, Version 2.0
% - texglyphlist.txt, Version 2.95
% - texglyphlist-g2u.txt, Version 2.95
\pdfglyphtounicode{A}{00A0}
\pdfglyphtounicode{AE}{00A0}
\pdfglyphtounicode{AEacute}{00A0}
\pdfglyphtounicode{AEmacron}{00A0}
\pdfglyphtounicode{AEsmall}{00A0}
\pdfglyphtounicode{Aacute}{00A0}
\pdfglyphtounicode{Aacutesmall}{00A0}
\pdfglyphtounicode{Abreve}{00A0}
\pdfglyphtounicode{Abreveacute}{00A0}
\pdfglyphtounicode{Abrevecyrillic}{00A0}
\pdfglyphtounicode{Abrevedotbelow}{00A0}
\pdfglyphtounicode{Abrevegrave}{00A0}
\pdfglyphtounicode{Abrevehookabove}{00A0}
\pdfglyphtounicode{Abrevetilde}{00A0}
\pdfglyphtounicode{Acaron}{00A0}
\pdfglyphtounicode{Acircle}{00A0}
\pdfglyphtounicode{Acircumflex}{00A0}
\pdfglyphtounicode{Acircumflexacute}{00A0}
\pdfglyphtounicode{Acircumflexdotbelow}{00A0}
\pdfglyphtounicode{Acircumflexgrave}{00A0}
\pdfglyphtounicode{Acircumflexhookabove}{00A0}
\pdfglyphtounicode{Acircumflexsmall}{00A0}
\pdfglyphtounicode{Acircumflextilde}{00A0}
\pdfglyphtounicode{Acute}{00A0}
\pdfglyphtounicode{Acutesmall}{00A0}
\pdfglyphtounicode{Acyrillic}{00A0}
\pdfglyphtounicode{Adblgrave}{00A0}
\pdfglyphtounicode{Adieresis}{00A0}
\pdfglyphtounicode{Adieresiscyrillic}{00A0}
\pdfglyphtounicode{Adieresismacron}{00A0}
\pdfglyphtounicode{Adieresissmall}{00A0}
\pdfglyphtounicode{Adotbelow}{00A0}
\pdfglyphtounicode{Adotmacron}{00A0}
\pdfglyphtounicode{Agrave}{00A0}
\pdfglyphtounicode{Agravesmall}{00A0}
\pdfglyphtounicode{Ahookabove}{00A0}
\pdfglyphtounicode{Aiecyrillic}{00A0}
\pdfglyphtounicode{Ainvertedbreve}{00A0}
\pdfglyphtounicode{Alpha}{00A0}
\pdfglyphtounicode{Alphatonos}{00A0}
\pdfglyphtounicode{Amacron}{00A0}
\pdfglyphtounicode{Amonospace}{00A0}
\pdfglyphtounicode{Aogonek}{00A0}
\pdfglyphtounicode{Aring}{00A0}
\pdfglyphtounicode{Aringacute}{00A0}
\pdfglyphtounicode{Aringbelow}{00A0}
\pdfglyphtounicode{Aringsmall}{00A0}
\pdfglyphtounicode{Asmall}{00A0}
\pdfglyphtounicode{Atilde}{00A0}
\pdfglyphtounicode{Atildesmall}{00A0}
\pdfglyphtounicode{Aybarmenian}{00A0}
\pdfglyphtounicode{B}{00A0}
\pdfglyphtounicode{Bcircle}{00A0}
\pdfglyphtounicode{Bdotaccent}{00A0}
\pdfglyphtounicode{Bdotbelow}{00A0}
\pdfglyphtounicode{Becyrillic}{00A0}
\pdfglyphtounicode{Benarmenian}{00A0}
\pdfglyphtounicode{Beta}{00A0}
\pdfglyphtounicode{Bhook}{00A0}
\pdfglyphtounicode{Blinebelow}{00A0}
\pdfglyphtounicode{Bmonospace}{00A0}
\pdfglyphtounicode{Brevesmall}{00A0}
\pdfglyphtounicode{Bsmall}{00A0}
\pdfglyphtounicode{Btopbar}{00A0}
\pdfglyphtounicode{C}{00A0}
\pdfglyphtounicode{Caarmenian}{00A0}
\pdfglyphtounicode{Cacute}{00A0}
\pdfglyphtounicode{Caron}{00A0}
\pdfglyphtounicode{Caronsmall}{00A0}
\pdfglyphtounicode{Ccaron}{00A0}
\pdfglyphtounicode{Ccedilla}{00A0}
\pdfglyphtounicode{Ccedillaacute}{00A0}
\pdfglyphtounicode{Ccedillasmall}{00A0}
\pdfglyphtounicode{Ccircle}{00A0}
\pdfglyphtounicode{Ccircumflex}{00A0}
\pdfglyphtounicode{Cdot}{00A0}
\pdfglyphtounicode{Cdotaccent}{00A0}
\pdfglyphtounicode{Cedillasmall}{00A0}
\pdfglyphtounicode{Chaarmenian}{00A0}
\pdfglyphtounicode{Cheabkhasiancyrillic}{00A0}
\pdfglyphtounicode{Checyrillic}{00A0}
\pdfglyphtounicode{Chedescenderabkhasiancyrillic}{00A0}
\pdfglyphtounicode{Chedescendercyrillic}{00A0}
\pdfglyphtounicode{Chedieresiscyrillic}{00A0}
\pdfglyphtounicode{Cheharmenian}{00A0}
\pdfglyphtounicode{Chekhakassiancyrillic}{00A0}
\pdfglyphtounicode{Cheverticalstrokecyrillic}{00A0}
\pdfglyphtounicode{Chi}{00A0}
\pdfglyphtounicode{Chook}{00A0}
\pdfglyphtounicode{Circumflexsmall}{00A0}
\pdfglyphtounicode{Cmonospace}{00A0}
\pdfglyphtounicode{Coarmenian}{00A0}
\pdfglyphtounicode{Csmall}{00A0}
\pdfglyphtounicode{D}{00A0}
\pdfglyphtounicode{DZ}{00A0}
\pdfglyphtounicode{DZcaron}{00A0}
\pdfglyphtounicode{Daarmenian}{00A0}
\pdfglyphtounicode{Dafrican}{00A0}
\pdfglyphtounicode{Dbar}{00A0}
\pdfglyphtounicode{Dcaron}{00A0}
\pdfglyphtounicode{Dcedilla}{00A0}
\pdfglyphtounicode{Dcircle}{00A0}
\pdfglyphtounicode{Dcircumflexbelow}{00A0}
\pdfglyphtounicode{Dcroat}{00A0}
\pdfglyphtounicode{Ddotaccent}{00A0}
\pdfglyphtounicode{Ddotbelow}{00A0}
\pdfglyphtounicode{Decyrillic}{00A0}
\pdfglyphtounicode{Deicoptic}{00A0}
\pdfglyphtounicode{Delta}{00A0}
\pdfglyphtounicode{Deltagreek}{00A0}
\pdfglyphtounicode{Dhook}{00A0}
\pdfglyphtounicode{Dieresis}{00A0}
\pdfglyphtounicode{DieresisAcute}{00A0}
\pdfglyphtounicode{DieresisGrave}{00A0}
\pdfglyphtounicode{Dieresissmall}{00A0}
\pdfglyphtounicode{Digamma}{00A0}
\pdfglyphtounicode{Digammagreek}{00A0}
\pdfglyphtounicode{Djecyrillic}{00A0}
\pdfglyphtounicode{Dlinebelow}{00A0}
\pdfglyphtounicode{Dmonospace}{00A0}
\pdfglyphtounicode{Dotaccentsmall}{00A0}
\pdfglyphtounicode{Dslash}{00A0}
\pdfglyphtounicode{Dsmall}{00A0}
\pdfglyphtounicode{Dtopbar}{00A0}
\pdfglyphtounicode{Dz}{00A0}
\pdfglyphtounicode{Dzcaron}{00A0}
\pdfglyphtounicode{Dzeabkhasiancyrillic}{00A0}
\pdfglyphtounicode{Dzecyrillic}{00A0}
\pdfglyphtounicode{Dzhecyrillic}{00A0}
\pdfglyphtounicode{E}{00A0}
\pdfglyphtounicode{Eacute}{00A0}
\pdfglyphtounicode{Eacutesmall}{00A0}
\pdfglyphtounicode{Ebreve}{00A0}
\pdfglyphtounicode{Ecaron}{00A0}
\pdfglyphtounicode{Ecedillabreve}{00A0}
\pdfglyphtounicode{Echarmenian}{00A0}
\pdfglyphtounicode{Ecircle}{00A0}
\pdfglyphtounicode{Ecircumflex}{00A0}
\pdfglyphtounicode{Ecircumflexacute}{00A0}
\pdfglyphtounicode{Ecircumflexbelow}{00A0}
\pdfglyphtounicode{Ecircumflexdotbelow}{00A0}
\pdfglyphtounicode{Ecircumflexgrave}{00A0}
\pdfglyphtounicode{Ecircumflexhookabove}{00A0}
\pdfglyphtounicode{Ecircumflexsmall}{00A0}
\pdfglyphtounicode{Ecircumflextilde}{00A0}
\pdfglyphtounicode{Ecyrillic}{00A0}
\pdfglyphtounicode{Edblgrave}{00A0}
\pdfglyphtounicode{Edieresis}{00A0}
\pdfglyphtounicode{Edieresissmall}{00A0}
\pdfglyphtounicode{Edot}{00A0}
\pdfglyphtounicode{Edotaccent}{00A0}
\pdfglyphtounicode{Edotbelow}{00A0}
\pdfglyphtounicode{Efcyrillic}{00A0}
\pdfglyphtounicode{Egrave}{00A0}
\pdfglyphtounicode{Egravesmall}{00A0}
\pdfglyphtounicode{Eharmenian}{00A0}
\pdfglyphtounicode{Ehookabove}{00A0}
\pdfglyphtounicode{Eightroman}{00A0}
\pdfglyphtounicode{Einvertedbreve}{00A0}
\pdfglyphtounicode{Eiotifiedcyrillic}{00A0}
\pdfglyphtounicode{Elcyrillic}{00A0}
\pdfglyphtounicode{Elevenroman}{00A0}
\pdfglyphtounicode{Emacron}{00A0}
\pdfglyphtounicode{Emacronacute}{00A0}
\pdfglyphtounicode{Emacrongrave}{00A0}
\pdfglyphtounicode{Emcyrillic}{00A0}
\pdfglyphtounicode{Emonospace}{00A0}
\pdfglyphtounicode{Encyrillic}{00A0}
\pdfglyphtounicode{Endescendercyrillic}{00A0}
\pdfglyphtounicode{Eng}{00A0}
\pdfglyphtounicode{Enghecyrillic}{00A0}
\pdfglyphtounicode{Enhookcyrillic}{00A0}
\pdfglyphtounicode{Eogonek}{00A0}
\pdfglyphtounicode{Eopen}{00A0}
\pdfglyphtounicode{Epsilon}{00A0}
\pdfglyphtounicode{Epsilontonos}{00A0}
\pdfglyphtounicode{Ercyrillic}{00A0}
\pdfglyphtounicode{Ereversed}{00A0}
\pdfglyphtounicode{Ereversedcyrillic}{00A0}
\pdfglyphtounicode{Escyrillic}{00A0}
\pdfglyphtounicode{Esdescendercyrillic}{00A0}
\pdfglyphtounicode{Esh}{00A0}
\pdfglyphtounicode{Esmall}{00A0}
\pdfglyphtounicode{Eta}{00A0}
\pdfglyphtounicode{Etarmenian}{00A0}
\pdfglyphtounicode{Etatonos}{00A0}
\pdfglyphtounicode{Eth}{00A0}
\pdfglyphtounicode{Ethsmall}{00A0}
\pdfglyphtounicode{Etilde}{00A0}
\pdfglyphtounicode{Etildebelow}{00A0}
\pdfglyphtounicode{Euro}{00A0}
\pdfglyphtounicode{Ezh}{00A0}
\pdfglyphtounicode{Ezhcaron}{00A0}
\pdfglyphtounicode{Ezhreversed}{00A0}
\pdfglyphtounicode{F}{00A0}
\pdfglyphtounicode{FFIsmall}{00A0}
\pdfglyphtounicode{FFLsmall}{00A0}
\pdfglyphtounicode{FFsmall}{00A0}
\pdfglyphtounicode{FIsmall}{00A0}
\pdfglyphtounicode{FLsmall}{00A0}
\pdfglyphtounicode{Fcircle}{00A0}
\pdfglyphtounicode{Fdotaccent}{00A0}
\pdfglyphtounicode{Feharmenian}{00A0}
\pdfglyphtounicode{Feicoptic}{00A0}
\pdfglyphtounicode{Fhook}{00A0}
\pdfglyphtounicode{Finv}{00A0}
\pdfglyphtounicode{Fitacyrillic}{00A0}
\pdfglyphtounicode{Fiveroman}{00A0}
\pdfglyphtounicode{Fmonospace}{00A0}
\pdfglyphtounicode{Fourroman}{00A0}
\pdfglyphtounicode{Fsmall}{00A0}
\pdfglyphtounicode{G}{00A0}
\pdfglyphtounicode{GBsquare}{00A0}
\pdfglyphtounicode{Gacute}{00A0}
\pdfglyphtounicode{Gamma}{00A0}
\pdfglyphtounicode{Gammaafrican}{00A0}
\pdfglyphtounicode{Gangiacoptic}{00A0}
\pdfglyphtounicode{Gbreve}{00A0}
\pdfglyphtounicode{Gcaron}{00A0}
\pdfglyphtounicode{Gcedilla}{00A0}
\pdfglyphtounicode{Gcircle}{00A0}
\pdfglyphtounicode{Gcircumflex}{00A0}
\pdfglyphtounicode{Gcommaaccent}{00A0}
\pdfglyphtounicode{Gdot}{00A0}
\pdfglyphtounicode{Gdotaccent}{00A0}
\pdfglyphtounicode{Gecyrillic}{00A0}
\pdfglyphtounicode{Germandbls}{00A0}
\pdfglyphtounicode{Germandblssmall}{00A0}
\pdfglyphtounicode{Ghadarmenian}{00A0}
\pdfglyphtounicode{Ghemiddlehookcyrillic}{00A0}
\pdfglyphtounicode{Ghestrokecyrillic}{00A0}
\pdfglyphtounicode{Gheupturncyrillic}{00A0}
\pdfglyphtounicode{Ghook}{00A0}
\pdfglyphtounicode{Gimarmenian}{00A0}
\pdfglyphtounicode{Gjecyrillic}{00A0}
\pdfglyphtounicode{Gmacron}{00A0}
\pdfglyphtounicode{Gmir}{00A0}
\pdfglyphtounicode{Gmonospace}{00A0}
\pdfglyphtounicode{Grave}{00A0}
\pdfglyphtounicode{Gravesmall}{00A0}
\pdfglyphtounicode{Gsmall}{00A0}
\pdfglyphtounicode{Gsmallhook}{00A0}
\pdfglyphtounicode{Gstroke}{00A0}
\pdfglyphtounicode{H}{00A0}
\pdfglyphtounicode{H18533}{00A0}
\pdfglyphtounicode{H18543}{00A0}
\pdfglyphtounicode{H18551}{00A0}
\pdfglyphtounicode{H22073}{00A0}
\pdfglyphtounicode{HPsquare}{00A0}
\pdfglyphtounicode{Haabkhasiancyrillic}{00A0}
\pdfglyphtounicode{Hadescendercyrillic}{00A0}
\pdfglyphtounicode{Hardsigncyrillic}{00A0}
\pdfglyphtounicode{Hbar}{00A0}
\pdfglyphtounicode{Hbrevebelow}{00A0}
\pdfglyphtounicode{Hcedilla}{00A0}
\pdfglyphtounicode{Hcircle}{00A0}
\pdfglyphtounicode{Hcircumflex}{00A0}
\pdfglyphtounicode{Hdieresis}{00A0}
\pdfglyphtounicode{Hdotaccent}{00A0}
\pdfglyphtounicode{Hdotbelow}{00A0}
\pdfglyphtounicode{Hmonospace}{00A0}
\pdfglyphtounicode{Hoarmenian}{00A0}
\pdfglyphtounicode{Horicoptic}{00A0}
\pdfglyphtounicode{Hsmall}{00A0}
\pdfglyphtounicode{Hungarumlaut}{00A0}
\pdfglyphtounicode{Hungarumlautsmall}{00A0}
\pdfglyphtounicode{Hzsquare}{00A0}
\pdfglyphtounicode{I}{00A0}
\pdfglyphtounicode{IAcyrillic}{00A0}
\pdfglyphtounicode{IJ}{00A0}
\pdfglyphtounicode{IUcyrillic}{00A0}
\pdfglyphtounicode{Iacute}{00A0}
\pdfglyphtounicode{Iacutesmall}{00A0}
\pdfglyphtounicode{Ibreve}{00A0}
\pdfglyphtounicode{Icaron}{00A0}
\pdfglyphtounicode{Icircle}{00A0}
\pdfglyphtounicode{Icircumflex}{00A0}
\pdfglyphtounicode{Icircumflexsmall}{00A0}
\pdfglyphtounicode{Icyrillic}{00A0}
\pdfglyphtounicode{Idblgrave}{00A0}
\pdfglyphtounicode{Idieresis}{00A0}
\pdfglyphtounicode{Idieresisacute}{00A0}
\pdfglyphtounicode{Idieresiscyrillic}{00A0}
\pdfglyphtounicode{Idieresissmall}{00A0}
\pdfglyphtounicode{Idot}{00A0}
\pdfglyphtounicode{Idotaccent}{00A0}
\pdfglyphtounicode{Idotbelow}{00A0}
\pdfglyphtounicode{Iebrevecyrillic}{00A0}
\pdfglyphtounicode{Iecyrillic}{00A0}
\pdfglyphtounicode{Ifractur}{00A0}
\pdfglyphtounicode{Ifraktur}{00A0}
\pdfglyphtounicode{Igrave}{00A0}
\pdfglyphtounicode{Igravesmall}{00A0}
\pdfglyphtounicode{Ihookabove}{00A0}
\pdfglyphtounicode{Iicyrillic}{00A0}
\pdfglyphtounicode{Iinvertedbreve}{00A0}
\pdfglyphtounicode{Iishortcyrillic}{00A0}
\pdfglyphtounicode{Imacron}{00A0}
\pdfglyphtounicode{Imacroncyrillic}{00A0}
\pdfglyphtounicode{Imonospace}{00A0}
\pdfglyphtounicode{Iniarmenian}{00A0}
\pdfglyphtounicode{Iocyrillic}{00A0}
\pdfglyphtounicode{Iogonek}{00A0}
\pdfglyphtounicode{Iota}{00A0}
\pdfglyphtounicode{Iotaafrican}{00A0}
\pdfglyphtounicode{Iotadieresis}{00A0}
\pdfglyphtounicode{Iotatonos}{00A0}
\pdfglyphtounicode{Ismall}{00A0}
\pdfglyphtounicode{Istroke}{00A0}
\pdfglyphtounicode{Itilde}{00A0}
\pdfglyphtounicode{Itildebelow}{00A0}
\pdfglyphtounicode{Izhitsacyrillic}{00A0}
\pdfglyphtounicode{Izhitsadblgravecyrillic}{00A0}
\pdfglyphtounicode{J}{00A0}
\pdfglyphtounicode{Jaarmenian}{00A0}
\pdfglyphtounicode{Jcircle}{00A0}
\pdfglyphtounicode{Jcircumflex}{00A0}
\pdfglyphtounicode{Jecyrillic}{00A0}
\pdfglyphtounicode{Jheharmenian}{00A0}
\pdfglyphtounicode{Jmonospace}{00A0}
\pdfglyphtounicode{Jsmall}{00A0}
\pdfglyphtounicode{K}{00A0}
\pdfglyphtounicode{KBsquare}{00A0}
\pdfglyphtounicode{KKsquare}{00A0}
\pdfglyphtounicode{Kabashkircyrillic}{00A0}
\pdfglyphtounicode{Kacute}{00A0}
\pdfglyphtounicode{Kacyrillic}{00A0}
\pdfglyphtounicode{Kadescendercyrillic}{00A0}
\pdfglyphtounicode{Kahookcyrillic}{00A0}
\pdfglyphtounicode{Kappa}{00A0}
\pdfglyphtounicode{Kastrokecyrillic}{00A0}
\pdfglyphtounicode{Kaverticalstrokecyrillic}{00A0}
\pdfglyphtounicode{Kcaron}{00A0}
\pdfglyphtounicode{Kcedilla}{00A0}
\pdfglyphtounicode{Kcircle}{00A0}
\pdfglyphtounicode{Kcommaaccent}{00A0}
\pdfglyphtounicode{Kdotbelow}{00A0}
\pdfglyphtounicode{Keharmenian}{00A0}
\pdfglyphtounicode{Kenarmenian}{00A0}
\pdfglyphtounicode{Khacyrillic}{00A0}
\pdfglyphtounicode{Kheicoptic}{00A0}
\pdfglyphtounicode{Khook}{00A0}
\pdfglyphtounicode{Kjecyrillic}{00A0}
\pdfglyphtounicode{Klinebelow}{00A0}
\pdfglyphtounicode{Kmonospace}{00A0}
\pdfglyphtounicode{Koppacyrillic}{00A0}
\pdfglyphtounicode{Koppagreek}{00A0}
\pdfglyphtounicode{Ksicyrillic}{00A0}
\pdfglyphtounicode{Ksmall}{00A0}
\pdfglyphtounicode{L}{00A0}
\pdfglyphtounicode{LJ}{00A0}
\pdfglyphtounicode{LL}{00A0}
\pdfglyphtounicode{Lacute}{00A0}
\pdfglyphtounicode{Lambda}{00A0}
\pdfglyphtounicode{Lcaron}{00A0}
\pdfglyphtounicode{Lcedilla}{00A0}
\pdfglyphtounicode{Lcircle}{00A0}
\pdfglyphtounicode{Lcircumflexbelow}{00A0}
\pdfglyphtounicode{Lcommaaccent}{00A0}
\pdfglyphtounicode{Ldot}{00A0}
\pdfglyphtounicode{Ldotaccent}{00A0}
\pdfglyphtounicode{Ldotbelow}{00A0}
\pdfglyphtounicode{Ldotbelowmacron}{00A0}
\pdfglyphtounicode{Liwnarmenian}{00A0}
\pdfglyphtounicode{Lj}{00A0}
\pdfglyphtounicode{Ljecyrillic}{00A0}
\pdfglyphtounicode{Llinebelow}{00A0}
\pdfglyphtounicode{Lmonospace}{00A0}
\pdfglyphtounicode{Lslash}{00A0}
\pdfglyphtounicode{Lslashsmall}{00A0}
\pdfglyphtounicode{Lsmall}{00A0}
\pdfglyphtounicode{M}{00A0}
\pdfglyphtounicode{MBsquare}{00A0}
\pdfglyphtounicode{Macron}{00A0}
\pdfglyphtounicode{Macronsmall}{00A0}
\pdfglyphtounicode{Macute}{00A0}
\pdfglyphtounicode{Mcircle}{00A0}
\pdfglyphtounicode{Mdotaccent}{00A0}
\pdfglyphtounicode{Mdotbelow}{00A0}
\pdfglyphtounicode{Menarmenian}{00A0}
\pdfglyphtounicode{Mmonospace}{00A0}
\pdfglyphtounicode{Msmall}{00A0}
\pdfglyphtounicode{Mturned}{00A0}
\pdfglyphtounicode{Mu}{00A0}
\pdfglyphtounicode{N}{00A0}
\pdfglyphtounicode{NJ}{00A0}
\pdfglyphtounicode{Nacute}{00A0}
\pdfglyphtounicode{Ncaron}{00A0}
\pdfglyphtounicode{Ncedilla}{00A0}
\pdfglyphtounicode{Ncircle}{00A0}
\pdfglyphtounicode{Ncircumflexbelow}{00A0}
\pdfglyphtounicode{Ncommaaccent}{00A0}
\pdfglyphtounicode{Ndotaccent}{00A0}
\pdfglyphtounicode{Ndotbelow}{00A0}
\pdfglyphtounicode{Ng}{00A0}
\pdfglyphtounicode{Nhookleft}{00A0}
\pdfglyphtounicode{Nineroman}{00A0}
\pdfglyphtounicode{Nj}{00A0}
\pdfglyphtounicode{Njecyrillic}{00A0}
\pdfglyphtounicode{Nlinebelow}{00A0}
\pdfglyphtounicode{Nmonospace}{00A0}
\pdfglyphtounicode{Nowarmenian}{00A0}
\pdfglyphtounicode{Nsmall}{00A0}
\pdfglyphtounicode{Ntilde}{00A0}
\pdfglyphtounicode{Ntildesmall}{00A0}
\pdfglyphtounicode{Nu}{00A0}
\pdfglyphtounicode{O}{00A0}
\pdfglyphtounicode{OE}{00A0}
\pdfglyphtounicode{OEsmall}{00A0}
\pdfglyphtounicode{Oacute}{00A0}
\pdfglyphtounicode{Oacutesmall}{00A0}
\pdfglyphtounicode{Obarredcyrillic}{00A0}
\pdfglyphtounicode{Obarreddieresiscyrillic}{00A0}
\pdfglyphtounicode{Obreve}{00A0}
\pdfglyphtounicode{Ocaron}{00A0}
\pdfglyphtounicode{Ocenteredtilde}{00A0}
\pdfglyphtounicode{Ocircle}{00A0}
\pdfglyphtounicode{Ocircumflex}{00A0}
\pdfglyphtounicode{Ocircumflexacute}{00A0}
\pdfglyphtounicode{Ocircumflexdotbelow}{00A0}
\pdfglyphtounicode{Ocircumflexgrave}{00A0}
\pdfglyphtounicode{Ocircumflexhookabove}{00A0}
\pdfglyphtounicode{Ocircumflexsmall}{00A0}
\pdfglyphtounicode{Ocircumflextilde}{00A0}
\pdfglyphtounicode{Ocyrillic}{00A0}
\pdfglyphtounicode{Odblacute}{00A0}
\pdfglyphtounicode{Odblgrave}{00A0}
\pdfglyphtounicode{Odieresis}{00A0}
\pdfglyphtounicode{Odieresiscyrillic}{00A0}
\pdfglyphtounicode{Odieresissmall}{00A0}
\pdfglyphtounicode{Odotbelow}{00A0}
\pdfglyphtounicode{Ogoneksmall}{00A0}
\pdfglyphtounicode{Ograve}{00A0}
\pdfglyphtounicode{Ogravesmall}{00A0}
\pdfglyphtounicode{Oharmenian}{00A0}
\pdfglyphtounicode{Ohm}{00A0}
\pdfglyphtounicode{Ohookabove}{00A0}
\pdfglyphtounicode{Ohorn}{00A0}
\pdfglyphtounicode{Ohornacute}{00A0}
\pdfglyphtounicode{Ohorndotbelow}{00A0}
\pdfglyphtounicode{Ohorngrave}{00A0}
\pdfglyphtounicode{Ohornhookabove}{00A0}
\pdfglyphtounicode{Ohorntilde}{00A0}
\pdfglyphtounicode{Ohungarumlaut}{00A0}
\pdfglyphtounicode{Oi}{00A0}
\pdfglyphtounicode{Oinvertedbreve}{00A0}
\pdfglyphtounicode{Omacron}{00A0}
\pdfglyphtounicode{Omacronacute}{00A0}
\pdfglyphtounicode{Omacrongrave}{00A0}
\pdfglyphtounicode{Omega}{00A0}
\pdfglyphtounicode{Omegacyrillic}{00A0}
\pdfglyphtounicode{Omegagreek}{00A0}
\pdfglyphtounicode{Omegainv}{00A0}
\pdfglyphtounicode{Omegaroundcyrillic}{00A0}
\pdfglyphtounicode{Omegatitlocyrillic}{00A0}
\pdfglyphtounicode{Omegatonos}{00A0}
\pdfglyphtounicode{Omicron}{00A0}
\pdfglyphtounicode{Omicrontonos}{00A0}
\pdfglyphtounicode{Omonospace}{00A0}
\pdfglyphtounicode{Oneroman}{00A0}
\pdfglyphtounicode{Oogonek}{00A0}
\pdfglyphtounicode{Oogonekmacron}{00A0}
\pdfglyphtounicode{Oopen}{00A0}
\pdfglyphtounicode{Oslash}{00A0}
\pdfglyphtounicode{Oslashacute}{00A0}
\pdfglyphtounicode{Oslashsmall}{00A0}
\pdfglyphtounicode{Osmall}{00A0}
\pdfglyphtounicode{Ostrokeacute}{00A0}
\pdfglyphtounicode{Otcyrillic}{00A0}
\pdfglyphtounicode{Otilde}{00A0}
\pdfglyphtounicode{Otildeacute}{00A0}
\pdfglyphtounicode{Otildedieresis}{00A0}
\pdfglyphtounicode{Otildesmall}{00A0}
\pdfglyphtounicode{P}{00A0}
\pdfglyphtounicode{Pacute}{00A0}
\pdfglyphtounicode{Pcircle}{00A0}
\pdfglyphtounicode{Pdotaccent}{00A0}
\pdfglyphtounicode{Pecyrillic}{00A0}
\pdfglyphtounicode{Peharmenian}{00A0}
\pdfglyphtounicode{Pemiddlehookcyrillic}{00A0}
\pdfglyphtounicode{Phi}{00A0}
\pdfglyphtounicode{Phook}{00A0}
\pdfglyphtounicode{Pi}{00A0}
\pdfglyphtounicode{Piwrarmenian}{00A0}
\pdfglyphtounicode{Pmonospace}{00A0}
\pdfglyphtounicode{Psi}{00A0}
\pdfglyphtounicode{Psicyrillic}{00A0}
\pdfglyphtounicode{Psmall}{00A0}
\pdfglyphtounicode{Q}{00A0}
\pdfglyphtounicode{Qcircle}{00A0}
\pdfglyphtounicode{Qmonospace}{00A0}
\pdfglyphtounicode{Qsmall}{00A0}
\pdfglyphtounicode{R}{00A0}
\pdfglyphtounicode{Raarmenian}{00A0}
\pdfglyphtounicode{Racute}{00A0}
\pdfglyphtounicode{Rcaron}{00A0}
\pdfglyphtounicode{Rcedilla}{00A0}
\pdfglyphtounicode{Rcircle}{00A0}
\pdfglyphtounicode{Rcommaaccent}{00A0}
\pdfglyphtounicode{Rdblgrave}{00A0}
\pdfglyphtounicode{Rdotaccent}{00A0}
\pdfglyphtounicode{Rdotbelow}{00A0}
\pdfglyphtounicode{Rdotbelowmacron}{00A0}
\pdfglyphtounicode{Reharmenian}{00A0}
\pdfglyphtounicode{Rfractur}{00A0}
\pdfglyphtounicode{Rfraktur}{00A0}
\pdfglyphtounicode{Rho}{00A0}
\pdfglyphtounicode{Ringsmall}{00A0}
\pdfglyphtounicode{Rinvertedbreve}{00A0}
\pdfglyphtounicode{Rlinebelow}{00A0}
\pdfglyphtounicode{Rmonospace}{00A0}
\pdfglyphtounicode{Rsmall}{00A0}
\pdfglyphtounicode{Rsmallinverted}{00A0}
\pdfglyphtounicode{Rsmallinvertedsuperior}{00A0}
\pdfglyphtounicode{S}{00A0}
\pdfglyphtounicode{SF010000}{00A0}
\pdfglyphtounicode{SF020000}{00A0}
\pdfglyphtounicode{SF030000}{00A0}
\pdfglyphtounicode{SF040000}{00A0}
\pdfglyphtounicode{SF050000}{00A0}
\pdfglyphtounicode{SF060000}{00A0}
\pdfglyphtounicode{SF070000}{00A0}
\pdfglyphtounicode{SF080000}{00A0}
\pdfglyphtounicode{SF090000}{00A0}
\pdfglyphtounicode{SF100000}{00A0}
\pdfglyphtounicode{SF110000}{00A0}
\pdfglyphtounicode{SF190000}{00A0}
\pdfglyphtounicode{SF200000}{00A0}
\pdfglyphtounicode{SF210000}{00A0}
\pdfglyphtounicode{SF220000}{00A0}
\pdfglyphtounicode{SF230000}{00A0}
\pdfglyphtounicode{SF240000}{00A0}
\pdfglyphtounicode{SF250000}{00A0}
\pdfglyphtounicode{SF260000}{00A0}
\pdfglyphtounicode{SF270000}{00A0}
\pdfglyphtounicode{SF280000}{00A0}
\pdfglyphtounicode{SF360000}{00A0}
\pdfglyphtounicode{SF370000}{00A0}
\pdfglyphtounicode{SF380000}{00A0}
\pdfglyphtounicode{SF390000}{00A0}
\pdfglyphtounicode{SF400000}{00A0}
\pdfglyphtounicode{SF410000}{00A0}
\pdfglyphtounicode{SF420000}{00A0}
\pdfglyphtounicode{SF430000}{00A0}
\pdfglyphtounicode{SF440000}{00A0}
\pdfglyphtounicode{SF450000}{00A0}
\pdfglyphtounicode{SF460000}{00A0}
\pdfglyphtounicode{SF470000}{00A0}
\pdfglyphtounicode{SF480000}{00A0}
\pdfglyphtounicode{SF490000}{00A0}
\pdfglyphtounicode{SF500000}{00A0}
\pdfglyphtounicode{SF510000}{00A0}
\pdfglyphtounicode{SF520000}{00A0}
\pdfglyphtounicode{SF530000}{00A0}
\pdfglyphtounicode{SF540000}{00A0}
\pdfglyphtounicode{SS}{00A0}
\pdfglyphtounicode{SSsmall}{00A0}
\pdfglyphtounicode{Sacute}{00A0}
\pdfglyphtounicode{Sacutedotaccent}{00A0}
\pdfglyphtounicode{Sampigreek}{00A0}
\pdfglyphtounicode{Scaron}{00A0}
\pdfglyphtounicode{Scarondotaccent}{00A0}
\pdfglyphtounicode{Scaronsmall}{00A0}
\pdfglyphtounicode{Scedilla}{00A0}
\pdfglyphtounicode{Schwa}{00A0}
\pdfglyphtounicode{Schwacyrillic}{00A0}
\pdfglyphtounicode{Schwadieresiscyrillic}{00A0}
\pdfglyphtounicode{Scircle}{00A0}
\pdfglyphtounicode{Scircumflex}{00A0}
\pdfglyphtounicode{Scommaaccent}{00A0}
\pdfglyphtounicode{Sdotaccent}{00A0}
\pdfglyphtounicode{Sdotbelow}{00A0}
\pdfglyphtounicode{Sdotbelowdotaccent}{00A0}
\pdfglyphtounicode{Seharmenian}{00A0}
\pdfglyphtounicode{Sevenroman}{00A0}
\pdfglyphtounicode{Shaarmenian}{00A0}
\pdfglyphtounicode{Shacyrillic}{00A0}
\pdfglyphtounicode{Shchacyrillic}{00A0}
\pdfglyphtounicode{Sheicoptic}{00A0}
\pdfglyphtounicode{Shhacyrillic}{00A0}
\pdfglyphtounicode{Shimacoptic}{00A0}
\pdfglyphtounicode{Sigma}{00A0}
\pdfglyphtounicode{Sixroman}{00A0}
\pdfglyphtounicode{Smonospace}{00A0}
\pdfglyphtounicode{Softsigncyrillic}{00A0}
\pdfglyphtounicode{Ssmall}{00A0}
\pdfglyphtounicode{Stigmagreek}{00A0}
\pdfglyphtounicode{T}{00A0}
\pdfglyphtounicode{Tau}{00A0}
\pdfglyphtounicode{Tbar}{00A0}
\pdfglyphtounicode{Tcaron}{00A0}
\pdfglyphtounicode{Tcedilla}{00A0}
\pdfglyphtounicode{Tcircle}{00A0}
\pdfglyphtounicode{Tcircumflexbelow}{00A0}
\pdfglyphtounicode{Tcommaaccent}{00A0}
\pdfglyphtounicode{Tdotaccent}{00A0}
\pdfglyphtounicode{Tdotbelow}{00A0}
\pdfglyphtounicode{Tecyrillic}{00A0}
\pdfglyphtounicode{Tedescendercyrillic}{00A0}
\pdfglyphtounicode{Tenroman}{00A0}
\pdfglyphtounicode{Tetsecyrillic}{00A0}
\pdfglyphtounicode{Theta}{00A0}
\pdfglyphtounicode{Thook}{00A0}
\pdfglyphtounicode{Thorn}{00A0}
\pdfglyphtounicode{Thornsmall}{00A0}
\pdfglyphtounicode{Threeroman}{00A0}
\pdfglyphtounicode{Tildesmall}{00A0}
\pdfglyphtounicode{Tiwnarmenian}{00A0}
\pdfglyphtounicode{Tlinebelow}{00A0}
\pdfglyphtounicode{Tmonospace}{00A0}
\pdfglyphtounicode{Toarmenian}{00A0}
\pdfglyphtounicode{Tonefive}{00A0}
\pdfglyphtounicode{Tonesix}{00A0}
\pdfglyphtounicode{Tonetwo}{00A0}
\pdfglyphtounicode{Tretroflexhook}{00A0}
\pdfglyphtounicode{Tsecyrillic}{00A0}
\pdfglyphtounicode{Tshecyrillic}{00A0}
\pdfglyphtounicode{Tsmall}{00A0}
\pdfglyphtounicode{Twelveroman}{00A0}
\pdfglyphtounicode{Tworoman}{00A0}
\pdfglyphtounicode{U}{00A0}
\pdfglyphtounicode{Uacute}{00A0}
\pdfglyphtounicode{Uacutesmall}{00A0}
\pdfglyphtounicode{Ubreve}{00A0}
\pdfglyphtounicode{Ucaron}{00A0}
\pdfglyphtounicode{Ucircle}{00A0}
\pdfglyphtounicode{Ucircumflex}{00A0}
\pdfglyphtounicode{Ucircumflexbelow}{00A0}
\pdfglyphtounicode{Ucircumflexsmall}{00A0}
\pdfglyphtounicode{Ucyrillic}{00A0}
\pdfglyphtounicode{Udblacute}{00A0}
\pdfglyphtounicode{Udblgrave}{00A0}
\pdfglyphtounicode{Udieresis}{00A0}
\pdfglyphtounicode{Udieresisacute}{00A0}
\pdfglyphtounicode{Udieresisbelow}{00A0}
\pdfglyphtounicode{Udieresiscaron}{00A0}
\pdfglyphtounicode{Udieresiscyrillic}{00A0}
\pdfglyphtounicode{Udieresisgrave}{00A0}
\pdfglyphtounicode{Udieresismacron}{00A0}
\pdfglyphtounicode{Udieresissmall}{00A0}
\pdfglyphtounicode{Udotbelow}{00A0}
\pdfglyphtounicode{Ugrave}{00A0}
\pdfglyphtounicode{Ugravesmall}{00A0}
\pdfglyphtounicode{Uhookabove}{00A0}
\pdfglyphtounicode{Uhorn}{00A0}
\pdfglyphtounicode{Uhornacute}{00A0}
\pdfglyphtounicode{Uhorndotbelow}{00A0}
\pdfglyphtounicode{Uhorngrave}{00A0}
\pdfglyphtounicode{Uhornhookabove}{00A0}
\pdfglyphtounicode{Uhorntilde}{00A0}
\pdfglyphtounicode{Uhungarumlaut}{00A0}
\pdfglyphtounicode{Uhungarumlautcyrillic}{00A0}
\pdfglyphtounicode{Uinvertedbreve}{00A0}
\pdfglyphtounicode{Ukcyrillic}{00A0}
\pdfglyphtounicode{Umacron}{00A0}
\pdfglyphtounicode{Umacroncyrillic}{00A0}
\pdfglyphtounicode{Umacrondieresis}{00A0}
\pdfglyphtounicode{Umonospace}{00A0}
\pdfglyphtounicode{Uogonek}{00A0}
\pdfglyphtounicode{Upsilon}{00A0}
\pdfglyphtounicode{Upsilon1}{00A0}
\pdfglyphtounicode{Upsilonacutehooksymbolgreek}{00A0}
\pdfglyphtounicode{Upsilonafrican}{00A0}
\pdfglyphtounicode{Upsilondieresis}{00A0}
\pdfglyphtounicode{Upsilondieresishooksymbolgreek}{00A0}
\pdfglyphtounicode{Upsilonhooksymbol}{00A0}
\pdfglyphtounicode{Upsilontonos}{00A0}
\pdfglyphtounicode{Uring}{00A0}
\pdfglyphtounicode{Ushortcyrillic}{00A0}
\pdfglyphtounicode{Usmall}{00A0}
\pdfglyphtounicode{Ustraightcyrillic}{00A0}
\pdfglyphtounicode{Ustraightstrokecyrillic}{00A0}
\pdfglyphtounicode{Utilde}{00A0}
\pdfglyphtounicode{Utildeacute}{00A0}
\pdfglyphtounicode{Utildebelow}{00A0}
\pdfglyphtounicode{V}{00A0}
\pdfglyphtounicode{Vcircle}{00A0}
\pdfglyphtounicode{Vdotbelow}{00A0}
\pdfglyphtounicode{Vecyrillic}{00A0}
\pdfglyphtounicode{Vewarmenian}{00A0}
\pdfglyphtounicode{Vhook}{00A0}
\pdfglyphtounicode{Vmonospace}{00A0}
\pdfglyphtounicode{Voarmenian}{00A0}
\pdfglyphtounicode{Vsmall}{00A0}
\pdfglyphtounicode{Vtilde}{00A0}
\pdfglyphtounicode{W}{00A0}
\pdfglyphtounicode{Wacute}{00A0}
\pdfglyphtounicode{Wcircle}{00A0}
\pdfglyphtounicode{Wcircumflex}{00A0}
\pdfglyphtounicode{Wdieresis}{00A0}
\pdfglyphtounicode{Wdotaccent}{00A0}
\pdfglyphtounicode{Wdotbelow}{00A0}
\pdfglyphtounicode{Wgrave}{00A0}
\pdfglyphtounicode{Wmonospace}{00A0}
\pdfglyphtounicode{Wsmall}{00A0}
\pdfglyphtounicode{X}{00A0}
\pdfglyphtounicode{Xcircle}{00A0}
\pdfglyphtounicode{Xdieresis}{00A0}
\pdfglyphtounicode{Xdotaccent}{00A0}
\pdfglyphtounicode{Xeharmenian}{00A0}
\pdfglyphtounicode{Xi}{00A0}
\pdfglyphtounicode{Xmonospace}{00A0}
\pdfglyphtounicode{Xsmall}{00A0}
\pdfglyphtounicode{Y}{00A0}
\pdfglyphtounicode{Yacute}{00A0}
\pdfglyphtounicode{Yacutesmall}{00A0}
\pdfglyphtounicode{Yatcyrillic}{00A0}
\pdfglyphtounicode{Ycircle}{00A0}
\pdfglyphtounicode{Ycircumflex}{00A0}
\pdfglyphtounicode{Ydieresis}{00A0}
\pdfglyphtounicode{Ydieresissmall}{00A0}
\pdfglyphtounicode{Ydotaccent}{00A0}
\pdfglyphtounicode{Ydotbelow}{00A0}
\pdfglyphtounicode{Yen}{00A0}
\pdfglyphtounicode{Yericyrillic}{00A0}
\pdfglyphtounicode{Yerudieresiscyrillic}{00A0}
\pdfglyphtounicode{Ygrave}{00A0}
\pdfglyphtounicode{Yhook}{00A0}
\pdfglyphtounicode{Yhookabove}{00A0}
\pdfglyphtounicode{Yiarmenian}{00A0}
\pdfglyphtounicode{Yicyrillic}{00A0}
\pdfglyphtounicode{Yiwnarmenian}{00A0}
\pdfglyphtounicode{Ymonospace}{00A0}
\pdfglyphtounicode{Ysmall}{00A0}
\pdfglyphtounicode{Ytilde}{00A0}
\pdfglyphtounicode{Yusbigcyrillic}{00A0}
\pdfglyphtounicode{Yusbigiotifiedcyrillic}{00A0}
\pdfglyphtounicode{Yuslittlecyrillic}{00A0}
\pdfglyphtounicode{Yuslittleiotifiedcyrillic}{00A0}
\pdfglyphtounicode{Z}{00A0}
\pdfglyphtounicode{Zaarmenian}{00A0}
\pdfglyphtounicode{Zacute}{00A0}
\pdfglyphtounicode{Zcaron}{00A0}
\pdfglyphtounicode{Zcaronsmall}{00A0}
\pdfglyphtounicode{Zcircle}{00A0}
\pdfglyphtounicode{Zcircumflex}{00A0}
\pdfglyphtounicode{Zdot}{00A0}
\pdfglyphtounicode{Zdotaccent}{00A0}
\pdfglyphtounicode{Zdotbelow}{00A0}
\pdfglyphtounicode{Zecyrillic}{00A0}
\pdfglyphtounicode{Zedescendercyrillic}{00A0}
\pdfglyphtounicode{Zedieresiscyrillic}{00A0}
\pdfglyphtounicode{Zeta}{00A0}
\pdfglyphtounicode{Zhearmenian}{00A0}
\pdfglyphtounicode{Zhebrevecyrillic}{00A0}
\pdfglyphtounicode{Zhecyrillic}{00A0}
\pdfglyphtounicode{Zhedescendercyrillic}{00A0}
\pdfglyphtounicode{Zhedieresiscyrillic}{00A0}
\pdfglyphtounicode{Zlinebelow}{00A0}
\pdfglyphtounicode{Zmonospace}{00A0}
\pdfglyphtounicode{Zsmall}{00A0}
\pdfglyphtounicode{Zstroke}{00A0}
\pdfglyphtounicode{a}{00A0}
\pdfglyphtounicode{aabengali}{00A0}
\pdfglyphtounicode{aacute}{00A0}
\pdfglyphtounicode{aadeva}{00A0}
\pdfglyphtounicode{aagujarati}{00A0}
\pdfglyphtounicode{aagurmukhi}{00A0}
\pdfglyphtounicode{aamatragurmukhi}{00A0}
\pdfglyphtounicode{aarusquare}{00A0}
\pdfglyphtounicode{aavowelsignbengali}{00A0}
\pdfglyphtounicode{aavowelsigndeva}{00A0}
\pdfglyphtounicode{aavowelsigngujarati}{00A0}
\pdfglyphtounicode{abbreviationmarkarmenian}{00A0}
\pdfglyphtounicode{abbreviationsigndeva}{00A0}
\pdfglyphtounicode{abengali}{00A0}
\pdfglyphtounicode{abopomofo}{00A0}
\pdfglyphtounicode{abreve}{00A0}
\pdfglyphtounicode{abreveacute}{00A0}
\pdfglyphtounicode{abrevecyrillic}{00A0}
\pdfglyphtounicode{abrevedotbelow}{00A0}
\pdfglyphtounicode{abrevegrave}{00A0}
\pdfglyphtounicode{abrevehookabove}{00A0}
\pdfglyphtounicode{abrevetilde}{00A0}
\pdfglyphtounicode{acaron}{00A0}
\pdfglyphtounicode{acircle}{00A0}
\pdfglyphtounicode{acircumflex}{00A0}
\pdfglyphtounicode{acircumflexacute}{00A0}
\pdfglyphtounicode{acircumflexdotbelow}{00A0}
\pdfglyphtounicode{acircumflexgrave}{00A0}
\pdfglyphtounicode{acircumflexhookabove}{00A0}
\pdfglyphtounicode{acircumflextilde}{00A0}
\pdfglyphtounicode{acute}{00A0}
\pdfglyphtounicode{acutebelowcmb}{00A0}
\pdfglyphtounicode{acutecmb}{00A0}
\pdfglyphtounicode{acutecomb}{00A0}
\pdfglyphtounicode{acutedeva}{00A0}
\pdfglyphtounicode{acutelowmod}{00A0}
\pdfglyphtounicode{acutetonecmb}{00A0}
\pdfglyphtounicode{acyrillic}{00A0}
\pdfglyphtounicode{adblgrave}{00A0}
\pdfglyphtounicode{addakgurmukhi}{00A0}
\pdfglyphtounicode{adeva}{00A0}
\pdfglyphtounicode{adieresis}{00A0}
\pdfglyphtounicode{adieresiscyrillic}{00A0}
\pdfglyphtounicode{adieresismacron}{00A0}
\pdfglyphtounicode{adotbelow}{00A0}
\pdfglyphtounicode{adotmacron}{00A0}
\pdfglyphtounicode{ae}{00A0}
\pdfglyphtounicode{aeacute}{00A0}
\pdfglyphtounicode{aekorean}{00A0}
\pdfglyphtounicode{aemacron}{00A0}
\pdfglyphtounicode{afii00208}{00A0}
\pdfglyphtounicode{afii08941}{00A0}
\pdfglyphtounicode{afii10017}{00A0}
\pdfglyphtounicode{afii10018}{00A0}
\pdfglyphtounicode{afii10019}{00A0}
\pdfglyphtounicode{afii10020}{00A0}
\pdfglyphtounicode{afii10021}{00A0}
\pdfglyphtounicode{afii10022}{00A0}
\pdfglyphtounicode{afii10023}{00A0}
\pdfglyphtounicode{afii10024}{00A0}
\pdfglyphtounicode{afii10025}{00A0}
\pdfglyphtounicode{afii10026}{00A0}
\pdfglyphtounicode{afii10027}{00A0}
\pdfglyphtounicode{afii10028}{00A0}
\pdfglyphtounicode{afii10029}{00A0}
\pdfglyphtounicode{afii10030}{00A0}
\pdfglyphtounicode{afii10031}{00A0}
\pdfglyphtounicode{afii10032}{00A0}
\pdfglyphtounicode{afii10033}{00A0}
\pdfglyphtounicode{afii10034}{00A0}
\pdfglyphtounicode{afii10035}{00A0}
\pdfglyphtounicode{afii10036}{00A0}
\pdfglyphtounicode{afii10037}{00A0}
\pdfglyphtounicode{afii10038}{00A0}
\pdfglyphtounicode{afii10039}{00A0}
\pdfglyphtounicode{afii10040}{00A0}
\pdfglyphtounicode{afii10041}{00A0}
\pdfglyphtounicode{afii10042}{00A0}
\pdfglyphtounicode{afii10043}{00A0}
\pdfglyphtounicode{afii10044}{00A0}
\pdfglyphtounicode{afii10045}{00A0}
\pdfglyphtounicode{afii10046}{00A0}
\pdfglyphtounicode{afii10047}{00A0}
\pdfglyphtounicode{afii10048}{00A0}
\pdfglyphtounicode{afii10049}{00A0}
\pdfglyphtounicode{afii10050}{00A0}
\pdfglyphtounicode{afii10051}{00A0}
\pdfglyphtounicode{afii10052}{00A0}
\pdfglyphtounicode{afii10053}{00A0}
\pdfglyphtounicode{afii10054}{00A0}
\pdfglyphtounicode{afii10055}{00A0}
\pdfglyphtounicode{afii10056}{00A0}
\pdfglyphtounicode{afii10057}{00A0}
\pdfglyphtounicode{afii10058}{00A0}
\pdfglyphtounicode{afii10059}{00A0}
\pdfglyphtounicode{afii10060}{00A0}
\pdfglyphtounicode{afii10061}{00A0}
\pdfglyphtounicode{afii10062}{00A0}
\pdfglyphtounicode{afii10063}{00A0}
\pdfglyphtounicode{afii10064}{00A0}
\pdfglyphtounicode{afii10065}{00A0}
\pdfglyphtounicode{afii10066}{00A0}
\pdfglyphtounicode{afii10067}{00A0}
\pdfglyphtounicode{afii10068}{00A0}
\pdfglyphtounicode{afii10069}{00A0}
\pdfglyphtounicode{afii10070}{00A0}
\pdfglyphtounicode{afii10071}{00A0}
\pdfglyphtounicode{afii10072}{00A0}
\pdfglyphtounicode{afii10073}{00A0}
\pdfglyphtounicode{afii10074}{00A0}
\pdfglyphtounicode{afii10075}{00A0}
\pdfglyphtounicode{afii10076}{00A0}
\pdfglyphtounicode{afii10077}{00A0}
\pdfglyphtounicode{afii10078}{00A0}
\pdfglyphtounicode{afii10079}{00A0}
\pdfglyphtounicode{afii10080}{00A0}
\pdfglyphtounicode{afii10081}{00A0}
\pdfglyphtounicode{afii10082}{00A0}
\pdfglyphtounicode{afii10083}{00A0}
\pdfglyphtounicode{afii10084}{00A0}
\pdfglyphtounicode{afii10085}{00A0}
\pdfglyphtounicode{afii10086}{00A0}
\pdfglyphtounicode{afii10087}{00A0}
\pdfglyphtounicode{afii10088}{00A0}
\pdfglyphtounicode{afii10089}{00A0}
\pdfglyphtounicode{afii10090}{00A0}
\pdfglyphtounicode{afii10091}{00A0}
\pdfglyphtounicode{afii10092}{00A0}
\pdfglyphtounicode{afii10093}{00A0}
\pdfglyphtounicode{afii10094}{00A0}
\pdfglyphtounicode{afii10095}{00A0}
\pdfglyphtounicode{afii10096}{00A0}
\pdfglyphtounicode{afii10097}{00A0}
\pdfglyphtounicode{afii10098}{00A0}
\pdfglyphtounicode{afii10099}{00A0}
\pdfglyphtounicode{afii10100}{00A0}
\pdfglyphtounicode{afii10101}{00A0}
\pdfglyphtounicode{afii10102}{00A0}
\pdfglyphtounicode{afii10103}{00A0}
\pdfglyphtounicode{afii10104}{00A0}
\pdfglyphtounicode{afii10105}{00A0}
\pdfglyphtounicode{afii10106}{00A0}
\pdfglyphtounicode{afii10107}{00A0}
\pdfglyphtounicode{afii10108}{00A0}
\pdfglyphtounicode{afii10109}{00A0}
\pdfglyphtounicode{afii10110}{00A0}
\pdfglyphtounicode{afii10145}{00A0}
\pdfglyphtounicode{afii10146}{00A0}
\pdfglyphtounicode{afii10147}{00A0}
\pdfglyphtounicode{afii10148}{00A0}
\pdfglyphtounicode{afii10192}{00A0}
\pdfglyphtounicode{afii10193}{00A0}
\pdfglyphtounicode{afii10194}{00A0}
\pdfglyphtounicode{afii10195}{00A0}
\pdfglyphtounicode{afii10196}{00A0}
\pdfglyphtounicode{afii10831}{00A0}
\pdfglyphtounicode{afii10832}{00A0}
\pdfglyphtounicode{afii10846}{00A0}
\pdfglyphtounicode{afii299}{00A0}
\pdfglyphtounicode{afii300}{00A0}
\pdfglyphtounicode{afii301}{00A0}
\pdfglyphtounicode{afii57381}{00A0}
\pdfglyphtounicode{afii57388}{00A0}
\pdfglyphtounicode{afii57392}{00A0}
\pdfglyphtounicode{afii57393}{00A0}
\pdfglyphtounicode{afii57394}{00A0}
\pdfglyphtounicode{afii57395}{00A0}
\pdfglyphtounicode{afii57396}{00A0}
\pdfglyphtounicode{afii57397}{00A0}
\pdfglyphtounicode{afii57398}{00A0}
\pdfglyphtounicode{afii57399}{00A0}
\pdfglyphtounicode{afii57400}{00A0}
\pdfglyphtounicode{afii57401}{00A0}
\pdfglyphtounicode{afii57403}{00A0}
\pdfglyphtounicode{afii57407}{00A0}
\pdfglyphtounicode{afii57409}{00A0}
\pdfglyphtounicode{afii57410}{00A0}
\pdfglyphtounicode{afii57411}{00A0}
\pdfglyphtounicode{afii57412}{00A0}
\pdfglyphtounicode{afii57413}{00A0}
\pdfglyphtounicode{afii57414}{00A0}
\pdfglyphtounicode{afii57415}{00A0}
\pdfglyphtounicode{afii57416}{00A0}
\pdfglyphtounicode{afii57417}{00A0}
\pdfglyphtounicode{afii57418}{00A0}
\pdfglyphtounicode{afii57419}{00A0}
\pdfglyphtounicode{afii57420}{00A0}
\pdfglyphtounicode{afii57421}{00A0}
\pdfglyphtounicode{afii57422}{00A0}
\pdfglyphtounicode{afii57423}{00A0}
\pdfglyphtounicode{afii57424}{00A0}
\pdfglyphtounicode{afii57425}{00A0}
\pdfglyphtounicode{afii57426}{00A0}
\pdfglyphtounicode{afii57427}{00A0}
\pdfglyphtounicode{afii57428}{00A0}
\pdfglyphtounicode{afii57429}{00A0}
\pdfglyphtounicode{afii57430}{00A0}
\pdfglyphtounicode{afii57431}{00A0}
\pdfglyphtounicode{afii57432}{00A0}
\pdfglyphtounicode{afii57433}{00A0}
\pdfglyphtounicode{afii57434}{00A0}
\pdfglyphtounicode{afii57440}{00A0}
\pdfglyphtounicode{afii57441}{00A0}
\pdfglyphtounicode{afii57442}{00A0}
\pdfglyphtounicode{afii57443}{00A0}
\pdfglyphtounicode{afii57444}{00A0}
\pdfglyphtounicode{afii57445}{00A0}
\pdfglyphtounicode{afii57446}{00A0}
\pdfglyphtounicode{afii57448}{00A0}
\pdfglyphtounicode{afii57449}{00A0}
\pdfglyphtounicode{afii57450}{00A0}
\pdfglyphtounicode{afii57451}{00A0}
\pdfglyphtounicode{afii57452}{00A0}
\pdfglyphtounicode{afii57453}{00A0}
\pdfglyphtounicode{afii57454}{00A0}
\pdfglyphtounicode{afii57455}{00A0}
\pdfglyphtounicode{afii57456}{00A0}
\pdfglyphtounicode{afii57457}{00A0}
\pdfglyphtounicode{afii57458}{00A0}
\pdfglyphtounicode{afii57470}{00A0}
\pdfglyphtounicode{afii57505}{00A0}
\pdfglyphtounicode{afii57506}{00A0}
\pdfglyphtounicode{afii57507}{00A0}
\pdfglyphtounicode{afii57508}{00A0}
\pdfglyphtounicode{afii57509}{00A0}
\pdfglyphtounicode{afii57511}{00A0}
\pdfglyphtounicode{afii57512}{00A0}
\pdfglyphtounicode{afii57513}{00A0}
\pdfglyphtounicode{afii57514}{00A0}
\pdfglyphtounicode{afii57519}{00A0}
\pdfglyphtounicode{afii57534}{00A0}
\pdfglyphtounicode{afii57636}{00A0}
\pdfglyphtounicode{afii57645}{00A0}
\pdfglyphtounicode{afii57658}{00A0}
\pdfglyphtounicode{afii57664}{00A0}
\pdfglyphtounicode{afii57665}{00A0}
\pdfglyphtounicode{afii57666}{00A0}
\pdfglyphtounicode{afii57667}{00A0}
\pdfglyphtounicode{afii57668}{00A0}
\pdfglyphtounicode{afii57669}{00A0}
\pdfglyphtounicode{afii57670}{00A0}
\pdfglyphtounicode{afii57671}{00A0}
\pdfglyphtounicode{afii57672}{00A0}
\pdfglyphtounicode{afii57673}{00A0}
\pdfglyphtounicode{afii57674}{00A0}
\pdfglyphtounicode{afii57675}{00A0}
\pdfglyphtounicode{afii57676}{00A0}
\pdfglyphtounicode{afii57677}{00A0}
\pdfglyphtounicode{afii57678}{00A0}
\pdfglyphtounicode{afii57679}{00A0}
\pdfglyphtounicode{afii57680}{00A0}
\pdfglyphtounicode{afii57681}{00A0}
\pdfglyphtounicode{afii57682}{00A0}
\pdfglyphtounicode{afii57683}{00A0}
\pdfglyphtounicode{afii57684}{00A0}
\pdfglyphtounicode{afii57685}{00A0}
\pdfglyphtounicode{afii57686}{00A0}
\pdfglyphtounicode{afii57687}{00A0}
\pdfglyphtounicode{afii57688}{00A0}
\pdfglyphtounicode{afii57689}{00A0}
\pdfglyphtounicode{afii57690}{00A0}
\pdfglyphtounicode{afii57694}{00A0}
\pdfglyphtounicode{afii57695}{00A0}
\pdfglyphtounicode{afii57700}{00A0}
\pdfglyphtounicode{afii57705}{00A0}
\pdfglyphtounicode{afii57716}{00A0}
\pdfglyphtounicode{afii57717}{00A0}
\pdfglyphtounicode{afii57718}{00A0}
\pdfglyphtounicode{afii57723}{00A0}
\pdfglyphtounicode{afii57793}{00A0}
\pdfglyphtounicode{afii57794}{00A0}
\pdfglyphtounicode{afii57795}{00A0}
\pdfglyphtounicode{afii57796}{00A0}
\pdfglyphtounicode{afii57797}{00A0}
\pdfglyphtounicode{afii57798}{00A0}
\pdfglyphtounicode{afii57799}{00A0}
\pdfglyphtounicode{afii57800}{00A0}
\pdfglyphtounicode{afii57801}{00A0}
\pdfglyphtounicode{afii57802}{00A0}
\pdfglyphtounicode{afii57803}{00A0}
\pdfglyphtounicode{afii57804}{00A0}
\pdfglyphtounicode{afii57806}{00A0}
\pdfglyphtounicode{afii57807}{00A0}
\pdfglyphtounicode{afii57839}{00A0}
\pdfglyphtounicode{afii57841}{00A0}
\pdfglyphtounicode{afii57842}{00A0}
\pdfglyphtounicode{afii57929}{00A0}
\pdfglyphtounicode{afii61248}{00A0}
\pdfglyphtounicode{afii61289}{00A0}
\pdfglyphtounicode{afii61352}{00A0}
\pdfglyphtounicode{afii61573}{00A0}
\pdfglyphtounicode{afii61574}{00A0}
\pdfglyphtounicode{afii61575}{00A0}
\pdfglyphtounicode{afii61664}{00A0}
\pdfglyphtounicode{afii63167}{00A0}
\pdfglyphtounicode{afii64937}{00A0}
\pdfglyphtounicode{agrave}{00A0}
\pdfglyphtounicode{agujarati}{00A0}
\pdfglyphtounicode{agurmukhi}{00A0}
\pdfglyphtounicode{ahiragana}{00A0}
\pdfglyphtounicode{ahookabove}{00A0}
\pdfglyphtounicode{aibengali}{00A0}
\pdfglyphtounicode{aibopomofo}{00A0}
\pdfglyphtounicode{aideva}{00A0}
\pdfglyphtounicode{aiecyrillic}{00A0}
\pdfglyphtounicode{aigujarati}{00A0}
\pdfglyphtounicode{aigurmukhi}{00A0}
\pdfglyphtounicode{aimatragurmukhi}{00A0}
\pdfglyphtounicode{ainarabic}{00A0}
\pdfglyphtounicode{ainfinalarabic}{00A0}
\pdfglyphtounicode{aininitialarabic}{00A0}
\pdfglyphtounicode{ainmedialarabic}{00A0}
\pdfglyphtounicode{ainvertedbreve}{00A0}
\pdfglyphtounicode{aivowelsignbengali}{00A0}
\pdfglyphtounicode{aivowelsigndeva}{00A0}
\pdfglyphtounicode{aivowelsigngujarati}{00A0}
\pdfglyphtounicode{akatakana}{00A0}
\pdfglyphtounicode{akatakanahalfwidth}{00A0}
\pdfglyphtounicode{akorean}{00A0}
\pdfglyphtounicode{alef}{00A0}
\pdfglyphtounicode{alefarabic}{00A0}
\pdfglyphtounicode{alefdageshhebrew}{00A0}
\pdfglyphtounicode{aleffinalarabic}{00A0}
\pdfglyphtounicode{alefhamzaabovearabic}{00A0}
\pdfglyphtounicode{alefhamzaabovefinalarabic}{00A0}
\pdfglyphtounicode{alefhamzabelowarabic}{00A0}
\pdfglyphtounicode{alefhamzabelowfinalarabic}{00A0}
\pdfglyphtounicode{alefhebrew}{00A0}
\pdfglyphtounicode{aleflamedhebrew}{00A0}
\pdfglyphtounicode{alefmaddaabovearabic}{00A0}
\pdfglyphtounicode{alefmaddaabovefinalarabic}{00A0}
\pdfglyphtounicode{alefmaksuraarabic}{00A0}
\pdfglyphtounicode{alefmaksurafinalarabic}{00A0}
\pdfglyphtounicode{alefmaksurainitialarabic}{00A0}
\pdfglyphtounicode{alefmaksuramedialarabic}{00A0}
\pdfglyphtounicode{alefpatahhebrew}{00A0}
\pdfglyphtounicode{alefqamatshebrew}{00A0}
\pdfglyphtounicode{aleph}{00A0}
\pdfglyphtounicode{allequal}{00A0}
\pdfglyphtounicode{alpha}{00A0}
\pdfglyphtounicode{alphatonos}{00A0}
\pdfglyphtounicode{amacron}{00A0}
\pdfglyphtounicode{amonospace}{00A0}
\pdfglyphtounicode{ampersand}{00A0}
\pdfglyphtounicode{ampersandmonospace}{00A0}
\pdfglyphtounicode{ampersandsmall}{00A0}
\pdfglyphtounicode{amsquare}{00A0}
\pdfglyphtounicode{anbopomofo}{00A0}
\pdfglyphtounicode{angbopomofo}{00A0}
\pdfglyphtounicode{angbracketleft}{00A0}
\pdfglyphtounicode{angbracketright}{00A0}
\pdfglyphtounicode{angkhankhuthai}{00A0}
\pdfglyphtounicode{angle}{00A0}
\pdfglyphtounicode{anglebracketleft}{00A0}
\pdfglyphtounicode{anglebracketleftvertical}{00A0}
\pdfglyphtounicode{anglebracketright}{00A0}
\pdfglyphtounicode{anglebracketrightvertical}{00A0}
\pdfglyphtounicode{angleleft}{00A0}
\pdfglyphtounicode{angleright}{00A0}
\pdfglyphtounicode{angstrom}{00A0}
\pdfglyphtounicode{anoteleia}{00A0}
\pdfglyphtounicode{anticlockwise}{00A0}
\pdfglyphtounicode{anudattadeva}{00A0}
\pdfglyphtounicode{anusvarabengali}{00A0}
\pdfglyphtounicode{anusvaradeva}{00A0}
\pdfglyphtounicode{anusvaragujarati}{00A0}
\pdfglyphtounicode{aogonek}{00A0}
\pdfglyphtounicode{apaatosquare}{00A0}
\pdfglyphtounicode{aparen}{00A0}
\pdfglyphtounicode{apostrophearmenian}{00A0}
\pdfglyphtounicode{apostrophemod}{00A0}
\pdfglyphtounicode{apple}{00A0}
\pdfglyphtounicode{approaches}{00A0}
\pdfglyphtounicode{approxequal}{00A0}
\pdfglyphtounicode{approxequalorimage}{00A0}
\pdfglyphtounicode{approximatelyequal}{00A0}
\pdfglyphtounicode{approxorequal}{00A0}
\pdfglyphtounicode{araeaekorean}{00A0}
\pdfglyphtounicode{araeakorean}{00A0}
\pdfglyphtounicode{arc}{00A0}
\pdfglyphtounicode{archleftdown}{00A0}
\pdfglyphtounicode{archrightdown}{00A0}
\pdfglyphtounicode{arighthalfring}{00A0}
\pdfglyphtounicode{aring}{00A0}
\pdfglyphtounicode{aringacute}{00A0}
\pdfglyphtounicode{aringbelow}{00A0}
\pdfglyphtounicode{arrowboth}{00A0}
\pdfglyphtounicode{arrowbothv}{00A0}
\pdfglyphtounicode{arrowdashdown}{00A0}
\pdfglyphtounicode{arrowdashleft}{00A0}
\pdfglyphtounicode{arrowdashright}{00A0}
\pdfglyphtounicode{arrowdashup}{00A0}
\pdfglyphtounicode{arrowdblboth}{00A0}
\pdfglyphtounicode{arrowdblbothv}{00A0}
\pdfglyphtounicode{arrowdbldown}{00A0}
\pdfglyphtounicode{arrowdblleft}{00A0}
\pdfglyphtounicode{arrowdblright}{00A0}
\pdfglyphtounicode{arrowdblup}{00A0}
\pdfglyphtounicode{arrowdown}{00A0}
\pdfglyphtounicode{arrowdownleft}{00A0}
\pdfglyphtounicode{arrowdownright}{00A0}
\pdfglyphtounicode{arrowdownwhite}{00A0}
\pdfglyphtounicode{arrowheaddownmod}{00A0}
\pdfglyphtounicode{arrowheadleftmod}{00A0}
\pdfglyphtounicode{arrowheadrightmod}{00A0}
\pdfglyphtounicode{arrowheadupmod}{00A0}
\pdfglyphtounicode{arrowhorizex}{00A0}
\pdfglyphtounicode{arrowleft}{00A0}
\pdfglyphtounicode{arrowleftbothalf}{00A0}
\pdfglyphtounicode{arrowleftdbl}{00A0}
\pdfglyphtounicode{arrowleftdblstroke}{00A0}
\pdfglyphtounicode{arrowleftoverright}{00A0}
\pdfglyphtounicode{arrowlefttophalf}{00A0}
\pdfglyphtounicode{arrowleftwhite}{00A0}
\pdfglyphtounicode{arrownortheast}{00A0}
\pdfglyphtounicode{arrownorthwest}{00A0}
\pdfglyphtounicode{arrowparrleftright}{00A0}
\pdfglyphtounicode{arrowparrrightleft}{00A0}
\pdfglyphtounicode{arrowright}{00A0}
\pdfglyphtounicode{arrowrightbothalf}{00A0}
\pdfglyphtounicode{arrowrightdblstroke}{00A0}
\pdfglyphtounicode{arrowrightheavy}{00A0}
\pdfglyphtounicode{arrowrightoverleft}{00A0}
\pdfglyphtounicode{arrowrighttophalf}{00A0}
\pdfglyphtounicode{arrowrightwhite}{00A0}
\pdfglyphtounicode{arrowsoutheast}{00A0}
\pdfglyphtounicode{arrowsouthwest}{00A0}
\pdfglyphtounicode{arrowtableft}{00A0}
\pdfglyphtounicode{arrowtabright}{00A0}
\pdfglyphtounicode{arrowtailleft}{00A0}
\pdfglyphtounicode{arrowtailright}{00A0}
\pdfglyphtounicode{arrowtripleleft}{00A0}
\pdfglyphtounicode{arrowtripleright}{00A0}
\pdfglyphtounicode{arrowup}{00A0}
\pdfglyphtounicode{arrowupdn}{00A0}
\pdfglyphtounicode{arrowupdnbse}{00A0}
\pdfglyphtounicode{arrowupdownbase}{00A0}
\pdfglyphtounicode{arrowupleft}{00A0}
\pdfglyphtounicode{arrowupleftofdown}{00A0}
\pdfglyphtounicode{arrowupright}{00A0}
\pdfglyphtounicode{arrowupwhite}{00A0}
\pdfglyphtounicode{arrowvertex}{00A0}
\pdfglyphtounicode{asciicircum}{00A0}
\pdfglyphtounicode{asciicircummonospace}{00A0}
\pdfglyphtounicode{asciitilde}{00A0}
\pdfglyphtounicode{asciitildemonospace}{00A0}
\pdfglyphtounicode{ascript}{00A0}
\pdfglyphtounicode{ascriptturned}{00A0}
\pdfglyphtounicode{asmallhiragana}{00A0}
\pdfglyphtounicode{asmallkatakana}{00A0}
\pdfglyphtounicode{asmallkatakanahalfwidth}{00A0}
\pdfglyphtounicode{asterisk}{00A0}
\pdfglyphtounicode{asteriskaltonearabic}{00A0}
\pdfglyphtounicode{asteriskarabic}{00A0}
\pdfglyphtounicode{asteriskcentered}{00A0}
\pdfglyphtounicode{asteriskmath}{00A0}
\pdfglyphtounicode{asteriskmonospace}{00A0}
\pdfglyphtounicode{asterisksmall}{00A0}
\pdfglyphtounicode{asterism}{00A0}
\pdfglyphtounicode{asuperior}{00A0}
\pdfglyphtounicode{asymptoticallyequal}{00A0}
\pdfglyphtounicode{at}{00A0}
\pdfglyphtounicode{atilde}{00A0}
\pdfglyphtounicode{atmonospace}{00A0}
\pdfglyphtounicode{atsmall}{00A0}
\pdfglyphtounicode{aturned}{00A0}
\pdfglyphtounicode{aubengali}{00A0}
\pdfglyphtounicode{aubopomofo}{00A0}
\pdfglyphtounicode{audeva}{00A0}
\pdfglyphtounicode{augujarati}{00A0}
\pdfglyphtounicode{augurmukhi}{00A0}
\pdfglyphtounicode{aulengthmarkbengali}{00A0}
\pdfglyphtounicode{aumatragurmukhi}{00A0}
\pdfglyphtounicode{auvowelsignbengali}{00A0}
\pdfglyphtounicode{auvowelsigndeva}{00A0}
\pdfglyphtounicode{auvowelsigngujarati}{00A0}
\pdfglyphtounicode{avagrahadeva}{00A0}
\pdfglyphtounicode{aybarmenian}{00A0}
\pdfglyphtounicode{ayin}{00A0}
\pdfglyphtounicode{ayinaltonehebrew}{00A0}
\pdfglyphtounicode{ayinhebrew}{00A0}
\pdfglyphtounicode{b}{00A0}
\pdfglyphtounicode{babengali}{00A0}
\pdfglyphtounicode{backslash}{00A0}
\pdfglyphtounicode{backslashmonospace}{00A0}
\pdfglyphtounicode{badeva}{00A0}
\pdfglyphtounicode{bagujarati}{00A0}
\pdfglyphtounicode{bagurmukhi}{00A0}
\pdfglyphtounicode{bahiragana}{00A0}
\pdfglyphtounicode{bahtthai}{00A0}
\pdfglyphtounicode{bakatakana}{00A0}
\pdfglyphtounicode{bar}{00A0}
\pdfglyphtounicode{bardbl}{00A0}
\pdfglyphtounicode{barmonospace}{00A0}
\pdfglyphtounicode{bbopomofo}{00A0}
\pdfglyphtounicode{bcircle}{00A0}
\pdfglyphtounicode{bdotaccent}{00A0}
\pdfglyphtounicode{bdotbelow}{00A0}
\pdfglyphtounicode{beamedsixteenthnotes}{00A0}
\pdfglyphtounicode{because}{00A0}
\pdfglyphtounicode{becyrillic}{00A0}
\pdfglyphtounicode{beharabic}{00A0}
\pdfglyphtounicode{behfinalarabic}{00A0}
\pdfglyphtounicode{behinitialarabic}{00A0}
\pdfglyphtounicode{behiragana}{00A0}
\pdfglyphtounicode{behmedialarabic}{00A0}
\pdfglyphtounicode{behmeeminitialarabic}{00A0}
\pdfglyphtounicode{behmeemisolatedarabic}{00A0}
\pdfglyphtounicode{behnoonfinalarabic}{00A0}
\pdfglyphtounicode{bekatakana}{00A0}
\pdfglyphtounicode{benarmenian}{00A0}
\pdfglyphtounicode{bet}{00A0}
\pdfglyphtounicode{beta}{00A0}
\pdfglyphtounicode{betasymbolgreek}{00A0}
\pdfglyphtounicode{betdagesh}{00A0}
\pdfglyphtounicode{betdageshhebrew}{00A0}
\pdfglyphtounicode{beth}{00A0}
\pdfglyphtounicode{bethebrew}{00A0}
\pdfglyphtounicode{betrafehebrew}{00A0}
\pdfglyphtounicode{between}{00A0}
\pdfglyphtounicode{bhabengali}{00A0}
\pdfglyphtounicode{bhadeva}{00A0}
\pdfglyphtounicode{bhagujarati}{00A0}
\pdfglyphtounicode{bhagurmukhi}{00A0}
\pdfglyphtounicode{bhook}{00A0}
\pdfglyphtounicode{bihiragana}{00A0}
\pdfglyphtounicode{bikatakana}{00A0}
\pdfglyphtounicode{bilabialclick}{00A0}
\pdfglyphtounicode{bindigurmukhi}{00A0}
\pdfglyphtounicode{birusquare}{00A0}
\pdfglyphtounicode{blackcircle}{00A0}
\pdfglyphtounicode{blackdiamond}{00A0}
\pdfglyphtounicode{blackdownpointingtriangle}{00A0}
\pdfglyphtounicode{blackleftpointingpointer}{00A0}
\pdfglyphtounicode{blackleftpointingtriangle}{00A0}
\pdfglyphtounicode{blacklenticularbracketleft}{00A0}
\pdfglyphtounicode{blacklenticularbracketleftvertical}{00A0}
\pdfglyphtounicode{blacklenticularbracketright}{00A0}
\pdfglyphtounicode{blacklenticularbracketrightvertical}{00A0}
\pdfglyphtounicode{blacklowerlefttriangle}{00A0}
\pdfglyphtounicode{blacklowerrighttriangle}{00A0}
\pdfglyphtounicode{blackrectangle}{00A0}
\pdfglyphtounicode{blackrightpointingpointer}{00A0}
\pdfglyphtounicode{blackrightpointingtriangle}{00A0}
\pdfglyphtounicode{blacksmallsquare}{00A0}
\pdfglyphtounicode{blacksmilingface}{00A0}
\pdfglyphtounicode{blacksquare}{00A0}
\pdfglyphtounicode{blackstar}{00A0}
\pdfglyphtounicode{blackupperlefttriangle}{00A0}
\pdfglyphtounicode{blackupperrighttriangle}{00A0}
\pdfglyphtounicode{blackuppointingsmalltriangle}{00A0}
\pdfglyphtounicode{blackuppointingtriangle}{00A0}
\pdfglyphtounicode{blank}{00A0}
\pdfglyphtounicode{blinebelow}{00A0}
\pdfglyphtounicode{block}{00A0}
\pdfglyphtounicode{bmonospace}{00A0}
\pdfglyphtounicode{bobaimaithai}{00A0}
\pdfglyphtounicode{bohiragana}{00A0}
\pdfglyphtounicode{bokatakana}{00A0}
\pdfglyphtounicode{bparen}{00A0}
\pdfglyphtounicode{bqsquare}{00A0}
\pdfglyphtounicode{braceex}{00A0}
\pdfglyphtounicode{braceleft}{00A0}
\pdfglyphtounicode{braceleftbt}{00A0}
\pdfglyphtounicode{braceleftmid}{00A0}
\pdfglyphtounicode{braceleftmonospace}{00A0}
\pdfglyphtounicode{braceleftsmall}{00A0}
\pdfglyphtounicode{bracelefttp}{00A0}
\pdfglyphtounicode{braceleftvertical}{00A0}
\pdfglyphtounicode{braceright}{00A0}
\pdfglyphtounicode{bracerightbt}{00A0}
\pdfglyphtounicode{bracerightmid}{00A0}
\pdfglyphtounicode{bracerightmonospace}{00A0}
\pdfglyphtounicode{bracerightsmall}{00A0}
\pdfglyphtounicode{bracerighttp}{00A0}
\pdfglyphtounicode{bracerightvertical}{00A0}
\pdfglyphtounicode{bracketleft}{00A0}
\pdfglyphtounicode{bracketleftbt}{00A0}
\pdfglyphtounicode{bracketleftex}{00A0}
\pdfglyphtounicode{bracketleftmonospace}{00A0}
\pdfglyphtounicode{bracketlefttp}{00A0}
\pdfglyphtounicode{bracketright}{00A0}
\pdfglyphtounicode{bracketrightbt}{00A0}
\pdfglyphtounicode{bracketrightex}{00A0}
\pdfglyphtounicode{bracketrightmonospace}{00A0}
\pdfglyphtounicode{bracketrighttp}{00A0}
\pdfglyphtounicode{breve}{00A0}
\pdfglyphtounicode{brevebelowcmb}{00A0}
\pdfglyphtounicode{brevecmb}{00A0}
\pdfglyphtounicode{breveinvertedbelowcmb}{00A0}
\pdfglyphtounicode{breveinvertedcmb}{00A0}
\pdfglyphtounicode{breveinverteddoublecmb}{00A0}
\pdfglyphtounicode{bridgebelowcmb}{00A0}
\pdfglyphtounicode{bridgeinvertedbelowcmb}{00A0}
\pdfglyphtounicode{brokenbar}{00A0}
\pdfglyphtounicode{bstroke}{00A0}
\pdfglyphtounicode{bsuperior}{00A0}
\pdfglyphtounicode{btopbar}{00A0}
\pdfglyphtounicode{buhiragana}{00A0}
\pdfglyphtounicode{bukatakana}{00A0}
\pdfglyphtounicode{bullet}{00A0}
\pdfglyphtounicode{bulletinverse}{00A0}
\pdfglyphtounicode{bulletoperator}{00A0}
\pdfglyphtounicode{bullseye}{00A0}
\pdfglyphtounicode{c}{00A0}
\pdfglyphtounicode{caarmenian}{00A0}
\pdfglyphtounicode{cabengali}{00A0}
\pdfglyphtounicode{cacute}{00A0}
\pdfglyphtounicode{cadeva}{00A0}
\pdfglyphtounicode{cagujarati}{00A0}
\pdfglyphtounicode{cagurmukhi}{00A0}
\pdfglyphtounicode{calsquare}{00A0}
\pdfglyphtounicode{candrabindubengali}{00A0}
\pdfglyphtounicode{candrabinducmb}{00A0}
\pdfglyphtounicode{candrabindudeva}{00A0}
\pdfglyphtounicode{candrabindugujarati}{00A0}
\pdfglyphtounicode{capslock}{00A0}
\pdfglyphtounicode{careof}{00A0}
\pdfglyphtounicode{caron}{00A0}
\pdfglyphtounicode{caronbelowcmb}{00A0}
\pdfglyphtounicode{caroncmb}{00A0}
\pdfglyphtounicode{carriagereturn}{00A0}
\pdfglyphtounicode{cbopomofo}{00A0}
\pdfglyphtounicode{ccaron}{00A0}
\pdfglyphtounicode{ccedilla}{00A0}
\pdfglyphtounicode{ccedillaacute}{00A0}
\pdfglyphtounicode{ccircle}{00A0}
\pdfglyphtounicode{ccircumflex}{00A0}
\pdfglyphtounicode{ccurl}{00A0}
\pdfglyphtounicode{cdot}{00A0}
\pdfglyphtounicode{cdotaccent}{00A0}
\pdfglyphtounicode{cdsquare}{00A0}
\pdfglyphtounicode{cedilla}{00A0}
\pdfglyphtounicode{cedillacmb}{00A0}
\pdfglyphtounicode{ceilingleft}{00A0}
\pdfglyphtounicode{ceilingright}{00A0}
\pdfglyphtounicode{cent}{00A0}
\pdfglyphtounicode{centigrade}{00A0}
\pdfglyphtounicode{centinferior}{00A0}
\pdfglyphtounicode{centmonospace}{00A0}
\pdfglyphtounicode{centoldstyle}{00A0}
\pdfglyphtounicode{centsuperior}{00A0}
\pdfglyphtounicode{chaarmenian}{00A0}
\pdfglyphtounicode{chabengali}{00A0}
\pdfglyphtounicode{chadeva}{00A0}
\pdfglyphtounicode{chagujarati}{00A0}
\pdfglyphtounicode{chagurmukhi}{00A0}
\pdfglyphtounicode{chbopomofo}{00A0}
\pdfglyphtounicode{cheabkhasiancyrillic}{00A0}
\pdfglyphtounicode{check}{00A0}
\pdfglyphtounicode{checkmark}{00A0}
\pdfglyphtounicode{checyrillic}{00A0}
\pdfglyphtounicode{chedescenderabkhasiancyrillic}{00A0}
\pdfglyphtounicode{chedescendercyrillic}{00A0}
\pdfglyphtounicode{chedieresiscyrillic}{00A0}
\pdfglyphtounicode{cheharmenian}{00A0}
\pdfglyphtounicode{chekhakassiancyrillic}{00A0}
\pdfglyphtounicode{cheverticalstrokecyrillic}{00A0}
\pdfglyphtounicode{chi}{00A0}
\pdfglyphtounicode{chieuchacirclekorean}{00A0}
\pdfglyphtounicode{chieuchaparenkorean}{00A0}
\pdfglyphtounicode{chieuchcirclekorean}{00A0}
\pdfglyphtounicode{chieuchkorean}{00A0}
\pdfglyphtounicode{chieuchparenkorean}{00A0}
\pdfglyphtounicode{chochangthai}{00A0}
\pdfglyphtounicode{chochanthai}{00A0}
\pdfglyphtounicode{chochingthai}{00A0}
\pdfglyphtounicode{chochoethai}{00A0}
\pdfglyphtounicode{chook}{00A0}
\pdfglyphtounicode{cieucacirclekorean}{00A0}
\pdfglyphtounicode{cieucaparenkorean}{00A0}
\pdfglyphtounicode{cieuccirclekorean}{00A0}
\pdfglyphtounicode{cieuckorean}{00A0}
\pdfglyphtounicode{cieucparenkorean}{00A0}
\pdfglyphtounicode{cieucuparenkorean}{00A0}
\pdfglyphtounicode{circle}{00A0}
\pdfglyphtounicode{circleR}{00A0}
\pdfglyphtounicode{circleS}{00A0}
\pdfglyphtounicode{circleasterisk}{00A0}
\pdfglyphtounicode{circlecopyrt}{00A0}
\pdfglyphtounicode{circledivide}{00A0}
\pdfglyphtounicode{circledot}{00A0}
\pdfglyphtounicode{circleequal}{00A0}
\pdfglyphtounicode{circleminus}{00A0}
\pdfglyphtounicode{circlemultiply}{00A0}
\pdfglyphtounicode{circleot}{00A0}
\pdfglyphtounicode{circleplus}{00A0}
\pdfglyphtounicode{circlepostalmark}{00A0}
\pdfglyphtounicode{circlering}{00A0}
\pdfglyphtounicode{circlewithlefthalfblack}{00A0}
\pdfglyphtounicode{circlewithrighthalfblack}{00A0}
\pdfglyphtounicode{circumflex}{00A0}
\pdfglyphtounicode{circumflexbelowcmb}{00A0}
\pdfglyphtounicode{circumflexcmb}{00A0}
\pdfglyphtounicode{clear}{00A0}
\pdfglyphtounicode{clickalveolar}{00A0}
\pdfglyphtounicode{clickdental}{00A0}
\pdfglyphtounicode{clicklateral}{00A0}
\pdfglyphtounicode{clickretroflex}{00A0}
\pdfglyphtounicode{clockwise}{00A0}
\pdfglyphtounicode{club}{00A0}
\pdfglyphtounicode{clubsuitblack}{00A0}
\pdfglyphtounicode{clubsuitwhite}{00A0}
\pdfglyphtounicode{cmcubedsquare}{00A0}
\pdfglyphtounicode{cmonospace}{00A0}
\pdfglyphtounicode{cmsquaredsquare}{00A0}
\pdfglyphtounicode{coarmenian}{00A0}
\pdfglyphtounicode{colon}{00A0}
\pdfglyphtounicode{colonmonetary}{00A0}
\pdfglyphtounicode{colonmonospace}{00A0}
\pdfglyphtounicode{colonsign}{00A0}
\pdfglyphtounicode{colonsmall}{00A0}
\pdfglyphtounicode{colontriangularhalfmod}{00A0}
\pdfglyphtounicode{colontriangularmod}{00A0}
\pdfglyphtounicode{comma}{00A0}
\pdfglyphtounicode{commaabovecmb}{00A0}
\pdfglyphtounicode{commaaboverightcmb}{00A0}
\pdfglyphtounicode{commaaccent}{00A0}
\pdfglyphtounicode{commaarabic}{00A0}
\pdfglyphtounicode{commaarmenian}{00A0}
\pdfglyphtounicode{commainferior}{00A0}
\pdfglyphtounicode{commamonospace}{00A0}
\pdfglyphtounicode{commareversedabovecmb}{00A0}
\pdfglyphtounicode{commareversedmod}{00A0}
\pdfglyphtounicode{commasmall}{00A0}
\pdfglyphtounicode{commasuperior}{00A0}
\pdfglyphtounicode{commaturnedabovecmb}{00A0}
\pdfglyphtounicode{commaturnedmod}{00A0}
\pdfglyphtounicode{compass}{00A0}
\pdfglyphtounicode{complement}{00A0}
\pdfglyphtounicode{compwordmark}{00A0}
\pdfglyphtounicode{congruent}{00A0}
\pdfglyphtounicode{contourintegral}{00A0}
\pdfglyphtounicode{control}{00A0}
\pdfglyphtounicode{controlACK}{00A0}
\pdfglyphtounicode{controlBEL}{00A0}
\pdfglyphtounicode{controlBS}{00A0}
\pdfglyphtounicode{controlCAN}{00A0}
\pdfglyphtounicode{controlCR}{00A0}
\pdfglyphtounicode{controlDC1}{00A0}
\pdfglyphtounicode{controlDC2}{00A0}
\pdfglyphtounicode{controlDC3}{00A0}
\pdfglyphtounicode{controlDC4}{00A0}
\pdfglyphtounicode{controlDEL}{00A0}
\pdfglyphtounicode{controlDLE}{00A0}
\pdfglyphtounicode{controlEM}{00A0}
\pdfglyphtounicode{controlENQ}{00A0}
\pdfglyphtounicode{controlEOT}{00A0}
\pdfglyphtounicode{controlESC}{00A0}
\pdfglyphtounicode{controlETB}{00A0}
\pdfglyphtounicode{controlETX}{00A0}
\pdfglyphtounicode{controlFF}{00A0}
\pdfglyphtounicode{controlFS}{00A0}
\pdfglyphtounicode{controlGS}{00A0}
\pdfglyphtounicode{controlHT}{00A0}
\pdfglyphtounicode{controlLF}{00A0}
\pdfglyphtounicode{controlNAK}{00A0}
\pdfglyphtounicode{controlRS}{00A0}
\pdfglyphtounicode{controlSI}{00A0}
\pdfglyphtounicode{controlSO}{00A0}
\pdfglyphtounicode{controlSOT}{00A0}
\pdfglyphtounicode{controlSTX}{00A0}
\pdfglyphtounicode{controlSUB}{00A0}
\pdfglyphtounicode{controlSYN}{00A0}
\pdfglyphtounicode{controlUS}{00A0}
\pdfglyphtounicode{controlVT}{00A0}
\pdfglyphtounicode{coproduct}{00A0}
\pdfglyphtounicode{copyright}{00A0}
\pdfglyphtounicode{copyrightsans}{00A0}
\pdfglyphtounicode{copyrightserif}{00A0}
\pdfglyphtounicode{cornerbracketleft}{00A0}
\pdfglyphtounicode{cornerbracketlefthalfwidth}{00A0}
\pdfglyphtounicode{cornerbracketleftvertical}{00A0}
\pdfglyphtounicode{cornerbracketright}{00A0}
\pdfglyphtounicode{cornerbracketrighthalfwidth}{00A0}
\pdfglyphtounicode{cornerbracketrightvertical}{00A0}
\pdfglyphtounicode{corporationsquare}{00A0}
\pdfglyphtounicode{cosquare}{00A0}
\pdfglyphtounicode{coverkgsquare}{00A0}
\pdfglyphtounicode{cparen}{00A0}
\pdfglyphtounicode{cruzeiro}{00A0}
\pdfglyphtounicode{cstretched}{00A0}
\pdfglyphtounicode{ct}{00A0}
\pdfglyphtounicode{curlyand}{00A0}
\pdfglyphtounicode{curlyleft}{00A0}
\pdfglyphtounicode{curlyor}{00A0}
\pdfglyphtounicode{curlyright}{00A0}
\pdfglyphtounicode{currency}{00A0}
\pdfglyphtounicode{cwm}{00A0}
\pdfglyphtounicode{cyrBreve}{00A0}
\pdfglyphtounicode{cyrFlex}{00A0}
\pdfglyphtounicode{cyrbreve}{00A0}
\pdfglyphtounicode{cyrflex}{00A0}
\pdfglyphtounicode{d}{00A0}
\pdfglyphtounicode{daarmenian}{00A0}
\pdfglyphtounicode{dabengali}{00A0}
\pdfglyphtounicode{dadarabic}{00A0}
\pdfglyphtounicode{dadeva}{00A0}
\pdfglyphtounicode{dadfinalarabic}{00A0}
\pdfglyphtounicode{dadinitialarabic}{00A0}
\pdfglyphtounicode{dadmedialarabic}{00A0}
\pdfglyphtounicode{dagesh}{00A0}
\pdfglyphtounicode{dageshhebrew}{00A0}
\pdfglyphtounicode{dagger}{00A0}
\pdfglyphtounicode{daggerdbl}{00A0}
\pdfglyphtounicode{dagujarati}{00A0}
\pdfglyphtounicode{dagurmukhi}{00A0}
\pdfglyphtounicode{dahiragana}{00A0}
\pdfglyphtounicode{dakatakana}{00A0}
\pdfglyphtounicode{dalarabic}{00A0}
\pdfglyphtounicode{dalet}{00A0}
\pdfglyphtounicode{daletdagesh}{00A0}
\pdfglyphtounicode{daletdageshhebrew}{00A0}
\pdfglyphtounicode{daleth}{00A0}
\pdfglyphtounicode{dalethatafpatah}{00A0}
\pdfglyphtounicode{dalethatafpatahhebrew}{00A0}
\pdfglyphtounicode{dalethatafsegol}{00A0}
\pdfglyphtounicode{dalethatafsegolhebrew}{00A0}
\pdfglyphtounicode{dalethebrew}{00A0}
\pdfglyphtounicode{dalethiriq}{00A0}
\pdfglyphtounicode{dalethiriqhebrew}{00A0}
\pdfglyphtounicode{daletholam}{00A0}
\pdfglyphtounicode{daletholamhebrew}{00A0}
\pdfglyphtounicode{daletpatah}{00A0}
\pdfglyphtounicode{daletpatahhebrew}{00A0}
\pdfglyphtounicode{daletqamats}{00A0}
\pdfglyphtounicode{daletqamatshebrew}{00A0}
\pdfglyphtounicode{daletqubuts}{00A0}
\pdfglyphtounicode{daletqubutshebrew}{00A0}
\pdfglyphtounicode{daletsegol}{00A0}
\pdfglyphtounicode{daletsegolhebrew}{00A0}
\pdfglyphtounicode{daletsheva}{00A0}
\pdfglyphtounicode{daletshevahebrew}{00A0}
\pdfglyphtounicode{dalettsere}{00A0}
\pdfglyphtounicode{dalettserehebrew}{00A0}
\pdfglyphtounicode{dalfinalarabic}{00A0}
\pdfglyphtounicode{dammaarabic}{00A0}
\pdfglyphtounicode{dammalowarabic}{00A0}
\pdfglyphtounicode{dammatanaltonearabic}{00A0}
\pdfglyphtounicode{dammatanarabic}{00A0}
\pdfglyphtounicode{danda}{00A0}
\pdfglyphtounicode{dargahebrew}{00A0}
\pdfglyphtounicode{dargalefthebrew}{00A0}
\pdfglyphtounicode{dasiapneumatacyrilliccmb}{00A0}
\pdfglyphtounicode{dbar}{00A0}
\pdfglyphtounicode{dblGrave}{00A0}
\pdfglyphtounicode{dblanglebracketleft}{00A0}
\pdfglyphtounicode{dblanglebracketleftvertical}{00A0}
\pdfglyphtounicode{dblanglebracketright}{00A0}
\pdfglyphtounicode{dblanglebracketrightvertical}{00A0}
\pdfglyphtounicode{dblarchinvertedbelowcmb}{00A0}
\pdfglyphtounicode{dblarrowdwn}{00A0}
\pdfglyphtounicode{dblarrowheadleft}{00A0}
\pdfglyphtounicode{dblarrowheadright}{00A0}
\pdfglyphtounicode{dblarrowleft}{00A0}
\pdfglyphtounicode{dblarrowright}{00A0}
\pdfglyphtounicode{dblarrowup}{00A0}
\pdfglyphtounicode{dblbracketleft}{00A0}
\pdfglyphtounicode{dblbracketright}{00A0}
\pdfglyphtounicode{dbldanda}{00A0}
\pdfglyphtounicode{dblgrave}{00A0}
\pdfglyphtounicode{dblgravecmb}{00A0}
\pdfglyphtounicode{dblintegral}{00A0}
\pdfglyphtounicode{dbllowline}{00A0}
\pdfglyphtounicode{dbllowlinecmb}{00A0}
\pdfglyphtounicode{dbloverlinecmb}{00A0}
\pdfglyphtounicode{dblprimemod}{00A0}
\pdfglyphtounicode{dblverticalbar}{00A0}
\pdfglyphtounicode{dblverticallineabovecmb}{00A0}
\pdfglyphtounicode{dbopomofo}{00A0}
\pdfglyphtounicode{dbsquare}{00A0}
\pdfglyphtounicode{dcaron}{00A0}
\pdfglyphtounicode{dcedilla}{00A0}
\pdfglyphtounicode{dcircle}{00A0}
\pdfglyphtounicode{dcircumflexbelow}{00A0}
\pdfglyphtounicode{dcroat}{00A0}
\pdfglyphtounicode{ddabengali}{00A0}
\pdfglyphtounicode{ddadeva}{00A0}
\pdfglyphtounicode{ddagujarati}{00A0}
\pdfglyphtounicode{ddagurmukhi}{00A0}
\pdfglyphtounicode{ddalarabic}{00A0}
\pdfglyphtounicode{ddalfinalarabic}{00A0}
\pdfglyphtounicode{dddhadeva}{00A0}
\pdfglyphtounicode{ddhabengali}{00A0}
\pdfglyphtounicode{ddhadeva}{00A0}
\pdfglyphtounicode{ddhagujarati}{00A0}
\pdfglyphtounicode{ddhagurmukhi}{00A0}
\pdfglyphtounicode{ddotaccent}{00A0}
\pdfglyphtounicode{ddotbelow}{00A0}
\pdfglyphtounicode{decimalseparatorarabic}{00A0}
\pdfglyphtounicode{decimalseparatorpersian}{00A0}
\pdfglyphtounicode{decyrillic}{00A0}
\pdfglyphtounicode{defines}{00A0}
\pdfglyphtounicode{degree}{00A0}
\pdfglyphtounicode{dehihebrew}{00A0}
\pdfglyphtounicode{dehiragana}{00A0}
\pdfglyphtounicode{deicoptic}{00A0}
\pdfglyphtounicode{dekatakana}{00A0}
\pdfglyphtounicode{deleteleft}{00A0}
\pdfglyphtounicode{deleteright}{00A0}
\pdfglyphtounicode{delta}{00A0}
\pdfglyphtounicode{deltaturned}{00A0}
\pdfglyphtounicode{denominatorminusonenumeratorbengali}{00A0}
\pdfglyphtounicode{dezh}{00A0}
\pdfglyphtounicode{dhabengali}{00A0}
\pdfglyphtounicode{dhadeva}{00A0}
\pdfglyphtounicode{dhagujarati}{00A0}
\pdfglyphtounicode{dhagurmukhi}{00A0}
\pdfglyphtounicode{dhook}{00A0}
\pdfglyphtounicode{dialytikatonos}{00A0}
\pdfglyphtounicode{dialytikatonoscmb}{00A0}
\pdfglyphtounicode{diamond}{00A0}
\pdfglyphtounicode{diamondmath}{00A0}
\pdfglyphtounicode{diamondsolid}{00A0}
\pdfglyphtounicode{diamondsuitwhite}{00A0}
\pdfglyphtounicode{dieresis}{00A0}
\pdfglyphtounicode{dieresisacute}{00A0}
\pdfglyphtounicode{dieresisbelowcmb}{00A0}
\pdfglyphtounicode{dieresiscmb}{00A0}
\pdfglyphtounicode{dieresisgrave}{00A0}
\pdfglyphtounicode{dieresistonos}{00A0}
\pdfglyphtounicode{difference}{00A0}
\pdfglyphtounicode{dihiragana}{00A0}
\pdfglyphtounicode{dikatakana}{00A0}
\pdfglyphtounicode{dittomark}{00A0}
\pdfglyphtounicode{divide}{00A0}
\pdfglyphtounicode{dividemultiply}{00A0}
\pdfglyphtounicode{divides}{00A0}
\pdfglyphtounicode{divisionslash}{00A0}
\pdfglyphtounicode{djecyrillic}{00A0}
\pdfglyphtounicode{dkshade}{00A0}
\pdfglyphtounicode{dlinebelow}{00A0}
\pdfglyphtounicode{dlsquare}{00A0}
\pdfglyphtounicode{dmacron}{00A0}
\pdfglyphtounicode{dmonospace}{00A0}
\pdfglyphtounicode{dnblock}{00A0}
\pdfglyphtounicode{dochadathai}{00A0}
\pdfglyphtounicode{dodekthai}{00A0}
\pdfglyphtounicode{dohiragana}{00A0}
\pdfglyphtounicode{dokatakana}{00A0}
\pdfglyphtounicode{dollar}{00A0}
\pdfglyphtounicode{dollarinferior}{00A0}
\pdfglyphtounicode{dollarmonospace}{00A0}
\pdfglyphtounicode{dollaroldstyle}{00A0}
\pdfglyphtounicode{dollarsmall}{00A0}
\pdfglyphtounicode{dollarsuperior}{00A0}
\pdfglyphtounicode{dong}{00A0}
\pdfglyphtounicode{dorusquare}{00A0}
\pdfglyphtounicode{dotaccent}{00A0}
\pdfglyphtounicode{dotaccentcmb}{00A0}
\pdfglyphtounicode{dotbelowcmb}{00A0}
\pdfglyphtounicode{dotbelowcomb}{00A0}
\pdfglyphtounicode{dotkatakana}{00A0}
\pdfglyphtounicode{dotlessi}{00A0}
\pdfglyphtounicode{dotlessj}{00A0}
\pdfglyphtounicode{dotlessjstrokehook}{00A0}
\pdfglyphtounicode{dotmath}{00A0}
\pdfglyphtounicode{dotplus}{00A0}
\pdfglyphtounicode{dottedcircle}{00A0}
\pdfglyphtounicode{doubleyodpatah}{00A0}
\pdfglyphtounicode{doubleyodpatahhebrew}{00A0}
\pdfglyphtounicode{downfall}{00A0}
\pdfglyphtounicode{downslope}{00A0}
\pdfglyphtounicode{downtackbelowcmb}{00A0}
\pdfglyphtounicode{downtackmod}{00A0}
\pdfglyphtounicode{dparen}{00A0}
\pdfglyphtounicode{dsuperior}{00A0}
\pdfglyphtounicode{dtail}{00A0}
\pdfglyphtounicode{dtopbar}{00A0}
\pdfglyphtounicode{duhiragana}{00A0}
\pdfglyphtounicode{dukatakana}{00A0}
\pdfglyphtounicode{dz}{00A0}
\pdfglyphtounicode{dzaltone}{00A0}
\pdfglyphtounicode{dzcaron}{00A0}
\pdfglyphtounicode{dzcurl}{00A0}
\pdfglyphtounicode{dzeabkhasiancyrillic}{00A0}
\pdfglyphtounicode{dzecyrillic}{00A0}
\pdfglyphtounicode{dzhecyrillic}{00A0}
\pdfglyphtounicode{e}{00A0}
\pdfglyphtounicode{eacute}{00A0}
\pdfglyphtounicode{earth}{00A0}
\pdfglyphtounicode{ebengali}{00A0}
\pdfglyphtounicode{ebopomofo}{00A0}
\pdfglyphtounicode{ebreve}{00A0}
\pdfglyphtounicode{ecandradeva}{00A0}
\pdfglyphtounicode{ecandragujarati}{00A0}
\pdfglyphtounicode{ecandravowelsigndeva}{00A0}
\pdfglyphtounicode{ecandravowelsigngujarati}{00A0}
\pdfglyphtounicode{ecaron}{00A0}
\pdfglyphtounicode{ecedillabreve}{00A0}
\pdfglyphtounicode{echarmenian}{00A0}
\pdfglyphtounicode{echyiwnarmenian}{00A0}
\pdfglyphtounicode{ecircle}{00A0}
\pdfglyphtounicode{ecircumflex}{00A0}
\pdfglyphtounicode{ecircumflexacute}{00A0}
\pdfglyphtounicode{ecircumflexbelow}{00A0}
\pdfglyphtounicode{ecircumflexdotbelow}{00A0}
\pdfglyphtounicode{ecircumflexgrave}{00A0}
\pdfglyphtounicode{ecircumflexhookabove}{00A0}
\pdfglyphtounicode{ecircumflextilde}{00A0}
\pdfglyphtounicode{ecyrillic}{00A0}
\pdfglyphtounicode{edblgrave}{00A0}
\pdfglyphtounicode{edeva}{00A0}
\pdfglyphtounicode{edieresis}{00A0}
\pdfglyphtounicode{edot}{00A0}
\pdfglyphtounicode{edotaccent}{00A0}
\pdfglyphtounicode{edotbelow}{00A0}
\pdfglyphtounicode{eegurmukhi}{00A0}
\pdfglyphtounicode{eematragurmukhi}{00A0}
\pdfglyphtounicode{efcyrillic}{00A0}
\pdfglyphtounicode{egrave}{00A0}
\pdfglyphtounicode{egujarati}{00A0}
\pdfglyphtounicode{eharmenian}{00A0}
\pdfglyphtounicode{ehbopomofo}{00A0}
\pdfglyphtounicode{ehiragana}{00A0}
\pdfglyphtounicode{ehookabove}{00A0}
\pdfglyphtounicode{eibopomofo}{00A0}
\pdfglyphtounicode{eight}{00A0}
\pdfglyphtounicode{eightarabic}{00A0}
\pdfglyphtounicode{eightbengali}{00A0}
\pdfglyphtounicode{eightcircle}{00A0}
\pdfglyphtounicode{eightcircleinversesansserif}{00A0}
\pdfglyphtounicode{eightdeva}{00A0}
\pdfglyphtounicode{eighteencircle}{00A0}
\pdfglyphtounicode{eighteenparen}{00A0}
\pdfglyphtounicode{eighteenperiod}{00A0}
\pdfglyphtounicode{eightgujarati}{00A0}
\pdfglyphtounicode{eightgurmukhi}{00A0}
\pdfglyphtounicode{eighthackarabic}{00A0}
\pdfglyphtounicode{eighthangzhou}{00A0}
\pdfglyphtounicode{eighthnotebeamed}{00A0}
\pdfglyphtounicode{eightideographicparen}{00A0}
\pdfglyphtounicode{eightinferior}{00A0}
\pdfglyphtounicode{eightmonospace}{00A0}
\pdfglyphtounicode{eightoldstyle}{00A0}
\pdfglyphtounicode{eightparen}{00A0}
\pdfglyphtounicode{eightperiod}{00A0}
\pdfglyphtounicode{eightpersian}{00A0}
\pdfglyphtounicode{eightroman}{00A0}
\pdfglyphtounicode{eightsuperior}{00A0}
\pdfglyphtounicode{eightthai}{00A0}
\pdfglyphtounicode{einvertedbreve}{00A0}
\pdfglyphtounicode{eiotifiedcyrillic}{00A0}
\pdfglyphtounicode{ekatakana}{00A0}
\pdfglyphtounicode{ekatakanahalfwidth}{00A0}
\pdfglyphtounicode{ekonkargurmukhi}{00A0}
\pdfglyphtounicode{ekorean}{00A0}
\pdfglyphtounicode{elcyrillic}{00A0}
\pdfglyphtounicode{element}{00A0}
\pdfglyphtounicode{elevencircle}{00A0}
\pdfglyphtounicode{elevenparen}{00A0}
\pdfglyphtounicode{elevenperiod}{00A0}
\pdfglyphtounicode{elevenroman}{00A0}
\pdfglyphtounicode{ellipsis}{00A0}
\pdfglyphtounicode{ellipsisvertical}{00A0}
\pdfglyphtounicode{emacron}{00A0}
\pdfglyphtounicode{emacronacute}{00A0}
\pdfglyphtounicode{emacrongrave}{00A0}
\pdfglyphtounicode{emcyrillic}{00A0}
\pdfglyphtounicode{emdash}{00A0}
\pdfglyphtounicode{emdashvertical}{00A0}
\pdfglyphtounicode{emonospace}{00A0}
\pdfglyphtounicode{emphasismarkarmenian}{00A0}
\pdfglyphtounicode{emptyset}{00A0}
\pdfglyphtounicode{enbopomofo}{00A0}
\pdfglyphtounicode{encyrillic}{00A0}
\pdfglyphtounicode{endash}{00A0}
\pdfglyphtounicode{endashvertical}{00A0}
\pdfglyphtounicode{endescendercyrillic}{00A0}
\pdfglyphtounicode{eng}{00A0}
\pdfglyphtounicode{engbopomofo}{00A0}
\pdfglyphtounicode{enghecyrillic}{00A0}
\pdfglyphtounicode{enhookcyrillic}{00A0}
\pdfglyphtounicode{enspace}{00A0}
\pdfglyphtounicode{eogonek}{00A0}
\pdfglyphtounicode{eokorean}{00A0}
\pdfglyphtounicode{eopen}{00A0}
\pdfglyphtounicode{eopenclosed}{00A0}
\pdfglyphtounicode{eopenreversed}{00A0}
\pdfglyphtounicode{eopenreversedclosed}{00A0}
\pdfglyphtounicode{eopenreversedhook}{00A0}
\pdfglyphtounicode{eparen}{00A0}
\pdfglyphtounicode{epsilon}{00A0}
\pdfglyphtounicode{epsilon1}{00A0}
\pdfglyphtounicode{epsiloninv}{00A0}
\pdfglyphtounicode{epsilontonos}{00A0}
\pdfglyphtounicode{equal}{00A0}
\pdfglyphtounicode{equaldotleftright}{00A0}
\pdfglyphtounicode{equaldotrightleft}{00A0}
\pdfglyphtounicode{equalmonospace}{00A0}
\pdfglyphtounicode{equalorfollows}{00A0}
\pdfglyphtounicode{equalorgreater}{00A0}
\pdfglyphtounicode{equalorless}{00A0}
\pdfglyphtounicode{equalorprecedes}{00A0}
\pdfglyphtounicode{equalorsimilar}{00A0}
\pdfglyphtounicode{equalsdots}{00A0}
\pdfglyphtounicode{equalsmall}{00A0}
\pdfglyphtounicode{equalsuperior}{00A0}
\pdfglyphtounicode{equivalence}{00A0}
\pdfglyphtounicode{equivasymptotic}{00A0}
\pdfglyphtounicode{erbopomofo}{00A0}
\pdfglyphtounicode{ercyrillic}{00A0}
\pdfglyphtounicode{ereversed}{00A0}
\pdfglyphtounicode{ereversedcyrillic}{00A0}
\pdfglyphtounicode{escyrillic}{00A0}
\pdfglyphtounicode{esdescendercyrillic}{00A0}
\pdfglyphtounicode{esh}{00A0}
\pdfglyphtounicode{eshcurl}{00A0}
\pdfglyphtounicode{eshortdeva}{00A0}
\pdfglyphtounicode{eshortvowelsigndeva}{00A0}
\pdfglyphtounicode{eshreversedloop}{00A0}
\pdfglyphtounicode{eshsquatreversed}{00A0}
\pdfglyphtounicode{esmallhiragana}{00A0}
\pdfglyphtounicode{esmallkatakana}{00A0}
\pdfglyphtounicode{esmallkatakanahalfwidth}{00A0}
\pdfglyphtounicode{estimated}{00A0}
\pdfglyphtounicode{esuperior}{00A0}
\pdfglyphtounicode{eta}{00A0}
\pdfglyphtounicode{etarmenian}{00A0}
\pdfglyphtounicode{etatonos}{00A0}
\pdfglyphtounicode{eth}{00A0}
\pdfglyphtounicode{etilde}{00A0}
\pdfglyphtounicode{etildebelow}{00A0}
\pdfglyphtounicode{etnahtafoukhhebrew}{00A0}
\pdfglyphtounicode{etnahtafoukhlefthebrew}{00A0}
\pdfglyphtounicode{etnahtahebrew}{00A0}
\pdfglyphtounicode{etnahtalefthebrew}{00A0}
\pdfglyphtounicode{eturned}{00A0}
\pdfglyphtounicode{eukorean}{00A0}
\pdfglyphtounicode{euro}{00A0}
\pdfglyphtounicode{evowelsignbengali}{00A0}
\pdfglyphtounicode{evowelsigndeva}{00A0}
\pdfglyphtounicode{evowelsigngujarati}{00A0}
\pdfglyphtounicode{exclam}{00A0}
\pdfglyphtounicode{exclamarmenian}{00A0}
\pdfglyphtounicode{exclamdbl}{00A0}
\pdfglyphtounicode{exclamdown}{00A0}
\pdfglyphtounicode{exclamdownsmall}{00A0}
\pdfglyphtounicode{exclammonospace}{00A0}
\pdfglyphtounicode{exclamsmall}{00A0}
\pdfglyphtounicode{existential}{00A0}
\pdfglyphtounicode{ezh}{00A0}
\pdfglyphtounicode{ezhcaron}{00A0}
\pdfglyphtounicode{ezhcurl}{00A0}
\pdfglyphtounicode{ezhreversed}{00A0}
\pdfglyphtounicode{ezhtail}{00A0}
\pdfglyphtounicode{f}{00A0}
\pdfglyphtounicode{fadeva}{00A0}
\pdfglyphtounicode{fagurmukhi}{00A0}
\pdfglyphtounicode{fahrenheit}{00A0}
\pdfglyphtounicode{fathaarabic}{00A0}
\pdfglyphtounicode{fathalowarabic}{00A0}
\pdfglyphtounicode{fathatanarabic}{00A0}
\pdfglyphtounicode{fbopomofo}{00A0}
\pdfglyphtounicode{fcircle}{00A0}
\pdfglyphtounicode{fdotaccent}{00A0}
\pdfglyphtounicode{feharabic}{00A0}
\pdfglyphtounicode{feharmenian}{00A0}
\pdfglyphtounicode{fehfinalarabic}{00A0}
\pdfglyphtounicode{fehinitialarabic}{00A0}
\pdfglyphtounicode{fehmedialarabic}{00A0}
\pdfglyphtounicode{feicoptic}{00A0}
\pdfglyphtounicode{female}{00A0}
\pdfglyphtounicode{ff}{00A0}
\pdfglyphtounicode{ffi}{00A0}
\pdfglyphtounicode{ffl}{00A0}
\pdfglyphtounicode{fi}{00A0}
\pdfglyphtounicode{fifteencircle}{00A0}
\pdfglyphtounicode{fifteenparen}{00A0}
\pdfglyphtounicode{fifteenperiod}{00A0}
\pdfglyphtounicode{figuredash}{00A0}
\pdfglyphtounicode{filledbox}{00A0}
\pdfglyphtounicode{filledrect}{00A0}
\pdfglyphtounicode{finalkaf}{00A0}
\pdfglyphtounicode{finalkafdagesh}{00A0}
\pdfglyphtounicode{finalkafdageshhebrew}{00A0}
\pdfglyphtounicode{finalkafhebrew}{00A0}
\pdfglyphtounicode{finalkafqamats}{00A0}
\pdfglyphtounicode{finalkafqamatshebrew}{00A0}
\pdfglyphtounicode{finalkafsheva}{00A0}
\pdfglyphtounicode{finalkafshevahebrew}{00A0}
\pdfglyphtounicode{finalmem}{00A0}
\pdfglyphtounicode{finalmemhebrew}{00A0}
\pdfglyphtounicode{finalnun}{00A0}
\pdfglyphtounicode{finalnunhebrew}{00A0}
\pdfglyphtounicode{finalpe}{00A0}
\pdfglyphtounicode{finalpehebrew}{00A0}
\pdfglyphtounicode{finaltsadi}{00A0}
\pdfglyphtounicode{finaltsadihebrew}{00A0}
\pdfglyphtounicode{firsttonechinese}{00A0}
\pdfglyphtounicode{fisheye}{00A0}
\pdfglyphtounicode{fitacyrillic}{00A0}
\pdfglyphtounicode{five}{00A0}
\pdfglyphtounicode{fivearabic}{00A0}
\pdfglyphtounicode{fivebengali}{00A0}
\pdfglyphtounicode{fivecircle}{00A0}
\pdfglyphtounicode{fivecircleinversesansserif}{00A0}
\pdfglyphtounicode{fivedeva}{00A0}
\pdfglyphtounicode{fiveeighths}{00A0}
\pdfglyphtounicode{fivegujarati}{00A0}
\pdfglyphtounicode{fivegurmukhi}{00A0}
\pdfglyphtounicode{fivehackarabic}{00A0}
\pdfglyphtounicode{fivehangzhou}{00A0}
\pdfglyphtounicode{fiveideographicparen}{00A0}
\pdfglyphtounicode{fiveinferior}{00A0}
\pdfglyphtounicode{fivemonospace}{00A0}
\pdfglyphtounicode{fiveoldstyle}{00A0}
\pdfglyphtounicode{fiveparen}{00A0}
\pdfglyphtounicode{fiveperiod}{00A0}
\pdfglyphtounicode{fivepersian}{00A0}
\pdfglyphtounicode{fiveroman}{00A0}
\pdfglyphtounicode{fivesuperior}{00A0}
\pdfglyphtounicode{fivethai}{00A0}
\pdfglyphtounicode{fl}{00A0}
\pdfglyphtounicode{flat}{00A0}
\pdfglyphtounicode{floorleft}{00A0}
\pdfglyphtounicode{floorright}{00A0}
\pdfglyphtounicode{florin}{00A0}
\pdfglyphtounicode{fmonospace}{00A0}
\pdfglyphtounicode{fmsquare}{00A0}
\pdfglyphtounicode{fofanthai}{00A0}
\pdfglyphtounicode{fofathai}{00A0}
\pdfglyphtounicode{follownotdbleqv}{00A0}
\pdfglyphtounicode{follownotslnteql}{00A0}
\pdfglyphtounicode{followornoteqvlnt}{00A0}
\pdfglyphtounicode{follows}{00A0}
\pdfglyphtounicode{followsequal}{00A0}
\pdfglyphtounicode{followsorcurly}{00A0}
\pdfglyphtounicode{followsorequal}{00A0}
\pdfglyphtounicode{fongmanthai}{00A0}
\pdfglyphtounicode{forall}{00A0}
\pdfglyphtounicode{forces}{00A0}
\pdfglyphtounicode{forcesbar}{00A0}
\pdfglyphtounicode{fork}{00A0}
\pdfglyphtounicode{four}{00A0}
\pdfglyphtounicode{fourarabic}{00A0}
\pdfglyphtounicode{fourbengali}{00A0}
\pdfglyphtounicode{fourcircle}{00A0}
\pdfglyphtounicode{fourcircleinversesansserif}{00A0}
\pdfglyphtounicode{fourdeva}{00A0}
\pdfglyphtounicode{fourgujarati}{00A0}
\pdfglyphtounicode{fourgurmukhi}{00A0}
\pdfglyphtounicode{fourhackarabic}{00A0}
\pdfglyphtounicode{fourhangzhou}{00A0}
\pdfglyphtounicode{fourideographicparen}{00A0}
\pdfglyphtounicode{fourinferior}{00A0}
\pdfglyphtounicode{fourmonospace}{00A0}
\pdfglyphtounicode{fournumeratorbengali}{00A0}
\pdfglyphtounicode{fouroldstyle}{00A0}
\pdfglyphtounicode{fourparen}{00A0}
\pdfglyphtounicode{fourperiod}{00A0}
\pdfglyphtounicode{fourpersian}{00A0}
\pdfglyphtounicode{fourroman}{00A0}
\pdfglyphtounicode{foursuperior}{00A0}
\pdfglyphtounicode{fourteencircle}{00A0}
\pdfglyphtounicode{fourteenparen}{00A0}
\pdfglyphtounicode{fourteenperiod}{00A0}
\pdfglyphtounicode{fourthai}{00A0}
\pdfglyphtounicode{fourthtonechinese}{00A0}
\pdfglyphtounicode{fparen}{00A0}
\pdfglyphtounicode{fraction}{00A0}
\pdfglyphtounicode{franc}{00A0}
\pdfglyphtounicode{frown}{00A0}
\pdfglyphtounicode{g}{00A0}
\pdfglyphtounicode{gabengali}{00A0}
\pdfglyphtounicode{gacute}{00A0}
\pdfglyphtounicode{gadeva}{00A0}
\pdfglyphtounicode{gafarabic}{00A0}
\pdfglyphtounicode{gaffinalarabic}{00A0}
\pdfglyphtounicode{gafinitialarabic}{00A0}
\pdfglyphtounicode{gafmedialarabic}{00A0}
\pdfglyphtounicode{gagujarati}{00A0}
\pdfglyphtounicode{gagurmukhi}{00A0}
\pdfglyphtounicode{gahiragana}{00A0}
\pdfglyphtounicode{gakatakana}{00A0}
\pdfglyphtounicode{gamma}{00A0}
\pdfglyphtounicode{gammalatinsmall}{00A0}
\pdfglyphtounicode{gammasuperior}{00A0}
\pdfglyphtounicode{gangiacoptic}{00A0}
\pdfglyphtounicode{gbopomofo}{00A0}
\pdfglyphtounicode{gbreve}{00A0}
\pdfglyphtounicode{gcaron}{00A0}
\pdfglyphtounicode{gcedilla}{00A0}
\pdfglyphtounicode{gcircle}{00A0}
\pdfglyphtounicode{gcircumflex}{00A0}
\pdfglyphtounicode{gcommaaccent}{00A0}
\pdfglyphtounicode{gdot}{00A0}
\pdfglyphtounicode{gdotaccent}{00A0}
\pdfglyphtounicode{gecyrillic}{00A0}
\pdfglyphtounicode{gehiragana}{00A0}
\pdfglyphtounicode{gekatakana}{00A0}
\pdfglyphtounicode{geomequivalent}{00A0}
\pdfglyphtounicode{geometricallyequal}{00A0}
\pdfglyphtounicode{gereshaccenthebrew}{00A0}
\pdfglyphtounicode{gereshhebrew}{00A0}
\pdfglyphtounicode{gereshmuqdamhebrew}{00A0}
\pdfglyphtounicode{germandbls}{00A0}
\pdfglyphtounicode{gershayimaccenthebrew}{00A0}
\pdfglyphtounicode{gershayimhebrew}{00A0}
\pdfglyphtounicode{getamark}{00A0}
\pdfglyphtounicode{ghabengali}{00A0}
\pdfglyphtounicode{ghadarmenian}{00A0}
\pdfglyphtounicode{ghadeva}{00A0}
\pdfglyphtounicode{ghagujarati}{00A0}
\pdfglyphtounicode{ghagurmukhi}{00A0}
\pdfglyphtounicode{ghainarabic}{00A0}
\pdfglyphtounicode{ghainfinalarabic}{00A0}
\pdfglyphtounicode{ghaininitialarabic}{00A0}
\pdfglyphtounicode{ghainmedialarabic}{00A0}
\pdfglyphtounicode{ghemiddlehookcyrillic}{00A0}
\pdfglyphtounicode{ghestrokecyrillic}{00A0}
\pdfglyphtounicode{gheupturncyrillic}{00A0}
\pdfglyphtounicode{ghhadeva}{00A0}
\pdfglyphtounicode{ghhagurmukhi}{00A0}
\pdfglyphtounicode{ghook}{00A0}
\pdfglyphtounicode{ghzsquare}{00A0}
\pdfglyphtounicode{gihiragana}{00A0}
\pdfglyphtounicode{gikatakana}{00A0}
\pdfglyphtounicode{gimarmenian}{00A0}
\pdfglyphtounicode{gimel}{00A0}
\pdfglyphtounicode{gimeldagesh}{00A0}
\pdfglyphtounicode{gimeldageshhebrew}{00A0}
\pdfglyphtounicode{gimelhebrew}{00A0}
\pdfglyphtounicode{gjecyrillic}{00A0}
\pdfglyphtounicode{glottalinvertedstroke}{00A0}
\pdfglyphtounicode{glottalstop}{00A0}
\pdfglyphtounicode{glottalstopinverted}{00A0}
\pdfglyphtounicode{glottalstopmod}{00A0}
\pdfglyphtounicode{glottalstopreversed}{00A0}
\pdfglyphtounicode{glottalstopreversedmod}{00A0}
\pdfglyphtounicode{glottalstopreversedsuperior}{00A0}
\pdfglyphtounicode{glottalstopstroke}{00A0}
\pdfglyphtounicode{glottalstopstrokereversed}{00A0}
\pdfglyphtounicode{gmacron}{00A0}
\pdfglyphtounicode{gmonospace}{00A0}
\pdfglyphtounicode{gohiragana}{00A0}
\pdfglyphtounicode{gokatakana}{00A0}
\pdfglyphtounicode{gparen}{00A0}
\pdfglyphtounicode{gpasquare}{00A0}
\pdfglyphtounicode{gradient}{00A0}
\pdfglyphtounicode{grave}{00A0}
\pdfglyphtounicode{gravebelowcmb}{00A0}
\pdfglyphtounicode{gravecmb}{00A0}
\pdfglyphtounicode{gravecomb}{00A0}
\pdfglyphtounicode{gravedeva}{00A0}
\pdfglyphtounicode{gravelowmod}{00A0}
\pdfglyphtounicode{gravemonospace}{00A0}
\pdfglyphtounicode{gravetonecmb}{00A0}
\pdfglyphtounicode{greater}{00A0}
\pdfglyphtounicode{greaterdbleqlless}{00A0}
\pdfglyphtounicode{greaterdblequal}{00A0}
\pdfglyphtounicode{greaterdot}{00A0}
\pdfglyphtounicode{greaterequal}{00A0}
\pdfglyphtounicode{greaterequalorless}{00A0}
\pdfglyphtounicode{greaterlessequal}{00A0}
\pdfglyphtounicode{greatermonospace}{00A0}
\pdfglyphtounicode{greatermuch}{00A0}
\pdfglyphtounicode{greaternotdblequal}{00A0}
\pdfglyphtounicode{greaternotequal}{00A0}
\pdfglyphtounicode{greaterorapproxeql}{00A0}
\pdfglyphtounicode{greaterorequalslant}{00A0}
\pdfglyphtounicode{greaterorequivalent}{00A0}
\pdfglyphtounicode{greaterorless}{00A0}
\pdfglyphtounicode{greaterornotdbleql}{00A0}
\pdfglyphtounicode{greaterornotequal}{00A0}
\pdfglyphtounicode{greaterorsimilar}{00A0}
\pdfglyphtounicode{greateroverequal}{00A0}
\pdfglyphtounicode{greatersmall}{00A0}
\pdfglyphtounicode{gscript}{00A0}
\pdfglyphtounicode{gstroke}{00A0}
\pdfglyphtounicode{guhiragana}{00A0}
\pdfglyphtounicode{guillemotleft}{00A0}
\pdfglyphtounicode{guillemotright}{00A0}
\pdfglyphtounicode{guilsinglleft}{00A0}
\pdfglyphtounicode{guilsinglright}{00A0}
\pdfglyphtounicode{gukatakana}{00A0}
\pdfglyphtounicode{guramusquare}{00A0}
\pdfglyphtounicode{gysquare}{00A0}
\pdfglyphtounicode{h}{00A0}
\pdfglyphtounicode{haabkhasiancyrillic}{00A0}
\pdfglyphtounicode{haaltonearabic}{00A0}
\pdfglyphtounicode{habengali}{00A0}
\pdfglyphtounicode{hadescendercyrillic}{00A0}
\pdfglyphtounicode{hadeva}{00A0}
\pdfglyphtounicode{hagujarati}{00A0}
\pdfglyphtounicode{hagurmukhi}{00A0}
\pdfglyphtounicode{haharabic}{00A0}
\pdfglyphtounicode{hahfinalarabic}{00A0}
\pdfglyphtounicode{hahinitialarabic}{00A0}
\pdfglyphtounicode{hahiragana}{00A0}
\pdfglyphtounicode{hahmedialarabic}{00A0}
\pdfglyphtounicode{haitusquare}{00A0}
\pdfglyphtounicode{hakatakana}{00A0}
\pdfglyphtounicode{hakatakanahalfwidth}{00A0}
\pdfglyphtounicode{halantgurmukhi}{00A0}
\pdfglyphtounicode{hamzaarabic}{00A0}
\pdfglyphtounicode{hamzadammaarabic}{00A0}
\pdfglyphtounicode{hamzadammatanarabic}{00A0}
\pdfglyphtounicode{hamzafathaarabic}{00A0}
\pdfglyphtounicode{hamzafathatanarabic}{00A0}
\pdfglyphtounicode{hamzalowarabic}{00A0}
\pdfglyphtounicode{hamzalowkasraarabic}{00A0}
\pdfglyphtounicode{hamzalowkasratanarabic}{00A0}
\pdfglyphtounicode{hamzasukunarabic}{00A0}
\pdfglyphtounicode{hangulfiller}{00A0}
\pdfglyphtounicode{hardsigncyrillic}{00A0}
\pdfglyphtounicode{harpoondownleft}{00A0}
\pdfglyphtounicode{harpoondownright}{00A0}
\pdfglyphtounicode{harpoonleftbarbup}{00A0}
\pdfglyphtounicode{harpoonleftright}{00A0}
\pdfglyphtounicode{harpoonrightbarbup}{00A0}
\pdfglyphtounicode{harpoonrightleft}{00A0}
\pdfglyphtounicode{harpoonupleft}{00A0}
\pdfglyphtounicode{harpoonupright}{00A0}
\pdfglyphtounicode{hasquare}{00A0}
\pdfglyphtounicode{hatafpatah}{00A0}
\pdfglyphtounicode{hatafpatah16}{00A0}
\pdfglyphtounicode{hatafpatah23}{00A0}
\pdfglyphtounicode{hatafpatah2f}{00A0}
\pdfglyphtounicode{hatafpatahhebrew}{00A0}
\pdfglyphtounicode{hatafpatahnarrowhebrew}{00A0}
\pdfglyphtounicode{hatafpatahquarterhebrew}{00A0}
\pdfglyphtounicode{hatafpatahwidehebrew}{00A0}
\pdfglyphtounicode{hatafqamats}{00A0}
\pdfglyphtounicode{hatafqamats1b}{00A0}
\pdfglyphtounicode{hatafqamats28}{00A0}
\pdfglyphtounicode{hatafqamats34}{00A0}
\pdfglyphtounicode{hatafqamatshebrew}{00A0}
\pdfglyphtounicode{hatafqamatsnarrowhebrew}{00A0}
\pdfglyphtounicode{hatafqamatsquarterhebrew}{00A0}
\pdfglyphtounicode{hatafqamatswidehebrew}{00A0}
\pdfglyphtounicode{hatafsegol}{00A0}
\pdfglyphtounicode{hatafsegol17}{00A0}
\pdfglyphtounicode{hatafsegol24}{00A0}
\pdfglyphtounicode{hatafsegol30}{00A0}
\pdfglyphtounicode{hatafsegolhebrew}{00A0}
\pdfglyphtounicode{hatafsegolnarrowhebrew}{00A0}
\pdfglyphtounicode{hatafsegolquarterhebrew}{00A0}
\pdfglyphtounicode{hatafsegolwidehebrew}{00A0}
\pdfglyphtounicode{hbar}{00A0}
\pdfglyphtounicode{hbopomofo}{00A0}
\pdfglyphtounicode{hbrevebelow}{00A0}
\pdfglyphtounicode{hcedilla}{00A0}
\pdfglyphtounicode{hcircle}{00A0}
\pdfglyphtounicode{hcircumflex}{00A0}
\pdfglyphtounicode{hdieresis}{00A0}
\pdfglyphtounicode{hdotaccent}{00A0}
\pdfglyphtounicode{hdotbelow}{00A0}
\pdfglyphtounicode{he}{00A0}
\pdfglyphtounicode{heart}{00A0}
\pdfglyphtounicode{heartsuitblack}{00A0}
\pdfglyphtounicode{heartsuitwhite}{00A0}
\pdfglyphtounicode{hedagesh}{00A0}
\pdfglyphtounicode{hedageshhebrew}{00A0}
\pdfglyphtounicode{hehaltonearabic}{00A0}
\pdfglyphtounicode{heharabic}{00A0}
\pdfglyphtounicode{hehebrew}{00A0}
\pdfglyphtounicode{hehfinalaltonearabic}{00A0}
\pdfglyphtounicode{hehfinalalttwoarabic}{00A0}
\pdfglyphtounicode{hehfinalarabic}{00A0}
\pdfglyphtounicode{hehhamzaabovefinalarabic}{00A0}
\pdfglyphtounicode{hehhamzaaboveisolatedarabic}{00A0}
\pdfglyphtounicode{hehinitialaltonearabic}{00A0}
\pdfglyphtounicode{hehinitialarabic}{00A0}
\pdfglyphtounicode{hehiragana}{00A0}
\pdfglyphtounicode{hehmedialaltonearabic}{00A0}
\pdfglyphtounicode{hehmedialarabic}{00A0}
\pdfglyphtounicode{heiseierasquare}{00A0}
\pdfglyphtounicode{hekatakana}{00A0}
\pdfglyphtounicode{hekatakanahalfwidth}{00A0}
\pdfglyphtounicode{hekutaarusquare}{00A0}
\pdfglyphtounicode{henghook}{00A0}
\pdfglyphtounicode{herutusquare}{00A0}
\pdfglyphtounicode{het}{00A0}
\pdfglyphtounicode{hethebrew}{00A0}
\pdfglyphtounicode{hhook}{00A0}
\pdfglyphtounicode{hhooksuperior}{00A0}
\pdfglyphtounicode{hieuhacirclekorean}{00A0}
\pdfglyphtounicode{hieuhaparenkorean}{00A0}
\pdfglyphtounicode{hieuhcirclekorean}{00A0}
\pdfglyphtounicode{hieuhkorean}{00A0}
\pdfglyphtounicode{hieuhparenkorean}{00A0}
\pdfglyphtounicode{hihiragana}{00A0}
\pdfglyphtounicode{hikatakana}{00A0}
\pdfglyphtounicode{hikatakanahalfwidth}{00A0}
\pdfglyphtounicode{hiriq}{00A0}
\pdfglyphtounicode{hiriq14}{00A0}
\pdfglyphtounicode{hiriq21}{00A0}
\pdfglyphtounicode{hiriq2d}{00A0}
\pdfglyphtounicode{hiriqhebrew}{00A0}
\pdfglyphtounicode{hiriqnarrowhebrew}{00A0}
\pdfglyphtounicode{hiriqquarterhebrew}{00A0}
\pdfglyphtounicode{hiriqwidehebrew}{00A0}
\pdfglyphtounicode{hlinebelow}{00A0}
\pdfglyphtounicode{hmonospace}{00A0}
\pdfglyphtounicode{hoarmenian}{00A0}
\pdfglyphtounicode{hohipthai}{00A0}
\pdfglyphtounicode{hohiragana}{00A0}
\pdfglyphtounicode{hokatakana}{00A0}
\pdfglyphtounicode{hokatakanahalfwidth}{00A0}
\pdfglyphtounicode{holam}{00A0}
\pdfglyphtounicode{holam19}{00A0}
\pdfglyphtounicode{holam26}{00A0}
\pdfglyphtounicode{holam32}{00A0}
\pdfglyphtounicode{holamhebrew}{00A0}
\pdfglyphtounicode{holamnarrowhebrew}{00A0}
\pdfglyphtounicode{holamquarterhebrew}{00A0}
\pdfglyphtounicode{holamwidehebrew}{00A0}
\pdfglyphtounicode{honokhukthai}{00A0}
\pdfglyphtounicode{hookabovecomb}{00A0}
\pdfglyphtounicode{hookcmb}{00A0}
\pdfglyphtounicode{hookpalatalizedbelowcmb}{00A0}
\pdfglyphtounicode{hookretroflexbelowcmb}{00A0}
\pdfglyphtounicode{hoonsquare}{00A0}
\pdfglyphtounicode{horicoptic}{00A0}
\pdfglyphtounicode{horizontalbar}{00A0}
\pdfglyphtounicode{horncmb}{00A0}
\pdfglyphtounicode{hotsprings}{00A0}
\pdfglyphtounicode{house}{00A0}
\pdfglyphtounicode{hparen}{00A0}
\pdfglyphtounicode{hsuperior}{00A0}
\pdfglyphtounicode{hturned}{00A0}
\pdfglyphtounicode{huhiragana}{00A0}
\pdfglyphtounicode{huiitosquare}{00A0}
\pdfglyphtounicode{hukatakana}{00A0}
\pdfglyphtounicode{hukatakanahalfwidth}{00A0}
\pdfglyphtounicode{hungarumlaut}{00A0}
\pdfglyphtounicode{hungarumlautcmb}{00A0}
\pdfglyphtounicode{hv}{00A0}
\pdfglyphtounicode{hyphen}{00A0}
\pdfglyphtounicode{hyphenchar}{00A0}
\pdfglyphtounicode{hypheninferior}{00A0}
\pdfglyphtounicode{hyphenmonospace}{00A0}
\pdfglyphtounicode{hyphensmall}{00A0}
\pdfglyphtounicode{hyphensuperior}{00A0}
\pdfglyphtounicode{hyphentwo}{00A0}
\pdfglyphtounicode{i}{00A0}
\pdfglyphtounicode{iacute}{00A0}
\pdfglyphtounicode{iacyrillic}{00A0}
\pdfglyphtounicode{ibengali}{00A0}
\pdfglyphtounicode{ibopomofo}{00A0}
\pdfglyphtounicode{ibreve}{00A0}
\pdfglyphtounicode{icaron}{00A0}
\pdfglyphtounicode{icircle}{00A0}
\pdfglyphtounicode{icircumflex}{00A0}
\pdfglyphtounicode{icyrillic}{00A0}
\pdfglyphtounicode{idblgrave}{00A0}
\pdfglyphtounicode{ideographearthcircle}{00A0}
\pdfglyphtounicode{ideographfirecircle}{00A0}
\pdfglyphtounicode{ideographicallianceparen}{00A0}
\pdfglyphtounicode{ideographiccallparen}{00A0}
\pdfglyphtounicode{ideographiccentrecircle}{00A0}
\pdfglyphtounicode{ideographicclose}{00A0}
\pdfglyphtounicode{ideographiccomma}{00A0}
\pdfglyphtounicode{ideographiccommaleft}{00A0}
\pdfglyphtounicode{ideographiccongratulationparen}{00A0}
\pdfglyphtounicode{ideographiccorrectcircle}{00A0}
\pdfglyphtounicode{ideographicearthparen}{00A0}
\pdfglyphtounicode{ideographicenterpriseparen}{00A0}
\pdfglyphtounicode{ideographicexcellentcircle}{00A0}
\pdfglyphtounicode{ideographicfestivalparen}{00A0}
\pdfglyphtounicode{ideographicfinancialcircle}{00A0}
\pdfglyphtounicode{ideographicfinancialparen}{00A0}
\pdfglyphtounicode{ideographicfireparen}{00A0}
\pdfglyphtounicode{ideographichaveparen}{00A0}
\pdfglyphtounicode{ideographichighcircle}{00A0}
\pdfglyphtounicode{ideographiciterationmark}{00A0}
\pdfglyphtounicode{ideographiclaborcircle}{00A0}
\pdfglyphtounicode{ideographiclaborparen}{00A0}
\pdfglyphtounicode{ideographicleftcircle}{00A0}
\pdfglyphtounicode{ideographiclowcircle}{00A0}
\pdfglyphtounicode{ideographicmedicinecircle}{00A0}
\pdfglyphtounicode{ideographicmetalparen}{00A0}
\pdfglyphtounicode{ideographicmoonparen}{00A0}
\pdfglyphtounicode{ideographicnameparen}{00A0}
\pdfglyphtounicode{ideographicperiod}{00A0}
\pdfglyphtounicode{ideographicprintcircle}{00A0}
\pdfglyphtounicode{ideographicreachparen}{00A0}
\pdfglyphtounicode{ideographicrepresentparen}{00A0}
\pdfglyphtounicode{ideographicresourceparen}{00A0}
\pdfglyphtounicode{ideographicrightcircle}{00A0}
\pdfglyphtounicode{ideographicsecretcircle}{00A0}
\pdfglyphtounicode{ideographicselfparen}{00A0}
\pdfglyphtounicode{ideographicsocietyparen}{00A0}
\pdfglyphtounicode{ideographicspace}{00A0}
\pdfglyphtounicode{ideographicspecialparen}{00A0}
\pdfglyphtounicode{ideographicstockparen}{00A0}
\pdfglyphtounicode{ideographicstudyparen}{00A0}
\pdfglyphtounicode{ideographicsunparen}{00A0}
\pdfglyphtounicode{ideographicsuperviseparen}{00A0}
\pdfglyphtounicode{ideographicwaterparen}{00A0}
\pdfglyphtounicode{ideographicwoodparen}{00A0}
\pdfglyphtounicode{ideographiczero}{00A0}
\pdfglyphtounicode{ideographmetalcircle}{00A0}
\pdfglyphtounicode{ideographmooncircle}{00A0}
\pdfglyphtounicode{ideographnamecircle}{00A0}
\pdfglyphtounicode{ideographsuncircle}{00A0}
\pdfglyphtounicode{ideographwatercircle}{00A0}
\pdfglyphtounicode{ideographwoodcircle}{00A0}
\pdfglyphtounicode{ideva}{00A0}
\pdfglyphtounicode{idieresis}{00A0}
\pdfglyphtounicode{idieresisacute}{00A0}
\pdfglyphtounicode{idieresiscyrillic}{00A0}
\pdfglyphtounicode{idotbelow}{00A0}
\pdfglyphtounicode{iebrevecyrillic}{00A0}
\pdfglyphtounicode{iecyrillic}{00A0}
\pdfglyphtounicode{ieungacirclekorean}{00A0}
\pdfglyphtounicode{ieungaparenkorean}{00A0}
\pdfglyphtounicode{ieungcirclekorean}{00A0}
\pdfglyphtounicode{ieungkorean}{00A0}
\pdfglyphtounicode{ieungparenkorean}{00A0}
\pdfglyphtounicode{igrave}{00A0}
\pdfglyphtounicode{igujarati}{00A0}
\pdfglyphtounicode{igurmukhi}{00A0}
\pdfglyphtounicode{ihiragana}{00A0}
\pdfglyphtounicode{ihookabove}{00A0}
\pdfglyphtounicode{iibengali}{00A0}
\pdfglyphtounicode{iicyrillic}{00A0}
\pdfglyphtounicode{iideva}{00A0}
\pdfglyphtounicode{iigujarati}{00A0}
\pdfglyphtounicode{iigurmukhi}{00A0}
\pdfglyphtounicode{iimatragurmukhi}{00A0}
\pdfglyphtounicode{iinvertedbreve}{00A0}
\pdfglyphtounicode{iishortcyrillic}{00A0}
\pdfglyphtounicode{iivowelsignbengali}{00A0}
\pdfglyphtounicode{iivowelsigndeva}{00A0}
\pdfglyphtounicode{iivowelsigngujarati}{00A0}
\pdfglyphtounicode{ij}{00A0}
\pdfglyphtounicode{ikatakana}{00A0}
\pdfglyphtounicode{ikatakanahalfwidth}{00A0}
\pdfglyphtounicode{ikorean}{00A0}
\pdfglyphtounicode{ilde}{00A0}
\pdfglyphtounicode{iluyhebrew}{00A0}
\pdfglyphtounicode{imacron}{00A0}
\pdfglyphtounicode{imacroncyrillic}{00A0}
\pdfglyphtounicode{imageorapproximatelyequal}{00A0}
\pdfglyphtounicode{imatragurmukhi}{00A0}
\pdfglyphtounicode{imonospace}{00A0}
\pdfglyphtounicode{increment}{00A0}
\pdfglyphtounicode{infinity}{00A0}
\pdfglyphtounicode{iniarmenian}{00A0}
\pdfglyphtounicode{integerdivide}{00A0}
\pdfglyphtounicode{integral}{00A0}
\pdfglyphtounicode{integralbottom}{00A0}
\pdfglyphtounicode{integralbt}{00A0}
\pdfglyphtounicode{integralex}{00A0}
\pdfglyphtounicode{integraltop}{00A0}
\pdfglyphtounicode{integraltp}{00A0}
\pdfglyphtounicode{intercal}{00A0}
\pdfglyphtounicode{interrobang}{00A0}
\pdfglyphtounicode{interrobangdown}{00A0}
\pdfglyphtounicode{intersection}{00A0}
\pdfglyphtounicode{intersectiondbl}{00A0}
\pdfglyphtounicode{intersectionsq}{00A0}
\pdfglyphtounicode{intisquare}{00A0}
\pdfglyphtounicode{invbullet}{00A0}
\pdfglyphtounicode{invcircle}{00A0}
\pdfglyphtounicode{invsmileface}{00A0}
\pdfglyphtounicode{iocyrillic}{00A0}
\pdfglyphtounicode{iogonek}{00A0}
\pdfglyphtounicode{iota}{00A0}
\pdfglyphtounicode{iotadieresis}{00A0}
\pdfglyphtounicode{iotadieresistonos}{00A0}
\pdfglyphtounicode{iotalatin}{00A0}
\pdfglyphtounicode{iotatonos}{00A0}
\pdfglyphtounicode{iparen}{00A0}
\pdfglyphtounicode{irigurmukhi}{00A0}
\pdfglyphtounicode{ismallhiragana}{00A0}
\pdfglyphtounicode{ismallkatakana}{00A0}
\pdfglyphtounicode{ismallkatakanahalfwidth}{00A0}
\pdfglyphtounicode{issharbengali}{00A0}
\pdfglyphtounicode{istroke}{00A0}
\pdfglyphtounicode{isuperior}{00A0}
\pdfglyphtounicode{iterationhiragana}{00A0}
\pdfglyphtounicode{iterationkatakana}{00A0}
\pdfglyphtounicode{itilde}{00A0}
\pdfglyphtounicode{itildebelow}{00A0}
\pdfglyphtounicode{iubopomofo}{00A0}
\pdfglyphtounicode{iucyrillic}{00A0}
\pdfglyphtounicode{ivowelsignbengali}{00A0}
\pdfglyphtounicode{ivowelsigndeva}{00A0}
\pdfglyphtounicode{ivowelsigngujarati}{00A0}
\pdfglyphtounicode{izhitsacyrillic}{00A0}
\pdfglyphtounicode{izhitsadblgravecyrillic}{00A0}
\pdfglyphtounicode{j}{00A0}
\pdfglyphtounicode{jaarmenian}{00A0}
\pdfglyphtounicode{jabengali}{00A0}
\pdfglyphtounicode{jadeva}{00A0}
\pdfglyphtounicode{jagujarati}{00A0}
\pdfglyphtounicode{jagurmukhi}{00A0}
\pdfglyphtounicode{jbopomofo}{00A0}
\pdfglyphtounicode{jcaron}{00A0}
\pdfglyphtounicode{jcircle}{00A0}
\pdfglyphtounicode{jcircumflex}{00A0}
\pdfglyphtounicode{jcrossedtail}{00A0}
\pdfglyphtounicode{jdotlessstroke}{00A0}
\pdfglyphtounicode{jecyrillic}{00A0}
\pdfglyphtounicode{jeemarabic}{00A0}
\pdfglyphtounicode{jeemfinalarabic}{00A0}
\pdfglyphtounicode{jeeminitialarabic}{00A0}
\pdfglyphtounicode{jeemmedialarabic}{00A0}
\pdfglyphtounicode{jeharabic}{00A0}
\pdfglyphtounicode{jehfinalarabic}{00A0}
\pdfglyphtounicode{jhabengali}{00A0}
\pdfglyphtounicode{jhadeva}{00A0}
\pdfglyphtounicode{jhagujarati}{00A0}
\pdfglyphtounicode{jhagurmukhi}{00A0}
\pdfglyphtounicode{jheharmenian}{00A0}
\pdfglyphtounicode{jis}{00A0}
\pdfglyphtounicode{jmonospace}{00A0}
\pdfglyphtounicode{jparen}{00A0}
\pdfglyphtounicode{jsuperior}{00A0}
\pdfglyphtounicode{k}{00A0}
\pdfglyphtounicode{kabashkircyrillic}{00A0}
\pdfglyphtounicode{kabengali}{00A0}
\pdfglyphtounicode{kacute}{00A0}
\pdfglyphtounicode{kacyrillic}{00A0}
\pdfglyphtounicode{kadescendercyrillic}{00A0}
\pdfglyphtounicode{kadeva}{00A0}
\pdfglyphtounicode{kaf}{00A0}
\pdfglyphtounicode{kafarabic}{00A0}
\pdfglyphtounicode{kafdagesh}{00A0}
\pdfglyphtounicode{kafdageshhebrew}{00A0}
\pdfglyphtounicode{kaffinalarabic}{00A0}
\pdfglyphtounicode{kafhebrew}{00A0}
\pdfglyphtounicode{kafinitialarabic}{00A0}
\pdfglyphtounicode{kafmedialarabic}{00A0}
\pdfglyphtounicode{kafrafehebrew}{00A0}
\pdfglyphtounicode{kagujarati}{00A0}
\pdfglyphtounicode{kagurmukhi}{00A0}
\pdfglyphtounicode{kahiragana}{00A0}
\pdfglyphtounicode{kahookcyrillic}{00A0}
\pdfglyphtounicode{kakatakana}{00A0}
\pdfglyphtounicode{kakatakanahalfwidth}{00A0}
\pdfglyphtounicode{kappa}{00A0}
\pdfglyphtounicode{kappasymbolgreek}{00A0}
\pdfglyphtounicode{kapyeounmieumkorean}{00A0}
\pdfglyphtounicode{kapyeounphieuphkorean}{00A0}
\pdfglyphtounicode{kapyeounpieupkorean}{00A0}
\pdfglyphtounicode{kapyeounssangpieupkorean}{00A0}
\pdfglyphtounicode{karoriisquare}{00A0}
\pdfglyphtounicode{kashidaautoarabic}{00A0}
\pdfglyphtounicode{kashidaautonosidebearingarabic}{00A0}
\pdfglyphtounicode{kasmallkatakana}{00A0}
\pdfglyphtounicode{kasquare}{00A0}
\pdfglyphtounicode{kasraarabic}{00A0}
\pdfglyphtounicode{kasratanarabic}{00A0}
\pdfglyphtounicode{kastrokecyrillic}{00A0}
\pdfglyphtounicode{katahiraprolongmarkhalfwidth}{00A0}
\pdfglyphtounicode{kaverticalstrokecyrillic}{00A0}
\pdfglyphtounicode{kbopomofo}{00A0}
\pdfglyphtounicode{kcalsquare}{00A0}
\pdfglyphtounicode{kcaron}{00A0}
\pdfglyphtounicode{kcedilla}{00A0}
\pdfglyphtounicode{kcircle}{00A0}
\pdfglyphtounicode{kcommaaccent}{00A0}
\pdfglyphtounicode{kdotbelow}{00A0}
\pdfglyphtounicode{keharmenian}{00A0}
\pdfglyphtounicode{kehiragana}{00A0}
\pdfglyphtounicode{kekatakana}{00A0}
\pdfglyphtounicode{kekatakanahalfwidth}{00A0}
\pdfglyphtounicode{kenarmenian}{00A0}
\pdfglyphtounicode{kesmallkatakana}{00A0}
\pdfglyphtounicode{kgreenlandic}{00A0}
\pdfglyphtounicode{khabengali}{00A0}
\pdfglyphtounicode{khacyrillic}{00A0}
\pdfglyphtounicode{khadeva}{00A0}
\pdfglyphtounicode{khagujarati}{00A0}
\pdfglyphtounicode{khagurmukhi}{00A0}
\pdfglyphtounicode{khaharabic}{00A0}
\pdfglyphtounicode{khahfinalarabic}{00A0}
\pdfglyphtounicode{khahinitialarabic}{00A0}
\pdfglyphtounicode{khahmedialarabic}{00A0}
\pdfglyphtounicode{kheicoptic}{00A0}
\pdfglyphtounicode{khhadeva}{00A0}
\pdfglyphtounicode{khhagurmukhi}{00A0}
\pdfglyphtounicode{khieukhacirclekorean}{00A0}
\pdfglyphtounicode{khieukhaparenkorean}{00A0}
\pdfglyphtounicode{khieukhcirclekorean}{00A0}
\pdfglyphtounicode{khieukhkorean}{00A0}
\pdfglyphtounicode{khieukhparenkorean}{00A0}
\pdfglyphtounicode{khokhaithai}{00A0}
\pdfglyphtounicode{khokhonthai}{00A0}
\pdfglyphtounicode{khokhuatthai}{00A0}
\pdfglyphtounicode{khokhwaithai}{00A0}
\pdfglyphtounicode{khomutthai}{00A0}
\pdfglyphtounicode{khook}{00A0}
\pdfglyphtounicode{khorakhangthai}{00A0}
\pdfglyphtounicode{khzsquare}{00A0}
\pdfglyphtounicode{kihiragana}{00A0}
\pdfglyphtounicode{kikatakana}{00A0}
\pdfglyphtounicode{kikatakanahalfwidth}{00A0}
\pdfglyphtounicode{kiroguramusquare}{00A0}
\pdfglyphtounicode{kiromeetorusquare}{00A0}
\pdfglyphtounicode{kirosquare}{00A0}
\pdfglyphtounicode{kiyeokacirclekorean}{00A0}
\pdfglyphtounicode{kiyeokaparenkorean}{00A0}
\pdfglyphtounicode{kiyeokcirclekorean}{00A0}
\pdfglyphtounicode{kiyeokkorean}{00A0}
\pdfglyphtounicode{kiyeokparenkorean}{00A0}
\pdfglyphtounicode{kiyeoksioskorean}{00A0}
\pdfglyphtounicode{kjecyrillic}{00A0}
\pdfglyphtounicode{klinebelow}{00A0}
\pdfglyphtounicode{klsquare}{00A0}
\pdfglyphtounicode{kmcubedsquare}{00A0}
\pdfglyphtounicode{kmonospace}{00A0}
\pdfglyphtounicode{kmsquaredsquare}{00A0}
\pdfglyphtounicode{kohiragana}{00A0}
\pdfglyphtounicode{kohmsquare}{00A0}
\pdfglyphtounicode{kokaithai}{00A0}
\pdfglyphtounicode{kokatakana}{00A0}
\pdfglyphtounicode{kokatakanahalfwidth}{00A0}
\pdfglyphtounicode{kooposquare}{00A0}
\pdfglyphtounicode{koppacyrillic}{00A0}
\pdfglyphtounicode{koreanstandardsymbol}{00A0}
\pdfglyphtounicode{koroniscmb}{00A0}
\pdfglyphtounicode{kparen}{00A0}
\pdfglyphtounicode{kpasquare}{00A0}
\pdfglyphtounicode{ksicyrillic}{00A0}
\pdfglyphtounicode{ktsquare}{00A0}
\pdfglyphtounicode{kturned}{00A0}
\pdfglyphtounicode{kuhiragana}{00A0}
\pdfglyphtounicode{kukatakana}{00A0}
\pdfglyphtounicode{kukatakanahalfwidth}{00A0}
\pdfglyphtounicode{kvsquare}{00A0}
\pdfglyphtounicode{kwsquare}{00A0}
\pdfglyphtounicode{l}{00A0}
\pdfglyphtounicode{labengali}{00A0}
\pdfglyphtounicode{lacute}{00A0}
\pdfglyphtounicode{ladeva}{00A0}
\pdfglyphtounicode{lagujarati}{00A0}
\pdfglyphtounicode{lagurmukhi}{00A0}
\pdfglyphtounicode{lakkhangyaothai}{00A0}
\pdfglyphtounicode{lamaleffinalarabic}{00A0}
\pdfglyphtounicode{lamalefhamzaabovefinalarabic}{00A0}
\pdfglyphtounicode{lamalefhamzaaboveisolatedarabic}{00A0}
\pdfglyphtounicode{lamalefhamzabelowfinalarabic}{00A0}
\pdfglyphtounicode{lamalefhamzabelowisolatedarabic}{00A0}
\pdfglyphtounicode{lamalefisolatedarabic}{00A0}
\pdfglyphtounicode{lamalefmaddaabovefinalarabic}{00A0}
\pdfglyphtounicode{lamalefmaddaaboveisolatedarabic}{00A0}
\pdfglyphtounicode{lamarabic}{00A0}
\pdfglyphtounicode{lambda}{00A0}
\pdfglyphtounicode{lambdastroke}{00A0}
\pdfglyphtounicode{lamed}{00A0}
\pdfglyphtounicode{lameddagesh}{00A0}
\pdfglyphtounicode{lameddageshhebrew}{00A0}
\pdfglyphtounicode{lamedhebrew}{00A0}
\pdfglyphtounicode{lamedholam}{00A0}
\pdfglyphtounicode{lamedholamdagesh}{00A0}
\pdfglyphtounicode{lamedholamdageshhebrew}{00A0}
\pdfglyphtounicode{lamedholamhebrew}{00A0}
\pdfglyphtounicode{lamfinalarabic}{00A0}
\pdfglyphtounicode{lamhahinitialarabic}{00A0}
\pdfglyphtounicode{laminitialarabic}{00A0}
\pdfglyphtounicode{lamjeeminitialarabic}{00A0}
\pdfglyphtounicode{lamkhahinitialarabic}{00A0}
\pdfglyphtounicode{lamlamhehisolatedarabic}{00A0}
\pdfglyphtounicode{lammedialarabic}{00A0}
\pdfglyphtounicode{lammeemhahinitialarabic}{00A0}
\pdfglyphtounicode{lammeeminitialarabic}{00A0}
\pdfglyphtounicode{lammeemjeeminitialarabic}{00A0}
\pdfglyphtounicode{lammeemkhahinitialarabic}{00A0}
\pdfglyphtounicode{largecircle}{00A0}
\pdfglyphtounicode{latticetop}{00A0}
\pdfglyphtounicode{lbar}{00A0}
\pdfglyphtounicode{lbelt}{00A0}
\pdfglyphtounicode{lbopomofo}{00A0}
\pdfglyphtounicode{lcaron}{00A0}
\pdfglyphtounicode{lcedilla}{00A0}
\pdfglyphtounicode{lcircle}{00A0}
\pdfglyphtounicode{lcircumflexbelow}{00A0}
\pdfglyphtounicode{lcommaaccent}{00A0}
\pdfglyphtounicode{ldot}{00A0}
\pdfglyphtounicode{ldotaccent}{00A0}
\pdfglyphtounicode{ldotbelow}{00A0}
\pdfglyphtounicode{ldotbelowmacron}{00A0}
\pdfglyphtounicode{leftangleabovecmb}{00A0}
\pdfglyphtounicode{lefttackbelowcmb}{00A0}
\pdfglyphtounicode{less}{00A0}
\pdfglyphtounicode{lessdbleqlgreater}{00A0}
\pdfglyphtounicode{lessdblequal}{00A0}
\pdfglyphtounicode{lessdot}{00A0}
\pdfglyphtounicode{lessequal}{00A0}
\pdfglyphtounicode{lessequalgreater}{00A0}
\pdfglyphtounicode{lessequalorgreater}{00A0}
\pdfglyphtounicode{lessmonospace}{00A0}
\pdfglyphtounicode{lessmuch}{00A0}
\pdfglyphtounicode{lessnotdblequal}{00A0}
\pdfglyphtounicode{lessnotequal}{00A0}
\pdfglyphtounicode{lessorapproxeql}{00A0}
\pdfglyphtounicode{lessorequalslant}{00A0}
\pdfglyphtounicode{lessorequivalent}{00A0}
\pdfglyphtounicode{lessorgreater}{00A0}
\pdfglyphtounicode{lessornotdbleql}{00A0}
\pdfglyphtounicode{lessornotequal}{00A0}
\pdfglyphtounicode{lessorsimilar}{00A0}
\pdfglyphtounicode{lessoverequal}{00A0}
\pdfglyphtounicode{lesssmall}{00A0}
\pdfglyphtounicode{lezh}{00A0}
\pdfglyphtounicode{lfblock}{00A0}
\pdfglyphtounicode{lhookretroflex}{00A0}
\pdfglyphtounicode{lira}{00A0}
\pdfglyphtounicode{liwnarmenian}{00A0}
\pdfglyphtounicode{lj}{00A0}
\pdfglyphtounicode{ljecyrillic}{00A0}
\pdfglyphtounicode{ll}{00A0}
\pdfglyphtounicode{lladeva}{00A0}
\pdfglyphtounicode{llagujarati}{00A0}
\pdfglyphtounicode{llinebelow}{00A0}
\pdfglyphtounicode{llladeva}{00A0}
\pdfglyphtounicode{llvocalicbengali}{00A0}
\pdfglyphtounicode{llvocalicdeva}{00A0}
\pdfglyphtounicode{llvocalicvowelsignbengali}{00A0}
\pdfglyphtounicode{llvocalicvowelsigndeva}{00A0}
\pdfglyphtounicode{lmiddletilde}{00A0}
\pdfglyphtounicode{lmonospace}{00A0}
\pdfglyphtounicode{lmsquare}{00A0}
\pdfglyphtounicode{lochulathai}{00A0}
\pdfglyphtounicode{logicaland}{00A0}
\pdfglyphtounicode{logicalnot}{00A0}
\pdfglyphtounicode{logicalnotreversed}{00A0}
\pdfglyphtounicode{logicalor}{00A0}
\pdfglyphtounicode{lolingthai}{00A0}
\pdfglyphtounicode{longdbls}{00A0}
\pdfglyphtounicode{longs}{00A0}
\pdfglyphtounicode{longsh}{00A0}
\pdfglyphtounicode{longsi}{00A0}
\pdfglyphtounicode{longsl}{00A0}
\pdfglyphtounicode{longst}{00A0}
\pdfglyphtounicode{lowlinecenterline}{00A0}
\pdfglyphtounicode{lowlinecmb}{00A0}
\pdfglyphtounicode{lowlinedashed}{00A0}
\pdfglyphtounicode{lozenge}{00A0}
\pdfglyphtounicode{lparen}{00A0}
\pdfglyphtounicode{lscript}{00A0}
\pdfglyphtounicode{lslash}{00A0}
\pdfglyphtounicode{lsquare}{00A0}
\pdfglyphtounicode{lsuperior}{00A0}
\pdfglyphtounicode{ltshade}{00A0}
\pdfglyphtounicode{luthai}{00A0}
\pdfglyphtounicode{lvocalicbengali}{00A0}
\pdfglyphtounicode{lvocalicdeva}{00A0}
\pdfglyphtounicode{lvocalicvowelsignbengali}{00A0}
\pdfglyphtounicode{lvocalicvowelsigndeva}{00A0}
\pdfglyphtounicode{lxsquare}{00A0}
\pdfglyphtounicode{m}{00A0}
\pdfglyphtounicode{mabengali}{00A0}
\pdfglyphtounicode{macron}{00A0}
\pdfglyphtounicode{macronbelowcmb}{00A0}
\pdfglyphtounicode{macroncmb}{00A0}
\pdfglyphtounicode{macronlowmod}{00A0}
\pdfglyphtounicode{macronmonospace}{00A0}
\pdfglyphtounicode{macute}{00A0}
\pdfglyphtounicode{madeva}{00A0}
\pdfglyphtounicode{magujarati}{00A0}
\pdfglyphtounicode{magurmukhi}{00A0}
\pdfglyphtounicode{mahapakhhebrew}{00A0}
\pdfglyphtounicode{mahapakhlefthebrew}{00A0}
\pdfglyphtounicode{mahiragana}{00A0}
\pdfglyphtounicode{maichattawalowleftthai}{00A0}
\pdfglyphtounicode{maichattawalowrightthai}{00A0}
\pdfglyphtounicode{maichattawathai}{00A0}
\pdfglyphtounicode{maichattawaupperleftthai}{00A0}
\pdfglyphtounicode{maieklowleftthai}{00A0}
\pdfglyphtounicode{maieklowrightthai}{00A0}
\pdfglyphtounicode{maiekthai}{00A0}
\pdfglyphtounicode{maiekupperleftthai}{00A0}
\pdfglyphtounicode{maihanakatleftthai}{00A0}
\pdfglyphtounicode{maihanakatthai}{00A0}
\pdfglyphtounicode{maitaikhuleftthai}{00A0}
\pdfglyphtounicode{maitaikhuthai}{00A0}
\pdfglyphtounicode{maitholowleftthai}{00A0}
\pdfglyphtounicode{maitholowrightthai}{00A0}
\pdfglyphtounicode{maithothai}{00A0}
\pdfglyphtounicode{maithoupperleftthai}{00A0}
\pdfglyphtounicode{maitrilowleftthai}{00A0}
\pdfglyphtounicode{maitrilowrightthai}{00A0}
\pdfglyphtounicode{maitrithai}{00A0}
\pdfglyphtounicode{maitriupperleftthai}{00A0}
\pdfglyphtounicode{maiyamokthai}{00A0}
\pdfglyphtounicode{makatakana}{00A0}
\pdfglyphtounicode{makatakanahalfwidth}{00A0}
\pdfglyphtounicode{male}{00A0}
\pdfglyphtounicode{maltesecross}{00A0}
\pdfglyphtounicode{mansyonsquare}{00A0}
\pdfglyphtounicode{maqafhebrew}{00A0}
\pdfglyphtounicode{mars}{00A0}
\pdfglyphtounicode{masoracirclehebrew}{00A0}
\pdfglyphtounicode{masquare}{00A0}
\pdfglyphtounicode{mbopomofo}{00A0}
\pdfglyphtounicode{mbsquare}{00A0}
\pdfglyphtounicode{mcircle}{00A0}
\pdfglyphtounicode{mcubedsquare}{00A0}
\pdfglyphtounicode{mdotaccent}{00A0}
\pdfglyphtounicode{mdotbelow}{00A0}
\pdfglyphtounicode{measuredangle}{00A0}
\pdfglyphtounicode{meemarabic}{00A0}
\pdfglyphtounicode{meemfinalarabic}{00A0}
\pdfglyphtounicode{meeminitialarabic}{00A0}
\pdfglyphtounicode{meemmedialarabic}{00A0}
\pdfglyphtounicode{meemmeeminitialarabic}{00A0}
\pdfglyphtounicode{meemmeemisolatedarabic}{00A0}
\pdfglyphtounicode{meetorusquare}{00A0}
\pdfglyphtounicode{mehiragana}{00A0}
\pdfglyphtounicode{meizierasquare}{00A0}
\pdfglyphtounicode{mekatakana}{00A0}
\pdfglyphtounicode{mekatakanahalfwidth}{00A0}
\pdfglyphtounicode{mem}{00A0}
\pdfglyphtounicode{memdagesh}{00A0}
\pdfglyphtounicode{memdageshhebrew}{00A0}
\pdfglyphtounicode{memhebrew}{00A0}
\pdfglyphtounicode{menarmenian}{00A0}
\pdfglyphtounicode{merkhahebrew}{00A0}
\pdfglyphtounicode{merkhakefulahebrew}{00A0}
\pdfglyphtounicode{merkhakefulalefthebrew}{00A0}
\pdfglyphtounicode{merkhalefthebrew}{00A0}
\pdfglyphtounicode{mhook}{00A0}
\pdfglyphtounicode{mhzsquare}{00A0}
\pdfglyphtounicode{middledotkatakanahalfwidth}{00A0}
\pdfglyphtounicode{middot}{00A0}
\pdfglyphtounicode{mieumacirclekorean}{00A0}
\pdfglyphtounicode{mieumaparenkorean}{00A0}
\pdfglyphtounicode{mieumcirclekorean}{00A0}
\pdfglyphtounicode{mieumkorean}{00A0}
\pdfglyphtounicode{mieumpansioskorean}{00A0}
\pdfglyphtounicode{mieumparenkorean}{00A0}
\pdfglyphtounicode{mieumpieupkorean}{00A0}
\pdfglyphtounicode{mieumsioskorean}{00A0}
\pdfglyphtounicode{mihiragana}{00A0}
\pdfglyphtounicode{mikatakana}{00A0}
\pdfglyphtounicode{mikatakanahalfwidth}{00A0}
\pdfglyphtounicode{minus}{00A0}
\pdfglyphtounicode{minusbelowcmb}{00A0}
\pdfglyphtounicode{minuscircle}{00A0}
\pdfglyphtounicode{minusmod}{00A0}
\pdfglyphtounicode{minusplus}{00A0}
\pdfglyphtounicode{minute}{00A0}
\pdfglyphtounicode{miribaarusquare}{00A0}
\pdfglyphtounicode{mirisquare}{00A0}
\pdfglyphtounicode{mlonglegturned}{00A0}
\pdfglyphtounicode{mlsquare}{00A0}
\pdfglyphtounicode{mmcubedsquare}{00A0}
\pdfglyphtounicode{mmonospace}{00A0}
\pdfglyphtounicode{mmsquaredsquare}{00A0}
\pdfglyphtounicode{mohiragana}{00A0}
\pdfglyphtounicode{mohmsquare}{00A0}
\pdfglyphtounicode{mokatakana}{00A0}
\pdfglyphtounicode{mokatakanahalfwidth}{00A0}
\pdfglyphtounicode{molsquare}{00A0}
\pdfglyphtounicode{momathai}{00A0}
\pdfglyphtounicode{moverssquare}{00A0}
\pdfglyphtounicode{moverssquaredsquare}{00A0}
\pdfglyphtounicode{mparen}{00A0}
\pdfglyphtounicode{mpasquare}{00A0}
\pdfglyphtounicode{mssquare}{00A0}
\pdfglyphtounicode{msuperior}{00A0}
\pdfglyphtounicode{mturned}{00A0}
\pdfglyphtounicode{mu}{00A0}
\pdfglyphtounicode{mu1}{00A0}
\pdfglyphtounicode{muasquare}{00A0}
\pdfglyphtounicode{muchgreater}{00A0}
\pdfglyphtounicode{muchless}{00A0}
\pdfglyphtounicode{mufsquare}{00A0}
\pdfglyphtounicode{mugreek}{00A0}
\pdfglyphtounicode{mugsquare}{00A0}
\pdfglyphtounicode{muhiragana}{00A0}
\pdfglyphtounicode{mukatakana}{00A0}
\pdfglyphtounicode{mukatakanahalfwidth}{00A0}
\pdfglyphtounicode{mulsquare}{00A0}
\pdfglyphtounicode{multicloseleft}{00A0}
\pdfglyphtounicode{multicloseright}{00A0}
\pdfglyphtounicode{multimap}{00A0}
\pdfglyphtounicode{multiopenleft}{00A0}
\pdfglyphtounicode{multiopenright}{00A0}
\pdfglyphtounicode{multiply}{00A0}
\pdfglyphtounicode{mumsquare}{00A0}
\pdfglyphtounicode{munahhebrew}{00A0}
\pdfglyphtounicode{munahlefthebrew}{00A0}
\pdfglyphtounicode{musicalnote}{00A0}
\pdfglyphtounicode{musicalnotedbl}{00A0}
\pdfglyphtounicode{musicflatsign}{00A0}
\pdfglyphtounicode{musicsharpsign}{00A0}
\pdfglyphtounicode{mussquare}{00A0}
\pdfglyphtounicode{muvsquare}{00A0}
\pdfglyphtounicode{muwsquare}{00A0}
\pdfglyphtounicode{mvmegasquare}{00A0}
\pdfglyphtounicode{mvsquare}{00A0}
\pdfglyphtounicode{mwmegasquare}{00A0}
\pdfglyphtounicode{mwsquare}{00A0}
\pdfglyphtounicode{n}{00A0}
\pdfglyphtounicode{nabengali}{00A0}
\pdfglyphtounicode{nabla}{00A0}
\pdfglyphtounicode{nacute}{00A0}
\pdfglyphtounicode{nadeva}{00A0}
\pdfglyphtounicode{nagujarati}{00A0}
\pdfglyphtounicode{nagurmukhi}{00A0}
\pdfglyphtounicode{nahiragana}{00A0}
\pdfglyphtounicode{nakatakana}{00A0}
\pdfglyphtounicode{nakatakanahalfwidth}{00A0}
\pdfglyphtounicode{nand}{00A0}
\pdfglyphtounicode{napostrophe}{00A0}
\pdfglyphtounicode{nasquare}{00A0}
\pdfglyphtounicode{natural}{00A0}
\pdfglyphtounicode{nbopomofo}{00A0}
\pdfglyphtounicode{nbspace}{00A0}
\pdfglyphtounicode{ncaron}{00A0}
\pdfglyphtounicode{ncedilla}{00A0}
\pdfglyphtounicode{ncircle}{00A0}
\pdfglyphtounicode{ncircumflexbelow}{00A0}
\pdfglyphtounicode{ncommaaccent}{00A0}
\pdfglyphtounicode{ndotaccent}{00A0}
\pdfglyphtounicode{ndotbelow}{00A0}
\pdfglyphtounicode{negationslash}{00A0}
\pdfglyphtounicode{nehiragana}{00A0}
\pdfglyphtounicode{nekatakana}{00A0}
\pdfglyphtounicode{nekatakanahalfwidth}{00A0}
\pdfglyphtounicode{newsheqelsign}{00A0}
\pdfglyphtounicode{nfsquare}{00A0}
\pdfglyphtounicode{ng}{00A0}
\pdfglyphtounicode{ngabengali}{00A0}
\pdfglyphtounicode{ngadeva}{00A0}
\pdfglyphtounicode{ngagujarati}{00A0}
\pdfglyphtounicode{ngagurmukhi}{00A0}
\pdfglyphtounicode{ngonguthai}{00A0}
\pdfglyphtounicode{nhiragana}{00A0}
\pdfglyphtounicode{nhookleft}{00A0}
\pdfglyphtounicode{nhookretroflex}{00A0}
\pdfglyphtounicode{nieunacirclekorean}{00A0}
\pdfglyphtounicode{nieunaparenkorean}{00A0}
\pdfglyphtounicode{nieuncieuckorean}{00A0}
\pdfglyphtounicode{nieuncirclekorean}{00A0}
\pdfglyphtounicode{nieunhieuhkorean}{00A0}
\pdfglyphtounicode{nieunkorean}{00A0}
\pdfglyphtounicode{nieunpansioskorean}{00A0}
\pdfglyphtounicode{nieunparenkorean}{00A0}
\pdfglyphtounicode{nieunsioskorean}{00A0}
\pdfglyphtounicode{nieuntikeutkorean}{00A0}
\pdfglyphtounicode{nihiragana}{00A0}
\pdfglyphtounicode{nikatakana}{00A0}
\pdfglyphtounicode{nikatakanahalfwidth}{00A0}
\pdfglyphtounicode{nikhahitleftthai}{00A0}
\pdfglyphtounicode{nikhahitthai}{00A0}
\pdfglyphtounicode{nine}{00A0}
\pdfglyphtounicode{ninearabic}{00A0}
\pdfglyphtounicode{ninebengali}{00A0}
\pdfglyphtounicode{ninecircle}{00A0}
\pdfglyphtounicode{ninecircleinversesansserif}{00A0}
\pdfglyphtounicode{ninedeva}{00A0}
\pdfglyphtounicode{ninegujarati}{00A0}
\pdfglyphtounicode{ninegurmukhi}{00A0}
\pdfglyphtounicode{ninehackarabic}{00A0}
\pdfglyphtounicode{ninehangzhou}{00A0}
\pdfglyphtounicode{nineideographicparen}{00A0}
\pdfglyphtounicode{nineinferior}{00A0}
\pdfglyphtounicode{ninemonospace}{00A0}
\pdfglyphtounicode{nineoldstyle}{00A0}
\pdfglyphtounicode{nineparen}{00A0}
\pdfglyphtounicode{nineperiod}{00A0}
\pdfglyphtounicode{ninepersian}{00A0}
\pdfglyphtounicode{nineroman}{00A0}
\pdfglyphtounicode{ninesuperior}{00A0}
\pdfglyphtounicode{nineteencircle}{00A0}
\pdfglyphtounicode{nineteenparen}{00A0}
\pdfglyphtounicode{nineteenperiod}{00A0}
\pdfglyphtounicode{ninethai}{00A0}
\pdfglyphtounicode{nj}{00A0}
\pdfglyphtounicode{njecyrillic}{00A0}
\pdfglyphtounicode{nkatakana}{00A0}
\pdfglyphtounicode{nkatakanahalfwidth}{00A0}
\pdfglyphtounicode{nlegrightlong}{00A0}
\pdfglyphtounicode{nlinebelow}{00A0}
\pdfglyphtounicode{nmonospace}{00A0}
\pdfglyphtounicode{nmsquare}{00A0}
\pdfglyphtounicode{nnabengali}{00A0}
\pdfglyphtounicode{nnadeva}{00A0}
\pdfglyphtounicode{nnagujarati}{00A0}
\pdfglyphtounicode{nnagurmukhi}{00A0}
\pdfglyphtounicode{nnnadeva}{00A0}
\pdfglyphtounicode{nohiragana}{00A0}
\pdfglyphtounicode{nokatakana}{00A0}
\pdfglyphtounicode{nokatakanahalfwidth}{00A0}
\pdfglyphtounicode{nonbreakingspace}{00A0}
\pdfglyphtounicode{nonenthai}{00A0}
\pdfglyphtounicode{nonuthai}{00A0}
\pdfglyphtounicode{noonarabic}{00A0}
\pdfglyphtounicode{noonfinalarabic}{00A0}
\pdfglyphtounicode{noonghunnaarabic}{00A0}
\pdfglyphtounicode{noonghunnafinalarabic}{00A0}
\pdfglyphtounicode{noonhehinitialarabic}{00A0}
\pdfglyphtounicode{nooninitialarabic}{00A0}
\pdfglyphtounicode{noonjeeminitialarabic}{00A0}
\pdfglyphtounicode{noonjeemisolatedarabic}{00A0}
\pdfglyphtounicode{noonmedialarabic}{00A0}
\pdfglyphtounicode{noonmeeminitialarabic}{00A0}
\pdfglyphtounicode{noonmeemisolatedarabic}{00A0}
\pdfglyphtounicode{noonnoonfinalarabic}{00A0}
\pdfglyphtounicode{notapproxequal}{00A0}
\pdfglyphtounicode{notarrowboth}{00A0}
\pdfglyphtounicode{notarrowleft}{00A0}
\pdfglyphtounicode{notarrowright}{00A0}
\pdfglyphtounicode{notbar}{00A0}
\pdfglyphtounicode{notcontains}{00A0}
\pdfglyphtounicode{notdblarrowboth}{00A0}
\pdfglyphtounicode{notdblarrowleft}{00A0}
\pdfglyphtounicode{notdblarrowright}{00A0}
\pdfglyphtounicode{notelement}{00A0}
\pdfglyphtounicode{notelementof}{00A0}
\pdfglyphtounicode{notequal}{00A0}
\pdfglyphtounicode{notexistential}{00A0}
\pdfglyphtounicode{notfollows}{00A0}
\pdfglyphtounicode{notfollowsoreql}{00A0}
\pdfglyphtounicode{notforces}{00A0}
\pdfglyphtounicode{notforcesextra}{00A0}
\pdfglyphtounicode{notgreater}{00A0}
\pdfglyphtounicode{notgreaterdblequal}{00A0}
\pdfglyphtounicode{notgreaterequal}{00A0}
\pdfglyphtounicode{notgreaternorequal}{00A0}
\pdfglyphtounicode{notgreaternorless}{00A0}
\pdfglyphtounicode{notgreaterorslnteql}{00A0}
\pdfglyphtounicode{notidentical}{00A0}
\pdfglyphtounicode{notless}{00A0}
\pdfglyphtounicode{notlessdblequal}{00A0}
\pdfglyphtounicode{notlessequal}{00A0}
\pdfglyphtounicode{notlessnorequal}{00A0}
\pdfglyphtounicode{notlessorslnteql}{00A0}
\pdfglyphtounicode{notparallel}{00A0}
\pdfglyphtounicode{notprecedes}{00A0}
\pdfglyphtounicode{notprecedesoreql}{00A0}
\pdfglyphtounicode{notsatisfies}{00A0}
\pdfglyphtounicode{notsimilar}{00A0}
\pdfglyphtounicode{notsubset}{00A0}
\pdfglyphtounicode{notsubseteql}{00A0}
\pdfglyphtounicode{notsubsetordbleql}{00A0}
\pdfglyphtounicode{notsubsetoreql}{00A0}
\pdfglyphtounicode{notsucceeds}{00A0}
\pdfglyphtounicode{notsuperset}{00A0}
\pdfglyphtounicode{notsuperseteql}{00A0}
\pdfglyphtounicode{notsupersetordbleql}{00A0}
\pdfglyphtounicode{notsupersetoreql}{00A0}
\pdfglyphtounicode{nottriangeqlleft}{00A0}
\pdfglyphtounicode{nottriangeqlright}{00A0}
\pdfglyphtounicode{nottriangleleft}{00A0}
\pdfglyphtounicode{nottriangleright}{00A0}
\pdfglyphtounicode{notturnstile}{00A0}
\pdfglyphtounicode{nowarmenian}{00A0}
\pdfglyphtounicode{nparen}{00A0}
\pdfglyphtounicode{nssquare}{00A0}
\pdfglyphtounicode{nsuperior}{00A0}
\pdfglyphtounicode{ntilde}{00A0}
\pdfglyphtounicode{nu}{00A0}
\pdfglyphtounicode{nuhiragana}{00A0}
\pdfglyphtounicode{nukatakana}{00A0}
\pdfglyphtounicode{nukatakanahalfwidth}{00A0}
\pdfglyphtounicode{nuktabengali}{00A0}
\pdfglyphtounicode{nuktadeva}{00A0}
\pdfglyphtounicode{nuktagujarati}{00A0}
\pdfglyphtounicode{nuktagurmukhi}{00A0}
\pdfglyphtounicode{numbersign}{00A0}
\pdfglyphtounicode{numbersignmonospace}{00A0}
\pdfglyphtounicode{numbersignsmall}{00A0}
\pdfglyphtounicode{numeralsigngreek}{00A0}
\pdfglyphtounicode{numeralsignlowergreek}{00A0}
\pdfglyphtounicode{numero}{00A0}
\pdfglyphtounicode{nun}{00A0}
\pdfglyphtounicode{nundagesh}{00A0}
\pdfglyphtounicode{nundageshhebrew}{00A0}
\pdfglyphtounicode{nunhebrew}{00A0}
\pdfglyphtounicode{nvsquare}{00A0}
\pdfglyphtounicode{nwsquare}{00A0}
\pdfglyphtounicode{nyabengali}{00A0}
\pdfglyphtounicode{nyadeva}{00A0}
\pdfglyphtounicode{nyagujarati}{00A0}
\pdfglyphtounicode{nyagurmukhi}{00A0}
\pdfglyphtounicode{o}{00A0}
\pdfglyphtounicode{oacute}{00A0}
\pdfglyphtounicode{oangthai}{00A0}
\pdfglyphtounicode{obarred}{00A0}
\pdfglyphtounicode{obarredcyrillic}{00A0}
\pdfglyphtounicode{obarreddieresiscyrillic}{00A0}
\pdfglyphtounicode{obengali}{00A0}
\pdfglyphtounicode{obopomofo}{00A0}
\pdfglyphtounicode{obreve}{00A0}
\pdfglyphtounicode{ocandradeva}{00A0}
\pdfglyphtounicode{ocandragujarati}{00A0}
\pdfglyphtounicode{ocandravowelsigndeva}{00A0}
\pdfglyphtounicode{ocandravowelsigngujarati}{00A0}
\pdfglyphtounicode{ocaron}{00A0}
\pdfglyphtounicode{ocircle}{00A0}
\pdfglyphtounicode{ocircumflex}{00A0}
\pdfglyphtounicode{ocircumflexacute}{00A0}
\pdfglyphtounicode{ocircumflexdotbelow}{00A0}
\pdfglyphtounicode{ocircumflexgrave}{00A0}
\pdfglyphtounicode{ocircumflexhookabove}{00A0}
\pdfglyphtounicode{ocircumflextilde}{00A0}
\pdfglyphtounicode{ocyrillic}{00A0}
\pdfglyphtounicode{odblacute}{00A0}
\pdfglyphtounicode{odblgrave}{00A0}
\pdfglyphtounicode{odeva}{00A0}
\pdfglyphtounicode{odieresis}{00A0}
\pdfglyphtounicode{odieresiscyrillic}{00A0}
\pdfglyphtounicode{odotbelow}{00A0}
\pdfglyphtounicode{oe}{00A0}
\pdfglyphtounicode{oekorean}{00A0}
\pdfglyphtounicode{ogonek}{00A0}
\pdfglyphtounicode{ogonekcmb}{00A0}
\pdfglyphtounicode{ograve}{00A0}
\pdfglyphtounicode{ogujarati}{00A0}
\pdfglyphtounicode{oharmenian}{00A0}
\pdfglyphtounicode{ohiragana}{00A0}
\pdfglyphtounicode{ohookabove}{00A0}
\pdfglyphtounicode{ohorn}{00A0}
\pdfglyphtounicode{ohornacute}{00A0}
\pdfglyphtounicode{ohorndotbelow}{00A0}
\pdfglyphtounicode{ohorngrave}{00A0}
\pdfglyphtounicode{ohornhookabove}{00A0}
\pdfglyphtounicode{ohorntilde}{00A0}
\pdfglyphtounicode{ohungarumlaut}{00A0}
\pdfglyphtounicode{oi}{00A0}
\pdfglyphtounicode{oinvertedbreve}{00A0}
\pdfglyphtounicode{okatakana}{00A0}
\pdfglyphtounicode{okatakanahalfwidth}{00A0}
\pdfglyphtounicode{okorean}{00A0}
\pdfglyphtounicode{olehebrew}{00A0}
\pdfglyphtounicode{omacron}{00A0}
\pdfglyphtounicode{omacronacute}{00A0}
\pdfglyphtounicode{omacrongrave}{00A0}
\pdfglyphtounicode{omdeva}{00A0}
\pdfglyphtounicode{omega}{00A0}
\pdfglyphtounicode{omega1}{00A0}
\pdfglyphtounicode{omegacyrillic}{00A0}
\pdfglyphtounicode{omegalatinclosed}{00A0}
\pdfglyphtounicode{omegaroundcyrillic}{00A0}
\pdfglyphtounicode{omegatitlocyrillic}{00A0}
\pdfglyphtounicode{omegatonos}{00A0}
\pdfglyphtounicode{omgujarati}{00A0}
\pdfglyphtounicode{omicron}{00A0}
\pdfglyphtounicode{omicrontonos}{00A0}
\pdfglyphtounicode{omonospace}{00A0}
\pdfglyphtounicode{one}{00A0}
\pdfglyphtounicode{onearabic}{00A0}
\pdfglyphtounicode{onebengali}{00A0}
\pdfglyphtounicode{onecircle}{00A0}
\pdfglyphtounicode{onecircleinversesansserif}{00A0}
\pdfglyphtounicode{onedeva}{00A0}
\pdfglyphtounicode{onedotenleader}{00A0}
\pdfglyphtounicode{oneeighth}{00A0}
\pdfglyphtounicode{onefitted}{00A0}
\pdfglyphtounicode{onegujarati}{00A0}
\pdfglyphtounicode{onegurmukhi}{00A0}
\pdfglyphtounicode{onehackarabic}{00A0}
\pdfglyphtounicode{onehalf}{00A0}
\pdfglyphtounicode{onehangzhou}{00A0}
\pdfglyphtounicode{oneideographicparen}{00A0}
\pdfglyphtounicode{oneinferior}{00A0}
\pdfglyphtounicode{onemonospace}{00A0}
\pdfglyphtounicode{onenumeratorbengali}{00A0}
\pdfglyphtounicode{oneoldstyle}{00A0}
\pdfglyphtounicode{oneparen}{00A0}
\pdfglyphtounicode{oneperiod}{00A0}
\pdfglyphtounicode{onepersian}{00A0}
\pdfglyphtounicode{onequarter}{00A0}
\pdfglyphtounicode{oneroman}{00A0}
\pdfglyphtounicode{onesuperior}{00A0}
\pdfglyphtounicode{onethai}{00A0}
\pdfglyphtounicode{onethird}{00A0}
\pdfglyphtounicode{oogonek}{00A0}
\pdfglyphtounicode{oogonekmacron}{00A0}
\pdfglyphtounicode{oogurmukhi}{00A0}
\pdfglyphtounicode{oomatragurmukhi}{00A0}
\pdfglyphtounicode{oopen}{00A0}
\pdfglyphtounicode{oparen}{00A0}
\pdfglyphtounicode{openbullet}{00A0}
\pdfglyphtounicode{option}{00A0}
\pdfglyphtounicode{ordfeminine}{00A0}
\pdfglyphtounicode{ordmasculine}{00A0}
\pdfglyphtounicode{orthogonal}{00A0}
\pdfglyphtounicode{orunderscore}{00A0}
\pdfglyphtounicode{oshortdeva}{00A0}
\pdfglyphtounicode{oshortvowelsigndeva}{00A0}
\pdfglyphtounicode{oslash}{00A0}
\pdfglyphtounicode{oslashacute}{00A0}
\pdfglyphtounicode{osmallhiragana}{00A0}
\pdfglyphtounicode{osmallkatakana}{00A0}
\pdfglyphtounicode{osmallkatakanahalfwidth}{00A0}
\pdfglyphtounicode{ostrokeacute}{00A0}
\pdfglyphtounicode{osuperior}{00A0}
\pdfglyphtounicode{otcyrillic}{00A0}
\pdfglyphtounicode{otilde}{00A0}
\pdfglyphtounicode{otildeacute}{00A0}
\pdfglyphtounicode{otildedieresis}{00A0}
\pdfglyphtounicode{oubopomofo}{00A0}
\pdfglyphtounicode{overline}{00A0}
\pdfglyphtounicode{overlinecenterline}{00A0}
\pdfglyphtounicode{overlinecmb}{00A0}
\pdfglyphtounicode{overlinedashed}{00A0}
\pdfglyphtounicode{overlinedblwavy}{00A0}
\pdfglyphtounicode{overlinewavy}{00A0}
\pdfglyphtounicode{overscore}{00A0}
\pdfglyphtounicode{ovowelsignbengali}{00A0}
\pdfglyphtounicode{ovowelsigndeva}{00A0}
\pdfglyphtounicode{ovowelsigngujarati}{00A0}
\pdfglyphtounicode{owner}{00A0}
\pdfglyphtounicode{p}{00A0}
\pdfglyphtounicode{paampssquare}{00A0}
\pdfglyphtounicode{paasentosquare}{00A0}
\pdfglyphtounicode{pabengali}{00A0}
\pdfglyphtounicode{pacute}{00A0}
\pdfglyphtounicode{padeva}{00A0}
\pdfglyphtounicode{pagedown}{00A0}
\pdfglyphtounicode{pageup}{00A0}
\pdfglyphtounicode{pagujarati}{00A0}
\pdfglyphtounicode{pagurmukhi}{00A0}
\pdfglyphtounicode{pahiragana}{00A0}
\pdfglyphtounicode{paiyannoithai}{00A0}
\pdfglyphtounicode{pakatakana}{00A0}
\pdfglyphtounicode{palatalizationcyrilliccmb}{00A0}
\pdfglyphtounicode{palochkacyrillic}{00A0}
\pdfglyphtounicode{pansioskorean}{00A0}
\pdfglyphtounicode{paragraph}{00A0}
\pdfglyphtounicode{parallel}{00A0}
\pdfglyphtounicode{parenleft}{00A0}
\pdfglyphtounicode{parenleftaltonearabic}{00A0}
\pdfglyphtounicode{parenleftbt}{00A0}
\pdfglyphtounicode{parenleftex}{00A0}
\pdfglyphtounicode{parenleftinferior}{00A0}
\pdfglyphtounicode{parenleftmonospace}{00A0}
\pdfglyphtounicode{parenleftsmall}{00A0}
\pdfglyphtounicode{parenleftsuperior}{00A0}
\pdfglyphtounicode{parenlefttp}{00A0}
\pdfglyphtounicode{parenleftvertical}{00A0}
\pdfglyphtounicode{parenright}{00A0}
\pdfglyphtounicode{parenrightaltonearabic}{00A0}
\pdfglyphtounicode{parenrightbt}{00A0}
\pdfglyphtounicode{parenrightex}{00A0}
\pdfglyphtounicode{parenrightinferior}{00A0}
\pdfglyphtounicode{parenrightmonospace}{00A0}
\pdfglyphtounicode{parenrightsmall}{00A0}
\pdfglyphtounicode{parenrightsuperior}{00A0}
\pdfglyphtounicode{parenrighttp}{00A0}
\pdfglyphtounicode{parenrightvertical}{00A0}
\pdfglyphtounicode{partialdiff}{00A0}
\pdfglyphtounicode{paseqhebrew}{00A0}
\pdfglyphtounicode{pashtahebrew}{00A0}
\pdfglyphtounicode{pasquare}{00A0}
\pdfglyphtounicode{patah}{00A0}
\pdfglyphtounicode{patah11}{00A0}
\pdfglyphtounicode{patah1d}{00A0}
\pdfglyphtounicode{patah2a}{00A0}
\pdfglyphtounicode{patahhebrew}{00A0}
\pdfglyphtounicode{patahnarrowhebrew}{00A0}
\pdfglyphtounicode{patahquarterhebrew}{00A0}
\pdfglyphtounicode{patahwidehebrew}{00A0}
\pdfglyphtounicode{pazerhebrew}{00A0}
\pdfglyphtounicode{pbopomofo}{00A0}
\pdfglyphtounicode{pcircle}{00A0}
\pdfglyphtounicode{pdotaccent}{00A0}
\pdfglyphtounicode{pe}{00A0}
\pdfglyphtounicode{pecyrillic}{00A0}
\pdfglyphtounicode{pedagesh}{00A0}
\pdfglyphtounicode{pedageshhebrew}{00A0}
\pdfglyphtounicode{peezisquare}{00A0}
\pdfglyphtounicode{pefinaldageshhebrew}{00A0}
\pdfglyphtounicode{peharabic}{00A0}
\pdfglyphtounicode{peharmenian}{00A0}
\pdfglyphtounicode{pehebrew}{00A0}
\pdfglyphtounicode{pehfinalarabic}{00A0}
\pdfglyphtounicode{pehinitialarabic}{00A0}
\pdfglyphtounicode{pehiragana}{00A0}
\pdfglyphtounicode{pehmedialarabic}{00A0}
\pdfglyphtounicode{pekatakana}{00A0}
\pdfglyphtounicode{pemiddlehookcyrillic}{00A0}
\pdfglyphtounicode{perafehebrew}{00A0}
\pdfglyphtounicode{percent}{00A0}
\pdfglyphtounicode{percentarabic}{00A0}
\pdfglyphtounicode{percentmonospace}{00A0}
\pdfglyphtounicode{percentsmall}{00A0}
\pdfglyphtounicode{period}{00A0}
\pdfglyphtounicode{periodarmenian}{00A0}
\pdfglyphtounicode{periodcentered}{00A0}
\pdfglyphtounicode{periodhalfwidth}{00A0}
\pdfglyphtounicode{periodinferior}{00A0}
\pdfglyphtounicode{periodmonospace}{00A0}
\pdfglyphtounicode{periodsmall}{00A0}
\pdfglyphtounicode{periodsuperior}{00A0}
\pdfglyphtounicode{perispomenigreekcmb}{00A0}
\pdfglyphtounicode{perpcorrespond}{00A0}
\pdfglyphtounicode{perpendicular}{00A0}
\pdfglyphtounicode{pertenthousand}{00A0}
\pdfglyphtounicode{perthousand}{00A0}
\pdfglyphtounicode{peseta}{00A0}
\pdfglyphtounicode{pfsquare}{00A0}
\pdfglyphtounicode{phabengali}{00A0}
\pdfglyphtounicode{phadeva}{00A0}
\pdfglyphtounicode{phagujarati}{00A0}
\pdfglyphtounicode{phagurmukhi}{00A0}
\pdfglyphtounicode{phi}{00A0}
\pdfglyphtounicode{phi1}{00A0}
\pdfglyphtounicode{phieuphacirclekorean}{00A0}
\pdfglyphtounicode{phieuphaparenkorean}{00A0}
\pdfglyphtounicode{phieuphcirclekorean}{00A0}
\pdfglyphtounicode{phieuphkorean}{00A0}
\pdfglyphtounicode{phieuphparenkorean}{00A0}
\pdfglyphtounicode{philatin}{00A0}
\pdfglyphtounicode{phinthuthai}{00A0}
\pdfglyphtounicode{phisymbolgreek}{00A0}
\pdfglyphtounicode{phook}{00A0}
\pdfglyphtounicode{phophanthai}{00A0}
\pdfglyphtounicode{phophungthai}{00A0}
\pdfglyphtounicode{phosamphaothai}{00A0}
\pdfglyphtounicode{pi}{00A0}
\pdfglyphtounicode{pi1}{00A0}
\pdfglyphtounicode{pieupacirclekorean}{00A0}
\pdfglyphtounicode{pieupaparenkorean}{00A0}
\pdfglyphtounicode{pieupcieuckorean}{00A0}
\pdfglyphtounicode{pieupcirclekorean}{00A0}
\pdfglyphtounicode{pieupkiyeokkorean}{00A0}
\pdfglyphtounicode{pieupkorean}{00A0}
\pdfglyphtounicode{pieupparenkorean}{00A0}
\pdfglyphtounicode{pieupsioskiyeokkorean}{00A0}
\pdfglyphtounicode{pieupsioskorean}{00A0}
\pdfglyphtounicode{pieupsiostikeutkorean}{00A0}
\pdfglyphtounicode{pieupthieuthkorean}{00A0}
\pdfglyphtounicode{pieuptikeutkorean}{00A0}
\pdfglyphtounicode{pihiragana}{00A0}
\pdfglyphtounicode{pikatakana}{00A0}
\pdfglyphtounicode{pisymbolgreek}{00A0}
\pdfglyphtounicode{piwrarmenian}{00A0}
\pdfglyphtounicode{planckover2pi}{00A0}
\pdfglyphtounicode{planckover2pi1}{00A0}
\pdfglyphtounicode{plus}{00A0}
\pdfglyphtounicode{plusbelowcmb}{00A0}
\pdfglyphtounicode{pluscircle}{00A0}
\pdfglyphtounicode{plusminus}{00A0}
\pdfglyphtounicode{plusmod}{00A0}
\pdfglyphtounicode{plusmonospace}{00A0}
\pdfglyphtounicode{plussmall}{00A0}
\pdfglyphtounicode{plussuperior}{00A0}
\pdfglyphtounicode{pmonospace}{00A0}
\pdfglyphtounicode{pmsquare}{00A0}
\pdfglyphtounicode{pohiragana}{00A0}
\pdfglyphtounicode{pointingindexdownwhite}{00A0}
\pdfglyphtounicode{pointingindexleftwhite}{00A0}
\pdfglyphtounicode{pointingindexrightwhite}{00A0}
\pdfglyphtounicode{pointingindexupwhite}{00A0}
\pdfglyphtounicode{pokatakana}{00A0}
\pdfglyphtounicode{poplathai}{00A0}
\pdfglyphtounicode{postalmark}{00A0}
\pdfglyphtounicode{postalmarkface}{00A0}
\pdfglyphtounicode{pparen}{00A0}
\pdfglyphtounicode{precedenotdbleqv}{00A0}
\pdfglyphtounicode{precedenotslnteql}{00A0}
\pdfglyphtounicode{precedeornoteqvlnt}{00A0}
\pdfglyphtounicode{precedes}{00A0}
\pdfglyphtounicode{precedesequal}{00A0}
\pdfglyphtounicode{precedesorcurly}{00A0}
\pdfglyphtounicode{precedesorequal}{00A0}
\pdfglyphtounicode{prescription}{00A0}
\pdfglyphtounicode{prime}{00A0}
\pdfglyphtounicode{primemod}{00A0}
\pdfglyphtounicode{primereverse}{00A0}
\pdfglyphtounicode{primereversed}{00A0}
\pdfglyphtounicode{product}{00A0}
\pdfglyphtounicode{projective}{00A0}
\pdfglyphtounicode{prolongedkana}{00A0}
\pdfglyphtounicode{propellor}{00A0}
\pdfglyphtounicode{propersubset}{00A0}
\pdfglyphtounicode{propersuperset}{00A0}
\pdfglyphtounicode{proportion}{00A0}
\pdfglyphtounicode{proportional}{00A0}
\pdfglyphtounicode{psi}{00A0}
\pdfglyphtounicode{psicyrillic}{00A0}
\pdfglyphtounicode{psilipneumatacyrilliccmb}{00A0}
\pdfglyphtounicode{pssquare}{00A0}
\pdfglyphtounicode{puhiragana}{00A0}
\pdfglyphtounicode{pukatakana}{00A0}
\pdfglyphtounicode{punctdash}{00A0}
\pdfglyphtounicode{pvsquare}{00A0}
\pdfglyphtounicode{pwsquare}{00A0}
\pdfglyphtounicode{q}{00A0}
\pdfglyphtounicode{qadeva}{00A0}
\pdfglyphtounicode{qadmahebrew}{00A0}
\pdfglyphtounicode{qafarabic}{00A0}
\pdfglyphtounicode{qaffinalarabic}{00A0}
\pdfglyphtounicode{qafinitialarabic}{00A0}
\pdfglyphtounicode{qafmedialarabic}{00A0}
\pdfglyphtounicode{qamats}{00A0}
\pdfglyphtounicode{qamats10}{00A0}
\pdfglyphtounicode{qamats1a}{00A0}
\pdfglyphtounicode{qamats1c}{00A0}
\pdfglyphtounicode{qamats27}{00A0}
\pdfglyphtounicode{qamats29}{00A0}
\pdfglyphtounicode{qamats33}{00A0}
\pdfglyphtounicode{qamatsde}{00A0}
\pdfglyphtounicode{qamatshebrew}{00A0}
\pdfglyphtounicode{qamatsnarrowhebrew}{00A0}
\pdfglyphtounicode{qamatsqatanhebrew}{00A0}
\pdfglyphtounicode{qamatsqatannarrowhebrew}{00A0}
\pdfglyphtounicode{qamatsqatanquarterhebrew}{00A0}
\pdfglyphtounicode{qamatsqatanwidehebrew}{00A0}
\pdfglyphtounicode{qamatsquarterhebrew}{00A0}
\pdfglyphtounicode{qamatswidehebrew}{00A0}
\pdfglyphtounicode{qarneyparahebrew}{00A0}
\pdfglyphtounicode{qbopomofo}{00A0}
\pdfglyphtounicode{qcircle}{00A0}
\pdfglyphtounicode{qhook}{00A0}
\pdfglyphtounicode{qmonospace}{00A0}
\pdfglyphtounicode{qof}{00A0}
\pdfglyphtounicode{qofdagesh}{00A0}
\pdfglyphtounicode{qofdageshhebrew}{00A0}
\pdfglyphtounicode{qofhatafpatah}{00A0}
\pdfglyphtounicode{qofhatafpatahhebrew}{00A0}
\pdfglyphtounicode{qofhatafsegol}{00A0}
\pdfglyphtounicode{qofhatafsegolhebrew}{00A0}
\pdfglyphtounicode{qofhebrew}{00A0}
\pdfglyphtounicode{qofhiriq}{00A0}
\pdfglyphtounicode{qofhiriqhebrew}{00A0}
\pdfglyphtounicode{qofholam}{00A0}
\pdfglyphtounicode{qofholamhebrew}{00A0}
\pdfglyphtounicode{qofpatah}{00A0}
\pdfglyphtounicode{qofpatahhebrew}{00A0}
\pdfglyphtounicode{qofqamats}{00A0}
\pdfglyphtounicode{qofqamatshebrew}{00A0}
\pdfglyphtounicode{qofqubuts}{00A0}
\pdfglyphtounicode{qofqubutshebrew}{00A0}
\pdfglyphtounicode{qofsegol}{00A0}
\pdfglyphtounicode{qofsegolhebrew}{00A0}
\pdfglyphtounicode{qofsheva}{00A0}
\pdfglyphtounicode{qofshevahebrew}{00A0}
\pdfglyphtounicode{qoftsere}{00A0}
\pdfglyphtounicode{qoftserehebrew}{00A0}
\pdfglyphtounicode{qparen}{00A0}
\pdfglyphtounicode{quarternote}{00A0}
\pdfglyphtounicode{qubuts}{00A0}
\pdfglyphtounicode{qubuts18}{00A0}
\pdfglyphtounicode{qubuts25}{00A0}
\pdfglyphtounicode{qubuts31}{00A0}
\pdfglyphtounicode{qubutshebrew}{00A0}
\pdfglyphtounicode{qubutsnarrowhebrew}{00A0}
\pdfglyphtounicode{qubutsquarterhebrew}{00A0}
\pdfglyphtounicode{qubutswidehebrew}{00A0}
\pdfglyphtounicode{question}{00A0}
\pdfglyphtounicode{questionarabic}{00A0}
\pdfglyphtounicode{questionarmenian}{00A0}
\pdfglyphtounicode{questiondown}{00A0}
\pdfglyphtounicode{questiondownsmall}{00A0}
\pdfglyphtounicode{questiongreek}{00A0}
\pdfglyphtounicode{questionmonospace}{00A0}
\pdfglyphtounicode{questionsmall}{00A0}
\pdfglyphtounicode{quotedbl}{00A0}
\pdfglyphtounicode{quotedblbase}{00A0}
\pdfglyphtounicode{quotedblleft}{00A0}
\pdfglyphtounicode{quotedblmonospace}{00A0}
\pdfglyphtounicode{quotedblprime}{00A0}
\pdfglyphtounicode{quotedblprimereversed}{00A0}
\pdfglyphtounicode{quotedblright}{00A0}
\pdfglyphtounicode{quoteleft}{00A0}
\pdfglyphtounicode{quoteleftreversed}{00A0}
\pdfglyphtounicode{quotereversed}{00A0}
\pdfglyphtounicode{quoteright}{00A0}
\pdfglyphtounicode{quoterightn}{00A0}
\pdfglyphtounicode{quotesinglbase}{00A0}
\pdfglyphtounicode{quotesingle}{00A0}
\pdfglyphtounicode{quotesinglemonospace}{00A0}
\pdfglyphtounicode{r}{00A0}
\pdfglyphtounicode{raarmenian}{00A0}
\pdfglyphtounicode{rabengali}{00A0}
\pdfglyphtounicode{racute}{00A0}
\pdfglyphtounicode{radeva}{00A0}
\pdfglyphtounicode{radical}{00A0}
\pdfglyphtounicode{radicalex}{00A0}
\pdfglyphtounicode{radoverssquare}{00A0}
\pdfglyphtounicode{radoverssquaredsquare}{00A0}
\pdfglyphtounicode{radsquare}{00A0}
\pdfglyphtounicode{rafe}{00A0}
\pdfglyphtounicode{rafehebrew}{00A0}
\pdfglyphtounicode{ragujarati}{00A0}
\pdfglyphtounicode{ragurmukhi}{00A0}
\pdfglyphtounicode{rahiragana}{00A0}
\pdfglyphtounicode{rakatakana}{00A0}
\pdfglyphtounicode{rakatakanahalfwidth}{00A0}
\pdfglyphtounicode{ralowerdiagonalbengali}{00A0}
\pdfglyphtounicode{ramiddlediagonalbengali}{00A0}
\pdfglyphtounicode{ramshorn}{00A0}
\pdfglyphtounicode{rangedash}{00A0}
\pdfglyphtounicode{ratio}{00A0}
\pdfglyphtounicode{rbopomofo}{00A0}
\pdfglyphtounicode{rcaron}{00A0}
\pdfglyphtounicode{rcedilla}{00A0}
\pdfglyphtounicode{rcircle}{00A0}
\pdfglyphtounicode{rcommaaccent}{00A0}
\pdfglyphtounicode{rdblgrave}{00A0}
\pdfglyphtounicode{rdotaccent}{00A0}
\pdfglyphtounicode{rdotbelow}{00A0}
\pdfglyphtounicode{rdotbelowmacron}{00A0}
\pdfglyphtounicode{referencemark}{00A0}
\pdfglyphtounicode{reflexsubset}{00A0}
\pdfglyphtounicode{reflexsuperset}{00A0}
\pdfglyphtounicode{registered}{00A0}
\pdfglyphtounicode{registersans}{00A0}
\pdfglyphtounicode{registerserif}{00A0}
\pdfglyphtounicode{reharabic}{00A0}
\pdfglyphtounicode{reharmenian}{00A0}
\pdfglyphtounicode{rehfinalarabic}{00A0}
\pdfglyphtounicode{rehiragana}{00A0}
\pdfglyphtounicode{rehyehaleflamarabic}{00A0}
\pdfglyphtounicode{rekatakana}{00A0}
\pdfglyphtounicode{rekatakanahalfwidth}{00A0}
\pdfglyphtounicode{resh}{00A0}
\pdfglyphtounicode{reshdageshhebrew}{00A0}
\pdfglyphtounicode{reshhatafpatah}{00A0}
\pdfglyphtounicode{reshhatafpatahhebrew}{00A0}
\pdfglyphtounicode{reshhatafsegol}{00A0}
\pdfglyphtounicode{reshhatafsegolhebrew}{00A0}
\pdfglyphtounicode{reshhebrew}{00A0}
\pdfglyphtounicode{reshhiriq}{00A0}
\pdfglyphtounicode{reshhiriqhebrew}{00A0}
\pdfglyphtounicode{reshholam}{00A0}
\pdfglyphtounicode{reshholamhebrew}{00A0}
\pdfglyphtounicode{reshpatah}{00A0}
\pdfglyphtounicode{reshpatahhebrew}{00A0}
\pdfglyphtounicode{reshqamats}{00A0}
\pdfglyphtounicode{reshqamatshebrew}{00A0}
\pdfglyphtounicode{reshqubuts}{00A0}
\pdfglyphtounicode{reshqubutshebrew}{00A0}
\pdfglyphtounicode{reshsegol}{00A0}
\pdfglyphtounicode{reshsegolhebrew}{00A0}
\pdfglyphtounicode{reshsheva}{00A0}
\pdfglyphtounicode{reshshevahebrew}{00A0}
\pdfglyphtounicode{reshtsere}{00A0}
\pdfglyphtounicode{reshtserehebrew}{00A0}
\pdfglyphtounicode{revasymptequal}{00A0}
\pdfglyphtounicode{reversedtilde}{00A0}
\pdfglyphtounicode{reviahebrew}{00A0}
\pdfglyphtounicode{reviamugrashhebrew}{00A0}
\pdfglyphtounicode{revlogicalnot}{00A0}
\pdfglyphtounicode{revsimilar}{00A0}
\pdfglyphtounicode{rfishhook}{00A0}
\pdfglyphtounicode{rfishhookreversed}{00A0}
\pdfglyphtounicode{rhabengali}{00A0}
\pdfglyphtounicode{rhadeva}{00A0}
\pdfglyphtounicode{rho}{00A0}
\pdfglyphtounicode{rho1}{00A0}
\pdfglyphtounicode{rhook}{00A0}
\pdfglyphtounicode{rhookturned}{00A0}
\pdfglyphtounicode{rhookturnedsuperior}{00A0}
\pdfglyphtounicode{rhosymbolgreek}{00A0}
\pdfglyphtounicode{rhotichookmod}{00A0}
\pdfglyphtounicode{rieulacirclekorean}{00A0}
\pdfglyphtounicode{rieulaparenkorean}{00A0}
\pdfglyphtounicode{rieulcirclekorean}{00A0}
\pdfglyphtounicode{rieulhieuhkorean}{00A0}
\pdfglyphtounicode{rieulkiyeokkorean}{00A0}
\pdfglyphtounicode{rieulkiyeoksioskorean}{00A0}
\pdfglyphtounicode{rieulkorean}{00A0}
\pdfglyphtounicode{rieulmieumkorean}{00A0}
\pdfglyphtounicode{rieulpansioskorean}{00A0}
\pdfglyphtounicode{rieulparenkorean}{00A0}
\pdfglyphtounicode{rieulphieuphkorean}{00A0}
\pdfglyphtounicode{rieulpieupkorean}{00A0}
\pdfglyphtounicode{rieulpieupsioskorean}{00A0}
\pdfglyphtounicode{rieulsioskorean}{00A0}
\pdfglyphtounicode{rieulthieuthkorean}{00A0}
\pdfglyphtounicode{rieultikeutkorean}{00A0}
\pdfglyphtounicode{rieulyeorinhieuhkorean}{00A0}
\pdfglyphtounicode{rightangle}{00A0}
\pdfglyphtounicode{rightanglene}{00A0}
\pdfglyphtounicode{rightanglenw}{00A0}
\pdfglyphtounicode{rightanglese}{00A0}
\pdfglyphtounicode{rightanglesw}{00A0}
\pdfglyphtounicode{righttackbelowcmb}{00A0}
\pdfglyphtounicode{righttriangle}{00A0}
\pdfglyphtounicode{rihiragana}{00A0}
\pdfglyphtounicode{rikatakana}{00A0}
\pdfglyphtounicode{rikatakanahalfwidth}{00A0}
\pdfglyphtounicode{ring}{00A0}
\pdfglyphtounicode{ringbelowcmb}{00A0}
\pdfglyphtounicode{ringcmb}{00A0}
\pdfglyphtounicode{ringhalfleft}{00A0}
\pdfglyphtounicode{ringhalfleftarmenian}{00A0}
\pdfglyphtounicode{ringhalfleftbelowcmb}{00A0}
\pdfglyphtounicode{ringhalfleftcentered}{00A0}
\pdfglyphtounicode{ringhalfright}{00A0}
\pdfglyphtounicode{ringhalfrightbelowcmb}{00A0}
\pdfglyphtounicode{ringhalfrightcentered}{00A0}
\pdfglyphtounicode{ringinequal}{00A0}
\pdfglyphtounicode{rinvertedbreve}{00A0}
\pdfglyphtounicode{rittorusquare}{00A0}
\pdfglyphtounicode{rlinebelow}{00A0}
\pdfglyphtounicode{rlongleg}{00A0}
\pdfglyphtounicode{rlonglegturned}{00A0}
\pdfglyphtounicode{rmonospace}{00A0}
\pdfglyphtounicode{rohiragana}{00A0}
\pdfglyphtounicode{rokatakana}{00A0}
\pdfglyphtounicode{rokatakanahalfwidth}{00A0}
\pdfglyphtounicode{roruathai}{00A0}
\pdfglyphtounicode{rparen}{00A0}
\pdfglyphtounicode{rrabengali}{00A0}
\pdfglyphtounicode{rradeva}{00A0}
\pdfglyphtounicode{rragurmukhi}{00A0}
\pdfglyphtounicode{rreharabic}{00A0}
\pdfglyphtounicode{rrehfinalarabic}{00A0}
\pdfglyphtounicode{rrvocalicbengali}{00A0}
\pdfglyphtounicode{rrvocalicdeva}{00A0}
\pdfglyphtounicode{rrvocalicgujarati}{00A0}
\pdfglyphtounicode{rrvocalicvowelsignbengali}{00A0}
\pdfglyphtounicode{rrvocalicvowelsigndeva}{00A0}
\pdfglyphtounicode{rrvocalicvowelsigngujarati}{00A0}
\pdfglyphtounicode{rsuperior}{00A0}
\pdfglyphtounicode{rtblock}{00A0}
\pdfglyphtounicode{rturned}{00A0}
\pdfglyphtounicode{rturnedsuperior}{00A0}
\pdfglyphtounicode{ruhiragana}{00A0}
\pdfglyphtounicode{rukatakana}{00A0}
\pdfglyphtounicode{rukatakanahalfwidth}{00A0}
\pdfglyphtounicode{rupeemarkbengali}{00A0}
\pdfglyphtounicode{rupeesignbengali}{00A0}
\pdfglyphtounicode{rupiah}{00A0}
\pdfglyphtounicode{ruthai}{00A0}
\pdfglyphtounicode{rvocalicbengali}{00A0}
\pdfglyphtounicode{rvocalicdeva}{00A0}
\pdfglyphtounicode{rvocalicgujarati}{00A0}
\pdfglyphtounicode{rvocalicvowelsignbengali}{00A0}
\pdfglyphtounicode{rvocalicvowelsigndeva}{00A0}
\pdfglyphtounicode{rvocalicvowelsigngujarati}{00A0}
\pdfglyphtounicode{s}{00A0}
\pdfglyphtounicode{sabengali}{00A0}
\pdfglyphtounicode{sacute}{00A0}
\pdfglyphtounicode{sacutedotaccent}{00A0}
\pdfglyphtounicode{sadarabic}{00A0}
\pdfglyphtounicode{sadeva}{00A0}
\pdfglyphtounicode{sadfinalarabic}{00A0}
\pdfglyphtounicode{sadinitialarabic}{00A0}
\pdfglyphtounicode{sadmedialarabic}{00A0}
\pdfglyphtounicode{sagujarati}{00A0}
\pdfglyphtounicode{sagurmukhi}{00A0}
\pdfglyphtounicode{sahiragana}{00A0}
\pdfglyphtounicode{sakatakana}{00A0}
\pdfglyphtounicode{sakatakanahalfwidth}{00A0}
\pdfglyphtounicode{sallallahoualayhewasallamarabic}{00A0}
\pdfglyphtounicode{samekh}{00A0}
\pdfglyphtounicode{samekhdagesh}{00A0}
\pdfglyphtounicode{samekhdageshhebrew}{00A0}
\pdfglyphtounicode{samekhhebrew}{00A0}
\pdfglyphtounicode{saraaathai}{00A0}
\pdfglyphtounicode{saraaethai}{00A0}
\pdfglyphtounicode{saraaimaimalaithai}{00A0}
\pdfglyphtounicode{saraaimaimuanthai}{00A0}
\pdfglyphtounicode{saraamthai}{00A0}
\pdfglyphtounicode{saraathai}{00A0}
\pdfglyphtounicode{saraethai}{00A0}
\pdfglyphtounicode{saraiileftthai}{00A0}
\pdfglyphtounicode{saraiithai}{00A0}
\pdfglyphtounicode{saraileftthai}{00A0}
\pdfglyphtounicode{saraithai}{00A0}
\pdfglyphtounicode{saraothai}{00A0}
\pdfglyphtounicode{saraueeleftthai}{00A0}
\pdfglyphtounicode{saraueethai}{00A0}
\pdfglyphtounicode{saraueleftthai}{00A0}
\pdfglyphtounicode{sarauethai}{00A0}
\pdfglyphtounicode{sarauthai}{00A0}
\pdfglyphtounicode{sarauuthai}{00A0}
\pdfglyphtounicode{satisfies}{00A0}
\pdfglyphtounicode{sbopomofo}{00A0}
\pdfglyphtounicode{scaron}{00A0}
\pdfglyphtounicode{scarondotaccent}{00A0}
\pdfglyphtounicode{scedilla}{00A0}
\pdfglyphtounicode{schwa}{00A0}
\pdfglyphtounicode{schwacyrillic}{00A0}
\pdfglyphtounicode{schwadieresiscyrillic}{00A0}
\pdfglyphtounicode{schwahook}{00A0}
\pdfglyphtounicode{scircle}{00A0}
\pdfglyphtounicode{scircumflex}{00A0}
\pdfglyphtounicode{scommaaccent}{00A0}
\pdfglyphtounicode{sdotaccent}{00A0}
\pdfglyphtounicode{sdotbelow}{00A0}
\pdfglyphtounicode{sdotbelowdotaccent}{00A0}
\pdfglyphtounicode{seagullbelowcmb}{00A0}
\pdfglyphtounicode{second}{00A0}
\pdfglyphtounicode{secondtonechinese}{00A0}
\pdfglyphtounicode{section}{00A0}
\pdfglyphtounicode{seenarabic}{00A0}
\pdfglyphtounicode{seenfinalarabic}{00A0}
\pdfglyphtounicode{seeninitialarabic}{00A0}
\pdfglyphtounicode{seenmedialarabic}{00A0}
\pdfglyphtounicode{segol}{00A0}
\pdfglyphtounicode{segol13}{00A0}
\pdfglyphtounicode{segol1f}{00A0}
\pdfglyphtounicode{segol2c}{00A0}
\pdfglyphtounicode{segolhebrew}{00A0}
\pdfglyphtounicode{segolnarrowhebrew}{00A0}
\pdfglyphtounicode{segolquarterhebrew}{00A0}
\pdfglyphtounicode{segoltahebrew}{00A0}
\pdfglyphtounicode{segolwidehebrew}{00A0}
\pdfglyphtounicode{seharmenian}{00A0}
\pdfglyphtounicode{sehiragana}{00A0}
\pdfglyphtounicode{sekatakana}{00A0}
\pdfglyphtounicode{sekatakanahalfwidth}{00A0}
\pdfglyphtounicode{semicolon}{00A0}
\pdfglyphtounicode{semicolonarabic}{00A0}
\pdfglyphtounicode{semicolonmonospace}{00A0}
\pdfglyphtounicode{semicolonsmall}{00A0}
\pdfglyphtounicode{semivoicedmarkkana}{00A0}
\pdfglyphtounicode{semivoicedmarkkanahalfwidth}{00A0}
\pdfglyphtounicode{sentisquare}{00A0}
\pdfglyphtounicode{sentosquare}{00A0}
\pdfglyphtounicode{seven}{00A0}
\pdfglyphtounicode{sevenarabic}{00A0}
\pdfglyphtounicode{sevenbengali}{00A0}
\pdfglyphtounicode{sevencircle}{00A0}
\pdfglyphtounicode{sevencircleinversesansserif}{00A0}
\pdfglyphtounicode{sevendeva}{00A0}
\pdfglyphtounicode{seveneighths}{00A0}
\pdfglyphtounicode{sevengujarati}{00A0}
\pdfglyphtounicode{sevengurmukhi}{00A0}
\pdfglyphtounicode{sevenhackarabic}{00A0}
\pdfglyphtounicode{sevenhangzhou}{00A0}
\pdfglyphtounicode{sevenideographicparen}{00A0}
\pdfglyphtounicode{seveninferior}{00A0}
\pdfglyphtounicode{sevenmonospace}{00A0}
\pdfglyphtounicode{sevenoldstyle}{00A0}
\pdfglyphtounicode{sevenparen}{00A0}
\pdfglyphtounicode{sevenperiod}{00A0}
\pdfglyphtounicode{sevenpersian}{00A0}
\pdfglyphtounicode{sevenroman}{00A0}
\pdfglyphtounicode{sevensuperior}{00A0}
\pdfglyphtounicode{seventeencircle}{00A0}
\pdfglyphtounicode{seventeenparen}{00A0}
\pdfglyphtounicode{seventeenperiod}{00A0}
\pdfglyphtounicode{seventhai}{00A0}
\pdfglyphtounicode{sfthyphen}{00A0}
\pdfglyphtounicode{shaarmenian}{00A0}
\pdfglyphtounicode{shabengali}{00A0}
\pdfglyphtounicode{shacyrillic}{00A0}
\pdfglyphtounicode{shaddaarabic}{00A0}
\pdfglyphtounicode{shaddadammaarabic}{00A0}
\pdfglyphtounicode{shaddadammatanarabic}{00A0}
\pdfglyphtounicode{shaddafathaarabic}{00A0}
\pdfglyphtounicode{shaddafathatanarabic}{00A0}
\pdfglyphtounicode{shaddakasraarabic}{00A0}
\pdfglyphtounicode{shaddakasratanarabic}{00A0}
\pdfglyphtounicode{shade}{00A0}
\pdfglyphtounicode{shadedark}{00A0}
\pdfglyphtounicode{shadelight}{00A0}
\pdfglyphtounicode{shademedium}{00A0}
\pdfglyphtounicode{shadeva}{00A0}
\pdfglyphtounicode{shagujarati}{00A0}
\pdfglyphtounicode{shagurmukhi}{00A0}
\pdfglyphtounicode{shalshelethebrew}{00A0}
\pdfglyphtounicode{sharp}{00A0}
\pdfglyphtounicode{shbopomofo}{00A0}
\pdfglyphtounicode{shchacyrillic}{00A0}
\pdfglyphtounicode{sheenarabic}{00A0}
\pdfglyphtounicode{sheenfinalarabic}{00A0}
\pdfglyphtounicode{sheeninitialarabic}{00A0}
\pdfglyphtounicode{sheenmedialarabic}{00A0}
\pdfglyphtounicode{sheicoptic}{00A0}
\pdfglyphtounicode{sheqel}{00A0}
\pdfglyphtounicode{sheqelhebrew}{00A0}
\pdfglyphtounicode{sheva}{00A0}
\pdfglyphtounicode{sheva115}{00A0}
\pdfglyphtounicode{sheva15}{00A0}
\pdfglyphtounicode{sheva22}{00A0}
\pdfglyphtounicode{sheva2e}{00A0}
\pdfglyphtounicode{shevahebrew}{00A0}
\pdfglyphtounicode{shevanarrowhebrew}{00A0}
\pdfglyphtounicode{shevaquarterhebrew}{00A0}
\pdfglyphtounicode{shevawidehebrew}{00A0}
\pdfglyphtounicode{shhacyrillic}{00A0}
\pdfglyphtounicode{shiftleft}{00A0}
\pdfglyphtounicode{shiftright}{00A0}
\pdfglyphtounicode{shimacoptic}{00A0}
\pdfglyphtounicode{shin}{00A0}
\pdfglyphtounicode{shindagesh}{00A0}
\pdfglyphtounicode{shindageshhebrew}{00A0}
\pdfglyphtounicode{shindageshshindot}{00A0}
\pdfglyphtounicode{shindageshshindothebrew}{00A0}
\pdfglyphtounicode{shindageshsindot}{00A0}
\pdfglyphtounicode{shindageshsindothebrew}{00A0}
\pdfglyphtounicode{shindothebrew}{00A0}
\pdfglyphtounicode{shinhebrew}{00A0}
\pdfglyphtounicode{shinshindot}{00A0}
\pdfglyphtounicode{shinshindothebrew}{00A0}
\pdfglyphtounicode{shinsindot}{00A0}
\pdfglyphtounicode{shinsindothebrew}{00A0}
\pdfglyphtounicode{shook}{00A0}
\pdfglyphtounicode{sigma}{00A0}
\pdfglyphtounicode{sigma1}{00A0}
\pdfglyphtounicode{sigmafinal}{00A0}
\pdfglyphtounicode{sigmalunatesymbolgreek}{00A0}
\pdfglyphtounicode{sihiragana}{00A0}
\pdfglyphtounicode{sikatakana}{00A0}
\pdfglyphtounicode{sikatakanahalfwidth}{00A0}
\pdfglyphtounicode{siluqhebrew}{00A0}
\pdfglyphtounicode{siluqlefthebrew}{00A0}
\pdfglyphtounicode{similar}{00A0}
\pdfglyphtounicode{similarequal}{00A0}
\pdfglyphtounicode{sindothebrew}{00A0}
\pdfglyphtounicode{siosacirclekorean}{00A0}
\pdfglyphtounicode{siosaparenkorean}{00A0}
\pdfglyphtounicode{sioscieuckorean}{00A0}
\pdfglyphtounicode{sioscirclekorean}{00A0}
\pdfglyphtounicode{sioskiyeokkorean}{00A0}
\pdfglyphtounicode{sioskorean}{00A0}
\pdfglyphtounicode{siosnieunkorean}{00A0}
\pdfglyphtounicode{siosparenkorean}{00A0}
\pdfglyphtounicode{siospieupkorean}{00A0}
\pdfglyphtounicode{siostikeutkorean}{00A0}
\pdfglyphtounicode{six}{00A0}
\pdfglyphtounicode{sixarabic}{00A0}
\pdfglyphtounicode{sixbengali}{00A0}
\pdfglyphtounicode{sixcircle}{00A0}
\pdfglyphtounicode{sixcircleinversesansserif}{00A0}
\pdfglyphtounicode{sixdeva}{00A0}
\pdfglyphtounicode{sixgujarati}{00A0}
\pdfglyphtounicode{sixgurmukhi}{00A0}
\pdfglyphtounicode{sixhackarabic}{00A0}
\pdfglyphtounicode{sixhangzhou}{00A0}
\pdfglyphtounicode{sixideographicparen}{00A0}
\pdfglyphtounicode{sixinferior}{00A0}
\pdfglyphtounicode{sixmonospace}{00A0}
\pdfglyphtounicode{sixoldstyle}{00A0}
\pdfglyphtounicode{sixparen}{00A0}
\pdfglyphtounicode{sixperiod}{00A0}
\pdfglyphtounicode{sixpersian}{00A0}
\pdfglyphtounicode{sixroman}{00A0}
\pdfglyphtounicode{sixsuperior}{00A0}
\pdfglyphtounicode{sixteencircle}{00A0}
\pdfglyphtounicode{sixteencurrencydenominatorbengali}{00A0}
\pdfglyphtounicode{sixteenparen}{00A0}
\pdfglyphtounicode{sixteenperiod}{00A0}
\pdfglyphtounicode{sixthai}{00A0}
\pdfglyphtounicode{slash}{00A0}
\pdfglyphtounicode{slashmonospace}{00A0}
\pdfglyphtounicode{slong}{00A0}
\pdfglyphtounicode{slongdotaccent}{00A0}
\pdfglyphtounicode{slurabove}{00A0}
\pdfglyphtounicode{slurbelow}{00A0}
\pdfglyphtounicode{smile}{00A0}
\pdfglyphtounicode{smileface}{00A0}
\pdfglyphtounicode{smonospace}{00A0}
\pdfglyphtounicode{sofpasuqhebrew}{00A0}
\pdfglyphtounicode{softhyphen}{00A0}
\pdfglyphtounicode{softsigncyrillic}{00A0}
\pdfglyphtounicode{sohiragana}{00A0}
\pdfglyphtounicode{sokatakana}{00A0}
\pdfglyphtounicode{sokatakanahalfwidth}{00A0}
\pdfglyphtounicode{soliduslongoverlaycmb}{00A0}
\pdfglyphtounicode{solidusshortoverlaycmb}{00A0}
\pdfglyphtounicode{sorusithai}{00A0}
\pdfglyphtounicode{sosalathai}{00A0}
\pdfglyphtounicode{sosothai}{00A0}
\pdfglyphtounicode{sosuathai}{00A0}
\pdfglyphtounicode{space}{00A0}
\pdfglyphtounicode{spacehackarabic}{00A0}
\pdfglyphtounicode{spade}{00A0}
\pdfglyphtounicode{spadesuitblack}{00A0}
\pdfglyphtounicode{spadesuitwhite}{00A0}
\pdfglyphtounicode{sparen}{00A0}
\pdfglyphtounicode{sphericalangle}{00A0}
\pdfglyphtounicode{square}{00A0}
\pdfglyphtounicode{squarebelowcmb}{00A0}
\pdfglyphtounicode{squarecc}{00A0}
\pdfglyphtounicode{squarecm}{00A0}
\pdfglyphtounicode{squarediagonalcrosshatchfill}{00A0}
\pdfglyphtounicode{squaredot}{00A0}
\pdfglyphtounicode{squarehorizontalfill}{00A0}
\pdfglyphtounicode{squareimage}{00A0}
\pdfglyphtounicode{squarekg}{00A0}
\pdfglyphtounicode{squarekm}{00A0}
\pdfglyphtounicode{squarekmcapital}{00A0}
\pdfglyphtounicode{squareln}{00A0}
\pdfglyphtounicode{squarelog}{00A0}
\pdfglyphtounicode{squaremg}{00A0}
\pdfglyphtounicode{squaremil}{00A0}
\pdfglyphtounicode{squareminus}{00A0}
\pdfglyphtounicode{squaremm}{00A0}
\pdfglyphtounicode{squaremsquared}{00A0}
\pdfglyphtounicode{squaremultiply}{00A0}
\pdfglyphtounicode{squareoriginal}{00A0}
\pdfglyphtounicode{squareorthogonalcrosshatchfill}{00A0}
\pdfglyphtounicode{squareplus}{00A0}
\pdfglyphtounicode{squaresolid}{00A0}
\pdfglyphtounicode{squareupperlefttolowerrightfill}{00A0}
\pdfglyphtounicode{squareupperrighttolowerleftfill}{00A0}
\pdfglyphtounicode{squareverticalfill}{00A0}
\pdfglyphtounicode{squarewhitewithsmallblack}{00A0}
\pdfglyphtounicode{squiggleleftright}{00A0}
\pdfglyphtounicode{squiggleright}{00A0}
\pdfglyphtounicode{srsquare}{00A0}
\pdfglyphtounicode{ssabengali}{00A0}
\pdfglyphtounicode{ssadeva}{00A0}
\pdfglyphtounicode{ssagujarati}{00A0}
\pdfglyphtounicode{ssangcieuckorean}{00A0}
\pdfglyphtounicode{ssanghieuhkorean}{00A0}
\pdfglyphtounicode{ssangieungkorean}{00A0}
\pdfglyphtounicode{ssangkiyeokkorean}{00A0}
\pdfglyphtounicode{ssangnieunkorean}{00A0}
\pdfglyphtounicode{ssangpieupkorean}{00A0}
\pdfglyphtounicode{ssangsioskorean}{00A0}
\pdfglyphtounicode{ssangtikeutkorean}{00A0}
\pdfglyphtounicode{ssuperior}{00A0}
\pdfglyphtounicode{st}{00A0}
\pdfglyphtounicode{star}{00A0}
\pdfglyphtounicode{sterling}{00A0}
\pdfglyphtounicode{sterlingmonospace}{00A0}
\pdfglyphtounicode{strokelongoverlaycmb}{00A0}
\pdfglyphtounicode{strokeshortoverlaycmb}{00A0}
\pdfglyphtounicode{subset}{00A0}
\pdfglyphtounicode{subsetdbl}{00A0}
\pdfglyphtounicode{subsetdblequal}{00A0}
\pdfglyphtounicode{subsetnoteql}{00A0}
\pdfglyphtounicode{subsetnotequal}{00A0}
\pdfglyphtounicode{subsetorequal}{00A0}
\pdfglyphtounicode{subsetornotdbleql}{00A0}
\pdfglyphtounicode{subsetsqequal}{00A0}
\pdfglyphtounicode{succeeds}{00A0}
\pdfglyphtounicode{suchthat}{00A0}
\pdfglyphtounicode{suhiragana}{00A0}
\pdfglyphtounicode{sukatakana}{00A0}
\pdfglyphtounicode{sukatakanahalfwidth}{00A0}
\pdfglyphtounicode{sukunarabic}{00A0}
\pdfglyphtounicode{summation}{00A0}
\pdfglyphtounicode{sun}{00A0}
\pdfglyphtounicode{superset}{00A0}
\pdfglyphtounicode{supersetdbl}{00A0}
\pdfglyphtounicode{supersetdblequal}{00A0}
\pdfglyphtounicode{supersetnoteql}{00A0}
\pdfglyphtounicode{supersetnotequal}{00A0}
\pdfglyphtounicode{supersetorequal}{00A0}
\pdfglyphtounicode{supersetornotdbleql}{00A0}
\pdfglyphtounicode{supersetsqequal}{00A0}
\pdfglyphtounicode{svsquare}{00A0}
\pdfglyphtounicode{syouwaerasquare}{00A0}
\pdfglyphtounicode{t}{00A0}
\pdfglyphtounicode{tabengali}{00A0}
\pdfglyphtounicode{tackdown}{00A0}
\pdfglyphtounicode{tackleft}{00A0}
\pdfglyphtounicode{tadeva}{00A0}
\pdfglyphtounicode{tagujarati}{00A0}
\pdfglyphtounicode{tagurmukhi}{00A0}
\pdfglyphtounicode{taharabic}{00A0}
\pdfglyphtounicode{tahfinalarabic}{00A0}
\pdfglyphtounicode{tahinitialarabic}{00A0}
\pdfglyphtounicode{tahiragana}{00A0}
\pdfglyphtounicode{tahmedialarabic}{00A0}
\pdfglyphtounicode{taisyouerasquare}{00A0}
\pdfglyphtounicode{takatakana}{00A0}
\pdfglyphtounicode{takatakanahalfwidth}{00A0}
\pdfglyphtounicode{tatweelarabic}{00A0}
\pdfglyphtounicode{tau}{00A0}
\pdfglyphtounicode{tav}{00A0}
\pdfglyphtounicode{tavdages}{00A0}
\pdfglyphtounicode{tavdagesh}{00A0}
\pdfglyphtounicode{tavdageshhebrew}{00A0}
\pdfglyphtounicode{tavhebrew}{00A0}
\pdfglyphtounicode{tbar}{00A0}
\pdfglyphtounicode{tbopomofo}{00A0}
\pdfglyphtounicode{tcaron}{00A0}
\pdfglyphtounicode{tccurl}{00A0}
\pdfglyphtounicode{tcedilla}{00A0}
\pdfglyphtounicode{tcheharabic}{00A0}
\pdfglyphtounicode{tchehfinalarabic}{00A0}
\pdfglyphtounicode{tchehinitialarabic}{00A0}
\pdfglyphtounicode{tchehmedialarabic}{00A0}
\pdfglyphtounicode{tchehmeeminitialarabic}{00A0}
\pdfglyphtounicode{tcircle}{00A0}
\pdfglyphtounicode{tcircumflexbelow}{00A0}
\pdfglyphtounicode{tcommaaccent}{00A0}
\pdfglyphtounicode{tdieresis}{00A0}
\pdfglyphtounicode{tdotaccent}{00A0}
\pdfglyphtounicode{tdotbelow}{00A0}
\pdfglyphtounicode{tecyrillic}{00A0}
\pdfglyphtounicode{tedescendercyrillic}{00A0}
\pdfglyphtounicode{teharabic}{00A0}
\pdfglyphtounicode{tehfinalarabic}{00A0}
\pdfglyphtounicode{tehhahinitialarabic}{00A0}
\pdfglyphtounicode{tehhahisolatedarabic}{00A0}
\pdfglyphtounicode{tehinitialarabic}{00A0}
\pdfglyphtounicode{tehiragana}{00A0}
\pdfglyphtounicode{tehjeeminitialarabic}{00A0}
\pdfglyphtounicode{tehjeemisolatedarabic}{00A0}
\pdfglyphtounicode{tehmarbutaarabic}{00A0}
\pdfglyphtounicode{tehmarbutafinalarabic}{00A0}
\pdfglyphtounicode{tehmedialarabic}{00A0}
\pdfglyphtounicode{tehmeeminitialarabic}{00A0}
\pdfglyphtounicode{tehmeemisolatedarabic}{00A0}
\pdfglyphtounicode{tehnoonfinalarabic}{00A0}
\pdfglyphtounicode{tekatakana}{00A0}
\pdfglyphtounicode{tekatakanahalfwidth}{00A0}
\pdfglyphtounicode{telephone}{00A0}
\pdfglyphtounicode{telephoneblack}{00A0}
\pdfglyphtounicode{telishagedolahebrew}{00A0}
\pdfglyphtounicode{telishaqetanahebrew}{00A0}
\pdfglyphtounicode{tencircle}{00A0}
\pdfglyphtounicode{tenideographicparen}{00A0}
\pdfglyphtounicode{tenparen}{00A0}
\pdfglyphtounicode{tenperiod}{00A0}
\pdfglyphtounicode{tenroman}{00A0}
\pdfglyphtounicode{tesh}{00A0}
\pdfglyphtounicode{tet}{00A0}
\pdfglyphtounicode{tetdagesh}{00A0}
\pdfglyphtounicode{tetdageshhebrew}{00A0}
\pdfglyphtounicode{tethebrew}{00A0}
\pdfglyphtounicode{tetsecyrillic}{00A0}
\pdfglyphtounicode{tevirhebrew}{00A0}
\pdfglyphtounicode{tevirlefthebrew}{00A0}
\pdfglyphtounicode{tfm:cmbsy10/diamond}{00A0}
\pdfglyphtounicode{tfm:cmbsy10/heart}{00A0}
\pdfglyphtounicode{tfm:cmbsy5/diamond}{00A0}
\pdfglyphtounicode{tfm:cmbsy5/heart}{00A0}
\pdfglyphtounicode{tfm:cmbsy6/diamond}{00A0}
\pdfglyphtounicode{tfm:cmbsy6/heart}{00A0}
\pdfglyphtounicode{tfm:cmbsy7/diamond}{00A0}
\pdfglyphtounicode{tfm:cmbsy7/heart}{00A0}
\pdfglyphtounicode{tfm:cmbsy8/diamond}{00A0}
\pdfglyphtounicode{tfm:cmbsy8/heart}{00A0}
\pdfglyphtounicode{tfm:cmbsy9/diamond}{00A0}
\pdfglyphtounicode{tfm:cmbsy9/heart}{00A0}
\pdfglyphtounicode{tfm:cmmi10/phi}{00A0}
\pdfglyphtounicode{tfm:cmmi10/phi1}{00A0}
\pdfglyphtounicode{tfm:cmmi12/phi}{00A0}
\pdfglyphtounicode{tfm:cmmi12/phi1}{00A0}
\pdfglyphtounicode{tfm:cmmi5/phi}{00A0}
\pdfglyphtounicode{tfm:cmmi5/phi1}{00A0}
\pdfglyphtounicode{tfm:cmmi6/phi}{00A0}
\pdfglyphtounicode{tfm:cmmi6/phi1}{00A0}
\pdfglyphtounicode{tfm:cmmi7/phi}{00A0}
\pdfglyphtounicode{tfm:cmmi7/phi1}{00A0}
\pdfglyphtounicode{tfm:cmmi8/phi}{00A0}
\pdfglyphtounicode{tfm:cmmi8/phi1}{00A0}
\pdfglyphtounicode{tfm:cmmi9/phi}{00A0}
\pdfglyphtounicode{tfm:cmmi9/phi1}{00A0}
\pdfglyphtounicode{tfm:cmmib10/phi}{00A0}
\pdfglyphtounicode{tfm:cmmib10/phi1}{00A0}
\pdfglyphtounicode{tfm:cmmib5/phi}{00A0}
\pdfglyphtounicode{tfm:cmmib5/phi1}{00A0}
\pdfglyphtounicode{tfm:cmmib6/phi}{00A0}
\pdfglyphtounicode{tfm:cmmib6/phi1}{00A0}
\pdfglyphtounicode{tfm:cmmib7/phi}{00A0}
\pdfglyphtounicode{tfm:cmmib7/phi1}{00A0}
\pdfglyphtounicode{tfm:cmmib8/phi}{00A0}
\pdfglyphtounicode{tfm:cmmib8/phi1}{00A0}
\pdfglyphtounicode{tfm:cmmib9/phi}{00A0}
\pdfglyphtounicode{tfm:cmmib9/phi1}{00A0}
\pdfglyphtounicode{tfm:cmsy10/diamond}{00A0}
\pdfglyphtounicode{tfm:cmsy10/heart}{00A0}
\pdfglyphtounicode{tfm:cmsy5/heart}{00A0}
\pdfglyphtounicode{tfm:cmsy6/diamond}{00A0}
\pdfglyphtounicode{tfm:cmsy6/heart}{00A0}
\pdfglyphtounicode{tfm:cmsy7/diamond}{00A0}
\pdfglyphtounicode{tfm:cmsy7/heart}{00A0}
\pdfglyphtounicode{tfm:cmsy8/diamond}{00A0}
\pdfglyphtounicode{tfm:cmsy8/heart}{00A0}
\pdfglyphtounicode{tfm:cmsy9/diamond}{00A0}
\pdfglyphtounicode{tfm:cmsy9/heart}{00A0}
\pdfglyphtounicode{tfm:eurb10/phi}{00A0}
\pdfglyphtounicode{tfm:eurb10/phi1}{00A0}
\pdfglyphtounicode{tfm:eurb5/phi}{00A0}
\pdfglyphtounicode{tfm:eurb5/phi1}{00A0}
\pdfglyphtounicode{tfm:eurb6/phi}{00A0}
\pdfglyphtounicode{tfm:eurb6/phi1}{00A0}
\pdfglyphtounicode{tfm:eurb7/phi}{00A0}
\pdfglyphtounicode{tfm:eurb7/phi1}{00A0}
\pdfglyphtounicode{tfm:eurb8/phi}{00A0}
\pdfglyphtounicode{tfm:eurb8/phi1}{00A0}
\pdfglyphtounicode{tfm:eurb9/phi}{00A0}
\pdfglyphtounicode{tfm:eurb9/phi1}{00A0}
\pdfglyphtounicode{tfm:eurm10/phi}{00A0}
\pdfglyphtounicode{tfm:eurm10/phi1}{00A0}
\pdfglyphtounicode{tfm:eurm5/phi}{00A0}
\pdfglyphtounicode{tfm:eurm5/phi1}{00A0}
\pdfglyphtounicode{tfm:eurm6/phi}{00A0}
\pdfglyphtounicode{tfm:eurm6/phi1}{00A0}
\pdfglyphtounicode{tfm:eurm7/phi}{00A0}
\pdfglyphtounicode{tfm:eurm7/phi1}{00A0}
\pdfglyphtounicode{tfm:eurm8/phi}{00A0}
\pdfglyphtounicode{tfm:eurm8/phi1}{00A0}
\pdfglyphtounicode{tfm:eurm9/phi}{00A0}
\pdfglyphtounicode{tfm:eurm9/phi1}{00A0}
\pdfglyphtounicode{tfm:fplmbi/phi}{00A0}
\pdfglyphtounicode{tfm:fplmbi/phi1}{00A0}
\pdfglyphtounicode{tfm:fplmri/phi}{00A0}
\pdfglyphtounicode{tfm:fplmri/phi1}{00A0}
\pdfglyphtounicode{tfm:lmbsy10/diamond}{00A0}
\pdfglyphtounicode{tfm:lmbsy10/heart}{00A0}
\pdfglyphtounicode{tfm:lmbsy5/diamond}{00A0}
\pdfglyphtounicode{tfm:lmbsy5/heart}{00A0}
\pdfglyphtounicode{tfm:lmbsy7/diamond}{00A0}
\pdfglyphtounicode{tfm:lmbsy7/heart}{00A0}
\pdfglyphtounicode{tfm:lmmi10/phi}{00A0}
\pdfglyphtounicode{tfm:lmmi10/phi1}{00A0}
\pdfglyphtounicode{tfm:lmmi12/phi}{00A0}
\pdfglyphtounicode{tfm:lmmi12/phi1}{00A0}
\pdfglyphtounicode{tfm:lmmi5/phi}{00A0}
\pdfglyphtounicode{tfm:lmmi5/phi1}{00A0}
\pdfglyphtounicode{tfm:lmmi6/phi}{00A0}
\pdfglyphtounicode{tfm:lmmi6/phi1}{00A0}
\pdfglyphtounicode{tfm:lmmi7/phi}{00A0}
\pdfglyphtounicode{tfm:lmmi7/phi1}{00A0}
\pdfglyphtounicode{tfm:lmmi8/phi}{00A0}
\pdfglyphtounicode{tfm:lmmi8/phi1}{00A0}
\pdfglyphtounicode{tfm:lmmi9/phi}{00A0}
\pdfglyphtounicode{tfm:lmmi9/phi1}{00A0}
\pdfglyphtounicode{tfm:lmmib10/phi}{00A0}
\pdfglyphtounicode{tfm:lmmib10/phi1}{00A0}
\pdfglyphtounicode{tfm:lmmib5/phi}{00A0}
\pdfglyphtounicode{tfm:lmmib5/phi1}{00A0}
\pdfglyphtounicode{tfm:lmmib7/phi}{00A0}
\pdfglyphtounicode{tfm:lmmib7/phi1}{00A0}
\pdfglyphtounicode{tfm:lmsy10/diamond}{00A0}
\pdfglyphtounicode{tfm:lmsy10/heart}{00A0}
\pdfglyphtounicode{tfm:lmsy5/diamond}{00A0}
\pdfglyphtounicode{tfm:lmsy5/heart}{00A0}
\pdfglyphtounicode{tfm:lmsy6/diamond}{00A0}
\pdfglyphtounicode{tfm:lmsy6/heart}{00A0}
\pdfglyphtounicode{tfm:lmsy7/diamond}{00A0}
\pdfglyphtounicode{tfm:lmsy7/heart}{00A0}
\pdfglyphtounicode{tfm:lmsy8/diamond}{00A0}
\pdfglyphtounicode{tfm:lmsy8/heart}{00A0}
\pdfglyphtounicode{tfm:lmsy9/diamond}{00A0}
\pdfglyphtounicode{tfm:lmsy9/heart}{00A0}
\pdfglyphtounicode{tfm:msam10/diamond}{00A0}
\pdfglyphtounicode{tfm:msam5/diamond}{00A0}
\pdfglyphtounicode{tfm:msam6/diamond}{00A0}
\pdfglyphtounicode{tfm:msam7/diamond}{00A0}
\pdfglyphtounicode{tfm:msam8/diamond}{00A0}
\pdfglyphtounicode{tfm:msam9/diamond}{00A0}
\pdfglyphtounicode{tfm:pxbmia/phi}{00A0}
\pdfglyphtounicode{tfm:pxbmia/phi1}{00A0}
\pdfglyphtounicode{tfm:pxbsy/diamond}{00A0}
\pdfglyphtounicode{tfm:pxbsy/heart}{00A0}
\pdfglyphtounicode{tfm:pxbsya/diamond}{00A0}
\pdfglyphtounicode{tfm:pxmia/phi}{00A0}
\pdfglyphtounicode{tfm:pxmia/phi1}{00A0}
\pdfglyphtounicode{tfm:pxsy/diamond}{00A0}
\pdfglyphtounicode{tfm:pxsy/heart}{00A0}
\pdfglyphtounicode{tfm:pxsya/diamond}{00A0}
\pdfglyphtounicode{tfm:pzdr/a1}{00A0}
\pdfglyphtounicode{tfm:pzdr/a10}{00A0}
\pdfglyphtounicode{tfm:pzdr/a100}{00A0}
\pdfglyphtounicode{tfm:pzdr/a101}{00A0}
\pdfglyphtounicode{tfm:pzdr/a102}{00A0}
\pdfglyphtounicode{tfm:pzdr/a103}{00A0}
\pdfglyphtounicode{tfm:pzdr/a104}{00A0}
\pdfglyphtounicode{tfm:pzdr/a105}{00A0}
\pdfglyphtounicode{tfm:pzdr/a106}{00A0}
\pdfglyphtounicode{tfm:pzdr/a107}{00A0}
\pdfglyphtounicode{tfm:pzdr/a108}{00A0}
\pdfglyphtounicode{tfm:pzdr/a109}{00A0}
\pdfglyphtounicode{tfm:pzdr/a11}{00A0}
\pdfglyphtounicode{tfm:pzdr/a110}{00A0}
\pdfglyphtounicode{tfm:pzdr/a111}{00A0}
\pdfglyphtounicode{tfm:pzdr/a112}{00A0}
\pdfglyphtounicode{tfm:pzdr/a117}{00A0}
\pdfglyphtounicode{tfm:pzdr/a118}{00A0}
\pdfglyphtounicode{tfm:pzdr/a119}{00A0}
\pdfglyphtounicode{tfm:pzdr/a12}{00A0}
\pdfglyphtounicode{tfm:pzdr/a120}{00A0}
\pdfglyphtounicode{tfm:pzdr/a121}{00A0}
\pdfglyphtounicode{tfm:pzdr/a122}{00A0}
\pdfglyphtounicode{tfm:pzdr/a123}{00A0}
\pdfglyphtounicode{tfm:pzdr/a124}{00A0}
\pdfglyphtounicode{tfm:pzdr/a125}{00A0}
\pdfglyphtounicode{tfm:pzdr/a126}{00A0}
\pdfglyphtounicode{tfm:pzdr/a127}{00A0}
\pdfglyphtounicode{tfm:pzdr/a128}{00A0}
\pdfglyphtounicode{tfm:pzdr/a129}{00A0}
\pdfglyphtounicode{tfm:pzdr/a13}{00A0}
\pdfglyphtounicode{tfm:pzdr/a130}{00A0}
\pdfglyphtounicode{tfm:pzdr/a131}{00A0}
\pdfglyphtounicode{tfm:pzdr/a132}{00A0}
\pdfglyphtounicode{tfm:pzdr/a133}{00A0}
\pdfglyphtounicode{tfm:pzdr/a134}{00A0}
\pdfglyphtounicode{tfm:pzdr/a135}{00A0}
\pdfglyphtounicode{tfm:pzdr/a136}{00A0}
\pdfglyphtounicode{tfm:pzdr/a137}{00A0}
\pdfglyphtounicode{tfm:pzdr/a138}{00A0}
\pdfglyphtounicode{tfm:pzdr/a139}{00A0}
\pdfglyphtounicode{tfm:pzdr/a14}{00A0}
\pdfglyphtounicode{tfm:pzdr/a140}{00A0}
\pdfglyphtounicode{tfm:pzdr/a141}{00A0}
\pdfglyphtounicode{tfm:pzdr/a142}{00A0}
\pdfglyphtounicode{tfm:pzdr/a143}{00A0}
\pdfglyphtounicode{tfm:pzdr/a144}{00A0}
\pdfglyphtounicode{tfm:pzdr/a145}{00A0}
\pdfglyphtounicode{tfm:pzdr/a146}{00A0}
\pdfglyphtounicode{tfm:pzdr/a147}{00A0}
\pdfglyphtounicode{tfm:pzdr/a148}{00A0}
\pdfglyphtounicode{tfm:pzdr/a149}{00A0}
\pdfglyphtounicode{tfm:pzdr/a15}{00A0}
\pdfglyphtounicode{tfm:pzdr/a150}{00A0}
\pdfglyphtounicode{tfm:pzdr/a151}{00A0}
\pdfglyphtounicode{tfm:pzdr/a152}{00A0}
\pdfglyphtounicode{tfm:pzdr/a153}{00A0}
\pdfglyphtounicode{tfm:pzdr/a154}{00A0}
\pdfglyphtounicode{tfm:pzdr/a155}{00A0}
\pdfglyphtounicode{tfm:pzdr/a156}{00A0}
\pdfglyphtounicode{tfm:pzdr/a157}{00A0}
\pdfglyphtounicode{tfm:pzdr/a158}{00A0}
\pdfglyphtounicode{tfm:pzdr/a159}{00A0}
\pdfglyphtounicode{tfm:pzdr/a16}{00A0}
\pdfglyphtounicode{tfm:pzdr/a160}{00A0}
\pdfglyphtounicode{tfm:pzdr/a161}{00A0}
\pdfglyphtounicode{tfm:pzdr/a162}{00A0}
\pdfglyphtounicode{tfm:pzdr/a163}{00A0}
\pdfglyphtounicode{tfm:pzdr/a164}{00A0}
\pdfglyphtounicode{tfm:pzdr/a165}{00A0}
\pdfglyphtounicode{tfm:pzdr/a166}{00A0}
\pdfglyphtounicode{tfm:pzdr/a167}{00A0}
\pdfglyphtounicode{tfm:pzdr/a168}{00A0}
\pdfglyphtounicode{tfm:pzdr/a169}{00A0}
\pdfglyphtounicode{tfm:pzdr/a17}{00A0}
\pdfglyphtounicode{tfm:pzdr/a170}{00A0}
\pdfglyphtounicode{tfm:pzdr/a171}{00A0}
\pdfglyphtounicode{tfm:pzdr/a172}{00A0}
\pdfglyphtounicode{tfm:pzdr/a173}{00A0}
\pdfglyphtounicode{tfm:pzdr/a174}{00A0}
\pdfglyphtounicode{tfm:pzdr/a175}{00A0}
\pdfglyphtounicode{tfm:pzdr/a176}{00A0}
\pdfglyphtounicode{tfm:pzdr/a177}{00A0}
\pdfglyphtounicode{tfm:pzdr/a178}{00A0}
\pdfglyphtounicode{tfm:pzdr/a179}{00A0}
\pdfglyphtounicode{tfm:pzdr/a18}{00A0}
\pdfglyphtounicode{tfm:pzdr/a180}{00A0}
\pdfglyphtounicode{tfm:pzdr/a181}{00A0}
\pdfglyphtounicode{tfm:pzdr/a182}{00A0}
\pdfglyphtounicode{tfm:pzdr/a183}{00A0}
\pdfglyphtounicode{tfm:pzdr/a184}{00A0}
\pdfglyphtounicode{tfm:pzdr/a185}{00A0}
\pdfglyphtounicode{tfm:pzdr/a186}{00A0}
\pdfglyphtounicode{tfm:pzdr/a187}{00A0}
\pdfglyphtounicode{tfm:pzdr/a188}{00A0}
\pdfglyphtounicode{tfm:pzdr/a189}{00A0}
\pdfglyphtounicode{tfm:pzdr/a19}{00A0}
\pdfglyphtounicode{tfm:pzdr/a190}{00A0}
\pdfglyphtounicode{tfm:pzdr/a191}{00A0}
\pdfglyphtounicode{tfm:pzdr/a192}{00A0}
\pdfglyphtounicode{tfm:pzdr/a193}{00A0}
\pdfglyphtounicode{tfm:pzdr/a194}{00A0}
\pdfglyphtounicode{tfm:pzdr/a195}{00A0}
\pdfglyphtounicode{tfm:pzdr/a196}{00A0}
\pdfglyphtounicode{tfm:pzdr/a197}{00A0}
\pdfglyphtounicode{tfm:pzdr/a198}{00A0}
\pdfglyphtounicode{tfm:pzdr/a199}{00A0}
\pdfglyphtounicode{tfm:pzdr/a2}{00A0}
\pdfglyphtounicode{tfm:pzdr/a20}{00A0}
\pdfglyphtounicode{tfm:pzdr/a200}{00A0}
\pdfglyphtounicode{tfm:pzdr/a201}{00A0}
\pdfglyphtounicode{tfm:pzdr/a202}{00A0}
\pdfglyphtounicode{tfm:pzdr/a203}{00A0}
\pdfglyphtounicode{tfm:pzdr/a204}{00A0}
\pdfglyphtounicode{tfm:pzdr/a205}{00A0}
\pdfglyphtounicode{tfm:pzdr/a206}{00A0}
\pdfglyphtounicode{tfm:pzdr/a21}{00A0}
\pdfglyphtounicode{tfm:pzdr/a22}{00A0}
\pdfglyphtounicode{tfm:pzdr/a23}{00A0}
\pdfglyphtounicode{tfm:pzdr/a24}{00A0}
\pdfglyphtounicode{tfm:pzdr/a25}{00A0}
\pdfglyphtounicode{tfm:pzdr/a26}{00A0}
\pdfglyphtounicode{tfm:pzdr/a27}{00A0}
\pdfglyphtounicode{tfm:pzdr/a28}{00A0}
\pdfglyphtounicode{tfm:pzdr/a29}{00A0}
\pdfglyphtounicode{tfm:pzdr/a3}{00A0}
\pdfglyphtounicode{tfm:pzdr/a30}{00A0}
\pdfglyphtounicode{tfm:pzdr/a31}{00A0}
\pdfglyphtounicode{tfm:pzdr/a32}{00A0}
\pdfglyphtounicode{tfm:pzdr/a33}{00A0}
\pdfglyphtounicode{tfm:pzdr/a34}{00A0}
\pdfglyphtounicode{tfm:pzdr/a35}{00A0}
\pdfglyphtounicode{tfm:pzdr/a36}{00A0}
\pdfglyphtounicode{tfm:pzdr/a37}{00A0}
\pdfglyphtounicode{tfm:pzdr/a38}{00A0}
\pdfglyphtounicode{tfm:pzdr/a39}{00A0}
\pdfglyphtounicode{tfm:pzdr/a4}{00A0}
\pdfglyphtounicode{tfm:pzdr/a40}{00A0}
\pdfglyphtounicode{tfm:pzdr/a41}{00A0}
\pdfglyphtounicode{tfm:pzdr/a42}{00A0}
\pdfglyphtounicode{tfm:pzdr/a43}{00A0}
\pdfglyphtounicode{tfm:pzdr/a44}{00A0}
\pdfglyphtounicode{tfm:pzdr/a45}{00A0}
\pdfglyphtounicode{tfm:pzdr/a46}{00A0}
\pdfglyphtounicode{tfm:pzdr/a47}{00A0}
\pdfglyphtounicode{tfm:pzdr/a48}{00A0}
\pdfglyphtounicode{tfm:pzdr/a49}{00A0}
\pdfglyphtounicode{tfm:pzdr/a5}{00A0}
\pdfglyphtounicode{tfm:pzdr/a50}{00A0}
\pdfglyphtounicode{tfm:pzdr/a51}{00A0}
\pdfglyphtounicode{tfm:pzdr/a52}{00A0}
\pdfglyphtounicode{tfm:pzdr/a53}{00A0}
\pdfglyphtounicode{tfm:pzdr/a54}{00A0}
\pdfglyphtounicode{tfm:pzdr/a55}{00A0}
\pdfglyphtounicode{tfm:pzdr/a56}{00A0}
\pdfglyphtounicode{tfm:pzdr/a57}{00A0}
\pdfglyphtounicode{tfm:pzdr/a58}{00A0}
\pdfglyphtounicode{tfm:pzdr/a59}{00A0}
\pdfglyphtounicode{tfm:pzdr/a6}{00A0}
\pdfglyphtounicode{tfm:pzdr/a60}{00A0}
\pdfglyphtounicode{tfm:pzdr/a61}{00A0}
\pdfglyphtounicode{tfm:pzdr/a62}{00A0}
\pdfglyphtounicode{tfm:pzdr/a63}{00A0}
\pdfglyphtounicode{tfm:pzdr/a64}{00A0}
\pdfglyphtounicode{tfm:pzdr/a65}{00A0}
\pdfglyphtounicode{tfm:pzdr/a66}{00A0}
\pdfglyphtounicode{tfm:pzdr/a67}{00A0}
\pdfglyphtounicode{tfm:pzdr/a68}{00A0}
\pdfglyphtounicode{tfm:pzdr/a69}{00A0}
\pdfglyphtounicode{tfm:pzdr/a7}{00A0}
\pdfglyphtounicode{tfm:pzdr/a70}{00A0}
\pdfglyphtounicode{tfm:pzdr/a71}{00A0}
\pdfglyphtounicode{tfm:pzdr/a72}{00A0}
\pdfglyphtounicode{tfm:pzdr/a73}{00A0}
\pdfglyphtounicode{tfm:pzdr/a74}{00A0}
\pdfglyphtounicode{tfm:pzdr/a75}{00A0}
\pdfglyphtounicode{tfm:pzdr/a76}{00A0}
\pdfglyphtounicode{tfm:pzdr/a77}{00A0}
\pdfglyphtounicode{tfm:pzdr/a78}{00A0}
\pdfglyphtounicode{tfm:pzdr/a79}{00A0}
\pdfglyphtounicode{tfm:pzdr/a8}{00A0}
\pdfglyphtounicode{tfm:pzdr/a81}{00A0}
\pdfglyphtounicode{tfm:pzdr/a82}{00A0}
\pdfglyphtounicode{tfm:pzdr/a83}{00A0}
\pdfglyphtounicode{tfm:pzdr/a84}{00A0}
\pdfglyphtounicode{tfm:pzdr/a85}{00A0}
\pdfglyphtounicode{tfm:pzdr/a86}{00A0}
\pdfglyphtounicode{tfm:pzdr/a87}{00A0}
\pdfglyphtounicode{tfm:pzdr/a88}{00A0}
\pdfglyphtounicode{tfm:pzdr/a89}{00A0}
\pdfglyphtounicode{tfm:pzdr/a9}{00A0}
\pdfglyphtounicode{tfm:pzdr/a90}{00A0}
\pdfglyphtounicode{tfm:pzdr/a91}{00A0}
\pdfglyphtounicode{tfm:pzdr/a92}{00A0}
\pdfglyphtounicode{tfm:pzdr/a93}{00A0}
\pdfglyphtounicode{tfm:pzdr/a94}{00A0}
\pdfglyphtounicode{tfm:pzdr/a95}{00A0}
\pdfglyphtounicode{tfm:pzdr/a96}{00A0}
\pdfglyphtounicode{tfm:pzdr/a97}{00A0}
\pdfglyphtounicode{tfm:pzdr/a98}{00A0}
\pdfglyphtounicode{tfm:pzdr/a99}{00A0}
\pdfglyphtounicode{tfm:rpxbmi/phi}{00A0}
\pdfglyphtounicode{tfm:rpxbmi/phi1}{00A0}
\pdfglyphtounicode{tfm:rpxmi/phi}{00A0}
\pdfglyphtounicode{tfm:rpxmi/phi1}{00A0}
\pdfglyphtounicode{tfm:rpzdr/a1}{00A0}
\pdfglyphtounicode{tfm:rpzdr/a10}{00A0}
\pdfglyphtounicode{tfm:rpzdr/a100}{00A0}
\pdfglyphtounicode{tfm:rpzdr/a101}{00A0}
\pdfglyphtounicode{tfm:rpzdr/a102}{00A0}
\pdfglyphtounicode{tfm:rpzdr/a103}{00A0}
\pdfglyphtounicode{tfm:rpzdr/a104}{00A0}
\pdfglyphtounicode{tfm:rpzdr/a105}{00A0}
\pdfglyphtounicode{tfm:rpzdr/a106}{00A0}
\pdfglyphtounicode{tfm:rpzdr/a107}{00A0}
\pdfglyphtounicode{tfm:rpzdr/a108}{00A0}
\pdfglyphtounicode{tfm:rpzdr/a109}{00A0}
\pdfglyphtounicode{tfm:rpzdr/a11}{00A0}
\pdfglyphtounicode{tfm:rpzdr/a110}{00A0}
\pdfglyphtounicode{tfm:rpzdr/a111}{00A0}
\pdfglyphtounicode{tfm:rpzdr/a112}{00A0}
\pdfglyphtounicode{tfm:rpzdr/a117}{00A0}
\pdfglyphtounicode{tfm:rpzdr/a118}{00A0}
\pdfglyphtounicode{tfm:rpzdr/a119}{00A0}
\pdfglyphtounicode{tfm:rpzdr/a12}{00A0}
\pdfglyphtounicode{tfm:rpzdr/a120}{00A0}
\pdfglyphtounicode{tfm:rpzdr/a121}{00A0}
\pdfglyphtounicode{tfm:rpzdr/a122}{00A0}
\pdfglyphtounicode{tfm:rpzdr/a123}{00A0}
\pdfglyphtounicode{tfm:rpzdr/a124}{00A0}
\pdfglyphtounicode{tfm:rpzdr/a125}{00A0}
\pdfglyphtounicode{tfm:rpzdr/a126}{00A0}
\pdfglyphtounicode{tfm:rpzdr/a127}{00A0}
\pdfglyphtounicode{tfm:rpzdr/a128}{00A0}
\pdfglyphtounicode{tfm:rpzdr/a129}{00A0}
\pdfglyphtounicode{tfm:rpzdr/a13}{00A0}
\pdfglyphtounicode{tfm:rpzdr/a130}{00A0}
\pdfglyphtounicode{tfm:rpzdr/a131}{00A0}
\pdfglyphtounicode{tfm:rpzdr/a132}{00A0}
\pdfglyphtounicode{tfm:rpzdr/a133}{00A0}
\pdfglyphtounicode{tfm:rpzdr/a134}{00A0}
\pdfglyphtounicode{tfm:rpzdr/a135}{00A0}
\pdfglyphtounicode{tfm:rpzdr/a136}{00A0}
\pdfglyphtounicode{tfm:rpzdr/a137}{00A0}
\pdfglyphtounicode{tfm:rpzdr/a138}{00A0}
\pdfglyphtounicode{tfm:rpzdr/a139}{00A0}
\pdfglyphtounicode{tfm:rpzdr/a14}{00A0}
\pdfglyphtounicode{tfm:rpzdr/a140}{00A0}
\pdfglyphtounicode{tfm:rpzdr/a141}{00A0}
\pdfglyphtounicode{tfm:rpzdr/a142}{00A0}
\pdfglyphtounicode{tfm:rpzdr/a143}{00A0}
\pdfglyphtounicode{tfm:rpzdr/a144}{00A0}
\pdfglyphtounicode{tfm:rpzdr/a145}{00A0}
\pdfglyphtounicode{tfm:rpzdr/a146}{00A0}
\pdfglyphtounicode{tfm:rpzdr/a147}{00A0}
\pdfglyphtounicode{tfm:rpzdr/a148}{00A0}
\pdfglyphtounicode{tfm:rpzdr/a149}{00A0}
\pdfglyphtounicode{tfm:rpzdr/a15}{00A0}
\pdfglyphtounicode{tfm:rpzdr/a150}{00A0}
\pdfglyphtounicode{tfm:rpzdr/a151}{00A0}
\pdfglyphtounicode{tfm:rpzdr/a152}{00A0}
\pdfglyphtounicode{tfm:rpzdr/a153}{00A0}
\pdfglyphtounicode{tfm:rpzdr/a154}{00A0}
\pdfglyphtounicode{tfm:rpzdr/a155}{00A0}
\pdfglyphtounicode{tfm:rpzdr/a156}{00A0}
\pdfglyphtounicode{tfm:rpzdr/a157}{00A0}
\pdfglyphtounicode{tfm:rpzdr/a158}{00A0}
\pdfglyphtounicode{tfm:rpzdr/a159}{00A0}
\pdfglyphtounicode{tfm:rpzdr/a16}{00A0}
\pdfglyphtounicode{tfm:rpzdr/a160}{00A0}
\pdfglyphtounicode{tfm:rpzdr/a161}{00A0}
\pdfglyphtounicode{tfm:rpzdr/a162}{00A0}
\pdfglyphtounicode{tfm:rpzdr/a163}{00A0}
\pdfglyphtounicode{tfm:rpzdr/a164}{00A0}
\pdfglyphtounicode{tfm:rpzdr/a165}{00A0}
\pdfglyphtounicode{tfm:rpzdr/a166}{00A0}
\pdfglyphtounicode{tfm:rpzdr/a167}{00A0}
\pdfglyphtounicode{tfm:rpzdr/a168}{00A0}
\pdfglyphtounicode{tfm:rpzdr/a169}{00A0}
\pdfglyphtounicode{tfm:rpzdr/a17}{00A0}
\pdfglyphtounicode{tfm:rpzdr/a170}{00A0}
\pdfglyphtounicode{tfm:rpzdr/a171}{00A0}
\pdfglyphtounicode{tfm:rpzdr/a172}{00A0}
\pdfglyphtounicode{tfm:rpzdr/a173}{00A0}
\pdfglyphtounicode{tfm:rpzdr/a174}{00A0}
\pdfglyphtounicode{tfm:rpzdr/a175}{00A0}
\pdfglyphtounicode{tfm:rpzdr/a176}{00A0}
\pdfglyphtounicode{tfm:rpzdr/a177}{00A0}
\pdfglyphtounicode{tfm:rpzdr/a178}{00A0}
\pdfglyphtounicode{tfm:rpzdr/a179}{00A0}
\pdfglyphtounicode{tfm:rpzdr/a18}{00A0}
\pdfglyphtounicode{tfm:rpzdr/a180}{00A0}
\pdfglyphtounicode{tfm:rpzdr/a181}{00A0}
\pdfglyphtounicode{tfm:rpzdr/a182}{00A0}
\pdfglyphtounicode{tfm:rpzdr/a183}{00A0}
\pdfglyphtounicode{tfm:rpzdr/a184}{00A0}
\pdfglyphtounicode{tfm:rpzdr/a185}{00A0}
\pdfglyphtounicode{tfm:rpzdr/a186}{00A0}
\pdfglyphtounicode{tfm:rpzdr/a187}{00A0}
\pdfglyphtounicode{tfm:rpzdr/a188}{00A0}
\pdfglyphtounicode{tfm:rpzdr/a189}{00A0}
\pdfglyphtounicode{tfm:rpzdr/a19}{00A0}
\pdfglyphtounicode{tfm:rpzdr/a190}{00A0}
\pdfglyphtounicode{tfm:rpzdr/a191}{00A0}
\pdfglyphtounicode{tfm:rpzdr/a192}{00A0}
\pdfglyphtounicode{tfm:rpzdr/a193}{00A0}
\pdfglyphtounicode{tfm:rpzdr/a194}{00A0}
\pdfglyphtounicode{tfm:rpzdr/a195}{00A0}
\pdfglyphtounicode{tfm:rpzdr/a196}{00A0}
\pdfglyphtounicode{tfm:rpzdr/a197}{00A0}
\pdfglyphtounicode{tfm:rpzdr/a198}{00A0}
\pdfglyphtounicode{tfm:rpzdr/a199}{00A0}
\pdfglyphtounicode{tfm:rpzdr/a2}{00A0}
\pdfglyphtounicode{tfm:rpzdr/a20}{00A0}
\pdfglyphtounicode{tfm:rpzdr/a200}{00A0}
\pdfglyphtounicode{tfm:rpzdr/a201}{00A0}
\pdfglyphtounicode{tfm:rpzdr/a202}{00A0}
\pdfglyphtounicode{tfm:rpzdr/a203}{00A0}
\pdfglyphtounicode{tfm:rpzdr/a204}{00A0}
\pdfglyphtounicode{tfm:rpzdr/a205}{00A0}
\pdfglyphtounicode{tfm:rpzdr/a206}{00A0}
\pdfglyphtounicode{tfm:rpzdr/a21}{00A0}
\pdfglyphtounicode{tfm:rpzdr/a22}{00A0}
\pdfglyphtounicode{tfm:rpzdr/a23}{00A0}
\pdfglyphtounicode{tfm:rpzdr/a24}{00A0}
\pdfglyphtounicode{tfm:rpzdr/a25}{00A0}
\pdfglyphtounicode{tfm:rpzdr/a26}{00A0}
\pdfglyphtounicode{tfm:rpzdr/a27}{00A0}
\pdfglyphtounicode{tfm:rpzdr/a28}{00A0}
\pdfglyphtounicode{tfm:rpzdr/a29}{00A0}
\pdfglyphtounicode{tfm:rpzdr/a3}{00A0}
\pdfglyphtounicode{tfm:rpzdr/a30}{00A0}
\pdfglyphtounicode{tfm:rpzdr/a31}{00A0}
\pdfglyphtounicode{tfm:rpzdr/a32}{00A0}
\pdfglyphtounicode{tfm:rpzdr/a33}{00A0}
\pdfglyphtounicode{tfm:rpzdr/a34}{00A0}
\pdfglyphtounicode{tfm:rpzdr/a35}{00A0}
\pdfglyphtounicode{tfm:rpzdr/a36}{00A0}
\pdfglyphtounicode{tfm:rpzdr/a37}{00A0}
\pdfglyphtounicode{tfm:rpzdr/a38}{00A0}
\pdfglyphtounicode{tfm:rpzdr/a39}{00A0}
\pdfglyphtounicode{tfm:rpzdr/a4}{00A0}
\pdfglyphtounicode{tfm:rpzdr/a40}{00A0}
\pdfglyphtounicode{tfm:rpzdr/a41}{00A0}
\pdfglyphtounicode{tfm:rpzdr/a42}{00A0}
\pdfglyphtounicode{tfm:rpzdr/a43}{00A0}
\pdfglyphtounicode{tfm:rpzdr/a44}{00A0}
\pdfglyphtounicode{tfm:rpzdr/a45}{00A0}
\pdfglyphtounicode{tfm:rpzdr/a46}{00A0}
\pdfglyphtounicode{tfm:rpzdr/a47}{00A0}
\pdfglyphtounicode{tfm:rpzdr/a48}{00A0}
\pdfglyphtounicode{tfm:rpzdr/a49}{00A0}
\pdfglyphtounicode{tfm:rpzdr/a5}{00A0}
\pdfglyphtounicode{tfm:rpzdr/a50}{00A0}
\pdfglyphtounicode{tfm:rpzdr/a51}{00A0}
\pdfglyphtounicode{tfm:rpzdr/a52}{00A0}
\pdfglyphtounicode{tfm:rpzdr/a53}{00A0}
\pdfglyphtounicode{tfm:rpzdr/a54}{00A0}
\pdfglyphtounicode{tfm:rpzdr/a55}{00A0}
\pdfglyphtounicode{tfm:rpzdr/a56}{00A0}
\pdfglyphtounicode{tfm:rpzdr/a57}{00A0}
\pdfglyphtounicode{tfm:rpzdr/a58}{00A0}
\pdfglyphtounicode{tfm:rpzdr/a59}{00A0}
\pdfglyphtounicode{tfm:rpzdr/a6}{00A0}
\pdfglyphtounicode{tfm:rpzdr/a60}{00A0}
\pdfglyphtounicode{tfm:rpzdr/a61}{00A0}
\pdfglyphtounicode{tfm:rpzdr/a62}{00A0}
\pdfglyphtounicode{tfm:rpzdr/a63}{00A0}
\pdfglyphtounicode{tfm:rpzdr/a64}{00A0}
\pdfglyphtounicode{tfm:rpzdr/a65}{00A0}
\pdfglyphtounicode{tfm:rpzdr/a66}{00A0}
\pdfglyphtounicode{tfm:rpzdr/a67}{00A0}
\pdfglyphtounicode{tfm:rpzdr/a68}{00A0}
\pdfglyphtounicode{tfm:rpzdr/a69}{00A0}
\pdfglyphtounicode{tfm:rpzdr/a7}{00A0}
\pdfglyphtounicode{tfm:rpzdr/a70}{00A0}
\pdfglyphtounicode{tfm:rpzdr/a71}{00A0}
\pdfglyphtounicode{tfm:rpzdr/a72}{00A0}
\pdfglyphtounicode{tfm:rpzdr/a73}{00A0}
\pdfglyphtounicode{tfm:rpzdr/a74}{00A0}
\pdfglyphtounicode{tfm:rpzdr/a75}{00A0}
\pdfglyphtounicode{tfm:rpzdr/a76}{00A0}
\pdfglyphtounicode{tfm:rpzdr/a77}{00A0}
\pdfglyphtounicode{tfm:rpzdr/a78}{00A0}
\pdfglyphtounicode{tfm:rpzdr/a79}{00A0}
\pdfglyphtounicode{tfm:rpzdr/a8}{00A0}
\pdfglyphtounicode{tfm:rpzdr/a81}{00A0}
\pdfglyphtounicode{tfm:rpzdr/a82}{00A0}
\pdfglyphtounicode{tfm:rpzdr/a83}{00A0}
\pdfglyphtounicode{tfm:rpzdr/a84}{00A0}
\pdfglyphtounicode{tfm:rpzdr/a85}{00A0}
\pdfglyphtounicode{tfm:rpzdr/a86}{00A0}
\pdfglyphtounicode{tfm:rpzdr/a87}{00A0}
\pdfglyphtounicode{tfm:rpzdr/a88}{00A0}
\pdfglyphtounicode{tfm:rpzdr/a89}{00A0}
\pdfglyphtounicode{tfm:rpzdr/a9}{00A0}
\pdfglyphtounicode{tfm:rpzdr/a90}{00A0}
\pdfglyphtounicode{tfm:rpzdr/a91}{00A0}
\pdfglyphtounicode{tfm:rpzdr/a92}{00A0}
\pdfglyphtounicode{tfm:rpzdr/a93}{00A0}
\pdfglyphtounicode{tfm:rpzdr/a94}{00A0}
\pdfglyphtounicode{tfm:rpzdr/a95}{00A0}
\pdfglyphtounicode{tfm:rpzdr/a96}{00A0}
\pdfglyphtounicode{tfm:rpzdr/a97}{00A0}
\pdfglyphtounicode{tfm:rpzdr/a98}{00A0}
\pdfglyphtounicode{tfm:rpzdr/a99}{00A0}
\pdfglyphtounicode{tfm:rtxbmi/phi}{00A0}
\pdfglyphtounicode{tfm:rtxbmi/phi1}{00A0}
\pdfglyphtounicode{tfm:rtxmi/phi}{00A0}
\pdfglyphtounicode{tfm:rtxmi/phi1}{00A0}
\pdfglyphtounicode{tfm:txbmia/phi}{00A0}
\pdfglyphtounicode{tfm:txbmia/phi1}{00A0}
\pdfglyphtounicode{tfm:txbsy/diamond}{00A0}
\pdfglyphtounicode{tfm:txbsy/heart}{00A0}
\pdfglyphtounicode{tfm:txbsya/diamond}{00A0}
\pdfglyphtounicode{tfm:txmia/phi}{00A0}
\pdfglyphtounicode{tfm:txmia/phi1}{00A0}
\pdfglyphtounicode{tfm:txsy/diamond}{00A0}
\pdfglyphtounicode{tfm:txsy/heart}{00A0}
\pdfglyphtounicode{tfm:txsya/diamond}{00A0}
\pdfglyphtounicode{tfm:zd/a1}{00A0}
\pdfglyphtounicode{tfm:zd/a10}{00A0}
\pdfglyphtounicode{tfm:zd/a100}{00A0}
\pdfglyphtounicode{tfm:zd/a101}{00A0}
\pdfglyphtounicode{tfm:zd/a102}{00A0}
\pdfglyphtounicode{tfm:zd/a103}{00A0}
\pdfglyphtounicode{tfm:zd/a104}{00A0}
\pdfglyphtounicode{tfm:zd/a105}{00A0}
\pdfglyphtounicode{tfm:zd/a106}{00A0}
\pdfglyphtounicode{tfm:zd/a107}{00A0}
\pdfglyphtounicode{tfm:zd/a108}{00A0}
\pdfglyphtounicode{tfm:zd/a109}{00A0}
\pdfglyphtounicode{tfm:zd/a11}{00A0}
\pdfglyphtounicode{tfm:zd/a110}{00A0}
\pdfglyphtounicode{tfm:zd/a111}{00A0}
\pdfglyphtounicode{tfm:zd/a112}{00A0}
\pdfglyphtounicode{tfm:zd/a117}{00A0}
\pdfglyphtounicode{tfm:zd/a118}{00A0}
\pdfglyphtounicode{tfm:zd/a119}{00A0}
\pdfglyphtounicode{tfm:zd/a12}{00A0}
\pdfglyphtounicode{tfm:zd/a120}{00A0}
\pdfglyphtounicode{tfm:zd/a121}{00A0}
\pdfglyphtounicode{tfm:zd/a122}{00A0}
\pdfglyphtounicode{tfm:zd/a123}{00A0}
\pdfglyphtounicode{tfm:zd/a124}{00A0}
\pdfglyphtounicode{tfm:zd/a125}{00A0}
\pdfglyphtounicode{tfm:zd/a126}{00A0}
\pdfglyphtounicode{tfm:zd/a127}{00A0}
\pdfglyphtounicode{tfm:zd/a128}{00A0}
\pdfglyphtounicode{tfm:zd/a129}{00A0}
\pdfglyphtounicode{tfm:zd/a13}{00A0}
\pdfglyphtounicode{tfm:zd/a130}{00A0}
\pdfglyphtounicode{tfm:zd/a131}{00A0}
\pdfglyphtounicode{tfm:zd/a132}{00A0}
\pdfglyphtounicode{tfm:zd/a133}{00A0}
\pdfglyphtounicode{tfm:zd/a134}{00A0}
\pdfglyphtounicode{tfm:zd/a135}{00A0}
\pdfglyphtounicode{tfm:zd/a136}{00A0}
\pdfglyphtounicode{tfm:zd/a137}{00A0}
\pdfglyphtounicode{tfm:zd/a138}{00A0}
\pdfglyphtounicode{tfm:zd/a139}{00A0}
\pdfglyphtounicode{tfm:zd/a14}{00A0}
\pdfglyphtounicode{tfm:zd/a140}{00A0}
\pdfglyphtounicode{tfm:zd/a141}{00A0}
\pdfglyphtounicode{tfm:zd/a142}{00A0}
\pdfglyphtounicode{tfm:zd/a143}{00A0}
\pdfglyphtounicode{tfm:zd/a144}{00A0}
\pdfglyphtounicode{tfm:zd/a145}{00A0}
\pdfglyphtounicode{tfm:zd/a146}{00A0}
\pdfglyphtounicode{tfm:zd/a147}{00A0}
\pdfglyphtounicode{tfm:zd/a148}{00A0}
\pdfglyphtounicode{tfm:zd/a149}{00A0}
\pdfglyphtounicode{tfm:zd/a15}{00A0}
\pdfglyphtounicode{tfm:zd/a150}{00A0}
\pdfglyphtounicode{tfm:zd/a151}{00A0}
\pdfglyphtounicode{tfm:zd/a152}{00A0}
\pdfglyphtounicode{tfm:zd/a153}{00A0}
\pdfglyphtounicode{tfm:zd/a154}{00A0}
\pdfglyphtounicode{tfm:zd/a155}{00A0}
\pdfglyphtounicode{tfm:zd/a156}{00A0}
\pdfglyphtounicode{tfm:zd/a157}{00A0}
\pdfglyphtounicode{tfm:zd/a158}{00A0}
\pdfglyphtounicode{tfm:zd/a159}{00A0}
\pdfglyphtounicode{tfm:zd/a16}{00A0}
\pdfglyphtounicode{tfm:zd/a160}{00A0}
\pdfglyphtounicode{tfm:zd/a161}{00A0}
\pdfglyphtounicode{tfm:zd/a162}{00A0}
\pdfglyphtounicode{tfm:zd/a163}{00A0}
\pdfglyphtounicode{tfm:zd/a164}{00A0}
\pdfglyphtounicode{tfm:zd/a165}{00A0}
\pdfglyphtounicode{tfm:zd/a166}{00A0}
\pdfglyphtounicode{tfm:zd/a167}{00A0}
\pdfglyphtounicode{tfm:zd/a168}{00A0}
\pdfglyphtounicode{tfm:zd/a169}{00A0}
\pdfglyphtounicode{tfm:zd/a17}{00A0}
\pdfglyphtounicode{tfm:zd/a170}{00A0}
\pdfglyphtounicode{tfm:zd/a171}{00A0}
\pdfglyphtounicode{tfm:zd/a172}{00A0}
\pdfglyphtounicode{tfm:zd/a173}{00A0}
\pdfglyphtounicode{tfm:zd/a174}{00A0}
\pdfglyphtounicode{tfm:zd/a175}{00A0}
\pdfglyphtounicode{tfm:zd/a176}{00A0}
\pdfglyphtounicode{tfm:zd/a177}{00A0}
\pdfglyphtounicode{tfm:zd/a178}{00A0}
\pdfglyphtounicode{tfm:zd/a179}{00A0}
\pdfglyphtounicode{tfm:zd/a18}{00A0}
\pdfglyphtounicode{tfm:zd/a180}{00A0}
\pdfglyphtounicode{tfm:zd/a181}{00A0}
\pdfglyphtounicode{tfm:zd/a182}{00A0}
\pdfglyphtounicode{tfm:zd/a183}{00A0}
\pdfglyphtounicode{tfm:zd/a184}{00A0}
\pdfglyphtounicode{tfm:zd/a185}{00A0}
\pdfglyphtounicode{tfm:zd/a186}{00A0}
\pdfglyphtounicode{tfm:zd/a187}{00A0}
\pdfglyphtounicode{tfm:zd/a188}{00A0}
\pdfglyphtounicode{tfm:zd/a189}{00A0}
\pdfglyphtounicode{tfm:zd/a19}{00A0}
\pdfglyphtounicode{tfm:zd/a190}{00A0}
\pdfglyphtounicode{tfm:zd/a191}{00A0}
\pdfglyphtounicode{tfm:zd/a192}{00A0}
\pdfglyphtounicode{tfm:zd/a193}{00A0}
\pdfglyphtounicode{tfm:zd/a194}{00A0}
\pdfglyphtounicode{tfm:zd/a195}{00A0}
\pdfglyphtounicode{tfm:zd/a196}{00A0}
\pdfglyphtounicode{tfm:zd/a197}{00A0}
\pdfglyphtounicode{tfm:zd/a198}{00A0}
\pdfglyphtounicode{tfm:zd/a199}{00A0}
\pdfglyphtounicode{tfm:zd/a2}{00A0}
\pdfglyphtounicode{tfm:zd/a20}{00A0}
\pdfglyphtounicode{tfm:zd/a200}{00A0}
\pdfglyphtounicode{tfm:zd/a201}{00A0}
\pdfglyphtounicode{tfm:zd/a202}{00A0}
\pdfglyphtounicode{tfm:zd/a203}{00A0}
\pdfglyphtounicode{tfm:zd/a204}{00A0}
\pdfglyphtounicode{tfm:zd/a205}{00A0}
\pdfglyphtounicode{tfm:zd/a206}{00A0}
\pdfglyphtounicode{tfm:zd/a21}{00A0}
\pdfglyphtounicode{tfm:zd/a22}{00A0}
\pdfglyphtounicode{tfm:zd/a23}{00A0}
\pdfglyphtounicode{tfm:zd/a24}{00A0}
\pdfglyphtounicode{tfm:zd/a25}{00A0}
\pdfglyphtounicode{tfm:zd/a26}{00A0}
\pdfglyphtounicode{tfm:zd/a27}{00A0}
\pdfglyphtounicode{tfm:zd/a28}{00A0}
\pdfglyphtounicode{tfm:zd/a29}{00A0}
\pdfglyphtounicode{tfm:zd/a3}{00A0}
\pdfglyphtounicode{tfm:zd/a30}{00A0}
\pdfglyphtounicode{tfm:zd/a31}{00A0}
\pdfglyphtounicode{tfm:zd/a32}{00A0}
\pdfglyphtounicode{tfm:zd/a33}{00A0}
\pdfglyphtounicode{tfm:zd/a34}{00A0}
\pdfglyphtounicode{tfm:zd/a35}{00A0}
\pdfglyphtounicode{tfm:zd/a36}{00A0}
\pdfglyphtounicode{tfm:zd/a37}{00A0}
\pdfglyphtounicode{tfm:zd/a38}{00A0}
\pdfglyphtounicode{tfm:zd/a39}{00A0}
\pdfglyphtounicode{tfm:zd/a4}{00A0}
\pdfglyphtounicode{tfm:zd/a40}{00A0}
\pdfglyphtounicode{tfm:zd/a41}{00A0}
\pdfglyphtounicode{tfm:zd/a42}{00A0}
\pdfglyphtounicode{tfm:zd/a43}{00A0}
\pdfglyphtounicode{tfm:zd/a44}{00A0}
\pdfglyphtounicode{tfm:zd/a45}{00A0}
\pdfglyphtounicode{tfm:zd/a46}{00A0}
\pdfglyphtounicode{tfm:zd/a47}{00A0}
\pdfglyphtounicode{tfm:zd/a48}{00A0}
\pdfglyphtounicode{tfm:zd/a49}{00A0}
\pdfglyphtounicode{tfm:zd/a5}{00A0}
\pdfglyphtounicode{tfm:zd/a50}{00A0}
\pdfglyphtounicode{tfm:zd/a51}{00A0}
\pdfglyphtounicode{tfm:zd/a52}{00A0}
\pdfglyphtounicode{tfm:zd/a53}{00A0}
\pdfglyphtounicode{tfm:zd/a54}{00A0}
\pdfglyphtounicode{tfm:zd/a55}{00A0}
\pdfglyphtounicode{tfm:zd/a56}{00A0}
\pdfglyphtounicode{tfm:zd/a57}{00A0}
\pdfglyphtounicode{tfm:zd/a58}{00A0}
\pdfglyphtounicode{tfm:zd/a59}{00A0}
\pdfglyphtounicode{tfm:zd/a6}{00A0}
\pdfglyphtounicode{tfm:zd/a60}{00A0}
\pdfglyphtounicode{tfm:zd/a61}{00A0}
\pdfglyphtounicode{tfm:zd/a62}{00A0}
\pdfglyphtounicode{tfm:zd/a63}{00A0}
\pdfglyphtounicode{tfm:zd/a64}{00A0}
\pdfglyphtounicode{tfm:zd/a65}{00A0}
\pdfglyphtounicode{tfm:zd/a66}{00A0}
\pdfglyphtounicode{tfm:zd/a67}{00A0}
\pdfglyphtounicode{tfm:zd/a68}{00A0}
\pdfglyphtounicode{tfm:zd/a69}{00A0}
\pdfglyphtounicode{tfm:zd/a7}{00A0}
\pdfglyphtounicode{tfm:zd/a70}{00A0}
\pdfglyphtounicode{tfm:zd/a71}{00A0}
\pdfglyphtounicode{tfm:zd/a72}{00A0}
\pdfglyphtounicode{tfm:zd/a73}{00A0}
\pdfglyphtounicode{tfm:zd/a74}{00A0}
\pdfglyphtounicode{tfm:zd/a75}{00A0}
\pdfglyphtounicode{tfm:zd/a76}{00A0}
\pdfglyphtounicode{tfm:zd/a77}{00A0}
\pdfglyphtounicode{tfm:zd/a78}{00A0}
\pdfglyphtounicode{tfm:zd/a79}{00A0}
\pdfglyphtounicode{tfm:zd/a8}{00A0}
\pdfglyphtounicode{tfm:zd/a81}{00A0}
\pdfglyphtounicode{tfm:zd/a82}{00A0}
\pdfglyphtounicode{tfm:zd/a83}{00A0}
\pdfglyphtounicode{tfm:zd/a84}{00A0}
\pdfglyphtounicode{tfm:zd/a85}{00A0}
\pdfglyphtounicode{tfm:zd/a86}{00A0}
\pdfglyphtounicode{tfm:zd/a87}{00A0}
\pdfglyphtounicode{tfm:zd/a88}{00A0}
\pdfglyphtounicode{tfm:zd/a89}{00A0}
\pdfglyphtounicode{tfm:zd/a9}{00A0}
\pdfglyphtounicode{tfm:zd/a90}{00A0}
\pdfglyphtounicode{tfm:zd/a91}{00A0}
\pdfglyphtounicode{tfm:zd/a92}{00A0}
\pdfglyphtounicode{tfm:zd/a93}{00A0}
\pdfglyphtounicode{tfm:zd/a94}{00A0}
\pdfglyphtounicode{tfm:zd/a95}{00A0}
\pdfglyphtounicode{tfm:zd/a96}{00A0}
\pdfglyphtounicode{tfm:zd/a97}{00A0}
\pdfglyphtounicode{tfm:zd/a98}{00A0}
\pdfglyphtounicode{tfm:zd/a99}{00A0}
\pdfglyphtounicode{tfm:zpzdr-reversed/a1}{00A0}
\pdfglyphtounicode{tfm:zpzdr-reversed/a10}{00A0}
\pdfglyphtounicode{tfm:zpzdr-reversed/a100}{00A0}
\pdfglyphtounicode{tfm:zpzdr-reversed/a101}{00A0}
\pdfglyphtounicode{tfm:zpzdr-reversed/a102}{00A0}
\pdfglyphtounicode{tfm:zpzdr-reversed/a103}{00A0}
\pdfglyphtounicode{tfm:zpzdr-reversed/a104}{00A0}
\pdfglyphtounicode{tfm:zpzdr-reversed/a105}{00A0}
\pdfglyphtounicode{tfm:zpzdr-reversed/a106}{00A0}
\pdfglyphtounicode{tfm:zpzdr-reversed/a107}{00A0}
\pdfglyphtounicode{tfm:zpzdr-reversed/a108}{00A0}
\pdfglyphtounicode{tfm:zpzdr-reversed/a109}{00A0}
\pdfglyphtounicode{tfm:zpzdr-reversed/a11}{00A0}
\pdfglyphtounicode{tfm:zpzdr-reversed/a110}{00A0}
\pdfglyphtounicode{tfm:zpzdr-reversed/a111}{00A0}
\pdfglyphtounicode{tfm:zpzdr-reversed/a112}{00A0}
\pdfglyphtounicode{tfm:zpzdr-reversed/a117}{00A0}
\pdfglyphtounicode{tfm:zpzdr-reversed/a118}{00A0}
\pdfglyphtounicode{tfm:zpzdr-reversed/a119}{00A0}
\pdfglyphtounicode{tfm:zpzdr-reversed/a12}{00A0}
\pdfglyphtounicode{tfm:zpzdr-reversed/a120}{00A0}
\pdfglyphtounicode{tfm:zpzdr-reversed/a121}{00A0}
\pdfglyphtounicode{tfm:zpzdr-reversed/a122}{00A0}
\pdfglyphtounicode{tfm:zpzdr-reversed/a123}{00A0}
\pdfglyphtounicode{tfm:zpzdr-reversed/a124}{00A0}
\pdfglyphtounicode{tfm:zpzdr-reversed/a125}{00A0}
\pdfglyphtounicode{tfm:zpzdr-reversed/a126}{00A0}
\pdfglyphtounicode{tfm:zpzdr-reversed/a127}{00A0}
\pdfglyphtounicode{tfm:zpzdr-reversed/a128}{00A0}
\pdfglyphtounicode{tfm:zpzdr-reversed/a129}{00A0}
\pdfglyphtounicode{tfm:zpzdr-reversed/a13}{00A0}
\pdfglyphtounicode{tfm:zpzdr-reversed/a130}{00A0}
\pdfglyphtounicode{tfm:zpzdr-reversed/a131}{00A0}
\pdfglyphtounicode{tfm:zpzdr-reversed/a132}{00A0}
\pdfglyphtounicode{tfm:zpzdr-reversed/a133}{00A0}
\pdfglyphtounicode{tfm:zpzdr-reversed/a134}{00A0}
\pdfglyphtounicode{tfm:zpzdr-reversed/a135}{00A0}
\pdfglyphtounicode{tfm:zpzdr-reversed/a136}{00A0}
\pdfglyphtounicode{tfm:zpzdr-reversed/a137}{00A0}
\pdfglyphtounicode{tfm:zpzdr-reversed/a138}{00A0}
\pdfglyphtounicode{tfm:zpzdr-reversed/a139}{00A0}
\pdfglyphtounicode{tfm:zpzdr-reversed/a14}{00A0}
\pdfglyphtounicode{tfm:zpzdr-reversed/a140}{00A0}
\pdfglyphtounicode{tfm:zpzdr-reversed/a141}{00A0}
\pdfglyphtounicode{tfm:zpzdr-reversed/a142}{00A0}
\pdfglyphtounicode{tfm:zpzdr-reversed/a143}{00A0}
\pdfglyphtounicode{tfm:zpzdr-reversed/a144}{00A0}
\pdfglyphtounicode{tfm:zpzdr-reversed/a145}{00A0}
\pdfglyphtounicode{tfm:zpzdr-reversed/a146}{00A0}
\pdfglyphtounicode{tfm:zpzdr-reversed/a147}{00A0}
\pdfglyphtounicode{tfm:zpzdr-reversed/a148}{00A0}
\pdfglyphtounicode{tfm:zpzdr-reversed/a149}{00A0}
\pdfglyphtounicode{tfm:zpzdr-reversed/a15}{00A0}
\pdfglyphtounicode{tfm:zpzdr-reversed/a150}{00A0}
\pdfglyphtounicode{tfm:zpzdr-reversed/a151}{00A0}
\pdfglyphtounicode{tfm:zpzdr-reversed/a152}{00A0}
\pdfglyphtounicode{tfm:zpzdr-reversed/a153}{00A0}
\pdfglyphtounicode{tfm:zpzdr-reversed/a154}{00A0}
\pdfglyphtounicode{tfm:zpzdr-reversed/a155}{00A0}
\pdfglyphtounicode{tfm:zpzdr-reversed/a156}{00A0}
\pdfglyphtounicode{tfm:zpzdr-reversed/a157}{00A0}
\pdfglyphtounicode{tfm:zpzdr-reversed/a158}{00A0}
\pdfglyphtounicode{tfm:zpzdr-reversed/a159}{00A0}
\pdfglyphtounicode{tfm:zpzdr-reversed/a16}{00A0}
\pdfglyphtounicode{tfm:zpzdr-reversed/a160}{00A0}
\pdfglyphtounicode{tfm:zpzdr-reversed/a161}{00A0}
\pdfglyphtounicode{tfm:zpzdr-reversed/a162}{00A0}
\pdfglyphtounicode{tfm:zpzdr-reversed/a163}{00A0}
\pdfglyphtounicode{tfm:zpzdr-reversed/a164}{00A0}
\pdfglyphtounicode{tfm:zpzdr-reversed/a165}{00A0}
\pdfglyphtounicode{tfm:zpzdr-reversed/a166}{00A0}
\pdfglyphtounicode{tfm:zpzdr-reversed/a167}{00A0}
\pdfglyphtounicode{tfm:zpzdr-reversed/a168}{00A0}
\pdfglyphtounicode{tfm:zpzdr-reversed/a169}{00A0}
\pdfglyphtounicode{tfm:zpzdr-reversed/a17}{00A0}
\pdfglyphtounicode{tfm:zpzdr-reversed/a170}{00A0}
\pdfglyphtounicode{tfm:zpzdr-reversed/a171}{00A0}
\pdfglyphtounicode{tfm:zpzdr-reversed/a172}{00A0}
\pdfglyphtounicode{tfm:zpzdr-reversed/a173}{00A0}
\pdfglyphtounicode{tfm:zpzdr-reversed/a174}{00A0}
\pdfglyphtounicode{tfm:zpzdr-reversed/a175}{00A0}
\pdfglyphtounicode{tfm:zpzdr-reversed/a176}{00A0}
\pdfglyphtounicode{tfm:zpzdr-reversed/a177}{00A0}
\pdfglyphtounicode{tfm:zpzdr-reversed/a178}{00A0}
\pdfglyphtounicode{tfm:zpzdr-reversed/a179}{00A0}
\pdfglyphtounicode{tfm:zpzdr-reversed/a18}{00A0}
\pdfglyphtounicode{tfm:zpzdr-reversed/a180}{00A0}
\pdfglyphtounicode{tfm:zpzdr-reversed/a181}{00A0}
\pdfglyphtounicode{tfm:zpzdr-reversed/a182}{00A0}
\pdfglyphtounicode{tfm:zpzdr-reversed/a183}{00A0}
\pdfglyphtounicode{tfm:zpzdr-reversed/a184}{00A0}
\pdfglyphtounicode{tfm:zpzdr-reversed/a185}{00A0}
\pdfglyphtounicode{tfm:zpzdr-reversed/a186}{00A0}
\pdfglyphtounicode{tfm:zpzdr-reversed/a187}{00A0}
\pdfglyphtounicode{tfm:zpzdr-reversed/a188}{00A0}
\pdfglyphtounicode{tfm:zpzdr-reversed/a189}{00A0}
\pdfglyphtounicode{tfm:zpzdr-reversed/a19}{00A0}
\pdfglyphtounicode{tfm:zpzdr-reversed/a190}{00A0}
\pdfglyphtounicode{tfm:zpzdr-reversed/a191}{00A0}
\pdfglyphtounicode{tfm:zpzdr-reversed/a192}{00A0}
\pdfglyphtounicode{tfm:zpzdr-reversed/a193}{00A0}
\pdfglyphtounicode{tfm:zpzdr-reversed/a194}{00A0}
\pdfglyphtounicode{tfm:zpzdr-reversed/a195}{00A0}
\pdfglyphtounicode{tfm:zpzdr-reversed/a196}{00A0}
\pdfglyphtounicode{tfm:zpzdr-reversed/a197}{00A0}
\pdfglyphtounicode{tfm:zpzdr-reversed/a198}{00A0}
\pdfglyphtounicode{tfm:zpzdr-reversed/a199}{00A0}
\pdfglyphtounicode{tfm:zpzdr-reversed/a2}{00A0}
\pdfglyphtounicode{tfm:zpzdr-reversed/a20}{00A0}
\pdfglyphtounicode{tfm:zpzdr-reversed/a200}{00A0}
\pdfglyphtounicode{tfm:zpzdr-reversed/a201}{00A0}
\pdfglyphtounicode{tfm:zpzdr-reversed/a202}{00A0}
\pdfglyphtounicode{tfm:zpzdr-reversed/a203}{00A0}
\pdfglyphtounicode{tfm:zpzdr-reversed/a204}{00A0}
\pdfglyphtounicode{tfm:zpzdr-reversed/a205}{00A0}
\pdfglyphtounicode{tfm:zpzdr-reversed/a206}{00A0}
\pdfglyphtounicode{tfm:zpzdr-reversed/a21}{00A0}
\pdfglyphtounicode{tfm:zpzdr-reversed/a22}{00A0}
\pdfglyphtounicode{tfm:zpzdr-reversed/a23}{00A0}
\pdfglyphtounicode{tfm:zpzdr-reversed/a24}{00A0}
\pdfglyphtounicode{tfm:zpzdr-reversed/a25}{00A0}
\pdfglyphtounicode{tfm:zpzdr-reversed/a26}{00A0}
\pdfglyphtounicode{tfm:zpzdr-reversed/a27}{00A0}
\pdfglyphtounicode{tfm:zpzdr-reversed/a28}{00A0}
\pdfglyphtounicode{tfm:zpzdr-reversed/a29}{00A0}
\pdfglyphtounicode{tfm:zpzdr-reversed/a3}{00A0}
\pdfglyphtounicode{tfm:zpzdr-reversed/a30}{00A0}
\pdfglyphtounicode{tfm:zpzdr-reversed/a31}{00A0}
\pdfglyphtounicode{tfm:zpzdr-reversed/a32}{00A0}
\pdfglyphtounicode{tfm:zpzdr-reversed/a33}{00A0}
\pdfglyphtounicode{tfm:zpzdr-reversed/a34}{00A0}
\pdfglyphtounicode{tfm:zpzdr-reversed/a35}{00A0}
\pdfglyphtounicode{tfm:zpzdr-reversed/a36}{00A0}
\pdfglyphtounicode{tfm:zpzdr-reversed/a37}{00A0}
\pdfglyphtounicode{tfm:zpzdr-reversed/a38}{00A0}
\pdfglyphtounicode{tfm:zpzdr-reversed/a39}{00A0}
\pdfglyphtounicode{tfm:zpzdr-reversed/a4}{00A0}
\pdfglyphtounicode{tfm:zpzdr-reversed/a40}{00A0}
\pdfglyphtounicode{tfm:zpzdr-reversed/a41}{00A0}
\pdfglyphtounicode{tfm:zpzdr-reversed/a42}{00A0}
\pdfglyphtounicode{tfm:zpzdr-reversed/a43}{00A0}
\pdfglyphtounicode{tfm:zpzdr-reversed/a44}{00A0}
\pdfglyphtounicode{tfm:zpzdr-reversed/a45}{00A0}
\pdfglyphtounicode{tfm:zpzdr-reversed/a46}{00A0}
\pdfglyphtounicode{tfm:zpzdr-reversed/a47}{00A0}
\pdfglyphtounicode{tfm:zpzdr-reversed/a48}{00A0}
\pdfglyphtounicode{tfm:zpzdr-reversed/a49}{00A0}
\pdfglyphtounicode{tfm:zpzdr-reversed/a5}{00A0}
\pdfglyphtounicode{tfm:zpzdr-reversed/a50}{00A0}
\pdfglyphtounicode{tfm:zpzdr-reversed/a51}{00A0}
\pdfglyphtounicode{tfm:zpzdr-reversed/a52}{00A0}
\pdfglyphtounicode{tfm:zpzdr-reversed/a53}{00A0}
\pdfglyphtounicode{tfm:zpzdr-reversed/a54}{00A0}
\pdfglyphtounicode{tfm:zpzdr-reversed/a55}{00A0}
\pdfglyphtounicode{tfm:zpzdr-reversed/a56}{00A0}
\pdfglyphtounicode{tfm:zpzdr-reversed/a57}{00A0}
\pdfglyphtounicode{tfm:zpzdr-reversed/a58}{00A0}
\pdfglyphtounicode{tfm:zpzdr-reversed/a59}{00A0}
\pdfglyphtounicode{tfm:zpzdr-reversed/a6}{00A0}
\pdfglyphtounicode{tfm:zpzdr-reversed/a60}{00A0}
\pdfglyphtounicode{tfm:zpzdr-reversed/a61}{00A0}
\pdfglyphtounicode{tfm:zpzdr-reversed/a62}{00A0}
\pdfglyphtounicode{tfm:zpzdr-reversed/a63}{00A0}
\pdfglyphtounicode{tfm:zpzdr-reversed/a64}{00A0}
\pdfglyphtounicode{tfm:zpzdr-reversed/a65}{00A0}
\pdfglyphtounicode{tfm:zpzdr-reversed/a66}{00A0}
\pdfglyphtounicode{tfm:zpzdr-reversed/a67}{00A0}
\pdfglyphtounicode{tfm:zpzdr-reversed/a68}{00A0}
\pdfglyphtounicode{tfm:zpzdr-reversed/a69}{00A0}
\pdfglyphtounicode{tfm:zpzdr-reversed/a7}{00A0}
\pdfglyphtounicode{tfm:zpzdr-reversed/a70}{00A0}
\pdfglyphtounicode{tfm:zpzdr-reversed/a71}{00A0}
\pdfglyphtounicode{tfm:zpzdr-reversed/a72}{00A0}
\pdfglyphtounicode{tfm:zpzdr-reversed/a73}{00A0}
\pdfglyphtounicode{tfm:zpzdr-reversed/a74}{00A0}
\pdfglyphtounicode{tfm:zpzdr-reversed/a75}{00A0}
\pdfglyphtounicode{tfm:zpzdr-reversed/a76}{00A0}
\pdfglyphtounicode{tfm:zpzdr-reversed/a77}{00A0}
\pdfglyphtounicode{tfm:zpzdr-reversed/a78}{00A0}
\pdfglyphtounicode{tfm:zpzdr-reversed/a79}{00A0}
\pdfglyphtounicode{tfm:zpzdr-reversed/a8}{00A0}
\pdfglyphtounicode{tfm:zpzdr-reversed/a81}{00A0}
\pdfglyphtounicode{tfm:zpzdr-reversed/a82}{00A0}
\pdfglyphtounicode{tfm:zpzdr-reversed/a83}{00A0}
\pdfglyphtounicode{tfm:zpzdr-reversed/a84}{00A0}
\pdfglyphtounicode{tfm:zpzdr-reversed/a85}{00A0}
\pdfglyphtounicode{tfm:zpzdr-reversed/a86}{00A0}
\pdfglyphtounicode{tfm:zpzdr-reversed/a87}{00A0}
\pdfglyphtounicode{tfm:zpzdr-reversed/a88}{00A0}
\pdfglyphtounicode{tfm:zpzdr-reversed/a89}{00A0}
\pdfglyphtounicode{tfm:zpzdr-reversed/a9}{00A0}
\pdfglyphtounicode{tfm:zpzdr-reversed/a90}{00A0}
\pdfglyphtounicode{tfm:zpzdr-reversed/a91}{00A0}
\pdfglyphtounicode{tfm:zpzdr-reversed/a92}{00A0}
\pdfglyphtounicode{tfm:zpzdr-reversed/a93}{00A0}
\pdfglyphtounicode{tfm:zpzdr-reversed/a94}{00A0}
\pdfglyphtounicode{tfm:zpzdr-reversed/a95}{00A0}
\pdfglyphtounicode{tfm:zpzdr-reversed/a96}{00A0}
\pdfglyphtounicode{tfm:zpzdr-reversed/a97}{00A0}
\pdfglyphtounicode{tfm:zpzdr-reversed/a98}{00A0}
\pdfglyphtounicode{tfm:zpzdr-reversed/a99}{00A0}
\pdfglyphtounicode{thabengali}{00A0}
\pdfglyphtounicode{thadeva}{00A0}
\pdfglyphtounicode{thagujarati}{00A0}
\pdfglyphtounicode{thagurmukhi}{00A0}
\pdfglyphtounicode{thalarabic}{00A0}
\pdfglyphtounicode{thalfinalarabic}{00A0}
\pdfglyphtounicode{thanthakhatlowleftthai}{00A0}
\pdfglyphtounicode{thanthakhatlowrightthai}{00A0}
\pdfglyphtounicode{thanthakhatthai}{00A0}
\pdfglyphtounicode{thanthakhatupperleftthai}{00A0}
\pdfglyphtounicode{theharabic}{00A0}
\pdfglyphtounicode{thehfinalarabic}{00A0}
\pdfglyphtounicode{thehinitialarabic}{00A0}
\pdfglyphtounicode{thehmedialarabic}{00A0}
\pdfglyphtounicode{thereexists}{00A0}
\pdfglyphtounicode{therefore}{00A0}
\pdfglyphtounicode{theta}{00A0}
\pdfglyphtounicode{theta1}{00A0}
\pdfglyphtounicode{thetasymbolgreek}{00A0}
\pdfglyphtounicode{thieuthacirclekorean}{00A0}
\pdfglyphtounicode{thieuthaparenkorean}{00A0}
\pdfglyphtounicode{thieuthcirclekorean}{00A0}
\pdfglyphtounicode{thieuthkorean}{00A0}
\pdfglyphtounicode{thieuthparenkorean}{00A0}
\pdfglyphtounicode{thirteencircle}{00A0}
\pdfglyphtounicode{thirteenparen}{00A0}
\pdfglyphtounicode{thirteenperiod}{00A0}
\pdfglyphtounicode{thonangmonthothai}{00A0}
\pdfglyphtounicode{thook}{00A0}
\pdfglyphtounicode{thophuthaothai}{00A0}
\pdfglyphtounicode{thorn}{00A0}
\pdfglyphtounicode{thothahanthai}{00A0}
\pdfglyphtounicode{thothanthai}{00A0}
\pdfglyphtounicode{thothongthai}{00A0}
\pdfglyphtounicode{thothungthai}{00A0}
\pdfglyphtounicode{thousandcyrillic}{00A0}
\pdfglyphtounicode{thousandsseparatorarabic}{00A0}
\pdfglyphtounicode{thousandsseparatorpersian}{00A0}
\pdfglyphtounicode{three}{00A0}
\pdfglyphtounicode{threearabic}{00A0}
\pdfglyphtounicode{threebengali}{00A0}
\pdfglyphtounicode{threecircle}{00A0}
\pdfglyphtounicode{threecircleinversesansserif}{00A0}
\pdfglyphtounicode{threedeva}{00A0}
\pdfglyphtounicode{threeeighths}{00A0}
\pdfglyphtounicode{threegujarati}{00A0}
\pdfglyphtounicode{threegurmukhi}{00A0}
\pdfglyphtounicode{threehackarabic}{00A0}
\pdfglyphtounicode{threehangzhou}{00A0}
\pdfglyphtounicode{threeideographicparen}{00A0}
\pdfglyphtounicode{threeinferior}{00A0}
\pdfglyphtounicode{threemonospace}{00A0}
\pdfglyphtounicode{threenumeratorbengali}{00A0}
\pdfglyphtounicode{threeoldstyle}{00A0}
\pdfglyphtounicode{threeparen}{00A0}
\pdfglyphtounicode{threeperiod}{00A0}
\pdfglyphtounicode{threepersian}{00A0}
\pdfglyphtounicode{threequarters}{00A0}
\pdfglyphtounicode{threequartersemdash}{00A0}
\pdfglyphtounicode{threeroman}{00A0}
\pdfglyphtounicode{threesuperior}{00A0}
\pdfglyphtounicode{threethai}{00A0}
\pdfglyphtounicode{thzsquare}{00A0}
\pdfglyphtounicode{tihiragana}{00A0}
\pdfglyphtounicode{tikatakana}{00A0}
\pdfglyphtounicode{tikatakanahalfwidth}{00A0}
\pdfglyphtounicode{tikeutacirclekorean}{00A0}
\pdfglyphtounicode{tikeutaparenkorean}{00A0}
\pdfglyphtounicode{tikeutcirclekorean}{00A0}
\pdfglyphtounicode{tikeutkorean}{00A0}
\pdfglyphtounicode{tikeutparenkorean}{00A0}
\pdfglyphtounicode{tilde}{00A0}
\pdfglyphtounicode{tildebelowcmb}{00A0}
\pdfglyphtounicode{tildecmb}{00A0}
\pdfglyphtounicode{tildecomb}{00A0}
\pdfglyphtounicode{tildedoublecmb}{00A0}
\pdfglyphtounicode{tildeoperator}{00A0}
\pdfglyphtounicode{tildeoverlaycmb}{00A0}
\pdfglyphtounicode{tildeverticalcmb}{00A0}
\pdfglyphtounicode{timescircle}{00A0}
\pdfglyphtounicode{tipehahebrew}{00A0}
\pdfglyphtounicode{tipehalefthebrew}{00A0}
\pdfglyphtounicode{tippigurmukhi}{00A0}
\pdfglyphtounicode{titlocyrilliccmb}{00A0}
\pdfglyphtounicode{tiwnarmenian}{00A0}
\pdfglyphtounicode{tlinebelow}{00A0}
\pdfglyphtounicode{tmonospace}{00A0}
\pdfglyphtounicode{toarmenian}{00A0}
\pdfglyphtounicode{tohiragana}{00A0}
\pdfglyphtounicode{tokatakana}{00A0}
\pdfglyphtounicode{tokatakanahalfwidth}{00A0}
\pdfglyphtounicode{tonebarextrahighmod}{00A0}
\pdfglyphtounicode{tonebarextralowmod}{00A0}
\pdfglyphtounicode{tonebarhighmod}{00A0}
\pdfglyphtounicode{tonebarlowmod}{00A0}
\pdfglyphtounicode{tonebarmidmod}{00A0}
\pdfglyphtounicode{tonefive}{00A0}
\pdfglyphtounicode{tonesix}{00A0}
\pdfglyphtounicode{tonetwo}{00A0}
\pdfglyphtounicode{tonos}{00A0}
\pdfglyphtounicode{tonsquare}{00A0}
\pdfglyphtounicode{topatakthai}{00A0}
\pdfglyphtounicode{tortoiseshellbracketleft}{00A0}
\pdfglyphtounicode{tortoiseshellbracketleftsmall}{00A0}
\pdfglyphtounicode{tortoiseshellbracketleftvertical}{00A0}
\pdfglyphtounicode{tortoiseshellbracketright}{00A0}
\pdfglyphtounicode{tortoiseshellbracketrightsmall}{00A0}
\pdfglyphtounicode{tortoiseshellbracketrightvertical}{00A0}
\pdfglyphtounicode{totaothai}{00A0}
\pdfglyphtounicode{tpalatalhook}{00A0}
\pdfglyphtounicode{tparen}{00A0}
\pdfglyphtounicode{trademark}{00A0}
\pdfglyphtounicode{trademarksans}{00A0}
\pdfglyphtounicode{trademarkserif}{00A0}
\pdfglyphtounicode{tretroflexhook}{00A0}
\pdfglyphtounicode{triagdn}{00A0}
\pdfglyphtounicode{triaglf}{00A0}
\pdfglyphtounicode{triagrt}{00A0}
\pdfglyphtounicode{triagup}{00A0}
\pdfglyphtounicode{triangle}{00A0}
\pdfglyphtounicode{triangledownsld}{00A0}
\pdfglyphtounicode{triangleinv}{00A0}
\pdfglyphtounicode{triangleleft}{00A0}
\pdfglyphtounicode{triangleleftequal}{00A0}
\pdfglyphtounicode{triangleleftsld}{00A0}
\pdfglyphtounicode{triangleright}{00A0}
\pdfglyphtounicode{trianglerightequal}{00A0}
\pdfglyphtounicode{trianglerightsld}{00A0}
\pdfglyphtounicode{trianglesolid}{00A0}
\pdfglyphtounicode{ts}{00A0}
\pdfglyphtounicode{tsadi}{00A0}
\pdfglyphtounicode{tsadidagesh}{00A0}
\pdfglyphtounicode{tsadidageshhebrew}{00A0}
\pdfglyphtounicode{tsadihebrew}{00A0}
\pdfglyphtounicode{tsecyrillic}{00A0}
\pdfglyphtounicode{tsere}{00A0}
\pdfglyphtounicode{tsere12}{00A0}
\pdfglyphtounicode{tsere1e}{00A0}
\pdfglyphtounicode{tsere2b}{00A0}
\pdfglyphtounicode{tserehebrew}{00A0}
\pdfglyphtounicode{tserenarrowhebrew}{00A0}
\pdfglyphtounicode{tserequarterhebrew}{00A0}
\pdfglyphtounicode{tserewidehebrew}{00A0}
\pdfglyphtounicode{tshecyrillic}{00A0}
\pdfglyphtounicode{tsuperior}{00A0}
\pdfglyphtounicode{ttabengali}{00A0}
\pdfglyphtounicode{ttadeva}{00A0}
\pdfglyphtounicode{ttagujarati}{00A0}
\pdfglyphtounicode{ttagurmukhi}{00A0}
\pdfglyphtounicode{tteharabic}{00A0}
\pdfglyphtounicode{ttehfinalarabic}{00A0}
\pdfglyphtounicode{ttehinitialarabic}{00A0}
\pdfglyphtounicode{ttehmedialarabic}{00A0}
\pdfglyphtounicode{tthabengali}{00A0}
\pdfglyphtounicode{tthadeva}{00A0}
\pdfglyphtounicode{tthagujarati}{00A0}
\pdfglyphtounicode{tthagurmukhi}{00A0}
\pdfglyphtounicode{tturned}{00A0}
\pdfglyphtounicode{tuhiragana}{00A0}
\pdfglyphtounicode{tukatakana}{00A0}
\pdfglyphtounicode{tukatakanahalfwidth}{00A0}
\pdfglyphtounicode{turnstileleft}{00A0}
\pdfglyphtounicode{turnstileright}{00A0}
\pdfglyphtounicode{tusmallhiragana}{00A0}
\pdfglyphtounicode{tusmallkatakana}{00A0}
\pdfglyphtounicode{tusmallkatakanahalfwidth}{00A0}
\pdfglyphtounicode{twelvecircle}{00A0}
\pdfglyphtounicode{twelveparen}{00A0}
\pdfglyphtounicode{twelveperiod}{00A0}
\pdfglyphtounicode{twelveroman}{00A0}
\pdfglyphtounicode{twentycircle}{00A0}
\pdfglyphtounicode{twentyhangzhou}{00A0}
\pdfglyphtounicode{twentyparen}{00A0}
\pdfglyphtounicode{twentyperiod}{00A0}
\pdfglyphtounicode{two}{00A0}
\pdfglyphtounicode{twoarabic}{00A0}
\pdfglyphtounicode{twobengali}{00A0}
\pdfglyphtounicode{twocircle}{00A0}
\pdfglyphtounicode{twocircleinversesansserif}{00A0}
\pdfglyphtounicode{twodeva}{00A0}
\pdfglyphtounicode{twodotenleader}{00A0}
\pdfglyphtounicode{twodotleader}{00A0}
\pdfglyphtounicode{twodotleadervertical}{00A0}
\pdfglyphtounicode{twogujarati}{00A0}
\pdfglyphtounicode{twogurmukhi}{00A0}
\pdfglyphtounicode{twohackarabic}{00A0}
\pdfglyphtounicode{twohangzhou}{00A0}
\pdfglyphtounicode{twoideographicparen}{00A0}
\pdfglyphtounicode{twoinferior}{00A0}
\pdfglyphtounicode{twomonospace}{00A0}
\pdfglyphtounicode{twonumeratorbengali}{00A0}
\pdfglyphtounicode{twooldstyle}{00A0}
\pdfglyphtounicode{twoparen}{00A0}
\pdfglyphtounicode{twoperiod}{00A0}
\pdfglyphtounicode{twopersian}{00A0}
\pdfglyphtounicode{tworoman}{00A0}
\pdfglyphtounicode{twostroke}{00A0}
\pdfglyphtounicode{twosuperior}{00A0}
\pdfglyphtounicode{twothai}{00A0}
\pdfglyphtounicode{twothirds}{00A0}
\pdfglyphtounicode{u}{00A0}
\pdfglyphtounicode{uacute}{00A0}
\pdfglyphtounicode{ubar}{00A0}
\pdfglyphtounicode{ubengali}{00A0}
\pdfglyphtounicode{ubopomofo}{00A0}
\pdfglyphtounicode{ubreve}{00A0}
\pdfglyphtounicode{ucaron}{00A0}
\pdfglyphtounicode{ucircle}{00A0}
\pdfglyphtounicode{ucircumflex}{00A0}
\pdfglyphtounicode{ucircumflexbelow}{00A0}
\pdfglyphtounicode{ucyrillic}{00A0}
\pdfglyphtounicode{udattadeva}{00A0}
\pdfglyphtounicode{udblacute}{00A0}
\pdfglyphtounicode{udblgrave}{00A0}
\pdfglyphtounicode{udeva}{00A0}
\pdfglyphtounicode{udieresis}{00A0}
\pdfglyphtounicode{udieresisacute}{00A0}
\pdfglyphtounicode{udieresisbelow}{00A0}
\pdfglyphtounicode{udieresiscaron}{00A0}
\pdfglyphtounicode{udieresiscyrillic}{00A0}
\pdfglyphtounicode{udieresisgrave}{00A0}
\pdfglyphtounicode{udieresismacron}{00A0}
\pdfglyphtounicode{udotbelow}{00A0}
\pdfglyphtounicode{ugrave}{00A0}
\pdfglyphtounicode{ugujarati}{00A0}
\pdfglyphtounicode{ugurmukhi}{00A0}
\pdfglyphtounicode{uhiragana}{00A0}
\pdfglyphtounicode{uhookabove}{00A0}
\pdfglyphtounicode{uhorn}{00A0}
\pdfglyphtounicode{uhornacute}{00A0}
\pdfglyphtounicode{uhorndotbelow}{00A0}
\pdfglyphtounicode{uhorngrave}{00A0}
\pdfglyphtounicode{uhornhookabove}{00A0}
\pdfglyphtounicode{uhorntilde}{00A0}
\pdfglyphtounicode{uhungarumlaut}{00A0}
\pdfglyphtounicode{uhungarumlautcyrillic}{00A0}
\pdfglyphtounicode{uinvertedbreve}{00A0}
\pdfglyphtounicode{ukatakana}{00A0}
\pdfglyphtounicode{ukatakanahalfwidth}{00A0}
\pdfglyphtounicode{ukcyrillic}{00A0}
\pdfglyphtounicode{ukorean}{00A0}
\pdfglyphtounicode{umacron}{00A0}
\pdfglyphtounicode{umacroncyrillic}{00A0}
\pdfglyphtounicode{umacrondieresis}{00A0}
\pdfglyphtounicode{umatragurmukhi}{00A0}
\pdfglyphtounicode{umonospace}{00A0}
\pdfglyphtounicode{underscore}{00A0}
\pdfglyphtounicode{underscoredbl}{00A0}
\pdfglyphtounicode{underscoremonospace}{00A0}
\pdfglyphtounicode{underscorevertical}{00A0}
\pdfglyphtounicode{underscorewavy}{00A0}
\pdfglyphtounicode{union}{00A0}
\pdfglyphtounicode{uniondbl}{00A0}
\pdfglyphtounicode{unionmulti}{00A0}
\pdfglyphtounicode{unionsq}{00A0}
\pdfglyphtounicode{universal}{00A0}
\pdfglyphtounicode{uogonek}{00A0}
\pdfglyphtounicode{uparen}{00A0}
\pdfglyphtounicode{upblock}{00A0}
\pdfglyphtounicode{upperdothebrew}{00A0}
\pdfglyphtounicode{uprise}{00A0}
\pdfglyphtounicode{upsilon}{00A0}
\pdfglyphtounicode{upsilondieresis}{00A0}
\pdfglyphtounicode{upsilondieresistonos}{00A0}
\pdfglyphtounicode{upsilonlatin}{00A0}
\pdfglyphtounicode{upsilontonos}{00A0}
\pdfglyphtounicode{upslope}{00A0}
\pdfglyphtounicode{uptackbelowcmb}{00A0}
\pdfglyphtounicode{uptackmod}{00A0}
\pdfglyphtounicode{uragurmukhi}{00A0}
\pdfglyphtounicode{uring}{00A0}
\pdfglyphtounicode{ushortcyrillic}{00A0}
\pdfglyphtounicode{usmallhiragana}{00A0}
\pdfglyphtounicode{usmallkatakana}{00A0}
\pdfglyphtounicode{usmallkatakanahalfwidth}{00A0}
\pdfglyphtounicode{ustraightcyrillic}{00A0}
\pdfglyphtounicode{ustraightstrokecyrillic}{00A0}
\pdfglyphtounicode{utilde}{00A0}
\pdfglyphtounicode{utildeacute}{00A0}
\pdfglyphtounicode{utildebelow}{00A0}
\pdfglyphtounicode{uubengali}{00A0}
\pdfglyphtounicode{uudeva}{00A0}
\pdfglyphtounicode{uugujarati}{00A0}
\pdfglyphtounicode{uugurmukhi}{00A0}
\pdfglyphtounicode{uumatragurmukhi}{00A0}
\pdfglyphtounicode{uuvowelsignbengali}{00A0}
\pdfglyphtounicode{uuvowelsigndeva}{00A0}
\pdfglyphtounicode{uuvowelsigngujarati}{00A0}
\pdfglyphtounicode{uvowelsignbengali}{00A0}
\pdfglyphtounicode{uvowelsigndeva}{00A0}
\pdfglyphtounicode{uvowelsigngujarati}{00A0}
\pdfglyphtounicode{v}{00A0}
\pdfglyphtounicode{vadeva}{00A0}
\pdfglyphtounicode{vagujarati}{00A0}
\pdfglyphtounicode{vagurmukhi}{00A0}
\pdfglyphtounicode{vakatakana}{00A0}
\pdfglyphtounicode{vav}{00A0}
\pdfglyphtounicode{vavdagesh}{00A0}
\pdfglyphtounicode{vavdagesh65}{00A0}
\pdfglyphtounicode{vavdageshhebrew}{00A0}
\pdfglyphtounicode{vavhebrew}{00A0}
\pdfglyphtounicode{vavholam}{00A0}
\pdfglyphtounicode{vavholamhebrew}{00A0}
\pdfglyphtounicode{vavvavhebrew}{00A0}
\pdfglyphtounicode{vavyodhebrew}{00A0}
\pdfglyphtounicode{vcircle}{00A0}
\pdfglyphtounicode{vdotbelow}{00A0}
\pdfglyphtounicode{vector}{00A0}
\pdfglyphtounicode{vecyrillic}{00A0}
\pdfglyphtounicode{veharabic}{00A0}
\pdfglyphtounicode{vehfinalarabic}{00A0}
\pdfglyphtounicode{vehinitialarabic}{00A0}
\pdfglyphtounicode{vehmedialarabic}{00A0}
\pdfglyphtounicode{vekatakana}{00A0}
\pdfglyphtounicode{venus}{00A0}
\pdfglyphtounicode{verticalbar}{00A0}
\pdfglyphtounicode{verticallineabovecmb}{00A0}
\pdfglyphtounicode{verticallinebelowcmb}{00A0}
\pdfglyphtounicode{verticallinelowmod}{00A0}
\pdfglyphtounicode{verticallinemod}{00A0}
\pdfglyphtounicode{vewarmenian}{00A0}
\pdfglyphtounicode{vhook}{00A0}
\pdfglyphtounicode{vikatakana}{00A0}
\pdfglyphtounicode{viramabengali}{00A0}
\pdfglyphtounicode{viramadeva}{00A0}
\pdfglyphtounicode{viramagujarati}{00A0}
\pdfglyphtounicode{visargabengali}{00A0}
\pdfglyphtounicode{visargadeva}{00A0}
\pdfglyphtounicode{visargagujarati}{00A0}
\pdfglyphtounicode{visiblespace}{00A0}
\pdfglyphtounicode{visualspace}{00A0}
\pdfglyphtounicode{vmonospace}{00A0}
\pdfglyphtounicode{voarmenian}{00A0}
\pdfglyphtounicode{voicediterationhiragana}{00A0}
\pdfglyphtounicode{voicediterationkatakana}{00A0}
\pdfglyphtounicode{voicedmarkkana}{00A0}
\pdfglyphtounicode{voicedmarkkanahalfwidth}{00A0}
\pdfglyphtounicode{vokatakana}{00A0}
\pdfglyphtounicode{vparen}{00A0}
\pdfglyphtounicode{vtilde}{00A0}
\pdfglyphtounicode{vturned}{00A0}
\pdfglyphtounicode{vuhiragana}{00A0}
\pdfglyphtounicode{vukatakana}{00A0}
\pdfglyphtounicode{w}{00A0}
\pdfglyphtounicode{wacute}{00A0}
\pdfglyphtounicode{waekorean}{00A0}
\pdfglyphtounicode{wahiragana}{00A0}
\pdfglyphtounicode{wakatakana}{00A0}
\pdfglyphtounicode{wakatakanahalfwidth}{00A0}
\pdfglyphtounicode{wakorean}{00A0}
\pdfglyphtounicode{wasmallhiragana}{00A0}
\pdfglyphtounicode{wasmallkatakana}{00A0}
\pdfglyphtounicode{wattosquare}{00A0}
\pdfglyphtounicode{wavedash}{00A0}
\pdfglyphtounicode{wavyunderscorevertical}{00A0}
\pdfglyphtounicode{wawarabic}{00A0}
\pdfglyphtounicode{wawfinalarabic}{00A0}
\pdfglyphtounicode{wawhamzaabovearabic}{00A0}
\pdfglyphtounicode{wawhamzaabovefinalarabic}{00A0}
\pdfglyphtounicode{wbsquare}{00A0}
\pdfglyphtounicode{wcircle}{00A0}
\pdfglyphtounicode{wcircumflex}{00A0}
\pdfglyphtounicode{wdieresis}{00A0}
\pdfglyphtounicode{wdotaccent}{00A0}
\pdfglyphtounicode{wdotbelow}{00A0}
\pdfglyphtounicode{wehiragana}{00A0}
\pdfglyphtounicode{weierstrass}{00A0}
\pdfglyphtounicode{wekatakana}{00A0}
\pdfglyphtounicode{wekorean}{00A0}
\pdfglyphtounicode{weokorean}{00A0}
\pdfglyphtounicode{wgrave}{00A0}
\pdfglyphtounicode{whitebullet}{00A0}
\pdfglyphtounicode{whitecircle}{00A0}
\pdfglyphtounicode{whitecircleinverse}{00A0}
\pdfglyphtounicode{whitecornerbracketleft}{00A0}
\pdfglyphtounicode{whitecornerbracketleftvertical}{00A0}
\pdfglyphtounicode{whitecornerbracketright}{00A0}
\pdfglyphtounicode{whitecornerbracketrightvertical}{00A0}
\pdfglyphtounicode{whitediamond}{00A0}
\pdfglyphtounicode{whitediamondcontainingblacksmalldiamond}{00A0}
\pdfglyphtounicode{whitedownpointingsmalltriangle}{00A0}
\pdfglyphtounicode{whitedownpointingtriangle}{00A0}
\pdfglyphtounicode{whiteleftpointingsmalltriangle}{00A0}
\pdfglyphtounicode{whiteleftpointingtriangle}{00A0}
\pdfglyphtounicode{whitelenticularbracketleft}{00A0}
\pdfglyphtounicode{whitelenticularbracketright}{00A0}
\pdfglyphtounicode{whiterightpointingsmalltriangle}{00A0}
\pdfglyphtounicode{whiterightpointingtriangle}{00A0}
\pdfglyphtounicode{whitesmallsquare}{00A0}
\pdfglyphtounicode{whitesmilingface}{00A0}
\pdfglyphtounicode{whitesquare}{00A0}
\pdfglyphtounicode{whitestar}{00A0}
\pdfglyphtounicode{whitetelephone}{00A0}
\pdfglyphtounicode{whitetortoiseshellbracketleft}{00A0}
\pdfglyphtounicode{whitetortoiseshellbracketright}{00A0}
\pdfglyphtounicode{whiteuppointingsmalltriangle}{00A0}
\pdfglyphtounicode{whiteuppointingtriangle}{00A0}
\pdfglyphtounicode{wihiragana}{00A0}
\pdfglyphtounicode{wikatakana}{00A0}
\pdfglyphtounicode{wikorean}{00A0}
\pdfglyphtounicode{wmonospace}{00A0}
\pdfglyphtounicode{wohiragana}{00A0}
\pdfglyphtounicode{wokatakana}{00A0}
\pdfglyphtounicode{wokatakanahalfwidth}{00A0}
\pdfglyphtounicode{won}{00A0}
\pdfglyphtounicode{wonmonospace}{00A0}
\pdfglyphtounicode{wowaenthai}{00A0}
\pdfglyphtounicode{wparen}{00A0}
\pdfglyphtounicode{wreathproduct}{00A0}
\pdfglyphtounicode{wring}{00A0}
\pdfglyphtounicode{wsuperior}{00A0}
\pdfglyphtounicode{wturned}{00A0}
\pdfglyphtounicode{wynn}{00A0}
\pdfglyphtounicode{x}{00A0}
\pdfglyphtounicode{xabovecmb}{00A0}
\pdfglyphtounicode{xbopomofo}{00A0}
\pdfglyphtounicode{xcircle}{00A0}
\pdfglyphtounicode{xdieresis}{00A0}
\pdfglyphtounicode{xdotaccent}{00A0}
\pdfglyphtounicode{xeharmenian}{00A0}
\pdfglyphtounicode{xi}{00A0}
\pdfglyphtounicode{xmonospace}{00A0}
\pdfglyphtounicode{xparen}{00A0}
\pdfglyphtounicode{xsuperior}{00A0}
\pdfglyphtounicode{y}{00A0}
\pdfglyphtounicode{yaadosquare}{00A0}
\pdfglyphtounicode{yabengali}{00A0}
\pdfglyphtounicode{yacute}{00A0}
\pdfglyphtounicode{yadeva}{00A0}
\pdfglyphtounicode{yaekorean}{00A0}
\pdfglyphtounicode{yagujarati}{00A0}
\pdfglyphtounicode{yagurmukhi}{00A0}
\pdfglyphtounicode{yahiragana}{00A0}
\pdfglyphtounicode{yakatakana}{00A0}
\pdfglyphtounicode{yakatakanahalfwidth}{00A0}
\pdfglyphtounicode{yakorean}{00A0}
\pdfglyphtounicode{yamakkanthai}{00A0}
\pdfglyphtounicode{yasmallhiragana}{00A0}
\pdfglyphtounicode{yasmallkatakana}{00A0}
\pdfglyphtounicode{yasmallkatakanahalfwidth}{00A0}
\pdfglyphtounicode{yatcyrillic}{00A0}
\pdfglyphtounicode{ycircle}{00A0}
\pdfglyphtounicode{ycircumflex}{00A0}
\pdfglyphtounicode{ydieresis}{00A0}
\pdfglyphtounicode{ydotaccent}{00A0}
\pdfglyphtounicode{ydotbelow}{00A0}
\pdfglyphtounicode{yeharabic}{00A0}
\pdfglyphtounicode{yehbarreearabic}{00A0}
\pdfglyphtounicode{yehbarreefinalarabic}{00A0}
\pdfglyphtounicode{yehfinalarabic}{00A0}
\pdfglyphtounicode{yehhamzaabovearabic}{00A0}
\pdfglyphtounicode{yehhamzaabovefinalarabic}{00A0}
\pdfglyphtounicode{yehhamzaaboveinitialarabic}{00A0}
\pdfglyphtounicode{yehhamzaabovemedialarabic}{00A0}
\pdfglyphtounicode{yehinitialarabic}{00A0}
\pdfglyphtounicode{yehmedialarabic}{00A0}
\pdfglyphtounicode{yehmeeminitialarabic}{00A0}
\pdfglyphtounicode{yehmeemisolatedarabic}{00A0}
\pdfglyphtounicode{yehnoonfinalarabic}{00A0}
\pdfglyphtounicode{yehthreedotsbelowarabic}{00A0}
\pdfglyphtounicode{yekorean}{00A0}
\pdfglyphtounicode{yen}{00A0}
\pdfglyphtounicode{yenmonospace}{00A0}
\pdfglyphtounicode{yeokorean}{00A0}
\pdfglyphtounicode{yeorinhieuhkorean}{00A0}
\pdfglyphtounicode{yerahbenyomohebrew}{00A0}
\pdfglyphtounicode{yerahbenyomolefthebrew}{00A0}
\pdfglyphtounicode{yericyrillic}{00A0}
\pdfglyphtounicode{yerudieresiscyrillic}{00A0}
\pdfglyphtounicode{yesieungkorean}{00A0}
\pdfglyphtounicode{yesieungpansioskorean}{00A0}
\pdfglyphtounicode{yesieungsioskorean}{00A0}
\pdfglyphtounicode{yetivhebrew}{00A0}
\pdfglyphtounicode{ygrave}{00A0}
\pdfglyphtounicode{yhook}{00A0}
\pdfglyphtounicode{yhookabove}{00A0}
\pdfglyphtounicode{yiarmenian}{00A0}
\pdfglyphtounicode{yicyrillic}{00A0}
\pdfglyphtounicode{yikorean}{00A0}
\pdfglyphtounicode{yinyang}{00A0}
\pdfglyphtounicode{yiwnarmenian}{00A0}
\pdfglyphtounicode{ymonospace}{00A0}
\pdfglyphtounicode{yod}{00A0}
\pdfglyphtounicode{yoddagesh}{00A0}
\pdfglyphtounicode{yoddageshhebrew}{00A0}
\pdfglyphtounicode{yodhebrew}{00A0}
\pdfglyphtounicode{yodyodhebrew}{00A0}
\pdfglyphtounicode{yodyodpatahhebrew}{00A0}
\pdfglyphtounicode{yohiragana}{00A0}
\pdfglyphtounicode{yoikorean}{00A0}
\pdfglyphtounicode{yokatakana}{00A0}
\pdfglyphtounicode{yokatakanahalfwidth}{00A0}
\pdfglyphtounicode{yokorean}{00A0}
\pdfglyphtounicode{yosmallhiragana}{00A0}
\pdfglyphtounicode{yosmallkatakana}{00A0}
\pdfglyphtounicode{yosmallkatakanahalfwidth}{00A0}
\pdfglyphtounicode{yotgreek}{00A0}
\pdfglyphtounicode{yoyaekorean}{00A0}
\pdfglyphtounicode{yoyakorean}{00A0}
\pdfglyphtounicode{yoyakthai}{00A0}
\pdfglyphtounicode{yoyingthai}{00A0}
\pdfglyphtounicode{yparen}{00A0}
\pdfglyphtounicode{ypogegrammeni}{00A0}
\pdfglyphtounicode{ypogegrammenigreekcmb}{00A0}
\pdfglyphtounicode{yr}{00A0}
\pdfglyphtounicode{yring}{00A0}
\pdfglyphtounicode{ysuperior}{00A0}
\pdfglyphtounicode{ytilde}{00A0}
\pdfglyphtounicode{yturned}{00A0}
\pdfglyphtounicode{yuhiragana}{00A0}
\pdfglyphtounicode{yuikorean}{00A0}
\pdfglyphtounicode{yukatakana}{00A0}
\pdfglyphtounicode{yukatakanahalfwidth}{00A0}
\pdfglyphtounicode{yukorean}{00A0}
\pdfglyphtounicode{yusbigcyrillic}{00A0}
\pdfglyphtounicode{yusbigiotifiedcyrillic}{00A0}
\pdfglyphtounicode{yuslittlecyrillic}{00A0}
\pdfglyphtounicode{yuslittleiotifiedcyrillic}{00A0}
\pdfglyphtounicode{yusmallhiragana}{00A0}
\pdfglyphtounicode{yusmallkatakana}{00A0}
\pdfglyphtounicode{yusmallkatakanahalfwidth}{00A0}
\pdfglyphtounicode{yuyekorean}{00A0}
\pdfglyphtounicode{yuyeokorean}{00A0}
\pdfglyphtounicode{yyabengali}{00A0}
\pdfglyphtounicode{yyadeva}{00A0}
\pdfglyphtounicode{z}{00A0}
\pdfglyphtounicode{zaarmenian}{00A0}
\pdfglyphtounicode{zacute}{00A0}
\pdfglyphtounicode{zadeva}{00A0}
\pdfglyphtounicode{zagurmukhi}{00A0}
\pdfglyphtounicode{zaharabic}{00A0}
\pdfglyphtounicode{zahfinalarabic}{00A0}
\pdfglyphtounicode{zahinitialarabic}{00A0}
\pdfglyphtounicode{zahiragana}{00A0}
\pdfglyphtounicode{zahmedialarabic}{00A0}
\pdfglyphtounicode{zainarabic}{00A0}
\pdfglyphtounicode{zainfinalarabic}{00A0}
\pdfglyphtounicode{zakatakana}{00A0}
\pdfglyphtounicode{zaqefgadolhebrew}{00A0}
\pdfglyphtounicode{zaqefqatanhebrew}{00A0}
\pdfglyphtounicode{zarqahebrew}{00A0}
\pdfglyphtounicode{zayin}{00A0}
\pdfglyphtounicode{zayindagesh}{00A0}
\pdfglyphtounicode{zayindageshhebrew}{00A0}
\pdfglyphtounicode{zayinhebrew}{00A0}
\pdfglyphtounicode{zbopomofo}{00A0}
\pdfglyphtounicode{zcaron}{00A0}
\pdfglyphtounicode{zcircle}{00A0}
\pdfglyphtounicode{zcircumflex}{00A0}
\pdfglyphtounicode{zcurl}{00A0}
\pdfglyphtounicode{zdot}{00A0}
\pdfglyphtounicode{zdotaccent}{00A0}
\pdfglyphtounicode{zdotbelow}{00A0}
\pdfglyphtounicode{zecyrillic}{00A0}
\pdfglyphtounicode{zedescendercyrillic}{00A0}
\pdfglyphtounicode{zedieresiscyrillic}{00A0}
\pdfglyphtounicode{zehiragana}{00A0}
\pdfglyphtounicode{zekatakana}{00A0}
\pdfglyphtounicode{zero}{00A0}
\pdfglyphtounicode{zeroarabic}{00A0}
\pdfglyphtounicode{zerobengali}{00A0}
\pdfglyphtounicode{zerodeva}{00A0}
\pdfglyphtounicode{zerogujarati}{00A0}
\pdfglyphtounicode{zerogurmukhi}{00A0}
\pdfglyphtounicode{zerohackarabic}{00A0}
\pdfglyphtounicode{zeroinferior}{00A0}
\pdfglyphtounicode{zeromonospace}{00A0}
\pdfglyphtounicode{zerooldstyle}{00A0}
\pdfglyphtounicode{zeropersian}{00A0}
\pdfglyphtounicode{zerosuperior}{00A0}
\pdfglyphtounicode{zerothai}{00A0}
\pdfglyphtounicode{zerowidthjoiner}{00A0}
\pdfglyphtounicode{zerowidthnonjoiner}{00A0}
\pdfglyphtounicode{zerowidthspace}{00A0}
\pdfglyphtounicode{zeta}{00A0}
\pdfglyphtounicode{zhbopomofo}{00A0}
\pdfglyphtounicode{zhearmenian}{00A0}
\pdfglyphtounicode{zhebrevecyrillic}{00A0}
\pdfglyphtounicode{zhecyrillic}{00A0}
\pdfglyphtounicode{zhedescendercyrillic}{00A0}
\pdfglyphtounicode{zhedieresiscyrillic}{00A0}
\pdfglyphtounicode{zihiragana}{00A0}
\pdfglyphtounicode{zikatakana}{00A0}
\pdfglyphtounicode{zinorhebrew}{00A0}
\pdfglyphtounicode{zlinebelow}{00A0}
\pdfglyphtounicode{zmonospace}{00A0}
\pdfglyphtounicode{zohiragana}{00A0}
\pdfglyphtounicode{zokatakana}{00A0}
\pdfglyphtounicode{zparen}{00A0}
\pdfglyphtounicode{zretroflexhook}{00A0}
\pdfglyphtounicode{zstroke}{00A0}
\pdfglyphtounicode{zuhiragana}{00A0}
\pdfglyphtounicode{zukatakana}{00A0}
