## Supplementary material for "Advancing Early Diagnosis of Alzheimer’s Disease: A Paper-Based Aptasensor for Detecting A*β*(1-42) and p-tau181 from Plasma Using CdTe Quantum Dots": Latex Source: main.tex

\documentclass{article}
\usepackage{graphicx} % Required for inserting images

\title{Heart_Failure}
\author{Nihat Ahmadli}
\date{December 2023}

\begin{document}

\maketitle

\section{Introduction}

\end{document}
