## Supplementary material for "Advancing Early Diagnosis of Alzheimer’s Disease: A Paper-Based Aptasensor for Detecting A*β*(1-42) and p-tau181 from Plasma Using CdTe Quantum Dots": Latex Source: makefile

```
# $Id: makefile 37 2020-02-25 09:06:02Z rishi $

file=elsdoc-cas


all: pdf out 
	make pdf
	make pdf

out:
	if  [ -f $(file).out ] ; then cp $(file).out tmp.out; fi ;
	sed 's/BOOKMARK/dtxmark/g;' tmp.out > x.out; mv x.out tmp.out ;

pdf:
	pdflatex $(file).tex

index:
	makeindex -s gind.ist -o $(file).ind $(file).idx 

changes:
	makeindex -s gglo.ist -o $(file).gls $(file).glo

xview:
	xpdf -z 200 $(file).pdf &>/dev/null

view:
	open -a 'Adobe Reader.app' $(file).pdf

ins:
	latex $(file).ins 

diff:
	diff $(file).sty ../$(file).sty |less

copy:
	cp $(file).sty ../
```
