## Supplementary material for "Advancing Early Diagnosis of Alzheimer’s Disease: A Paper-Based Aptasensor for Detecting A*β*(1-42) and p-tau181 from Plasma Using CdTe Quantum Dots": Latex Source: manifest.txt

% Copyright 2019-2021 Elsevier Ltd
%
% This file is part of the 'CAS Bundle'.
% --------------------------------------
%
% It may be distributed and/or modified under the
% conditions of the LaTeX Project Public License, either version 1.2
% of this license or (at your option) any later version.
% The latest version of this license is in
%    http://www.latex-project.org/lppl.txt
% and version 1.2 or later is part of all distributions of LaTeX
% version 1999/12/01 or later.
%
% The list of all files belonging to the LaTeX 'CAS Bundle' is
% given in the file `manifest.txt'.
%
% CONTENTS OF THE CAS BUNDLE
% ==========================

Directory elsevier-cas-template/

cas-sc.cls
  Classfile to be used for single column format

cas-dc.cls
  Classfile to be used for double column format

cas-model2-names.bst
  BibTeX style file

cas-sc-template.tex 
  TeX template 
  
cas-dc-template.tex
  TeX template 

manifest.txt
  this file

README
  small readme documentation

Directory doc/

The following files are graphic files needed for creating pdf output
of the documentation from elsdoc.tex:

 dc-sample.pdf
 sc-sample.pdf

elsdoc-cas.tex  -- LaTeX source file of documentation
elsdoc-cas.pdf  -- documentation for elsarticle.cls

Directory thumbnails/

Contains thumbnail images which will be included in the
typeset PDF.

 cas-email.jpeg
 cas-facebook.jpeg
 cas-gplus.jpeg
 cas-linkedin.jpeg
 cas-twitter.jpeg
 cas-url.jpeg

The following files are written out every time elsdoc.tex is compiled:

 elsdoc-cas.aux
 elsdoc-cas.log
 elsdoc-cas.out
 tmp-cas.tex

Auxiliary packages needed to generate pdf output from elsdoc.tex:

 rvdtx.sty
 pdfwidgets.sty
 glyphtounicode.tex
