## Supplementary material for "Advancing Early Diagnosis of Alzheimer’s Disease: A Paper-Based Aptasensor for Detecting A*β*(1-42) and p-tau181 from Plasma Using CdTe Quantum Dots": Latex Source: README

This bundle provides two classfiles, namely cas-sc.cls and cas-dc.cls
and corresponding template files for typesetting journal articles
supposed to go through Elsevier's updated workflow. cas-sc.cls is meant
for one-column, the other cas-dc.cls for two-column layout. These are
now accepted for submitting articles both in Elsevier's electronic
submission system and elsewhere.

Bugs and problems with classfiles may be reported to the
developers of the class via.

The file manifest.txt provides a list of the files in the
elsarticle-cas bundle.  The following are the main files available:

- cas-sc.cls, classfile to typeset article in single column
- cas-dc.cls, classfile to typeset article in double column
- cas-sc-template.tex, template file for single column journals
- cas-dc-template.tex, template file for double column journals
- cas-common.sty, additional macro package for formatting
- cas-model2-names.bst, BibTeX style file
- cas-refs.bib, sample BibTeX database
- cas-twitter.jpeg, cas-linkedin.jpeg, cas-url.jpeg, cas-gplus.jpeg, 
  cas-facebook.jpeg, cas-email.jpeg - thumbnail images of social 
  networking sites

The PDF documentation elsdoc-cas.pdf is in the doc directory.  To
generate it:

 1. pdflatex elsdoc-cas
 2. pdflatex elsdoc-cas
 3. pdflatex elsdoc-cas

or

use the makefile. Use the target `all' (eg: make all) in doc
directory.

The above procedure will create a print version, namely elsdoc-cas.pdf.

Copyright 2019-2021, Elsevier. Bugs, feature requests, suggestions and
comments may be mailed to. cas-sc.cls, cas-dc.cls
related documentations and supporting packages are released under the
LATEX Project Public Licence, either version 1.2 or any later
version. This work has the LPPL maintenance status
'author-maintained'.
