## Supplementary figures and images for "Advancing Early Diagnosis of Alzheimer’s Disease: A Paper-Based Aptasensor for Detecting A*β*(1-42) and p-tau181 from Plasma Using CdTe Quantum Dots"

### absorp_emissi_qd.png

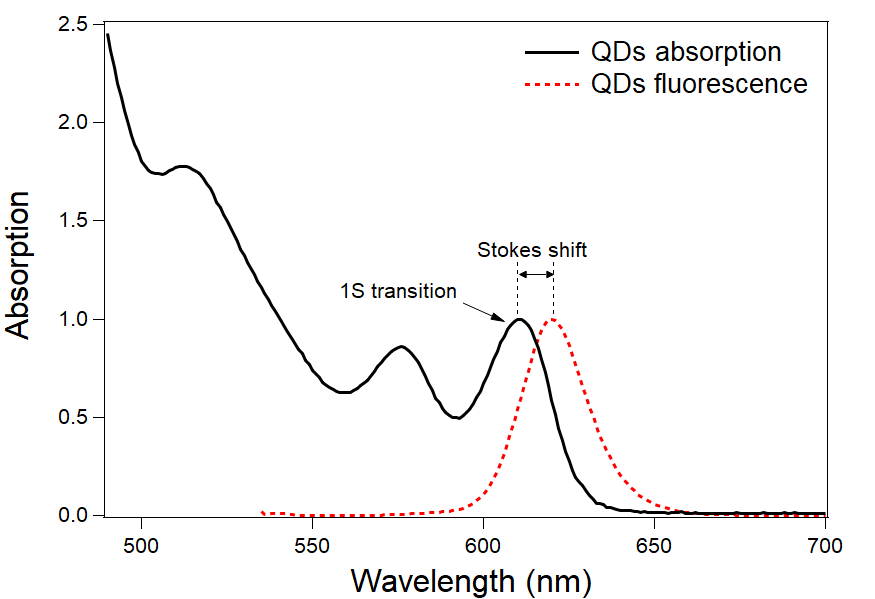

### complex.png

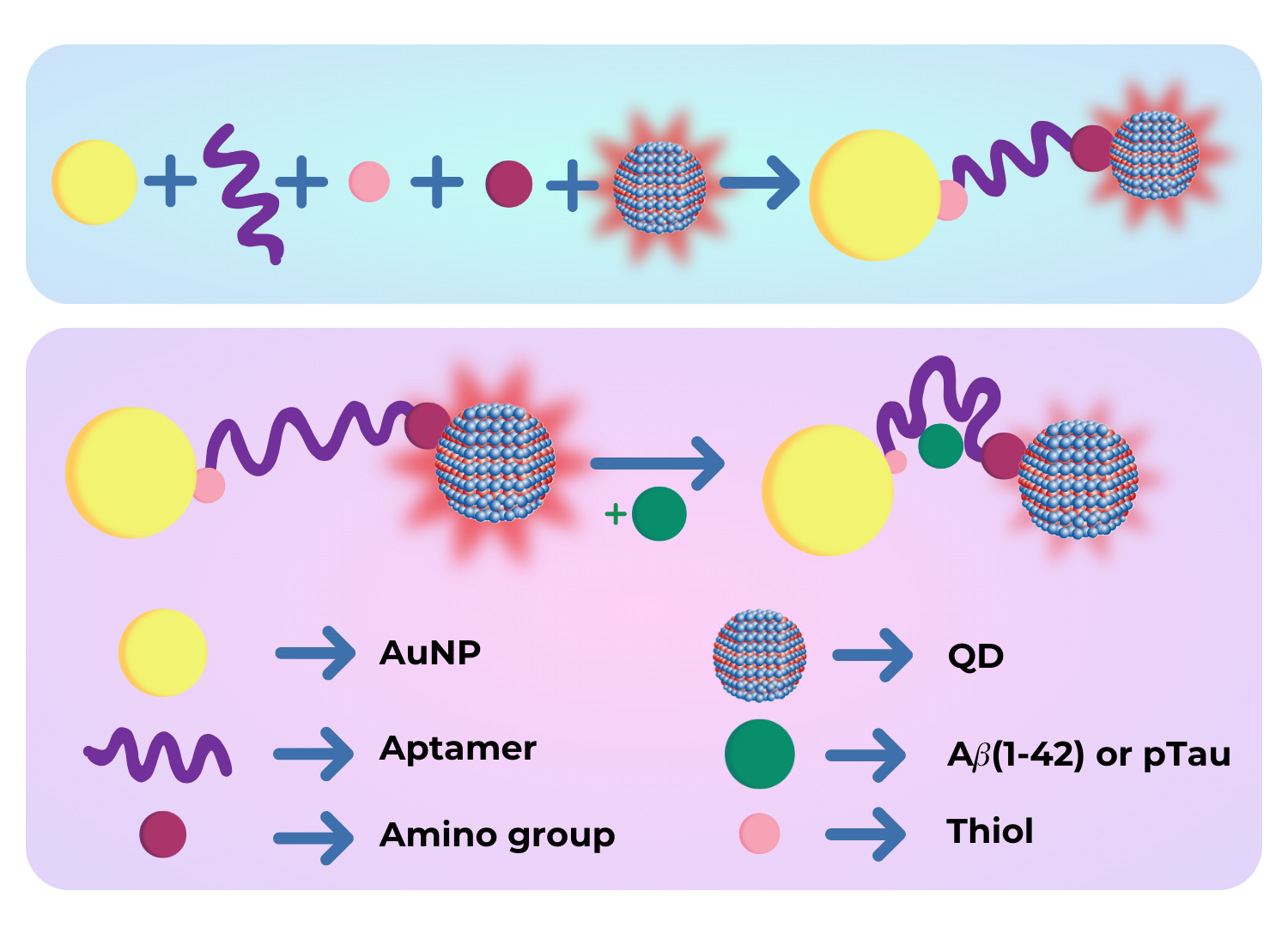

### desn.png

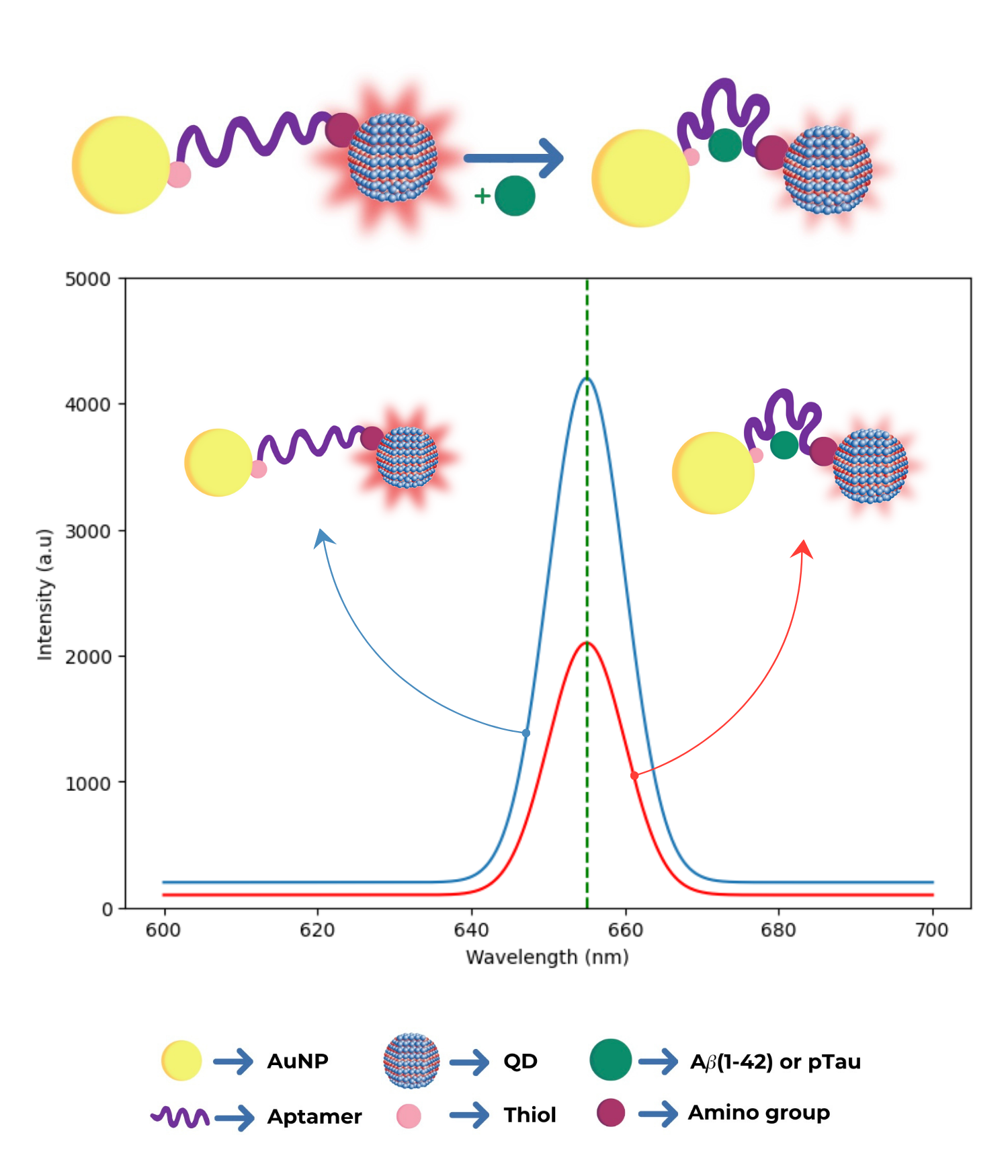

### emission.png

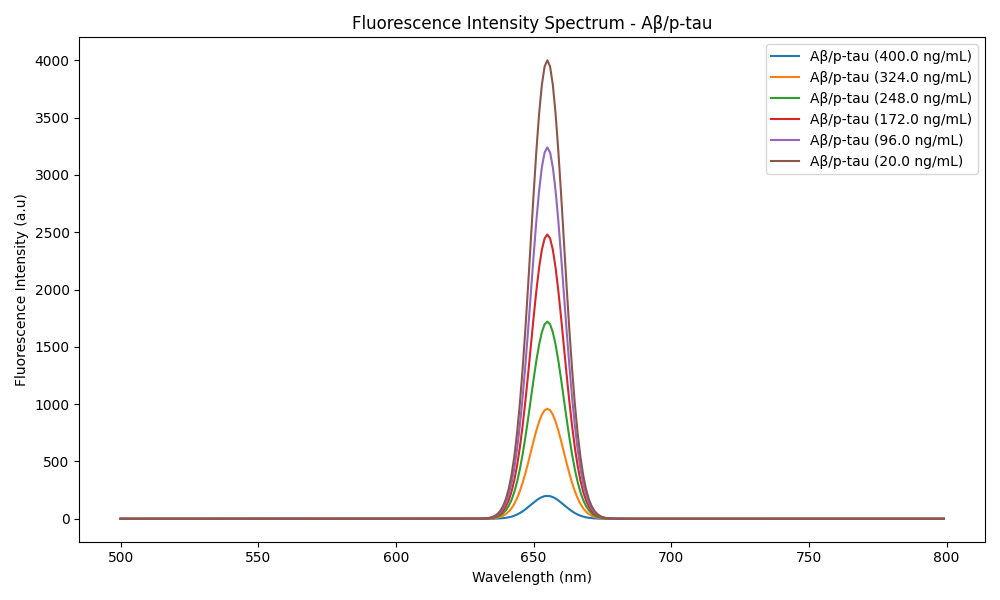

### meas1.png

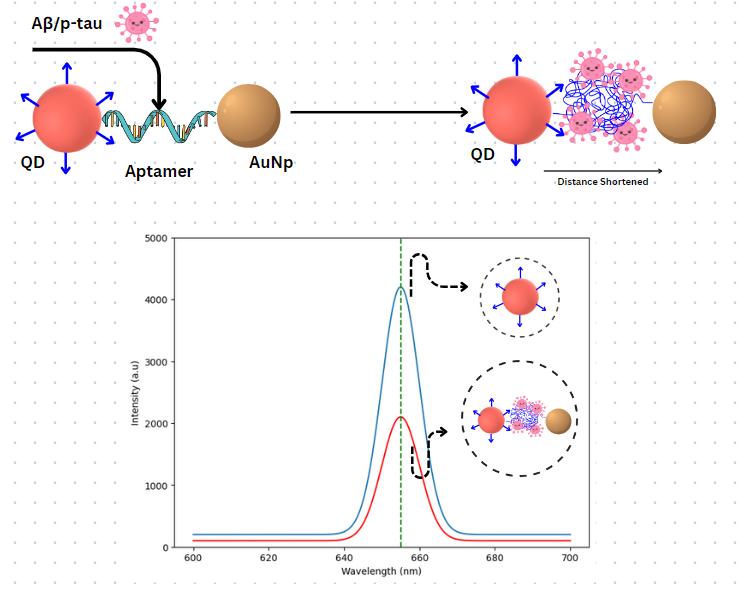

### paper.png

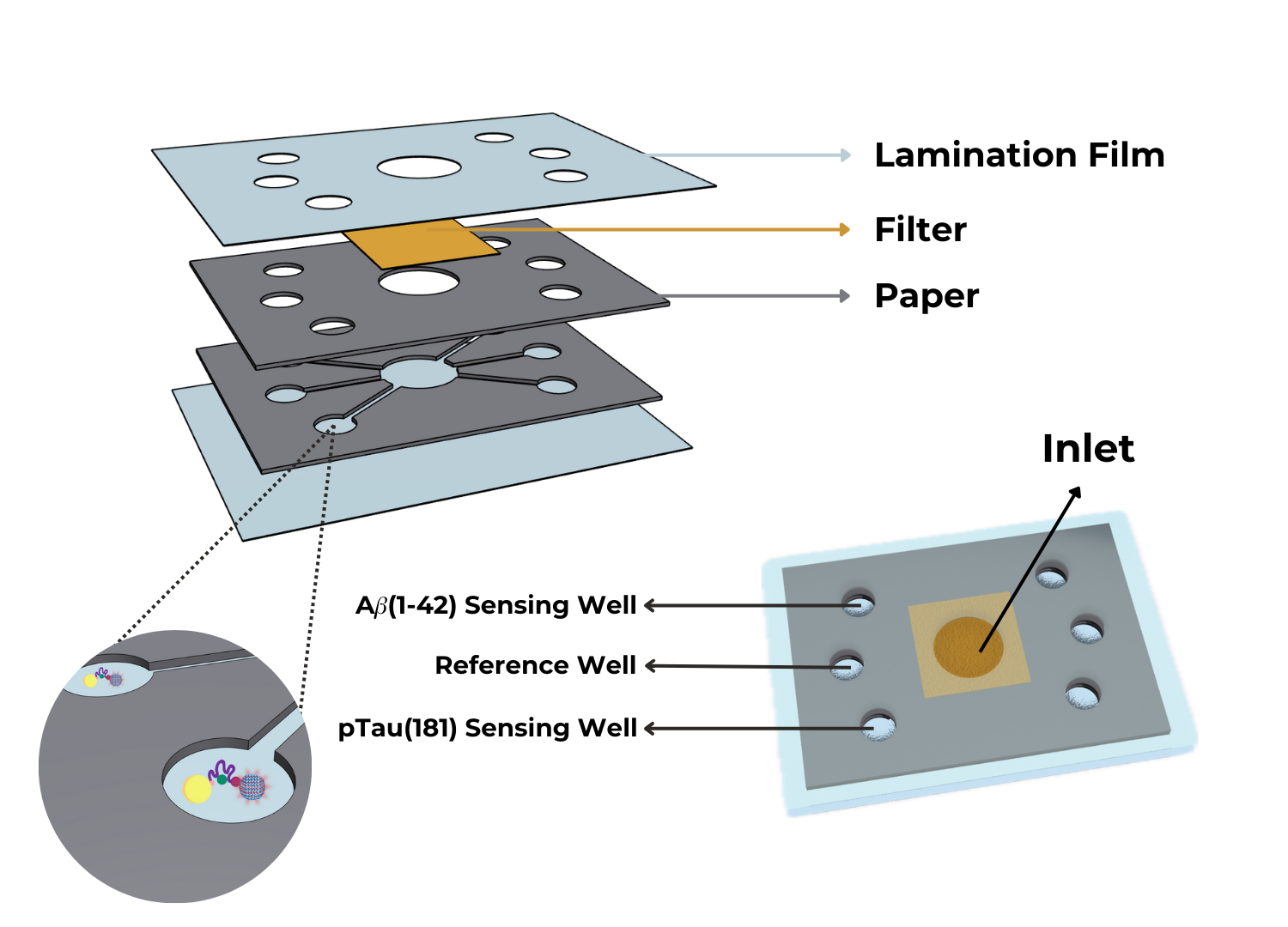

### qd.jpeg

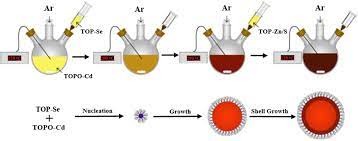

### sample.png

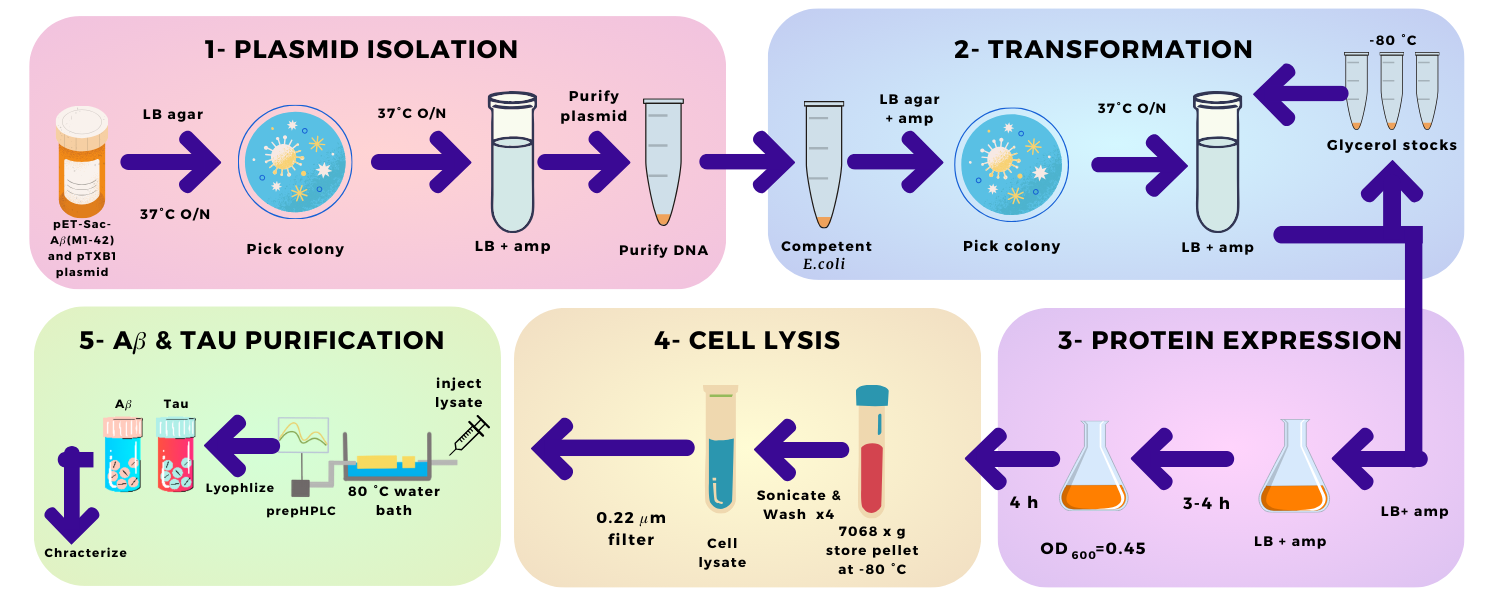

### tool.png

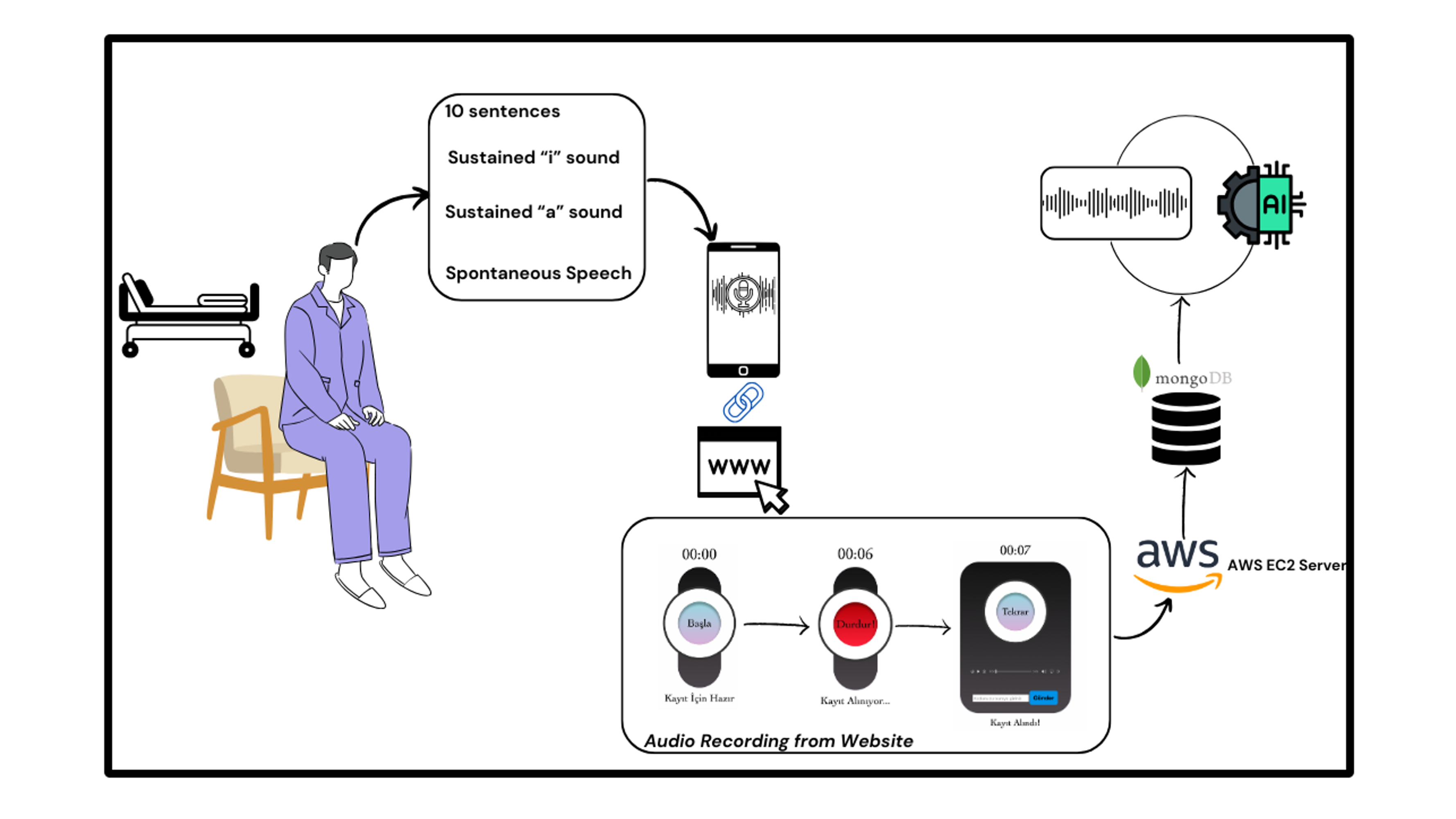
